# Cell line identity rather than medium composition determines transcriptomic profiles of HepaRG and HuH7 cells cultured in chemically defined or serum-based media: comparison with primary human hepatocytes

**DOI:** 10.64898/2026.03.09.710463

**Authors:** Ahmed S. M. Ali, Heike Sprenger, Albert Braeuning, Jens Kurreck

**Affiliations:** Technische Universität Berlin, Chair of Applied Biochemistry, Berlin, Germany; German Federal Institute for Risk Assessment (BfR), Berlin, Department Chemical and Product Safety, Germany

**Keywords:** chemically defined medium (CDM), serum-free medium, HepaRG, HuH7, primary human hepatocytes (PHH), RNA-seq, transcriptomic analysis, liver

## Abstract

The composition of culture medium is a major, yet frequently undercontrolled, determinant of hepatic cell state *in vitro*. For decades, fetal bovine serum (FBS) has been routinely incorporated into liver cell culture. Its undefined and lot-to-lot variable composition can, however, confound cell identity and experimental reproducibility. Serum-free, chemically defined media (CDM) represent an alternative approach that can improve standardization, but the consequences of transitioning from FBS-supplemented media (FBS-SM) to CDM remain insufficiently characterized in hepatic models, particularly with respect to metabolic and detoxification programs that govern xenobiotic handling and hepatotoxicity readouts. Here, we systematically assessed how replacing FBS-SM with CDM remodels transcriptomic profiles in two widely used human hepatic cell lines (HepaRG and HuH7 cells) and compared the results to that obtained from primary human hepatocytes (PHH). Global transcriptomic analyses indicated that cell type was the primary driver of variance, whereas medium induced a model-dependent secondary effect. Functional interpretation showed preferential enhancement of xenobiotic metabolism and transport-associated programs in HepaRG cells, while HuH7 cells response was dominated by lipid/sterol homeostasis and stress-linked processes. Benchmarking against PHH based on hepatic identity and detoxification gene panels further supported improved PHH alignment for HepaRG cells under CDM compared to cultures with FBS-SM, with limited improvement for HuH7 cells. Collectively, these findings address a key knowledge gap by defining how FBS-SM and CDM impact the transcriptomic profiles of HepaRG and HuH7 cells.

## Introduction

Mammalian cell culture is foundational to biomedical research, yet the field still relies heavily on classical media supplemented with FBS [1]. Since its introduction in the late 1950s, FBS has been widely adopted as a practical “gold standard” because it provides a complex mixture of growth factors, hormones, carrier proteins, and micronutrients that supports attachment and proliferation across diverse cell types [2].

Despite its widespread use, FBS introduces well-recognized limitations that can compromise reproducibility and translational interpretation [3]. Its composition is inherently ill-defined and varies substantially between lots, which can drive batch-dependent differences and weaken reproducibility [4]. In addition, serum can contain endotoxins [5], hemoglobin [6], and other inhibitory factors [4], and it is a recognized potential source of microbial contamination, including mycoplasma [7], pestiviruses [7, 8], fungi [6], and prions [9]. Beyond technical concerns, the xenogeneic origin of FBS and the ethical implications of its sourcing increasingly conflict with the goals of human-relevant and animal-free research practices.

Serum-free and chemically defined media (CDM) have therefore gained momentum as rational alternatives designed to replace the “black box” of serum with controlled and interpretable culture conditions [10]. In principle, CDM reduces qualitative and quantitative variability in medium composition by introducing components like insulin, transferrin, selenium, as well as essential nutrients in specific concentrations [11]. However, replacing FBS is not as simple as switching to commercial “serum-free” products. Many alternatives are compositionally undisclosed, preserving key interpretability limitations and introducing risks of undocumented formulation changes that undermine reproducibility. Recently, Rafnsdóttir et al. reported an animal product-free, chemically defined “universal” formulation (OUR medium) that sustained growth of multiple cell lines [12]. A subsequent paper by the authors provided a stepwise preparation protocol and practical guidance for mixing and using the medium [13]. However, the relatively high cost of universal media can limit the practical use in large-scale studies and high-throughput screening, motivating continued development of disclosed, fit-for-purpose CDM formulations.

Decades of serum-based work have raised the possibility that legacy datasets partly reflect serum-conditioned biology rather than physiology. Consequently, replacing FBS-SM with CDM can generate concern that future results will not be directly comparable with historical literature. These considerations underscore the need for systematic benchmarking of how CDM affects cellular programs relative to traditional serum-based conditions.

These issues are particularly salient in liver research, where *in vitro* systems are expected to represent xenobiotic handling, central to pharmacology and hepatotoxicity assessment. Primary human hepatocytes (PHH) are widely regarded as the gold standard for *in vitro* studies of hepatic metabolism and toxicity. However, their use is constrained by limited availability and pronounced donor-to-donor variability [14]. These limitations have driven the widespread adoption of human liver-derived cell lines as more practical alternatives for mechanistic studies and high-throughput screening applications [15–17]. Among these models, HepaRG and HuH7 cells remain extensively used due to their robustness and experimental accessibility. HepaRG cells, in particular, exhibit a comparatively differentiated hepatic phenotype, including the expression of drug-metabolizing enzymes and transporters [15], whereas HuH7 cells provide a genetically stable platform frequently employed in toxicological and viral infection research [18].

In this study, we performed a comprehensive transcriptomic analysis to systematically evaluate how replacing FBS-SM with CDM influences global and functional gene expression profiles in widely used hepatic cell lines (HepaRG and HuH7). RNA sequencing (RNA-seq), followed by differential expression analysis, pathway enrichment, and gene set variation analysis, was employed to evaluate both culture-dependent and cell type-dependent changes in hepatic transcriptional programs. The results were compared to transcriptomic analyses of PHH.

## Materials and Methods

### Cell culture

HepaRG and HuH7 cells were used as representative human hepatic *in vitro* models and were maintained under two parallel conditions: a conventional serum-supplemented medium and a serum-free CDM, with cell line-specific formulations for HepaRG and HuH7 cells. Serum-supplemented conditions were selected based on established literature protocols [19, 20], whereas the CDM condition was based on custom formulations previously reported to support both cell lines [21, 22].

HepaRG cells (Biopredic International) were cultured in William’s E Medium (12551-032, Gibco) as the basal medium under both serum-supplemented and chemically defined conditions. Unless otherwise specified, all HepaRG cell cultures were supplemented with L-glutamine (2 mM; X0550, Biowest), insulin (5 µg/mL; P07-04300, PAN-Biotech), hydrocortisone 21-hemisuccinate sodium salt (50 µM; H2270-100MG, Sigma-Aldrich), and penicillin/streptomycin (P/S; 1×; L0022, Biowest). For serum-supplemented culture, the basal medium was additionally supplemented with FBS (10%; fetal calf serum, FCS; S-14-L, c.c.pro GmbH). For serum-free culture, serum was replaced with a defined supplement comprising insulin-transferrin-selenium (ITS; 1×; 41400045, Gibco), sodium pyruvate (1 mM; S8636-100ML, Sigma-Aldrich), 2-mercaptoethanol (55 µM; 21985-023, Gibco), and recombinant human fibronectin (1.5 µg/mL; 1918-FN, R&D Systems). HepaRG cell differentiation toward a hepatocyte-like phenotype was induced using an identical protocol under both culture conditions while maintaining the same base formulation used for culture (serum-supplemented medium containing 10% FBS for the FBS-SM condition; serum-free CDM for the CDM condition): 14 days in growth medium, followed by 24 h in the corresponding medium supplemented with dimethyl sulfoxide (DMSO; 1%; D2650, Sigma-Aldrich), and a subsequent 14-day differentiation period in medium containing 1.7% DMSO prior to RNA isolation.

HuH7 cells (330156, CLS Cell Lines Service GmbH) were maintained under parallel serum-supplemented and chemically defined conditions. For serum-supplemented culture, cells were maintained in DMEM low glucose (L0064, Biowest) supplemented with FBS (10%; c.c.pro GmbH), L-glutamine (2 mM; Biowest), and P/S (1×; Biowest). D-(+)-glucose (G8769, Sigma-Aldrich) was added to a final concentration of 4.5 g/L. For serum-free culture, cells were maintained in a CDM based on DMEM-F12 (L0090, Biowest) supplemented with CD lipid concentrate (1:100; 11905-031, Gibco), MEM non-essential amino acids (NEAA; 1×; X0557-100, Biowest), HEPES (10 mM; L0180, Biowest), Pluronic F-68™ (0.1%; A1288, Applichem), GlutaMAX™ (2 mM; 35050, Gibco), D-(+)-glucose (4.5 g/L; Sigma-Aldrich), P/S (1×; Biowest), ITS (1×; Gibco), recombinant human epidermal growth factor (EGF; 10 ng/mL; PHG0313, Gibco), and recombinant human hepatocyte growth factor (HGF; 10 ng/mL; PHG0254, Gibco). For serum-free culture, HuH7 cells were seeded on culture vessels coated with human placental collagen type I (THT0102, THT Biomaterials GmbH) at a final coating concentration of 32 µg/mL, as previously reported [21].

HeLa cells (ACC 57, DSMZ, Braunschweig) were included as a non-hepatic comparator and were cultured according to previously reported conditions (Supporting Methods SM1) [23]. For each cell line, cultures maintained in FBS-SM and CDM were passaged in parallel for five consecutive passages prior to RNA isolation. Unless otherwise stated, all cultures were maintained at 37 °C in a humidified atmosphere containing 5% CO₂.

Cells were directly adapted to CDM without gradual serum withdrawal and were subcultured using TrypLE™ Express (12604021, Gibco). Pooled PHH were included as a physiological reference and were obtained commercially (HEP10™ Pooled Human Hepatocytes, HMCS10, Thermo Fisher Scientific), comprising hepatocytes pooled from ten individual donors (five male and five female). Cryopreserved PHH were processed directly for RNA isolation without *in vitro* culture.

### RNA isolation

Total RNA was isolated from HepaRG, HuH7, HeLa cells, and pooled PHH using the QIAwave RNA Kit (50) (74534, Qiagen) according to the manufacturer’s instructions. For HuH7 cells, RNA extraction was performed during the mid-exponential growth phase, whereas HepaRG cells were harvested following completion of the differentiation protocol. For Hep10™ pooled PHH, the cryovial was rapidly thawed in a 37 °C water bath for 2 min and immediately diluted in nine volumes of ice-cold phosphate-buffered saline (PBS; 13.5 mL) to minimize DMSO-associated cytotoxicity. Cells were pelleted by centrifugation at 100 × g for 5 min at 4 °C, after which the supernatant was carefully removed.

For HepaRG and HuH7 cells, monolayers were washed with PBS to remove residual culture medium and lysed directly in RLT buffer supplemented with β-mercaptoethanol (1% v/v), as recommended by the manufacturer. For PHH, cell pellets were lysed directly after removal of supernatant. RNA purification was then completed according to the manufacturer’s protocol. RNA concentration and purity (260/280 and 260/230 ratios) were assessed using a NanoDrop 2000 UV-Vis spectrophotometer (Thermo Scientific).

HeLa cells were processed for RNA isolation as shown in Supporting Methods SM1.

### RNA-seq and primary processing

RNA-seq was performed by Eurofins Genomics (INVIEW™ Transcriptome Discover service). Total RNA samples were processed using bead-based poly(A) selection for mRNA enrichment followed by library preparation. Libraries were sequenced using Illumina paired-end sequencing (2 × 150 bp) with a target yield of 30 million read pairs per sample (±3%). Raw reads were screened for rRNA-derived sequences using RiboDetector (v0.2.7), and reads classified as rRNA were removed prior to downstream processing. Quality filtering and adapter trimming of non-rRNA reads were performed using fastp (v0.20.0), applying sliding-window trimming with removal of bases below Phred Q20. High-quality reads were aligned to the human reference genome (UCSC hg38) using STAR (v2.7.10b) within the Sentieon (release 202503) secondary analysis framework using two-pass mapping and the parameterization reported by Eurofins (including splice junction and fusion read detection). Gene- and transcript-level abundance estimates were generated using RSEM (v1.3.3).

### Differential gene expression analysis

After removing genes with low expression (sum of reads across all samples below ten), differential gene expression analysis was performed in R using DESeq2 (v1.38.3) on gene-level integer count matrices for the retained 60,568 genes. DESeq2 size factors were estimated using the median-of-ratios method, followed by gene-wise dispersion estimation and model fitting under a negative binomial generalized linear model. Predefined contrasts captured medium effects within each model (CDM versus FBS-SM), cell type-specific comparisons, and transcriptomic benchmarking relative to PHH. False discovery rate (FDR) was used to control multiple testing [24]. For exploratory analyses and visualization, variance-stabilizing transformation (VST; DESeq2) was applied after expression filtering.

Global transcriptomic structure was assessed by principal component analysis (PCA) on VST-transformed expression values using the top 1,000 most variable genes ranked by row-wise variance (matrixStats v1.3.0) and visualized using ggplot2 (v3.5.1) and ggrepel (v0.9.5). In addition to sample-level distances, condition-mean VST profiles were computed to generate condition-to-condition distance matrices. Heatmaps were rendered using pheatmap (v1.0.13) and ComplexHeatmap (v2.14.0) with circlize (v0.4.16).

### Functional enrichment analyses

Functional enrichment analyses were performed on differential expression results using Gene Ontology Biological Process (GO:BP) over-representation testing implemented in clusterProfiler (v4.6.2). For each contrast, differentially expressed genes (DEGs) were stratified into three predefined sets: Upregulated (FDR < 0.05 and log2FC > 0.5), Downregulated (FDR < 0.05 and log2FC < −0.5), and Significant (FDR < 0.05 irrespective of direction). Enrichment testing was performed against the GO:BP ontology (GO.db v3.16.0), using Benjamini-Hochberg correction. To reduce semantic redundancy, results were simplified using Wang semantic similarity (GOSemSim v2.24.0) with a similarity cutoff of 0.7 and retention of terms with the lowest adjusted P value. Enrichment results were visualized using dot plots generated with enrichplot (v1.18.4) and ggplot2.

### Pathway-level analyses

Pathway activity was quantified using gene set variation analysis (GSVA) and pathway level analysis of gene expression (PLAGE) implemented in GSVA (v1.46.0) using DESeq2-derived VST expression values. Scores were computed at the per-sample level, with condition-level summaries generated downstream for visualization and benchmarking. Canonical pathway gene sets were obtained from MSigDB C2 (CP; Reactome, KEGG, and BioCarta subcollections) using msigdbr (v25.1.1). Prior to scoring, gene sets were restricted to genes present in the expression matrix.

For statistical inference on pathway scores, a condition-level design matrix was specified in limma (v3.54.2), with conditions defined as HepaRG_CDM, HepaRG_FBS-SM, HuH7_CDM, HuH7_FBS-SM, and PHH. To evaluate pathways reflecting hepatocyte functions and canonical injury mechanisms, including bile acid metabolism/transport (cholestatic liability) [25, 26], xenobiotic metabolism/transport and glutathione-dependent detoxification [27, 28], oxidative and endoplasmic reticulum (ER) stress responses [29], mitochondrial respiration/dysfunction [30], lipid/steroid/cholesterol homeostasis [31, 32], organic anion/acid transport [33], inflammatory signaling [34], and DNA damage responses [35], GO terms were curated a priori and queried in QuickGO (EMBL-EBI), with descendant terms included [25,26]. GO term-to-gene mappings were retrieved and standardized using AnnotationDbi (v1.70.0), org.Hs.eg.db (v3.16.0), and GO.db (v3.16.0). ssGSEA scores were computed per sample.

### PHH similarity analyses and identity marker panels

Three predefined detoxification gene panels were analyzed (Table S1): Phase I enzymes (44 genes), Phase II conjugation enzymes (35 genes), and transporters (23 genes) [36]. Condition-level heatmaps were generated using ComplexHeatmap with circlize and RColorBrewer (v1.1-3). Transcriptional similarity to pooled PHH was computed separately for each detoxification panel by calculating the root mean square error (RMSE) between each *in vitro* condition and the PHH reference vector across mapped genes, followed by normalization to a 0-1 scale as 1 − RMSE/max(RMSE) (PHH = 1 by definition). In addition, hepatocyte identity was assessed using predefined identity marker panels (22 genes, Table S2) [37–40]. Condition-mean VST expression values were extracted, mapped to gene symbols using org.Hs.eg.db, and summarized to compute PHH similarity scores using the same RMSE-based framework.

### Statistics and reproducibility

For RNA-seq analyses, HepaRG, HuH7 and HeLa cell culture conditions comprised three independent biological replicates per medium (n = 3), whereas PHH were included as technical duplicates (n = 2) of pooled cells from 5 female and 5 male donors, as stated by the supplier. Unless stated otherwise, statistical significance was defined as an adjusted P value (false discovery rate, FDR) < 0.05. Exploratory analyses, including PCA, sample- and condition-level distance assessments, and heatmap visualizations, were performed on VST expression values for descriptive and comparative purposes; no statistical inference was derived from these exploratory visualizations.

## Results

### RNA-seq quality control and alignment metrics

RNA-seq libraries generated by Eurofins Genomics exhibited consistently high quality across all HuH7, HepaRG and HeLa cell culture conditions, and pooled PHH replicates. Following rRNA screening and read cleaning, 63.05-64.85 million high-quality reads per sample were retained, corresponding to 95.53-98.25% read retention after filtering. Base-call quality was uniformly high (Q30: 92.24-96.93%), GC content was stable across libraries (48.60-49.88%), and the fraction of rRNA reads remained low overall (0.54-3.35%) (Table S3).

Alignment to the human reference genome using STAR achieved high mapping performance across all samples, with 97.19-99.24% of high-quality reads mapped and 92.83-97.50% uniquely mapped (Table S4). As expected, PHH libraries showed modestly lower mapping and unique alignment relative to the cell lines (mapped: 97.19-97.67%; unique: 92.83-93.06%), whereas HuH7 and HepaRG samples consistently exceeded 99% mapping with high unique alignment (HuH7 mapped: 99.05–99.24%, unique: 97.26–97.50%; HepaRG mapped: 99.02–99.16%, unique: 96.70–96.94%). HeLa libraries also showed high alignment performance (mapped: 98.11–99.13%; unique: 95.90–97.08%). Consistent with transcriptome-scale RNA-seq of intact mRNA, mapped reads were predominantly exonic (87.43–92.38%), with smaller intronic (6.56–11.42%) and intergenic fractions (1.01–1.93%), supporting robust gene-level quantification (Table S5).

### Global transcriptomic relationships across hepatic models and media

To evaluate global transcriptomic relationships independent of predefined contrasts, variance-stabilized expression profiles were examined using unsupervised clustering at both the condition and sample levels. Condition-level distances segregated samples primarily by cell model, with HepaRG cells clustering closer to pooled PHH than HuH7 cells, and both cell lines remaining distinct from PHH. Within each cell line, CDM and FBS-SM conditions were substantially more similar to one another than to the alternate cell line or PHH, indicating that lineage- and model-defining programs account for most global variance, while medium composition introduces a minimal displacement. HeLa cells, included as a non-hepatic outgroup, were clearly separated from HepaRG, HuH7 cells, and PHH in both the distance-based clustering and PCA, providing a non-hepatic reference for the transcriptomic space (Figure 1a, b).

**Figure 1.**
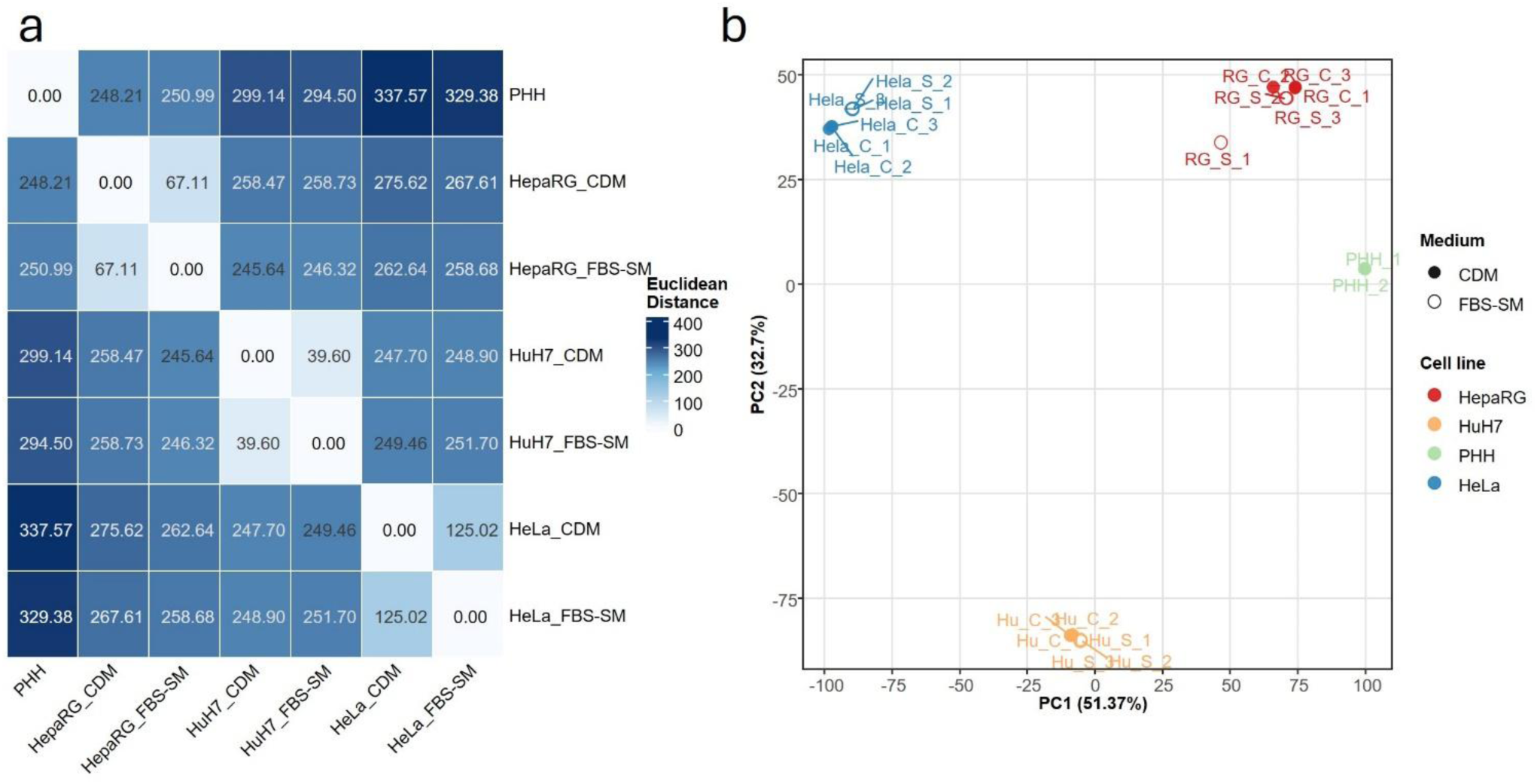
Global transcriptomic relationships across hepatic models, media, and reference samples. (a) Condition-level Euclidean distance heatmap computed from DESeq2 variance-stabilized expression values (VST), including HepaRG, HuH7 and Hela cells cultured in CDM or FBS-SM, pooled primary human hepatocytes (PHH), and HeLa cells as a non-hepatic outgroup. Distances were calculated on condition-mean VST profiles and clustered hierarchically. (b) Principal component analysis (PCA) of VST expression profiles across all samples (HepaRG, HuH7 and HeLa cells, n = 3 per medium; PHH, n = 2).

This structure was reinforced at the replicate level, where biological replicates clustered tightly within their respective groups, supporting high within-condition reproducibility and indicating that the observed separations reflect consistent biological differences rather than outlier-driven effects (Figure S1). These unsupervised analyses establish that medium composition modulates transcriptional state within a given hepatic model, but the dominant determinant of global transcriptomic structure remains cell identity.

### Medium-associated differential expression in HepaRG and HuH7 cells

To quantify medium-dependent transcriptional remodeling within each hepatic cell line, differential expression was assessed for CDM versus FBS-SM using a consistent statistical framework (adjusted P value < 0.05; |log2FC| > 0.5, corresponding to |fold change| > 1.41). Both HepaRG and HuH7 cells exhibited clear medium-responsive transcriptional changes, but the magnitude of the response was strongly model dependent (Figure 2a, b; Table 1). In HepaRG cells, CDM culture resulted in 2272 DEGs relative to FBS-SM, comprising 1104 upregulated (48.6%) and 1168 downregulated (51.4%) genes. In HuH7 cells, the medium effect was smaller, with 1017 DEGs (614 upregulated, 60.4%; 403 downregulated, 39.6%). Overlap of up- and downregulated DEGs across the medium contrasts is summarized using Venn diagram (Figure S2).

**Figure 2.**
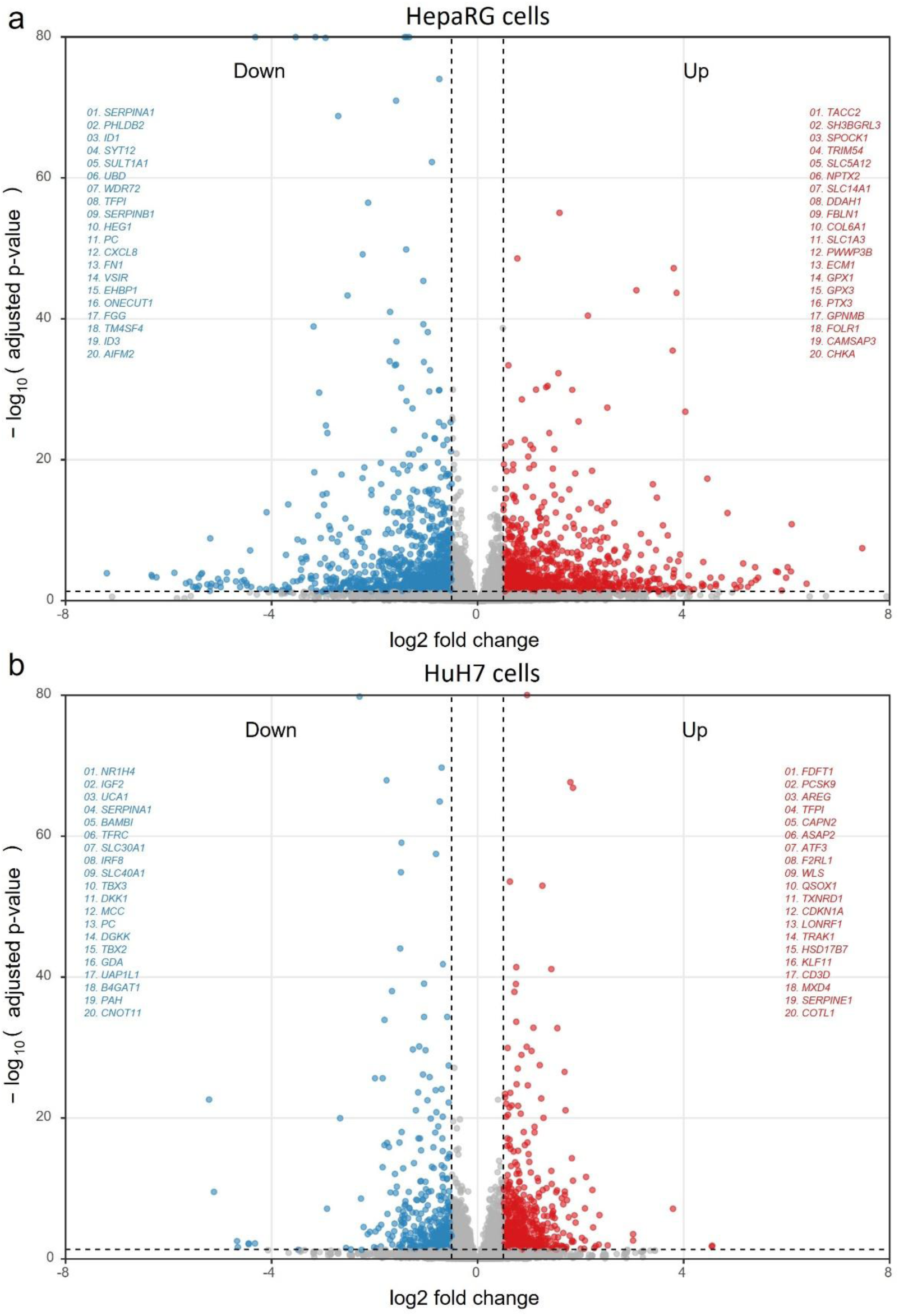
Medium-associated transcriptional responses in hepatic cell models. Volcano plots of DESeq2 results for CDM versus FBS-SM within (a) HepaRG and (b) HuH7 cells. Each point represents one gene; the x-axis denotes log2 fold change (CDM/ FBS-SM) and the y-axis denotes −log10 adjusted P value (Benjamini-Hochberg). Dashed lines indicate the significance thresholds used to define DEGs (FDR < 0.05 and |log2FC| > 0.5). Label strips report the top 20 upregulated and top 20 downregulated DEGs (ranked by adjusted P value; log2 fold change used as a secondary ordering criterion).

**Table 1.** Summary of differential gene expression across primary contrasts. Numbers of significantly upregulated (n_up) and downregulated (n_down) genes and total significant genes (n_sig) for each DESeq2 contrast, using a threshold of adjusted P value (Benjamini-Hochberg) < 0.05 and |log2 fold change| > 0.5. Full gene-level DESeq2 results are given in Table S6.

| contrast | N<sub>up</sub> | N<sub>down</sub> | N<sub>sig</sub> |
| --- | --- | --- | --- |
| HepaRG: CDM vs FBS-SM | 1104 | 1168 | 2272 |
| HuH7: CDM vs FBS-SM | 614 | 403 | 1017 |
| HepaRG CDM vs PHH | 6758 | 5962 | 12720 |
| HuH7 CDM vs PHH | 6536 | 7294 | 13830 |
| HepaRG vs HuH7 in CDM | 7927 | 6155 | 14082 |
| HuH7 in FBS-SM vs PHH | 6473 | 7150 | 13623 |
| HepaRG in FBS-SM vs PHH | 6681 | 5965 | 12646 |

Volcano plots showed that significantly changed genes were distributed across a broad range of fold changes and statistical support, with substantial numbers exceeding the fold-change threshold in both directions (Figure 2a, b). This bidirectional pattern indicates coordinated induction and repression of medium-sensitive programs rather than a uniform shift in expression. The label-strip annotations further highlight that medium-responsive transcripts include highly significant and relatively large-effect genes (Figure 2a, b), supporting the conclusion that CDM does not simply preserve an FBS-SM-like state, but instead drives a distinct transcriptional configuration. Full differential expression outputs for each contrast (all genes and significant subsets) are reported in Table S6.

Importantly, the scale of the CDM versus FBS-SM response remained moderate when contextualized against broader biological contrasts in the same dataset: comparisons to PHH and cross-cell line contrasts yielded ∼12,600-14,100 DEGs, far exceeding the medium-driven DEG burden in either cell line (Table 1; Table S6), Full-size volcano plots for the PHH benchmarking contrasts are provided in Figures S3 (HepaRG vs PHH) and S4 (HuH7 vs PHH). These data establish that medium composition produces a reproducible, statistically robust, and cell line-specific transcriptional shift, yet the dominant transcriptomic differences across the study remain attributable to cell identity and PHH divergence, rather than medium alone.

### Functional enrichment and pathway-level interpretation of medium-associated changes

To interpret the biological meaning of medium-dependent transcriptional changes, GO:BP enrichment was performed separately for genes upregulated and downregulated in CDM versus FBS-SM within each hepatic cell line.

In HepaRG cells, transcripts increased in CDM were strongly enriched for canonical hepatic metabolism and transport processes, including organic acid catabolic process, response to xenobiotic stimulus, xenobiotic metabolic process, and organic anion transport, together with broader lipid and small-molecule metabolic programs (e.g., fatty acid and triglyceride metabolism; Figure 3; Figure S5a). In contrast, genes reduced in CDM (i.e., higher in FBS-SM) were preferentially associated with processes less directly linked to hepatocyte metabolic specialization, including immune effector and adaptive immune response, cell adhesion and migration, and morphogenetic programs (e.g., epithelial morphogenesis; Figure 3; Figure S5b). Collectively, these enrichments indicate that replacing serum with a chemically defined formulation in HepaRG cells shifts the DEG landscape toward xenobiotic handling and metabolic functions while attenuating adhesion/immune-associated transcriptional programs that are commonly accentuated under serum-supplemented culture.

**Figure 3.**
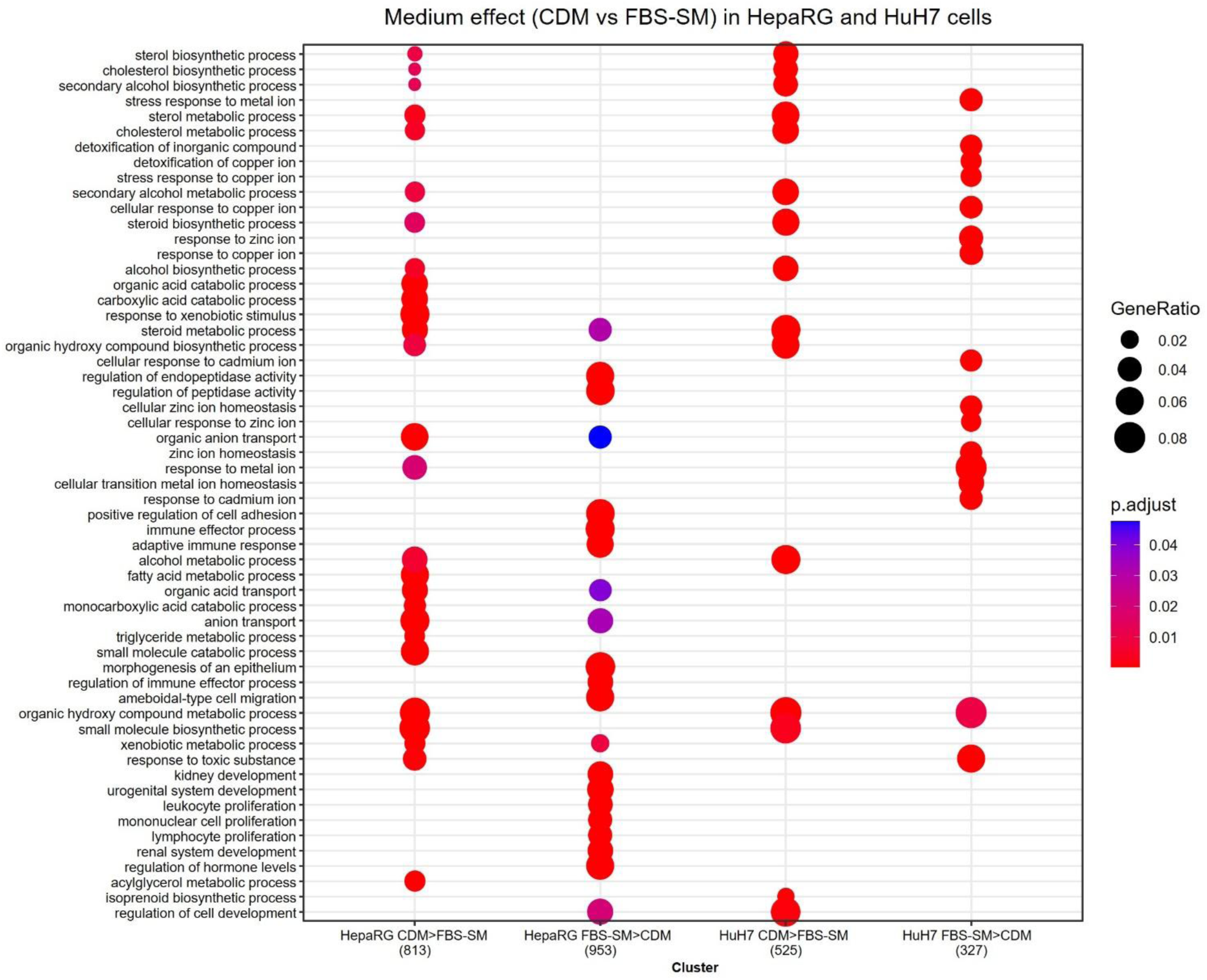
GO:BP enrichment of medium-dependent DEGs. CompareCluster dotplot summarizing Gene Ontology biological process (GO:BP) over-representation analysis for genes differentially expressed in CDM versus FBS-SM within HepaRG and HuH7 cells, stratified by direction (CDM > FBS-SM and FBS-SM > CDM; gene counts per cluster are shown on the x-axis). Dot size represents GeneRatio and dot color represents adjusted P value (Benjamini-Hochberg). Enriched terms were simplified to reduce semantic redundancy (Wang semantic similarity-based filtering), retaining representative terms.

In HuH7 cells, the CDM-induced signature differed qualitatively from HepaRG cells. Genes upregulated in CDM were dominated by lipid-associated biosynthetic programs, most prominently sterol biosynthetic process and sterol metabolic process-consistent with a medium-sensitive bias toward cholesterol/sterol pathway regulation in this model (Figure 3). Conversely, transcripts higher in FBS-SM were enriched for metal ion stress and detoxification-related processes (e.g., stress response to metal ion, detoxification of inorganic compounds, zinc and copper homeostasis), indicating a serum-associated shift toward ion-handling and stress-response programs in HuH7 cells (Figure 3; Figure S6).

Thus, although both cell lines responded to CDM, functional interpretation demonstrates that the direction and content of medium-driven reprogramming is model-dependent, with HepaRG cells preferentially engaging xenobiotic metabolism/transport programs, whereas HuH7 cells primarily exhibited modulation of sterol and stress/homeostasis-associated biology.

To complement over-representation analysis on filtered DEGs, we quantified pathway-level activity changes using single-sample gene set enrichment analysis (ssGSEA), which computes per-sample enrichment scores of genes within a given set. ssGSEA applied to curated QuickGO term sets grouped into hepatic-relevant themes (transport/secretion, xenobiotic detoxification, lipid metabolism, energy/mitochondria, and stress/homeostasis). The resulting theme-organized heatmap (Figure 4) provides a continuous, module-level view of how medium composition reshapes coordinated gene-set expression within each cell line. Consistent with GO enrichment, ssGSEA captured medium-associated modulation of xenobiotic-, lipid-, and transport-related modules while also highlighting broader stress/homeostasis signatures that may be incompletely represented by filtered DEG lists. The complete ssGSEA score matrices, delta outputs, and QuickGO term-set definitions are reported in Table S7.

**Figure 4.**
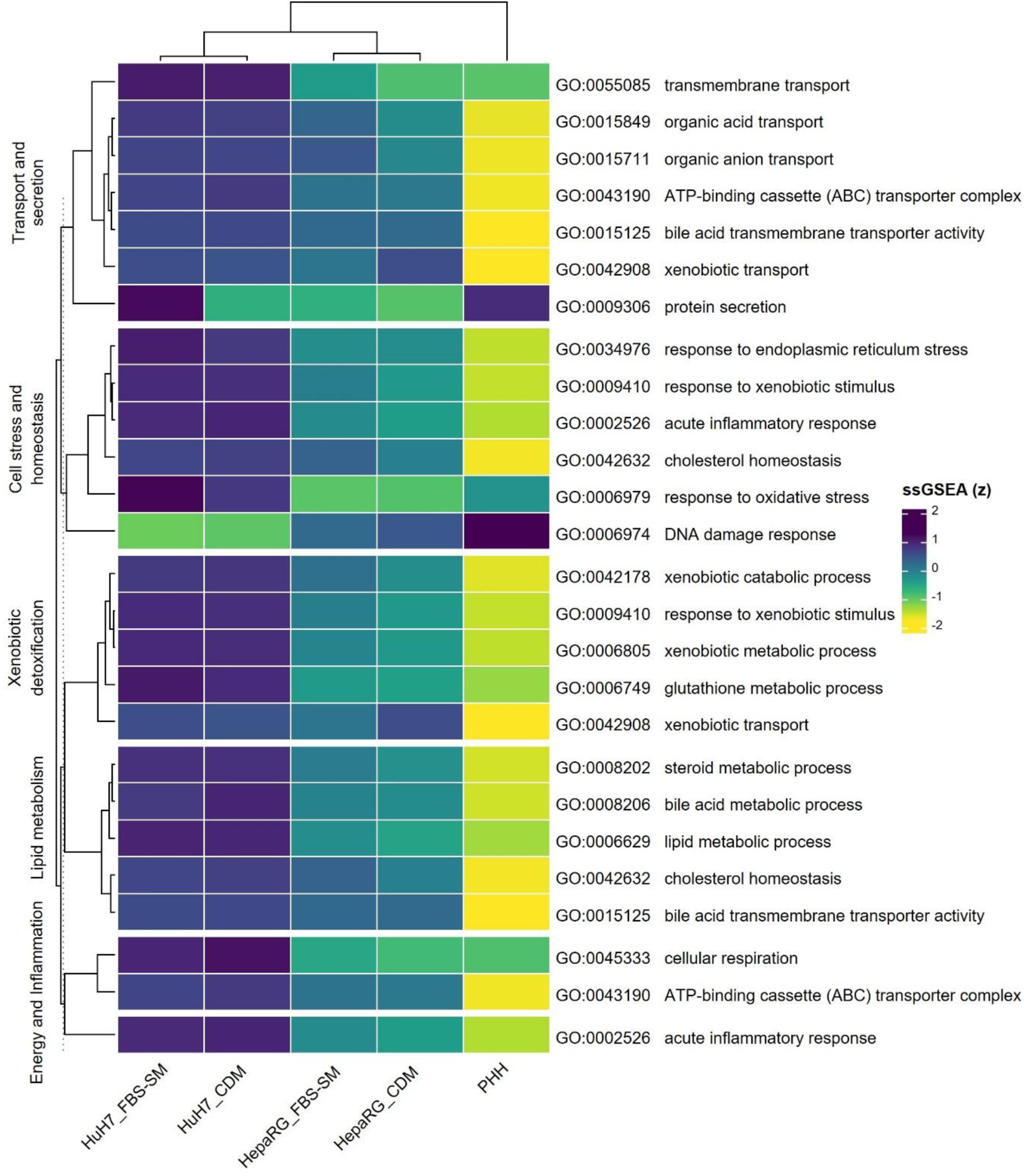
Theme-organized ssGSEA profiling using QuickGO-derived GO:BP term sets. Heatmap of single-sample gene set enrichment analysis (ssGSEA) scores (z-scored) computed from DESeq2 VST expression values using GSVA (ssGSEA method). GO:BP term sets curated from QuickGO were organized into hepatic-relevant themes (e.g., transport/secretion, stress/homeostasis, xenobiotic detoxification, lipid metabolism, energy/inflammation), and scores were summarized at the condition level to compare pathway activity patterns across HuH7, HepaRG cells, and PHH.

The GO:BP enrichment summary (Figure 3; Table S8); Figure S5a, b and S6a, b) and QuickGO ssGSEA profiling (Figure 4; Table S7) establish that CDM induces structured functional reprogramming that differs between HepaRG and HuH7 cells. Direction-resolved GO:BP enrichment for the PHH benchmarking contrasts (HepaRG CDM vs PHH and HuH7 CDM vs PHH) is provided in Figures S7 and S8.

### PHH-referenced benchmarking of hepatic identity and detoxification modules

To determine whether medium-dependent transcriptional remodeling translates into improved hepatic fidelity, we benchmarked both cell lines against PHH using curated marker panels capturing hepatic identity and core detoxification functions. This PHH-referenced framework evaluates whether shifting from FBS-SM to CDM moves each model toward an adult hepatocyte-like transcriptional state, rather than solely quantifying differential expression.

Across hepatic identity markers, HepaRG cells exhibited substantially higher concordance with PHH than HuH7 cells, and this alignment increased under CDM. The identity marker heatmap showed that HepaRG cells cultured in CDM more closely recapitulated PHH-associated expression patterns across multiple hepatic programs, whereas HuH7 cells remained comparatively divergent under both media conditions (Figure 5). Quantitative PHH similarity scoring (based on RMSA) corroborated these patterns, demonstrating consistently higher similarity for HepaRG cells than HuH7 cells across identity panels, with the largest improvement observed for HepaRG cells in CDM relative to FBS-SM (Figure S9). Complete panel-level similarity scores and marker definitions are provided in Table S2.

**Figure 5.**
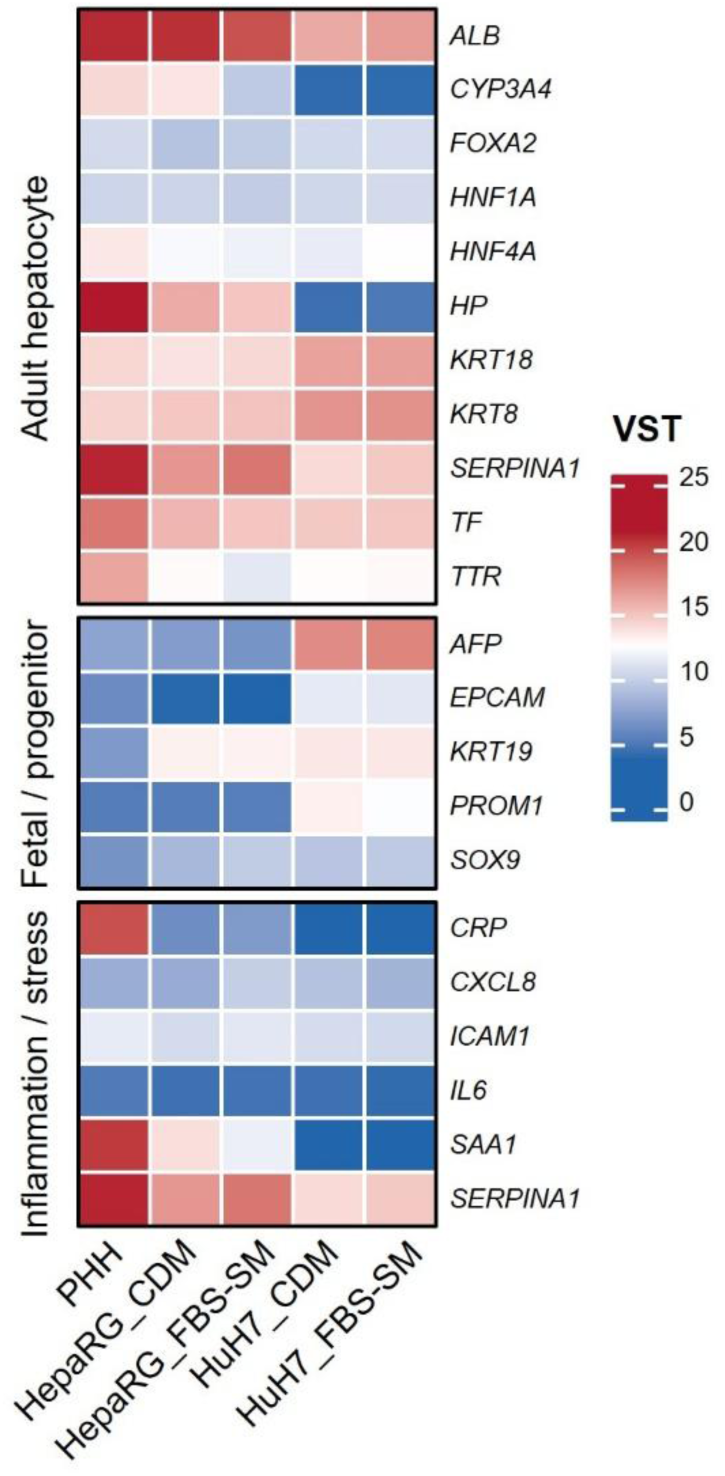
Hepatic identity marker expression across models and media, benchmarked to PHH. Heatmap of condition-mean DESeq2 VST expression values for curated hepatic identity markers across PHH, HepaRG cells (CDM or FBS-SM), and HuH7 cells (CDM or FBS-SM). Markers are grouped by biological category (adult hepatocyte, fetal/progenitor, inflammation/stress). Color indicates VST expression (scale shown).

We next evaluated detoxification competence using curated panels for Phase I enzymes, Phase II conjugation pathways, and drug transporters, which together capture key determinants of hepatic xenobiotic handling. Gene-level visualization of detox panel expression patterns is provided as dedicated heatmaps for Phase I, Phase II, and transporters (Figure 6a-c), PHH-referenced similarity scores again revealed a model-dependent medium effect: HepaRG cells showed improved alignment to PHH under CDM across detoxification panels, whereas HuH7 cells displayed minimal concordance with PHH in both CDM and FBS-SM (Figure 6d). The full similarity score outputs are reported in Table S1. These findings are consistent with the functional enrichment and ssGSEA results, in which HepaRG cells cultured in CDM preferentially engaged xenobiotic and transport-associated programs, while HuH7 cells exhibited medium sensitivity largely within sterol-related and stress/homeostasis processes.

**Figure 6.**
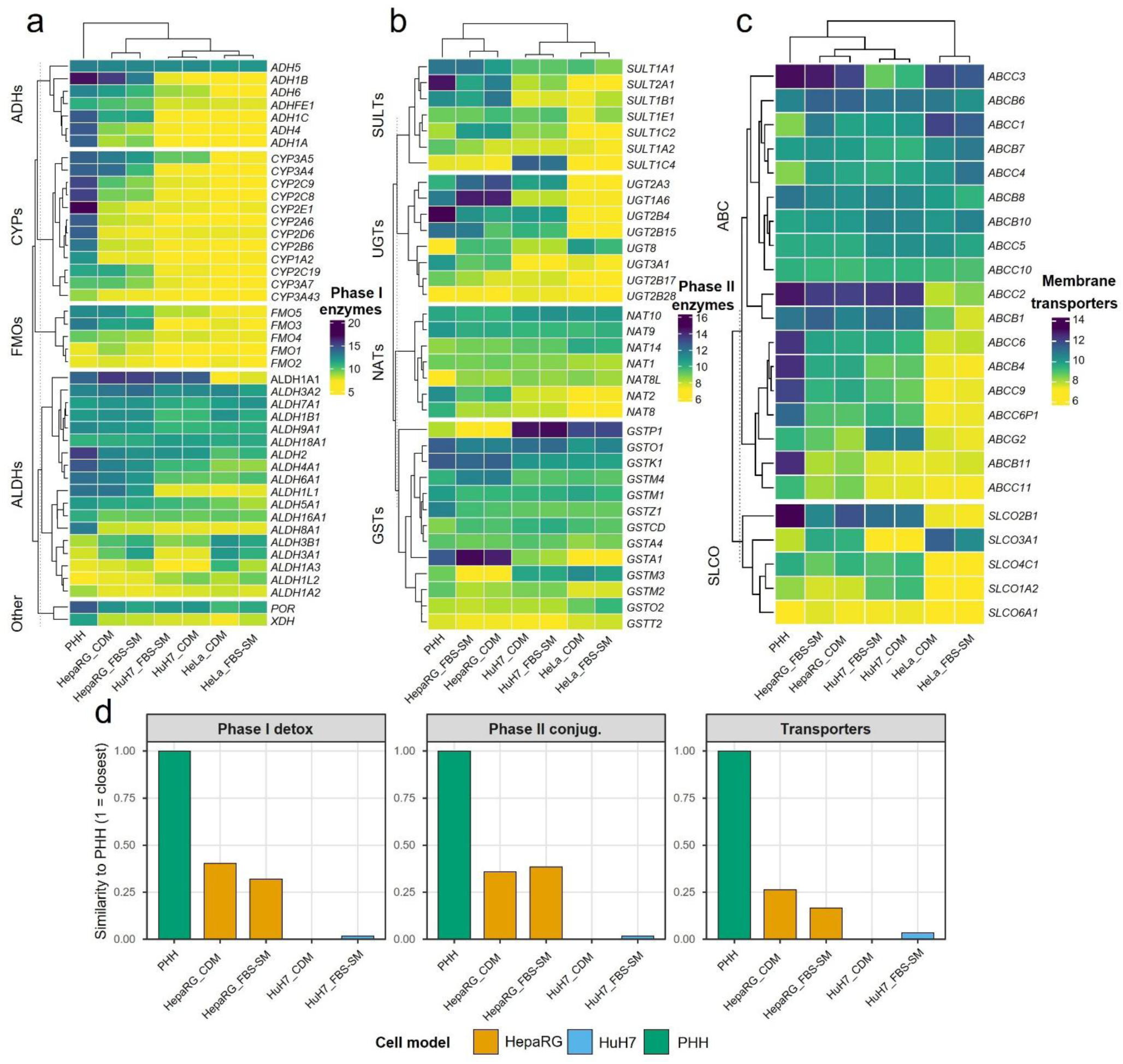
Detoxification gene expression and PHH similarity across Phase I/II enzymes and transporters. Heatmaps of condition-mean DESeq2 VST expression values for (a) Phase I enzymes, (b) Phase II conjugation enzymes, and (c) membrane transporters across PHH, HepaRG cells (CDM or FBS-SM), HuH7 cells (CDM or FBS-SM), and HeLa cells (CDM or FBS-SM). Genes are grouped by functional subfamilies within each panel, and columns represent condition-level profiles. (d) PHH-referenced similarity scores for each detoxification module, computed from gene-set expression deviations relative to PHH using an RMSE-based normalization (PHH fixed at 1).

The benchmarking analyses establish that CDM-driven transcriptional reprogramming in HepaRG cells is accompanied by increased PHH-like hepatic identity and detoxification module activity, supporting improved physiological relevance under chemically defined conditions. In contrast, HuH7 cells remain transcriptionally distant from PHH across both identity and detoxification gene panels, indicating that medium composition modulates specific pathways but does not overcome cell line-intrinsic limitations in adult hepatocyte fidelity. A full resolution ssGSEA pathway heatmap benchmarked to PHH is provided in Figure S10.

## Discussion

An ongoing challenge in hepatic *in vitro* modeling is that widely used “standard” culture conditions, most notably FBS supplementation, introduce an ill-defined and batch-variable biochemical environment. However, replacing FBS-SM with CDM can raise concern that future results will not be directly comparable with historical literature, even if legacy datasets partly reflect serum-conditioned biology rather than physiology. Against this backdrop, CDM should be viewed as a return to controlled biology, rather than an “artificial” condition that compromises relevance. In liver systems specifically, multiple lines of evidence support that defined, serum-free/CDM conditions can preserve or enhance adult hepatocyte-like biology. A landmark example is the 3D PHH spheroid system cultured in chemically defined, serum-free conditions, where whole-proteome analyses indicated that spheroids closely resembled the in vivo liver tissue from which they originated [41]. Another early study demonstrated maintenance of PHH morphology, transcription factors, and liver-specific functions for weeks in a chemically defined, serum-free setting [42]. For hepatic cell line models used in toxicology, a recent comparative proteomics study in HepG2 explicitly benchmarked serum-free cultivation against FBS-SM and reported coordinated upregulation of drug metabolism pathways and oxidative stress protection at the protein level [43]. Complementing this, an in-depth proteome analysis of differentiated HepaRG cells cultivated in a defined serum-free medium highlighted robust expression of xenobiotic metabolism and NRF2-mediated oxidative-stress response proteins that were generally closer to primary human hepatocytes than those observed in HepG2 [44].

Placed within this literature, our results support an “identity-first, medium-modulated” view of hepatic transcriptomes. Model identity remains the dominant constraint on global gene-expression state, while medium composition introduces a minimum secondary effect. In differentiated HepaRG cells, widely regarded as the metabolically most competent human hepatic cell line system for drug metabolism and hepatotoxicity applications, the switch from FBS-SM to CDM increased the expression of gene programs central to PHH biology, including xenobiotic handling and detoxification pathways, Phase I/Phase II metabolism, and hepatocyte-relevant transport functions. In parallel, PHH-referenced benchmarking indicated that HepaRG cells cultured in CDM aligned more closely with the PHH transcriptomic profile than HepaRG cells maintained in FBS-SM.

In contrast, HuH7 cells exhibited a qualitatively different mode of medium responsiveness. CDM-associated upregulated genes were dominated by sterol biosynthesis and sterol metabolic processes. This pattern is consistent with reported literature of serum-free adaptation, where serum removal reweighs lipid-handling programs in CHO cells [45]. Such a shift may represent conserved responses to altered lipid availability, carrier proteins, and growth-factor contexts when serum is removed, rather than being idiosyncratic artifacts. A similar pattern was observed in HuH7.5 cells culture when switched from FBS-supplemented culture to human serum-supplemented culture [46].

Several limitations should be noted. First, “CDM” formulations are not interchangeable; medium composition, lipid supplementation, and adaptation strategy can substantially influence metabolic set points. Second, the PHH reference used here comprised cryopreserved pooled hepatocytes processed without extended culture, which improves standardization but does not capture time-in-culture adaptation and phenotype drift observed in plated PHH systems; this should be considered when comparing across studies using different PHH configurations. Third, the CDM used here were cell line-optimized and did not constitute a simple “FBS-omission” condition; rather, they were designed as defined alternatives to (i) replace FBS-SM without disproportionately altering cell identity, (ii) improve reproducibility through compositional transparency, and (iii) reduce serum-driven signaling to bias cellular programs toward PHH-relevant biology.

In summary, the present study supports that adopting CDM should not be viewed as a disruptive break from serum-based literature, but rather as a controlled refinement that clarifies which programs reflect model-intrinsic biology and which are modulated by serum. Global analyses showed that cell type remained the dominant driver of variance, and CDM versus FBS-SM shifts within a given model were substantially smaller than contrasts between cell lines or versus PHH, supporting continued interpretability of legacy datasets while treating medium an explicitly controlled culture variable. Crucially, CDM provided a measurable gain in physiological relevance for differentiated HepaRG cells, where pathway and gene-panel benchmarking demonstrated increased alignment to PHH across hepatic identity and detoxification modules, consistent with reinforcement of hepatocyte-specialized programs under defined conditions. Therefore, HepaRG in CDM can improve hepatotoxicity testing by reducing serum-related confounding and yielding more consistent, interpretable toxicity readouts.

## Data availability

RNA-seq data are available in the NCBI GEO repository (GSE324107), and can be accessed at: https://www.ncbi.nlm.nih.gov/geo/query/acc.cgi?acc=GSE324107.

All data processing and downstream analyses were performed using a fully scripted, version-controlled R workflow. The complete analysis code is available on request.

## Authorship contribution statement

**Ahmed S. M. Ali**: Conceptualization, Writing - original draft, Data curation, Data Visualization, Writing - Review & Editing, Methodology, Investigation. **Heike Sprenger**: Conceptualization, Data curation, Data Visualization, Writing - Review & Editing, Methodology, Investigation. **Albert Braeuning**: Conceptualization, Funding acquisition, Writing - Review & Editing. **Jens Kurreck**: Conceptualization, Supervision, Funding acquisition, Writing - Review & Editing.

## Acknowledgments

We are thankful for the financial support of the research project by the Einstein Foundation Berlin (Einstein Center 3R, EZ-2020-597-2). We gratefully acknowledge the technical assistance by Viola Roehrs.

## Conflict of Interest

The authors declare no competing interests.

## Declaration of generative AI and AI-assisted technologies in the writing process

During the preparation of this manuscript, the authors used ChatGPT to improve readability. All content generated with this assistance was critically reviewed and revised by the authors as necessary, and the authors assume full responsibility for the accuracy, and final content of the publication.

## Notes

https://www.ncbi.nlm.nih.gov/geo/query/acc.cgi?acc=GSE324107

### Competing Interest Statement

The authors have declared no competing interest.

