## Supplementary material for "Cell line identity rather than medium composition determines transcriptomic profiles of HepaRG and HuH7 cells cultured in chemically defined or serum-based media: comparison with primary human hepatocytes": Figure S10

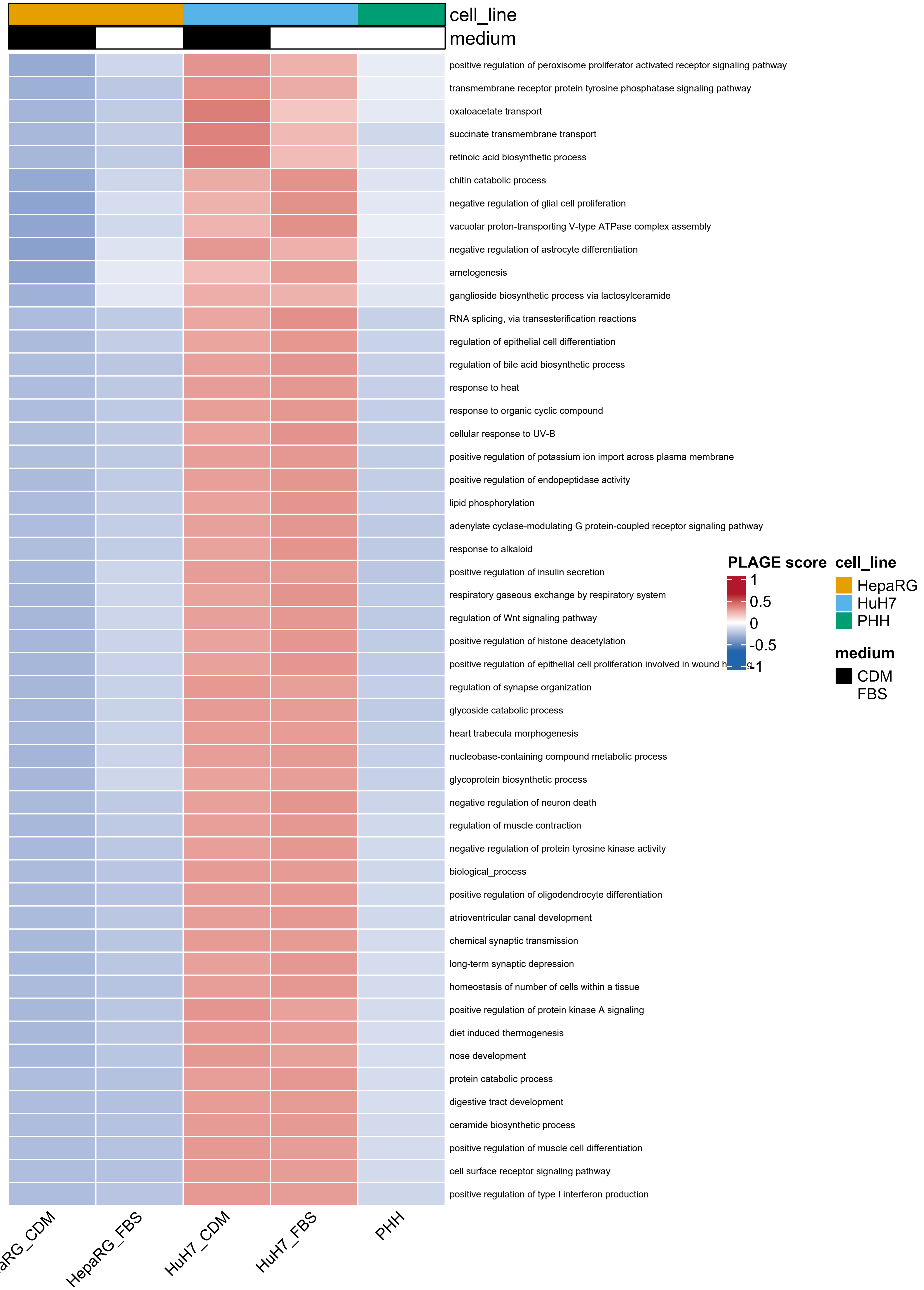

cell_line
medium
positive regulation of peroxisome proliferator activated receptor signaling pathway
transmembrane receptor protein tyrosine phosphatase signaling pathway
oxaloacetate transport
succinate transmembrane transport
retinoic acid biosynthetic process
chitin catabolic process
negative regulation of glial cell proliferation
vacuolar proton-transporting V-type ATPase complex assembly
negative regulation of astrocyte differentiation
amelogenesis
ganglioside biosynthetic process via lactosylceramide
RNA splicing, via transesterification reactions
regulation of epithelial cell differentiation
regulation of bile acid biosynthetic process
response to heat
response to organic cyclic compound
cellular response to UV-B
positive regulation of potassium ion import across plasma membrane
positive regulation of endopeptidase activity
lipid phosphorylation
adenylate cyclase-modulating G protein-coupled receptor signaling pathway
response to alkaloid
positive regulation of insulin secretion
respiratory gaseous exchange by respiratory system
regulation of Wnt signaling pathway
positive regulation of histone deacetylation
positive regulation of epithelial cell proliferation involved in wound h
regulation of synapse organization
glycoside catabolic process
heart trabecula morphogenesis
nucleobase-containing compound metabolic process
glycoprotein biosynthetic process
negative regulation of neuron death
regulation of muscle contraction
negative regulation of protein tyrosine kinase activity
biological_process
positive regulation of oligodendrocyte differentiation
atrioventricular canal development
chemical synaptic transmission
long-term synaptic depression
homeostasis of number of cells within a tissue
positive regulation of protein kinase A signaling
diet induced thermogenesis
nose development
protein catabolic process
digestive tract development
ceramide biosynthetic process
positive regulation of muscle cell differentiation
cell surface receptor signaling pathway
positive regulation of type I interferon production
PLAGE score
1
0.5
0
-0.5
-1
cell_line
HepaRG
HuH7
PHH
medium
CDM
FBS
HepaRG_FBS
HuH7_CDM
HuH7_FBS
PHH

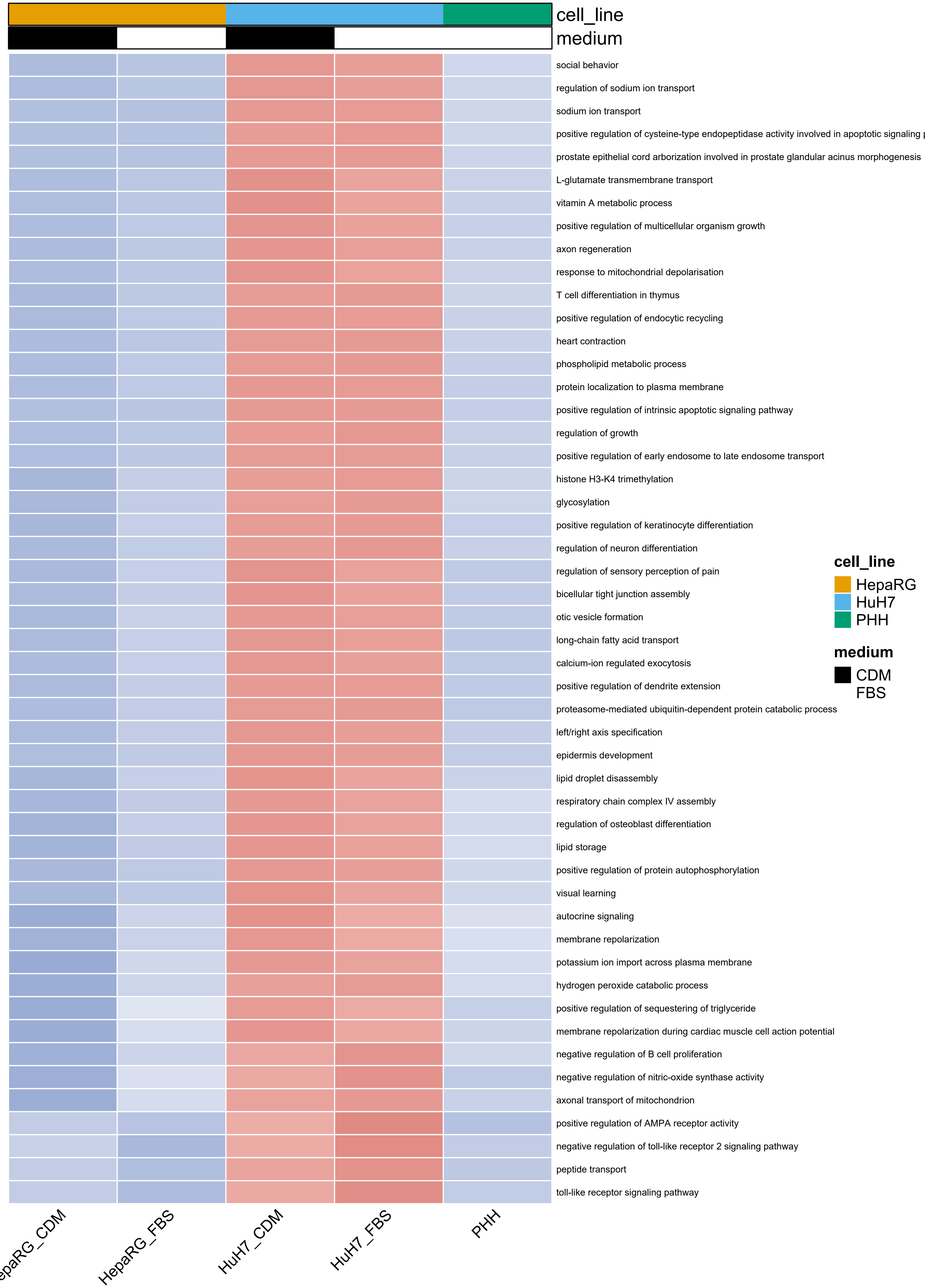
cell_line
medium
social behavior
regulation of sodium ion transport
sodium ion transport
positive regulation of cysteine-type endopeptidase activity involved in apoptotic signaling
prostate epithelial cord arborization involved in prostate glandular acinus morphogenesis
L-glutamate transmembrane transport
vitamin A metabolic process
positive regulation of multicellular organism growth
axon regeneration
response to mitochondrial depolarisation
T cell differentiation in thymus
positive regulation of endocytic recycling
heart contraction
phospholipid metabolic process
protein localization to plasma membrane
positive regulation of intrinsic apoptotic signaling pathway
regulation of growth
positive regulation of early endosome to late endosome transport
histone H3-K4 trimethylation
glycosylation
positive regulation of keratinocyte differentiation
regulation of neuron differentiation
regulation of sensory perception of pain
bicellular tight junction assembly
otic vesicle formation
long-chain fatty acid transport
calcium-ion regulated exocytosis
positive regulation of dendrite extension
proteasome-mediated ubiquitin-dependent protein catabolic process
left/right axis specification
epidermis development
lipid droplet disassembly
respiratory chain complex IV assembly
regulation of osteoblast differentiation
lipid storage
positive regulation of protein autophosphorylation
visual learning
autocrine signaling
membrane repolarization
potassium ion import across plasma membrane
hydrogen peroxide catabolic process
positive regulation of sequestering of triglyceride
membrane repolarization during cardiac muscle cell action potential
negative regulation of B cell proliferation
negative regulation of nitric-oxide synthase activity
axonal transport of mitochondrion
positive regulation of AMPA receptor activity
negative regulation of toll-like receptor 2 signaling pathway
peptide transport
toll-like receptor signaling pathway
cell_line
HepaRG
HuH7
PHH
medium
CDM
FBS
HepaRG_FBS
HuH7_CDM
HuH7_FBS
PHH

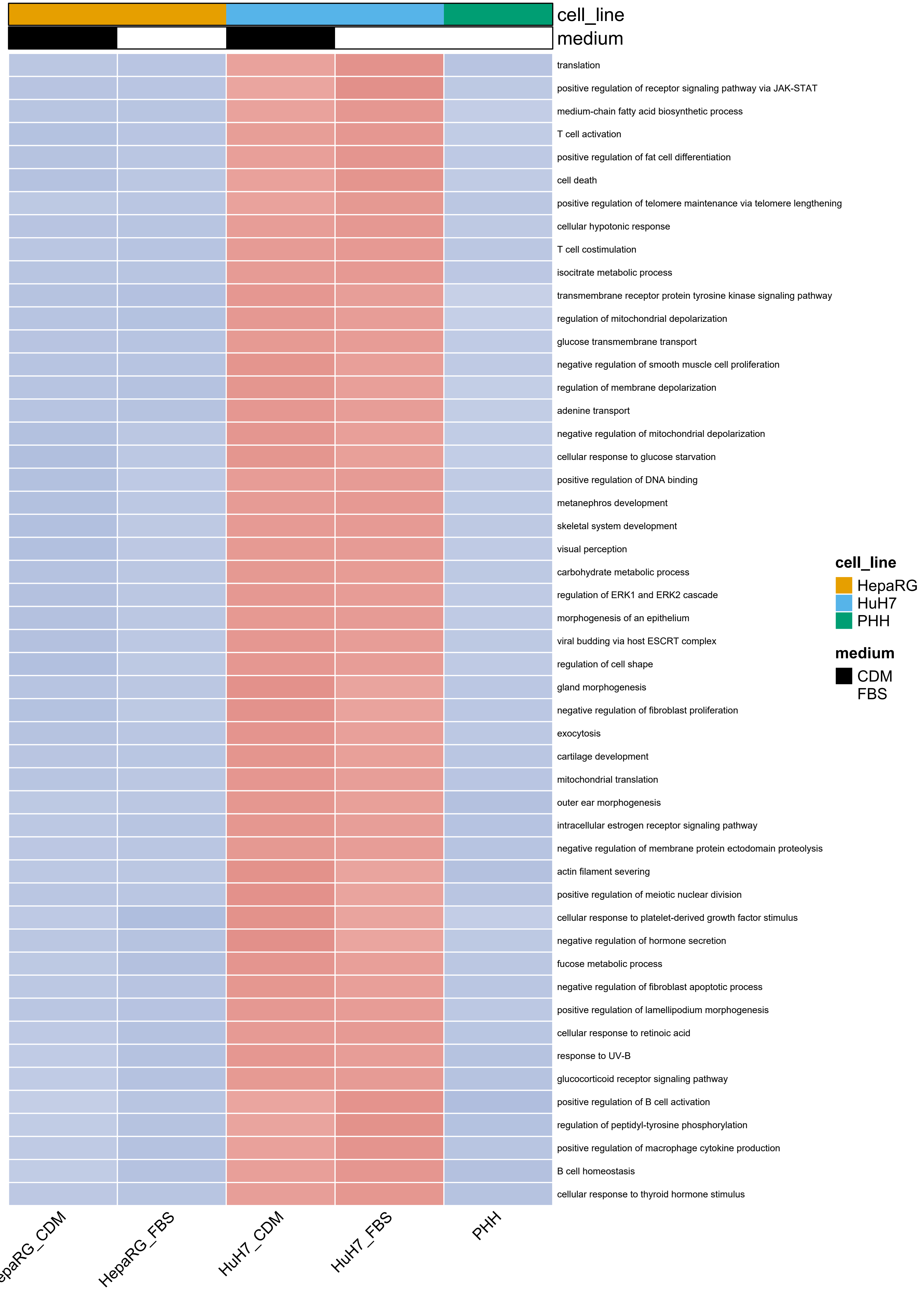
cell_line
medium
translation
positive regulation of receptor signaling pathway via JAK-STAT
medium-chain fatty acid biosynthetic process
T cell activation
positive regulation of fat cell differentiation
cell death
positive regulation of telomere maintenance via telomere lengthening
cellular hypotonic response
T cell costimulation
isocitrate metabolic process
transmembrane receptor protein tyrosine kinase signaling pathway
regulation of mitochondrial depolarization
glucose transmembrane transport
negative regulation of smooth muscle cell proliferation
regulation of membrane depolarization
adenine transport
negative regulation of mitochondrial depolarization
cellular response to glucose starvation
positive regulation of DNA binding
metanephros development
skeletal system development
visual perception
carbohydrate metabolic process
regulation of ERK1 and ERK2 cascade
morphogenesis of an epithelium
viral budding via host ESCRT complex
regulation of cell shape
gland morphogenesis
negative regulation of fibroblast proliferation
exocytosis
cartilage development
mitochondrial translation
outer ear morphogenesis
intracellular estrogen receptor signaling pathway
negative regulation of membrane protein ectodomain proteolysis
actin filament severing
positive regulation of meiotic nuclear division
cellular response to platelet-derived growth factor stimulus
negative regulation of hormone secretion
fucose metabolic process
negative regulation of fibroblast apoptotic process
positive regulation of lamellipodium morphogenesis
cellular response to retinoic acid
response to UV-B
glucocorticoid receptor signaling pathway
positive regulation of B cell activation
regulation of peptidyl-tyrosine phosphorylation
positive regulation of macrophage cytokine production
B cell homeostasis
cellular response to thyroid hormone stimulus
cell_line
HepaRG
HuH7
PHH
medium
CDM
FBS
HepaRG_FBS
HuH7_CDM
HuH7_FBS
PHH

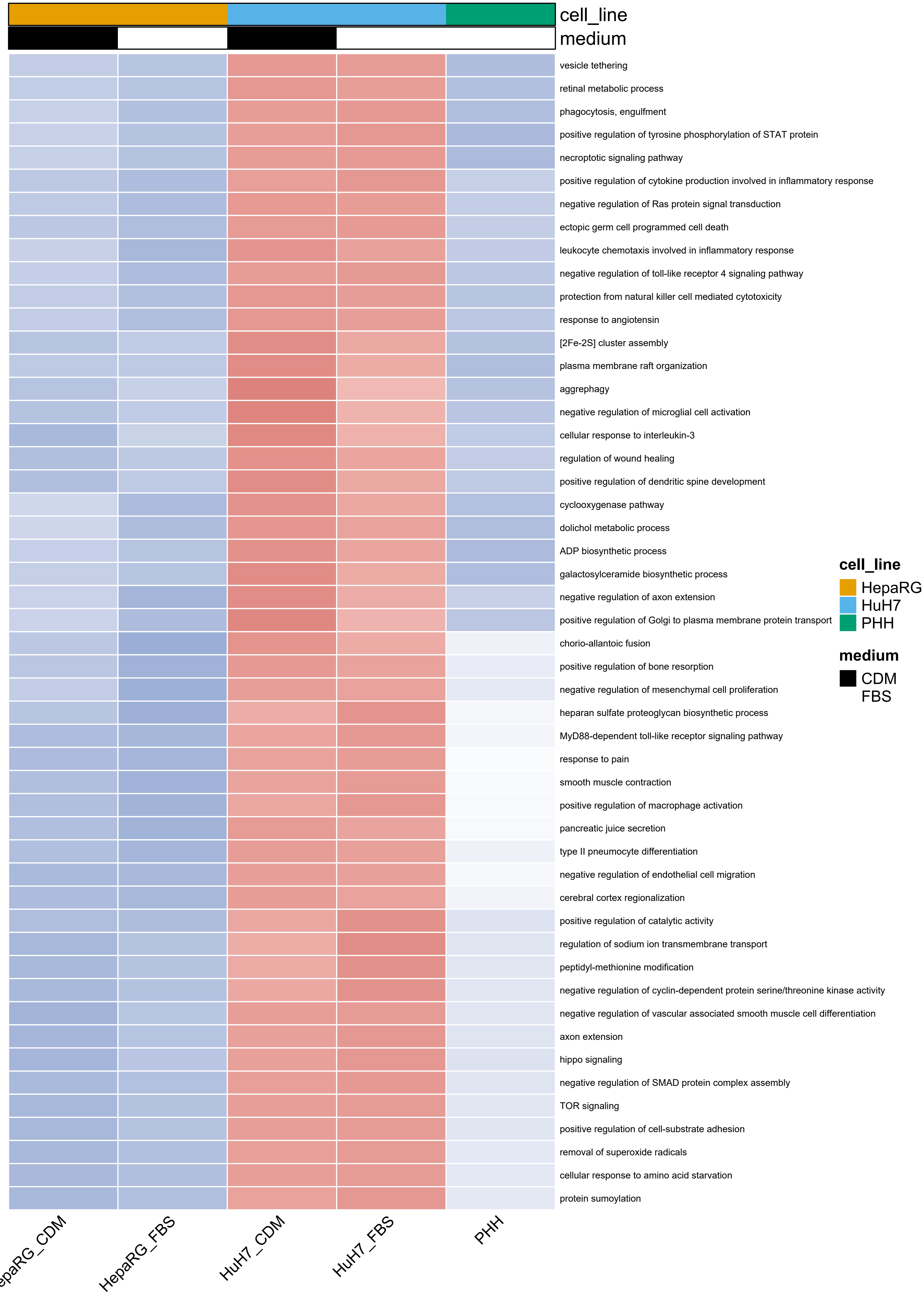
cell_line
medium
vesicle tethering
retinal metabolic process
phagocytosis, engulfment
positive regulation of tyrosine phosphorylation of STAT protein
necroptotic signaling pathway
positive regulation of cytokine production involved in inflammatory response
negative regulation of Ras protein signal transduction
ectopic germ cell programmed cell death
leukocyte chemotaxis involved in inflammatory response
negative regulation of toll-like receptor 4 signaling pathway
protection from natural killer cell mediated cytotoxicity
response to angiotensin
[2Fe-2S] cluster assembly
plasma membrane raft organization
aggrephagy
negative regulation of microglial cell activation
cellular response to interleukin-3
regulation of wound healing
positive regulation of dendritic spine development
cyclooxygenase pathway
dolichol metabolic process
ADP biosynthetic process
galactosylceramide biosynthetic process
negative regulation of axon extension
positive regulation of Golgi to plasma membrane protein transport
chorio-allantoic fusion
positive regulation of bone resorption
negative regulation of mesenchymal cell proliferation
heparan sulfate proteoglycan biosynthetic process
MyD88-dependent toll-like receptor signaling pathway
response to pain
smooth muscle contraction
positive regulation of macrophage activation
pancreatic juice secretion
type II pneumocyte differentiation
negative regulation of endothelial cell migration
cerebral cortex regionalization
positive regulation of catalytic activity
regulation of sodium ion transmembrane transport
peptidyl-methionine modification
negative regulation of cyclin-dependent protein serine/threonine kinase activity
negative regulation of vascular associated smooth muscle cell differentiation
axon extension
hippo signaling
negative regulation of SMAD protein complex assembly
TOR signaling
positive regulation of cell-substrate adhesion
removal of superoxide radicals
cellular response to amino acid starvation
protein sumoylation
HepaRG_FBS
HuH7_CDM
HuH7_FBS
PHH
cell_line
HepaRG
HuH7
PHH
medium
CDM
FBS

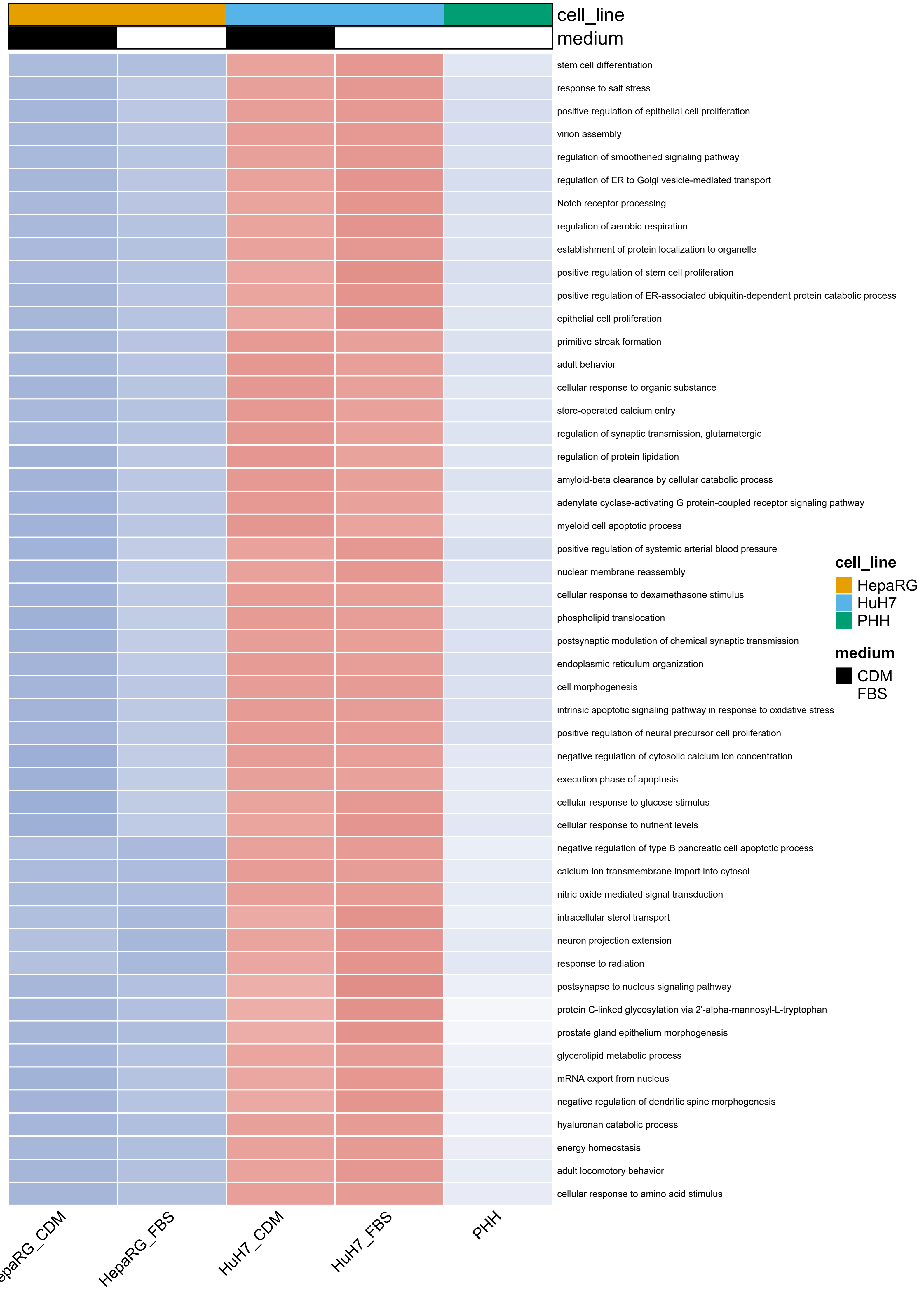
cell_line
medium
stem cell differentiation
response to salt stress
positive regulation of epithelial cell proliferation
virion assembly
regulation of smoothened signaling pathway
regulation of ER to Golgi vesicle-mediated transport
Notch receptor processing
regulation of aerobic respiration
establishment of protein localization to organelle
positive regulation of stem cell proliferation
positive regulation of ER-associated ubiquitin-dependent protein catabolic process
epithelial cell proliferation
primitive streak formation
adult behavior
cellular response to organic substance
store-operated calcium entry
regulation of synaptic transmission, glutamatergic
regulation of protein lipidation
amyloid-beta clearance by cellular catabolic process
adenylate cyclase-activating G protein-coupled receptor signaling pathway
myeloid cell apoptotic process
positive regulation of systemic arterial blood pressure
nuclear membrane reassembly
cellular response to dexamethasone stimulus
phospholipid translocation
postsynaptic modulation of chemical synaptic transmission
endoplasmic reticulum organization
cell morphogenesis
intrinsic apoptotic signaling pathway in response to oxidative stress
positive regulation of neural precursor cell proliferation
negative regulation of cytosolic calcium ion concentration
execution phase of apoptosis
cellular response to glucose stimulus
cellular response to nutrient levels
negative regulation of type B pancreatic cell apoptotic process
calcium ion transmembrane import into cytosol
nitric oxide mediated signal transduction
intracellular sterol transport
neuron projection extension
response to radiation
postsynapse to nucleus signaling pathway
protein C-linked glycosylation via 2'-alpha-mannosyl-L-tryptophan
prostate gland epithelium morphogenesis
glycerolipid metabolic process
mRNA export from nucleus
negative regulation of dendritic spine morphogenesis
hyaluronan catabolic process
energy homeostasis
adult locomotory behavior
cellular response to amino acid stimulus
cell_line
HepaRG
HuH7
PHH
medium
CDM
FBS
HepaRG_FBS
HuH7_CDM
HuH7_FBS
PHH

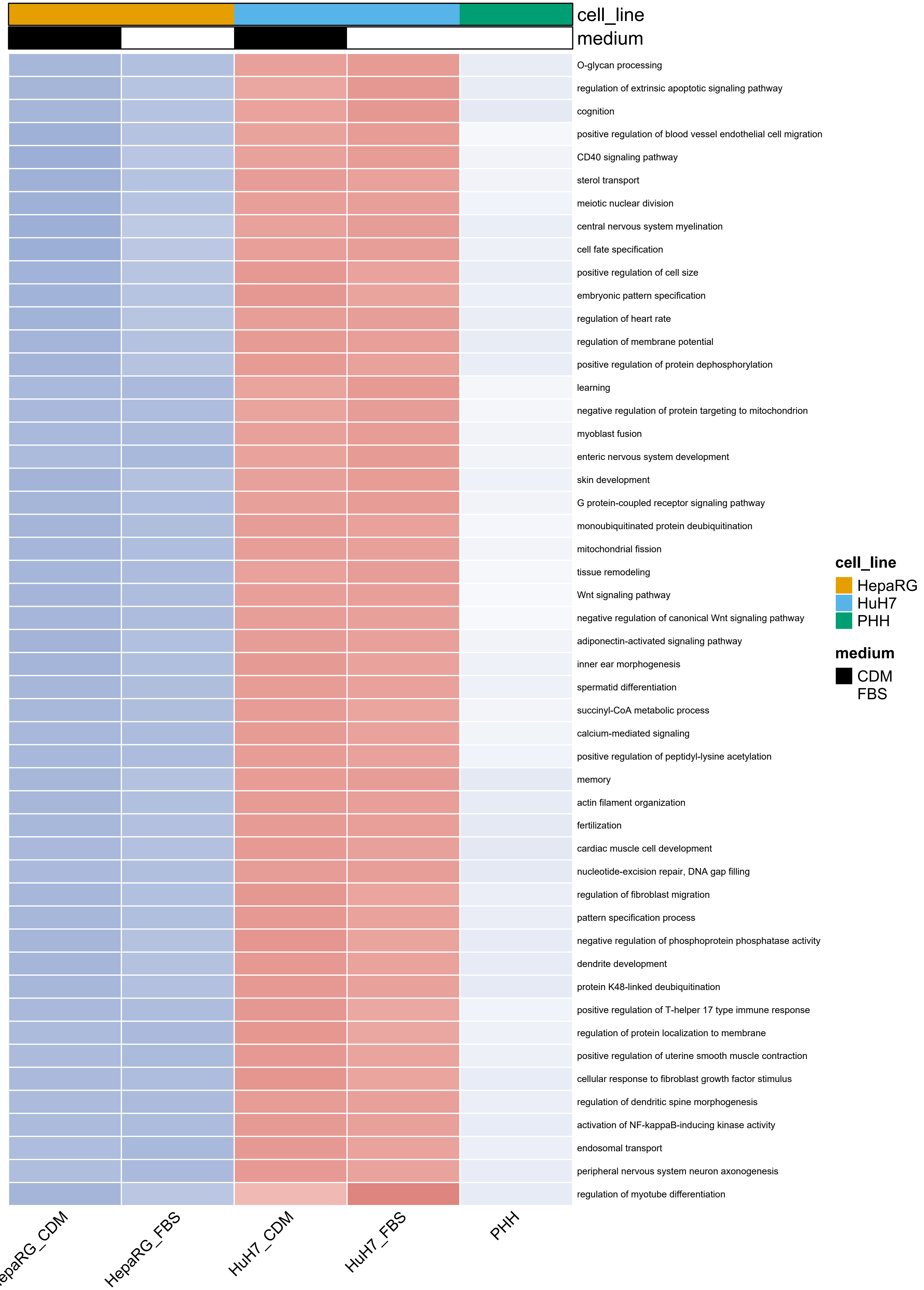

cell_line
medium
O-glycan processing
regulation of extrinsic apoptotic signaling pathway
cognition
positive regulation of blood vessel endothelial cell migration
CD40 signaling pathway
sterol transport
meiotic nuclear division
central nervous system myelination
cell fate specification
positive regulation of cell size
embryonic pattern specification
regulation of heart rate
regulation of membrane potential
positive regulation of protein dephosphorylation
learning
negative regulation of protein targeting to mitochondrion
myoblast fusion
enteric nervous system development
skin development
G protein-coupled receptor signaling pathway
monoubiquitinated protein deubiquitination
mitochondrial fission
tissue remodeling
Wnt signaling pathway
negative regulation of canonical Wnt signaling pathway
adiponectin-activated signaling pathway
inner ear morphogenesis
spermatid differentiation
succinyl-CoA metabolic process
calcium-mediated signaling
positive regulation of peptidyl-lysine acetylation
memory
actin filament organization
fertilization
cardiac muscle cell development
nucleotide-excision repair, DNA gap filling
regulation of fibroblast migration
pattern specification process
negative regulation of phosphoprotein phosphatase activity
dendrite development
protein K48-linked deubiquitination
positive regulation of T-helper 17 type immune response
regulation of protein localization to membrane
positive regulation of uterine smooth muscle contraction
cellular response to fibroblast growth factor stimulus
regulation of dendritic spine morphogenesis
activation of NF-kappaB-inducing kinase activity
endosomal transport
peripheral nervous system neuron axonogenesis
regulation of myotube differentiation
HepaRG_CDM
HepaRG_FBS
HuH7_CDM
HuH7_FBS
PHH
cell_line
HepaRG
HuH7
PHH
medium
CDM
FBS

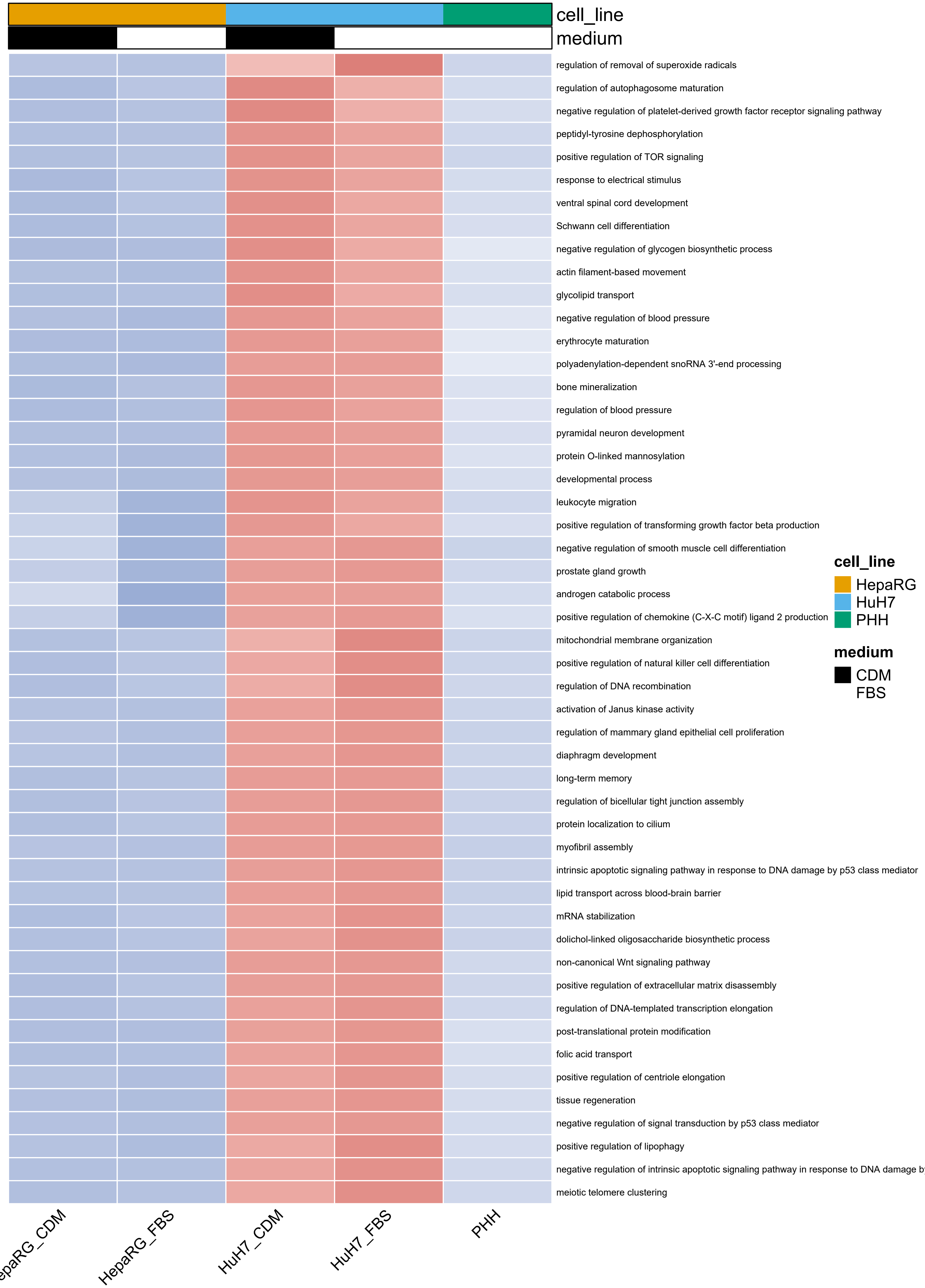
cell_line
medium
regulation of removal of superoxide radicals
regulation of autophagosome maturation
negative regulation of platelet-derived growth factor receptor signaling pathway
peptidyl-tyrosine dephosphorylation
positive regulation of TOR signaling
response to electrical stimulus
ventral spinal cord development
Schwann cell differentiation
negative regulation of glycogen biosynthetic process
actin filament-based movement
glycolipid transport
negative regulation of blood pressure
erythrocyte maturation
polyadenylation-dependent snoRNA 3'-end processing
bone mineralization
regulation of blood pressure
pyramidal neuron development
protein O-linked mannosylation
developmental process
leukocyte migration
positive regulation of transforming growth factor beta production
negative regulation of smooth muscle cell differentiation
prostate gland growth
androgen catabolic process
positive regulation of chemokine (C-X-C motif) ligand 2 production
mitochondrial membrane organization
positive regulation of natural killer cell differentiation
regulation of DNA recombination
activation of Janus kinase activity
regulation of mammary gland epithelial cell proliferation
diaphragm development
long-term memory
regulation of bicellular tight junction assembly
protein localization to cilium
myofibril assembly
intrinsic apoptotic signaling pathway in response to DNA damage by p53 class mediator
lipid transport across blood-brain barrier
mRNA stabilization
dolichol-linked oligosaccharide biosynthetic process
non-canonical Wnt signaling pathway
positive regulation of extracellular matrix disassembly
regulation of DNA-templated transcription elongation
post-translational protein modification
folic acid transport
positive regulation of centriole elongation
tissue regeneration
negative regulation of signal transduction by p53 class mediator
positive regulation of lipophagy
negative regulation of intrinsic apoptotic signaling pathway in response to DNA damage b
meiotic telomere clustering
cell_line
HepaRG
HuH7
PHH
medium
CDM
FBS
HepaRG_FBS
HuH7_CDM
HuH7_FBS
PHH

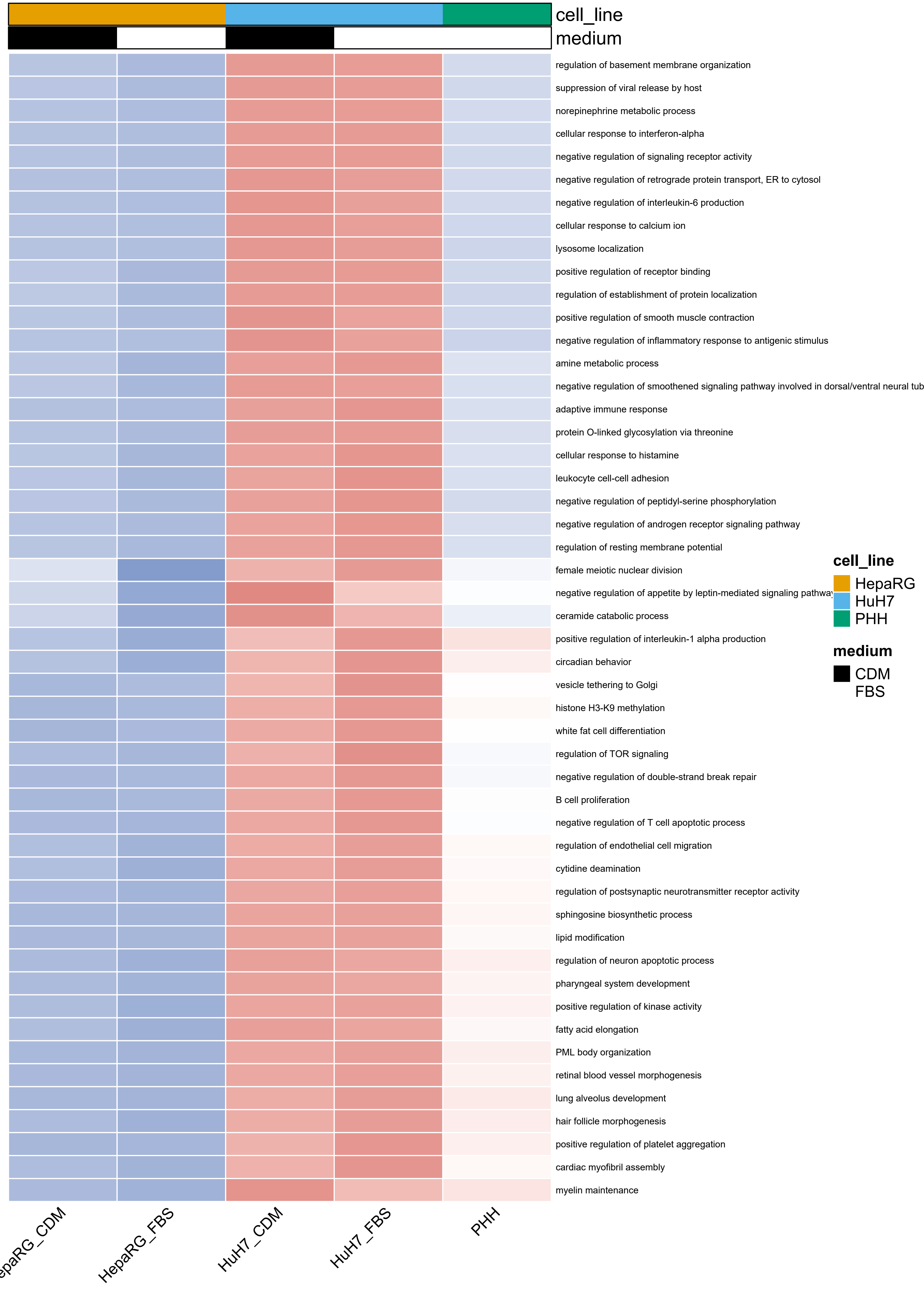

cell_line
medium
regulation of basement membrane organization
suppression of viral release by host
norepinephrine metabolic process
cellular response to interferon-alpha
negative regulation of signaling receptor activity
negative regulation of retrograde protein transport, ER to cytosol
negative regulation of interleukin-6 production
cellular response to calcium ion
lysosome localization
positive regulation of receptor binding
regulation of establishment of protein localization
positive regulation of smooth muscle contraction
negative regulation of inflammatory response to antigenic stimulus
amine metabolic process
negative regulation of smoothened signaling pathway involved in dorsal/ventral neural tub
adaptive immune response
protein O-linked glycosylation via threonine
cellular response to histamine
leukocyte cell-cell adhesion
negative regulation of peptidyl-serine phosphorylation
negative regulation of androgen receptor signaling pathway
regulation of resting membrane potential
female meiotic nuclear division
negative regulation of appetite by leptin-mediated signaling pathway
ceramide catabolic process
positive regulation of interleukin-1 alpha production
circadian behavior
vesicle tethering to Golgi
histone H3-K9 methylation
white fat cell differentiation
regulation of TOR signaling
negative regulation of double-strand break repair
B cell proliferation
negative regulation of T cell apoptotic process
regulation of endothelial cell migration
cytidine deamination
regulation of postsynaptic neurotransmitter receptor activity
sphingosine biosynthetic process
lipid modification
regulation of neuron apoptotic process
pharyngeal system development
positive regulation of kinase activity
fatty acid elongation
PML body organization
retinal blood vessel morphogenesis
lung alveolus development
hair follicle morphogenesis
positive regulation of platelet aggregation
cardiac myofibril assembly
myelin maintenance
cell_line
HepaRG
HuH7
PHH
medium
CDM
FBS
epaRG_CDM
HepaRG_FBS
HuH7_CDM
HuH7_FBS
PHH

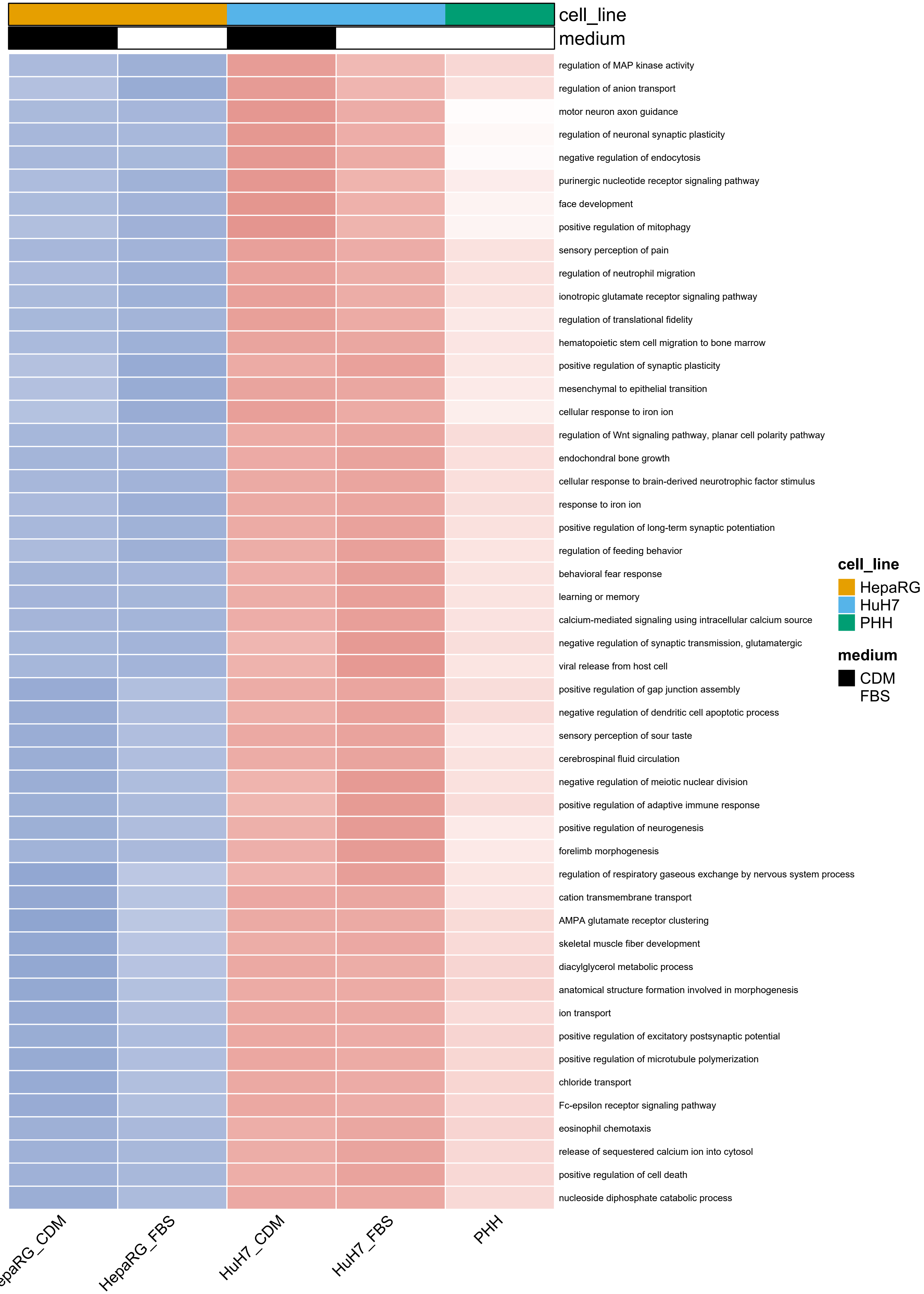
cell_line
medium
regulation of MAP kinase activity
regulation of anion transport
motor neuron axon guidance
regulation of neuronal synaptic plasticity
negative regulation of endocytosis
purinergic nucleotide receptor signaling pathway
face development
positive regulation of mitophagy
sensory perception of pain
regulation of neutrophil migration
ionotropic glutamate receptor signaling pathway
regulation of translational fidelity
hematopoietic stem cell migration to bone marrow
positive regulation of synaptic plasticity
mesenchymal to epithelial transition
cellular response to iron ion
regulation of Wnt signaling pathway, planar cell polarity pathway
endochondral bone growth
cellular response to brain-derived neurotrophic factor stimulus
response to iron ion
positive regulation of long-term synaptic potentiation
regulation of feeding behavior
behavioral fear response
learning or memory
calcium-mediated signaling using intracellular calcium source
negative regulation of synaptic transmission, glutamatergic
viral release from host cell
positive regulation of gap junction assembly
negative regulation of dendritic cell apoptotic process
sensory perception of sour taste
cerebrospinal fluid circulation
negative regulation of meiotic nuclear division
positive regulation of adaptive immune response
positive regulation of neurogenesis
forelimb morphogenesis
regulation of respiratory gaseous exchange by nervous system process
cation transmembrane transport
AMPA glutamate receptor clustering
skeletal muscle fiber development
diacylglycerol metabolic process
anatomical structure formation involved in morphogenesis
ion transport
positive regulation of excitatory postsynaptic potential
positive regulation of microtubule polymerization
chloride transport
Fc-epsilon receptor signaling pathway
eosinophil chemotaxis
release of sequestered calcium ion into cytosol
positive regulation of cell death
nucleoside diphosphate catabolic process
epaRG_CDM
HepaRG_FBS
HuH7_CDM
HuH7_FBS
PHH
cell_line
HepaRG
HuH7
PHH
medium
CDM
FBS

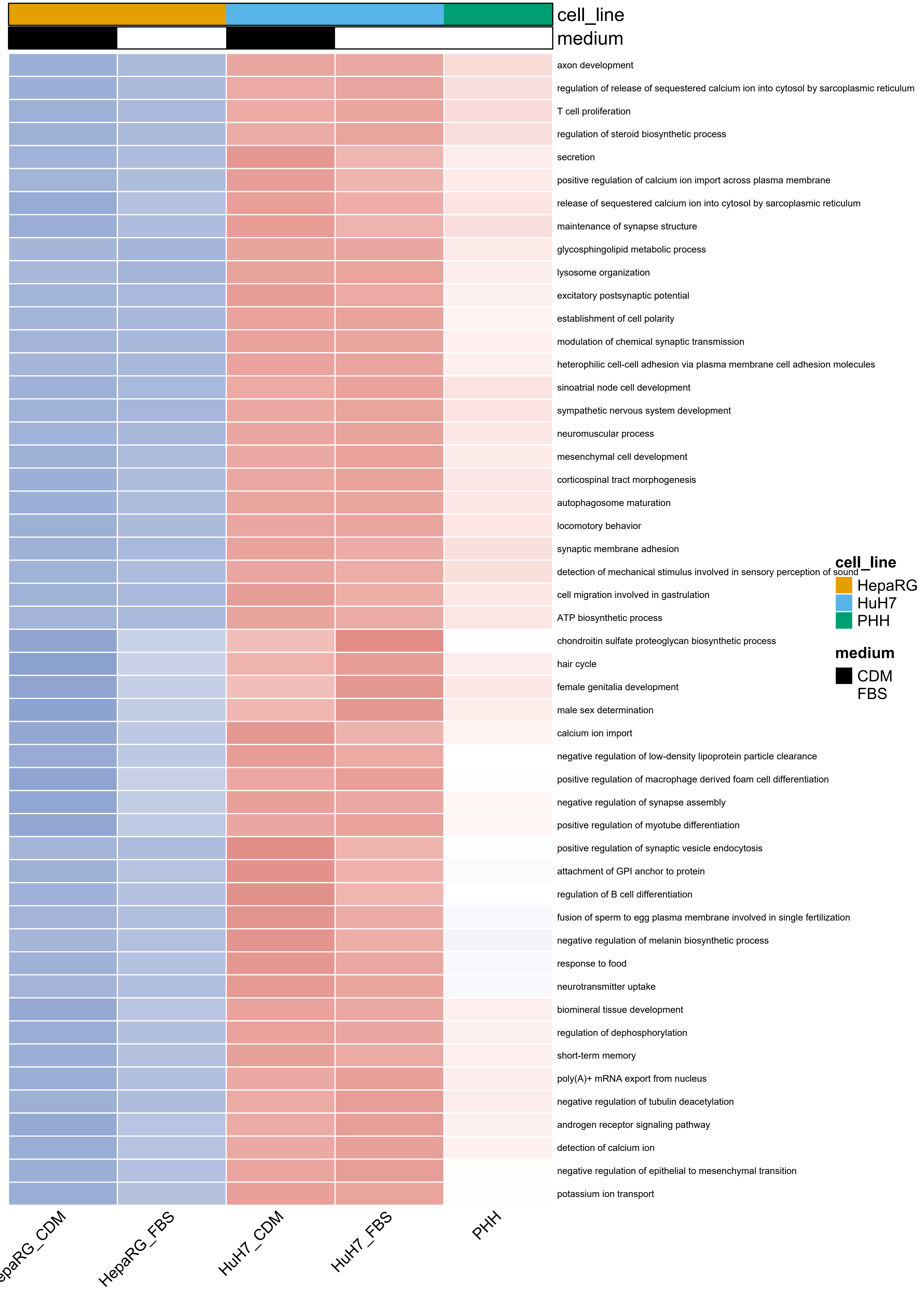
cell_line
medium
axon development
regulation of release of sequestered calcium ion into cytosol by sarcoplasmic reticulum
T cell proliferation
regulation of steroid biosynthetic process
secretion
positive regulation of calcium ion import across plasma membrane
release of sequestered calcium ion into cytosol by sarcoplasmic reticulum
maintenance of synapse structure
glycosphingolipid metabolic process
lysosome organization
excitatory postsynaptic potential
establishment of cell polarity
modulation of chemical synaptic transmission
heterophilic cell-cell adhesion via plasma membrane cell adhesion molecules
sinoatrial node cell development
sympathetic nervous system development
neuromuscular process
mesenchymal cell development
corticospinal tract morphogenesis
autophagosome maturation
locomotory behavior
synaptic membrane adhesion
detection of mechanical stimulus involved in sensory perception of sound
cell migration involved in gastrulation
ATP biosynthetic process
chondroitin sulfate proteoglycan biosynthetic process
hair cycle
female genitalia development
male sex determination
calcium ion import
negative regulation of low-density lipoprotein particle clearance
positive regulation of macrophage derived foam cell differentiation
negative regulation of synapse assembly
positive regulation of myotube differentiation
positive regulation of synaptic vesicle endocytosis
attachment of GPI anchor to protein
regulation of B cell differentiation
fusion of sperm to egg plasma membrane involved in single fertilization
negative regulation of melanin biosynthetic process
response to food
neurotransmitter uptake
biomineral tissue development
regulation of dephosphorylation
short-term memory
poly(A)+ mRNA export from nucleus
negative regulation of tubulin deacetylation
androgen receptor signaling pathway
detection of calcium ion
negative regulation of epithelial to mesenchymal transition
potassium ion transport
cell_line
HepaRG
HuH7
PHH
medium
CDM
FBS
HepaRG_CDM
HepaRG_FBS
HuH7_CDM
HuH7_FBS
PHH

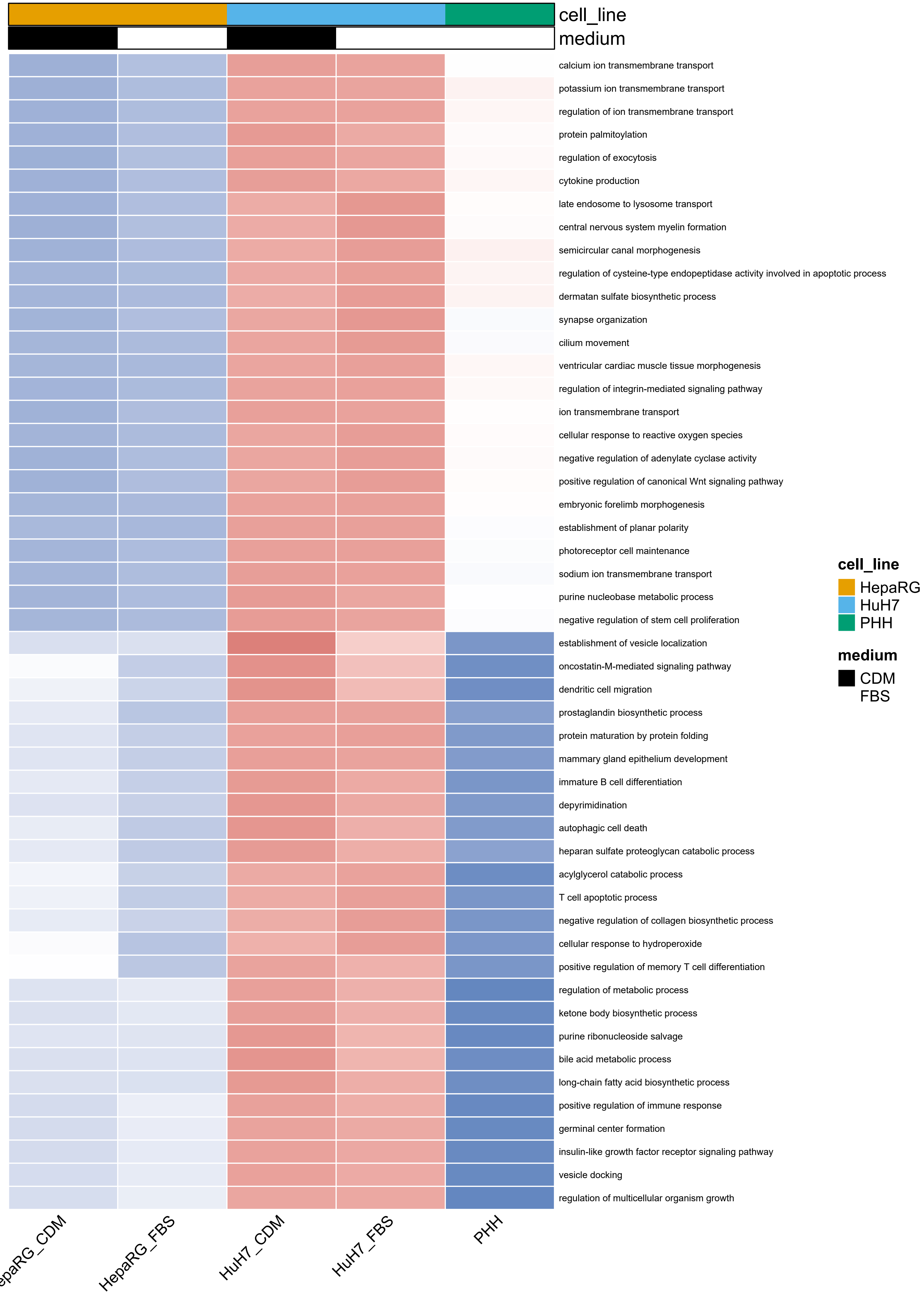

cell_line
medium
calcium ion transmembrane transport
potassium ion transmembrane transport
regulation of ion transmembrane transport
protein palmitoylation
regulation of exocytosis
cytokine production
late endosome to lysosome transport
central nervous system myelin formation
semicircular canal morphogenesis
regulation of cysteine-type endopeptidase activity involved in apoptotic process
dermatan sulfate biosynthetic process
synapse organization
cilium movement
ventricular cardiac muscle tissue morphogenesis
regulation of integrin-mediated signaling pathway
ion transmembrane transport
cellular response to reactive oxygen species
negative regulation of adenylate cyclase activity
positive regulation of canonical Wnt signaling pathway
embryonic forelimb morphogenesis
establishment of planar polarity
photoreceptor cell maintenance
sodium ion transmembrane transport
purine nucleobase metabolic process
negative regulation of stem cell proliferation
establishment of vesicle localization
oncostatin-M-mediated signaling pathway
dendritic cell migration
prostaglandin biosynthetic process
protein maturation by protein folding
mammary gland epithelium development
immature B cell differentiation
depyrimidination
autophagic cell death
heparan sulfate proteoglycan catabolic process
acylglycerol catabolic process
T cell apoptotic process
negative regulation of collagen biosynthetic process
cellular response to hydroperoxide
positive regulation of memory T cell differentiation
regulation of metabolic process
ketone body biosynthetic process
purine ribonucleoside salvage
bile acid metabolic process
long-chain fatty acid biosynthetic process
positive regulation of immune response
germinal center formation
insulin-like growth factor receptor signaling pathway
vesicle docking
regulation of multicellular organism growth
cell_line
HepaRG
HuH7
PHH
medium
CDM
FBS
HepaRG_FBS
HuH7_CDM
HuH7_FBS
PHH

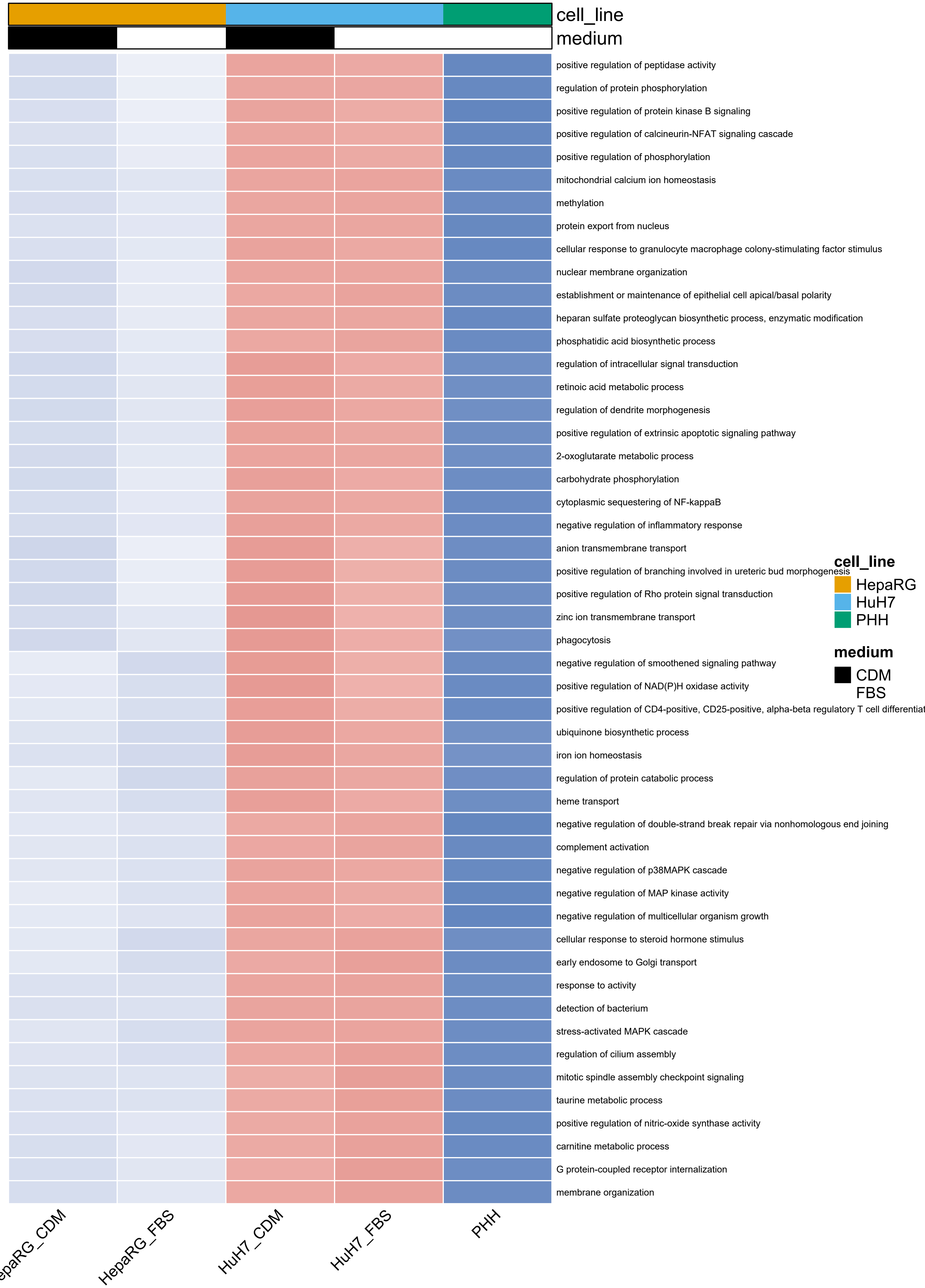
cell_line
medium
positive regulation of peptidase activity
regulation of protein phosphorylation
positive regulation of protein kinase B signaling
positive regulation of calcineurin-NFAT signaling cascade
positive regulation of phosphorylation
mitochondrial calcium ion homeostasis
methylation
protein export from nucleus
cellular response to granulocyte macrophage colony-stimulating factor stimulus
nuclear membrane organization
establishment or maintenance of epithelial cell apical/basal polarity
heparan sulfate proteoglycan biosynthetic process, enzymatic modification
phosphatidic acid biosynthetic process
regulation of intracellular signal transduction
retinoic acid metabolic process
regulation of dendrite morphogenesis
positive regulation of extrinsic apoptotic signaling pathway
2-oxoglutarate metabolic process
carbohydrate phosphorylation
cytoplasmic sequestering of NF-kappaB
negative regulation of inflammatory response
anion transmembrane transport
positive regulation of branching involved in ureteric bud morphogenesis
positive regulation of Rho protein signal transduction
zinc ion transmembrane transport
phagocytosis
negative regulation of smoothened signaling pathway
positive regulation of NAD(P)H oxidase activity
positive regulation of CD4-positive, CD25-positive, alpha-beta regulatory T cell differentiat
ubiquinone biosynthetic process
iron ion homeostasis
regulation of protein catabolic process
heme transport
negative regulation of double-strand break repair via nonhomologous end joining
complement activation
negative regulation of p38MAPK cascade
negative regulation of MAP kinase activity
negative regulation of multicellular organism growth
cellular response to steroid hormone stimulus
early endosome to Golgi transport
response to activity
detection of bacterium
stress-activated MAPK cascade
regulation of cilium assembly
mitotic spindle assembly checkpoint signaling
taurine metabolic process
positive regulation of nitric-oxide synthase activity
carnitine metabolic process
G protein-coupled receptor internalization
membrane organization
cell_line
HepaRG
HuH7
PHH
medium
CDM
FBS
epaRG_CDM
HepaRG_FBS
HuH7_CDM
HuH7_FBS
PHH

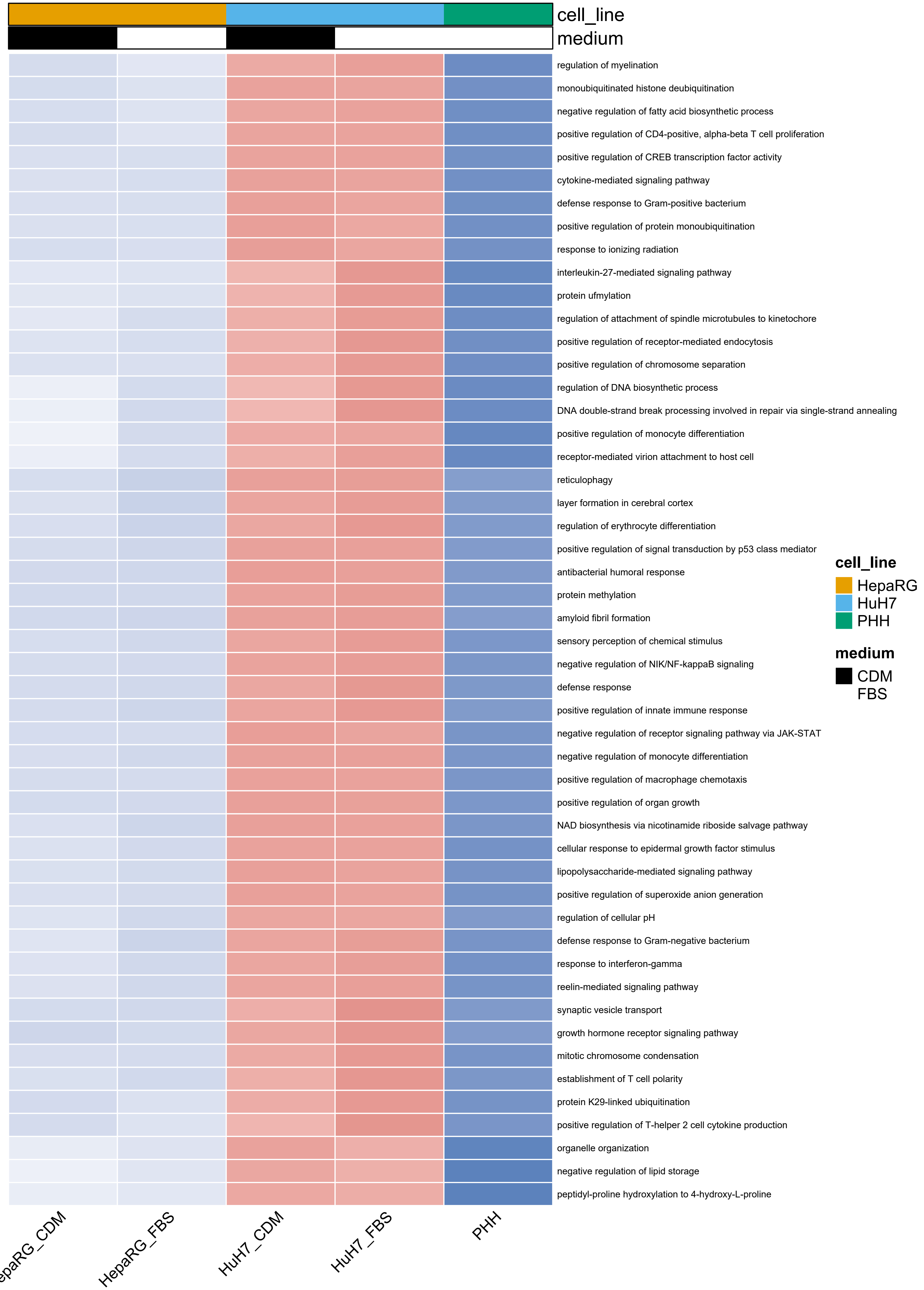
cell_line
medium
regulation of myelination
monoubiquitinated histone deubiquitination
negative regulation of fatty acid biosynthetic process
positive regulation of CD4-positive, alpha-beta T cell proliferation
positive regulation of CREB transcription factor activity
cytokine-mediated signaling pathway
defense response to Gram-positive bacterium
positive regulation of protein monoubiquitination
response to ionizing radiation
interleukin-27-mediated signaling pathway
protein ufmylation
regulation of attachment of spindle microtubules to kinetochore
positive regulation of receptor-mediated endocytosis
positive regulation of chromosome separation
regulation of DNA biosynthetic process
DNA double-strand break processing involved in repair via single-strand annealing
positive regulation of monocyte differentiation
receptor-mediated virion attachment to host cell
reticulophagy
layer formation in cerebral cortex
regulation of erythrocyte differentiation
positive regulation of signal transduction by p53 class mediator
antibacterial humoral response
protein methylation
amyloid fibril formation
sensory perception of chemical stimulus
negative regulation of NIK/NF-kappaB signaling
defense response
positive regulation of innate immune response
negative regulation of receptor signaling pathway via JAK-STAT
negative regulation of monocyte differentiation
positive regulation of macrophage chemotaxis
positive regulation of organ growth
NAD biosynthesis via nicotinamide riboside salvage pathway
cellular response to epidermal growth factor stimulus
lipopolysaccharide-mediated signaling pathway
positive regulation of superoxide anion generation
regulation of cellular pH
defense response to Gram-negative bacterium
response to interferon-gamma
reelin-mediated signaling pathway
synaptic vesicle transport
growth hormone receptor signaling pathway
mitotic chromosome condensation
establishment of T cell polarity
protein K29-linked ubiquitination
positive regulation of T-helper 2 cell cytokine production
organelle organization
negative regulation of lipid storage
peptidyl-proline hydroxylation to 4-hydroxy-L-proline
epaRG_CDM
HepaRG_FBS
HuH7_CDM
HuH7_FBS
PHH
cell_line
HepaRG
HuH7
PHH
medium
CDM
FBS

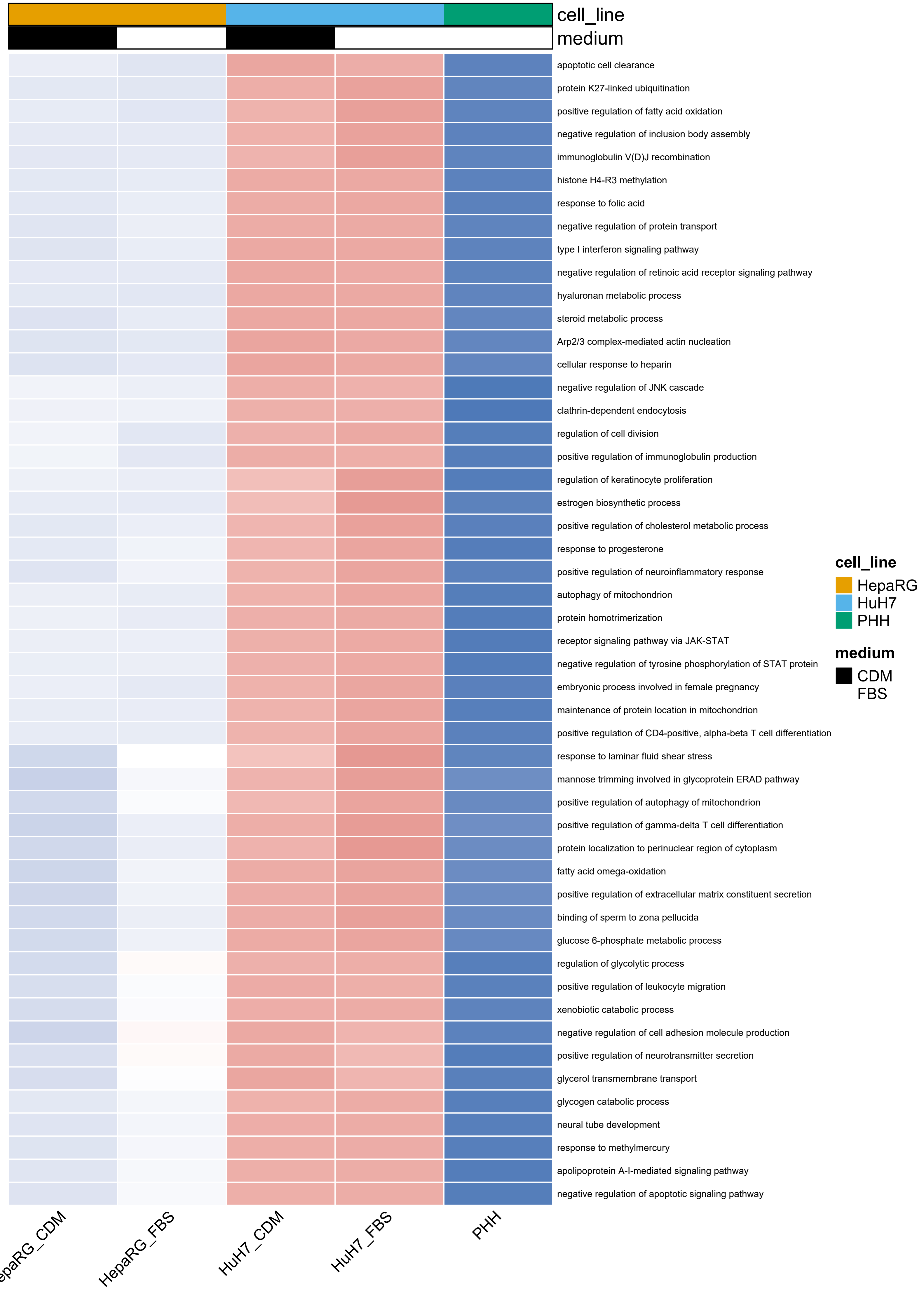

cell_line
medium
apoptotic cell clearance
protein K27-linked ubiquitination
positive regulation of fatty acid oxidation
negative regulation of inclusion body assembly
immunoglobulin V(D)J recombination
histone H4-R3 methylation
response to folic acid
negative regulation of protein transport
type I interferon signaling pathway
negative regulation of retinoic acid receptor signaling pathway
hyaluronan metabolic process
steroid metabolic process
Arp2/3 complex-mediated actin nucleation
cellular response to heparin
negative regulation of JNK cascade
clathrin-dependent endocytosis
regulation of cell division
positive regulation of immunoglobulin production
regulation of keratinocyte proliferation
estrogen biosynthetic process
positive regulation of cholesterol metabolic process
response to progesterone
positive regulation of neuroinflammatory response
autophagy of mitochondrion
protein homotrimerization
receptor signaling pathway via JAK-STAT
negative regulation of tyrosine phosphorylation of STAT protein
embryonic process involved in female pregnancy
maintenance of protein location in mitochondrion
positive regulation of CD4-positive, alpha-beta T cell differentiation
response to laminar fluid shear stress
mannose trimming involved in glycoprotein ERAD pathway
positive regulation of autophagy of mitochondrion
positive regulation of gamma-delta T cell differentiation
protein localization to perinuclear region of cytoplasm
fatty acid omega-oxidation
positive regulation of extracellular matrix constituent secretion
binding of sperm to zona pellucida
glucose 6-phosphate metabolic process
regulation of glycolytic process
positive regulation of leukocyte migration
xenobiotic catabolic process
negative regulation of cell adhesion molecule production
positive regulation of neurotransmitter secretion
glycerol transmembrane transport
glycogen catabolic process
neural tube development
response to methylmercury
apolipoprotein A-I-mediated signaling pathway
negative regulation of apoptotic signaling pathway
cell_line
HepaRG
HuH7
PHH
medium
CDM
FBS
epaRG_CDM
HepaRG_FBS
HuH7_CDM
HuH7_FBS
PHH

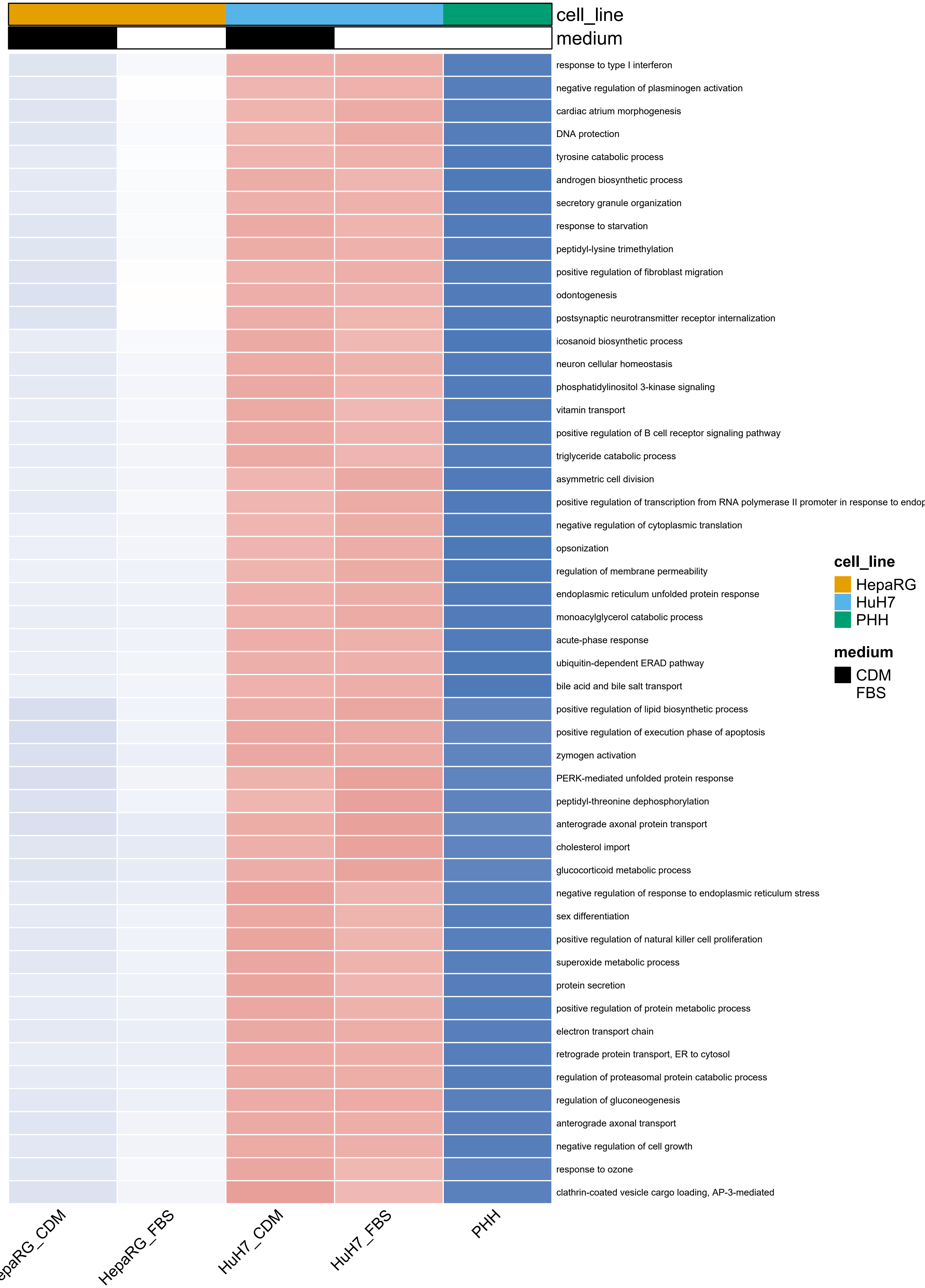
cell_line
medium
response to type I interferon
negative regulation of plasminogen activation
cardiac atrium morphogenesis
DNA protection
tyrosine catabolic process
androgen biosynthetic process
secretory granule organization
response to starvation
peptidyl-lysine trimethylation
positive regulation of fibroblast migration
odontogenesis
postsynaptic neurotransmitter receptor internalization
icosanoid biosynthetic process
neuron cellular homeostasis
phosphatidylinositol 3-kinase signaling
vitamin transport
positive regulation of B cell receptor signaling pathway
triglyceride catabolic process
asymmetric cell division
positive regulation of transcription from RNA polymerase II promoter in response to endop
negative regulation of cytoplasmic translation
opsonization
regulation of membrane permeability
endoplasmic reticulum unfolded protein response
monoacylglycerol catabolic process
acute-phase response
ubiquitin-dependent ERAD pathway
bile acid and bile salt transport
positive regulation of lipid biosynthetic process
positive regulation of execution phase of apoptosis
zymogen activation
PERK-mediated unfolded protein response
peptidyl-threonine dephosphorylation
anterograde axonal protein transport
cholesterol import
glucocorticoid metabolic process
negative regulation of response to endoplasmic reticulum stress
sex differentiation
positive regulation of natural killer cell proliferation
superoxide metabolic process
protein secretion
positive regulation of protein metabolic process
electron transport chain
retrograde protein transport, ER to cytosol
regulation of proteasomal protein catabolic process
regulation of gluconeogenesis
anterograde axonal transport
negative regulation of cell growth
response to ozone
clathrin-coated vesicle cargo loading, AP-3-mediated
cell_line
HepaRG
HuH7
PHH
medium
CDM
FBS
epaRG_CDM
HepaRG_FBS
HuH7_CDM
HuH7_FBS
PHH

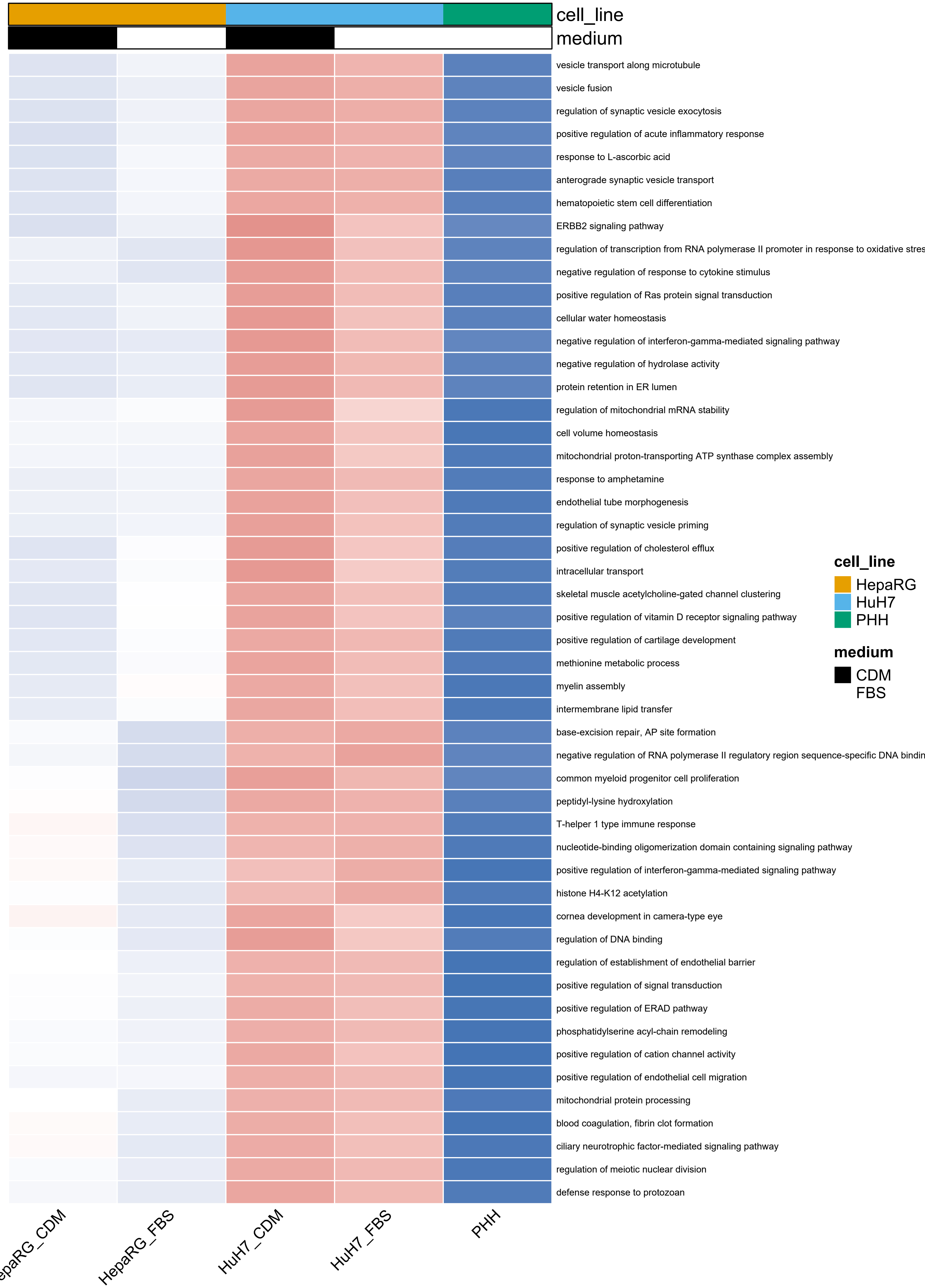

cell_line
medium
vesicle transport along microtubule
vesicle fusion
regulation of synaptic vesicle exocytosis
positive regulation of acute inflammatory response
response to L-ascorbic acid
anterograde synaptic vesicle transport
hematopoietic stem cell differentiation
ERBB2 signaling pathway
regulation of transcription from RNA polymerase II promoter in response to oxidative stres
negative regulation of response to cytokine stimulus
positive regulation of Ras protein signal transduction
cellular water homeostasis
negative regulation of interferon-gamma-mediated signaling pathway
negative regulation of hydrolase activity
protein retention in ER lumen
regulation of mitochondrial mRNA stability
cell volume homeostasis
mitochondrial proton-transporting ATP synthase complex assembly
response to amphetamine
endothelial tube morphogenesis
regulation of synaptic vesicle priming
positive regulation of cholesterol efflux
intracellular transport
skeletal muscle acetylcholine-gated channel clustering
positive regulation of vitamin D receptor signaling pathway
positive regulation of cartilage development
methionine metabolic process
myelin assembly
intermembrane lipid transfer
base-excision repair, AP site formation
negative regulation of RNA polymerase II regulatory region sequence-specific DNA bindin
common myeloid progenitor cell proliferation
peptidyl-lysine hydroxylation
T-helper 1 type immune response
nucleotide-binding oligomerization domain containing signaling pathway
positive regulation of interferon-gamma-mediated signaling pathway
histone H4-K12 acetylation
cornea development in camera-type eye
regulation of DNA binding
regulation of establishment of endothelial barrier
positive regulation of signal transduction
positive regulation of ERAD pathway
phosphatidylserine acyl-chain remodeling
positive regulation of cation channel activity
positive regulation of endothelial cell migration
mitochondrial protein processing
blood coagulation, fibrin clot formation
ciliary neurotrophic factor-mediated signaling pathway
regulation of meiotic nuclear division
defense response to protozoan
cell_line
HepaRG
HuH7
PHH
medium
CDM
FBS
epaRG_CDM
HepaRG_FBS
HuH7_CDM
HuH7_FBS
PHH

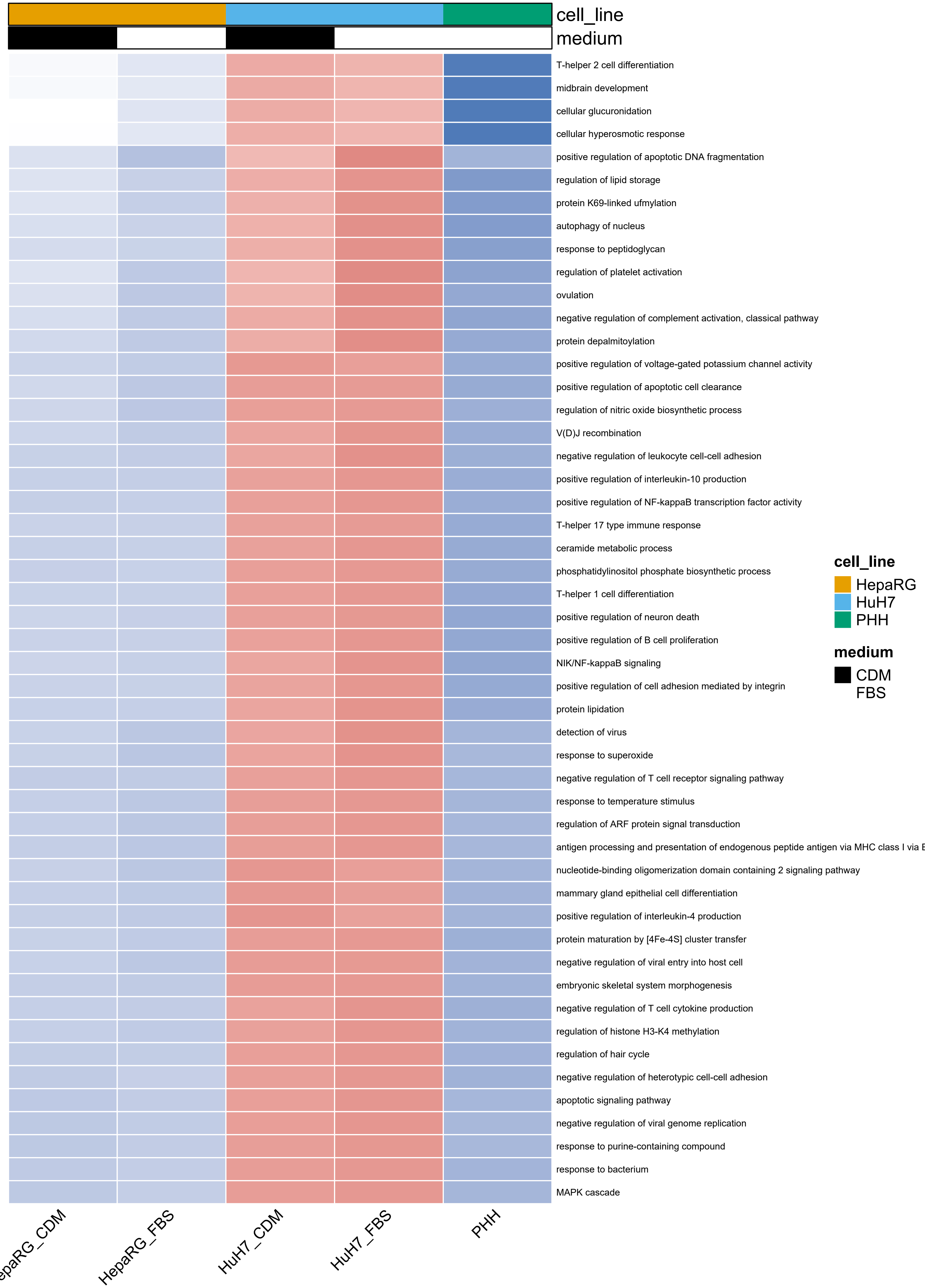
cell_line
medium
T-helper 2 cell differentiation
midbrain development
cellular glucuronidation
cellular hyperosmotic response
positive regulation of apoptotic DNA fragmentation
regulation of lipid storage
protein K69-linked ufmylation
autophagy of nucleus
response to peptidoglycan
regulation of platelet activation
ovulation
negative regulation of complement activation, classical pathway
protein depalmitoylation
positive regulation of voltage-gated potassium channel activity
positive regulation of apoptotic cell clearance
regulation of nitric oxide biosynthetic process
V(D)J recombination
negative regulation of leukocyte cell-cell adhesion
positive regulation of interleukin-10 production
positive regulation of NF-kappaB transcription factor activity
T-helper 17 type immune response
ceramide metabolic process
phosphatidylinositol phosphate biosynthetic process
T-helper 1 cell differentiation
positive regulation of neuron death
positive regulation of B cell proliferation
NIK/NF-kappaB signaling
positive regulation of cell adhesion mediated by integrin
protein lipidation
detection of virus
response to superoxide
negative regulation of T cell receptor signaling pathway
response to temperature stimulus
regulation of ARF protein signal transduction
antigen processing and presentation of endogenous peptide antigen via MHC class I via E
nucleotide-binding oligomerization domain containing 2 signaling pathway
mammary gland epithelial cell differentiation
positive regulation of interleukin-4 production
protein maturation by [4Fe-4S] cluster transfer
negative regulation of viral entry into host cell
embryonic skeletal system morphogenesis
negative regulation of T cell cytokine production
regulation of histone H3-K4 methylation
regulation of hair cycle
negative regulation of heterotypic cell-cell adhesion
apoptotic signaling pathway
negative regulation of viral genome replication
response to purine-containing compound
response to bacterium
MAPK cascade
cell_line
HepaRG
HuH7
PHH
medium
CDM
FBS
epaRG_CDM
HepaRG_FBS
HuH7_CDM
HuH7_FBS
PHH

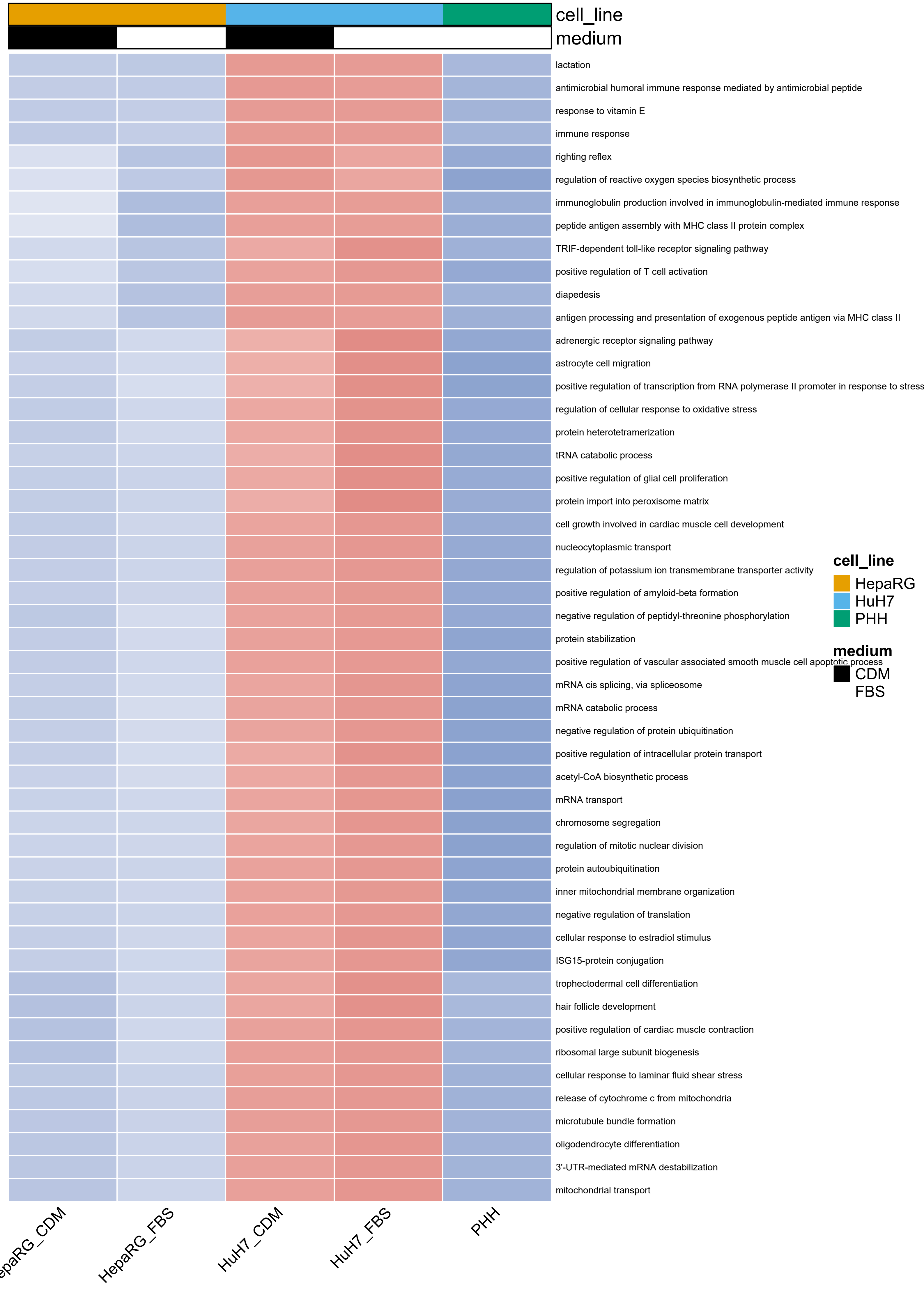
cell_line
medium
lactation
antimicrobial humoral immune response mediated by antimicrobial peptide
response to vitamin E
immune response
righting reflex
regulation of reactive oxygen species biosynthetic process
immunoglobulin production involved in immunoglobulin-mediated immune response
peptide antigen assembly with MHC class II protein complex
TRIF-dependent toll-like receptor signaling pathway
positive regulation of T cell activation
diapedesis
antigen processing and presentation of exogenous peptide antigen via MHC class II
adrenergic receptor signaling pathway
astrocyte cell migration
positive regulation of transcription from RNA polymerase II promoter in response to stress
regulation of cellular response to oxidative stress
protein heterotetramerization
tRNA catabolic process
positive regulation of glial cell proliferation
protein import into peroxisome matrix
cell growth involved in cardiac muscle cell development
nucleocytoplasmic transport
regulation of potassium ion transmembrane transporter activity
positive regulation of amyloid-beta formation
negative regulation of peptidyl-threonine phosphorylation
protein stabilization
positive regulation of vascular associated smooth muscle cell apoptotic process
mRNA cis splicing, via spliceosome
mRNA catabolic process
negative regulation of protein ubiquitination
positive regulation of intracellular protein transport
acetyl-CoA biosynthetic process
mRNA transport
chromosome segregation
regulation of mitotic nuclear division
protein autoubiquitination
inner mitochondrial membrane organization
negative regulation of translation
cellular response to estradiol stimulus
ISG15-protein conjugation
trophectodermal cell differentiation
hair follicle development
positive regulation of cardiac muscle contraction
ribosomal large subunit biogenesis
cellular response to laminar fluid shear stress
release of cytochrome c from mitochondria
microtubule bundle formation
oligodendrocyte differentiation
3'-UTR-mediated mRNA destabilization
mitochondrial transport
epaRG_CDM
HepaRG_FBS
HuH7_CDM
HuH7_FBS
PHH
cell_line
HepaRG
HuH7
PHH
medium
CDM
FBS

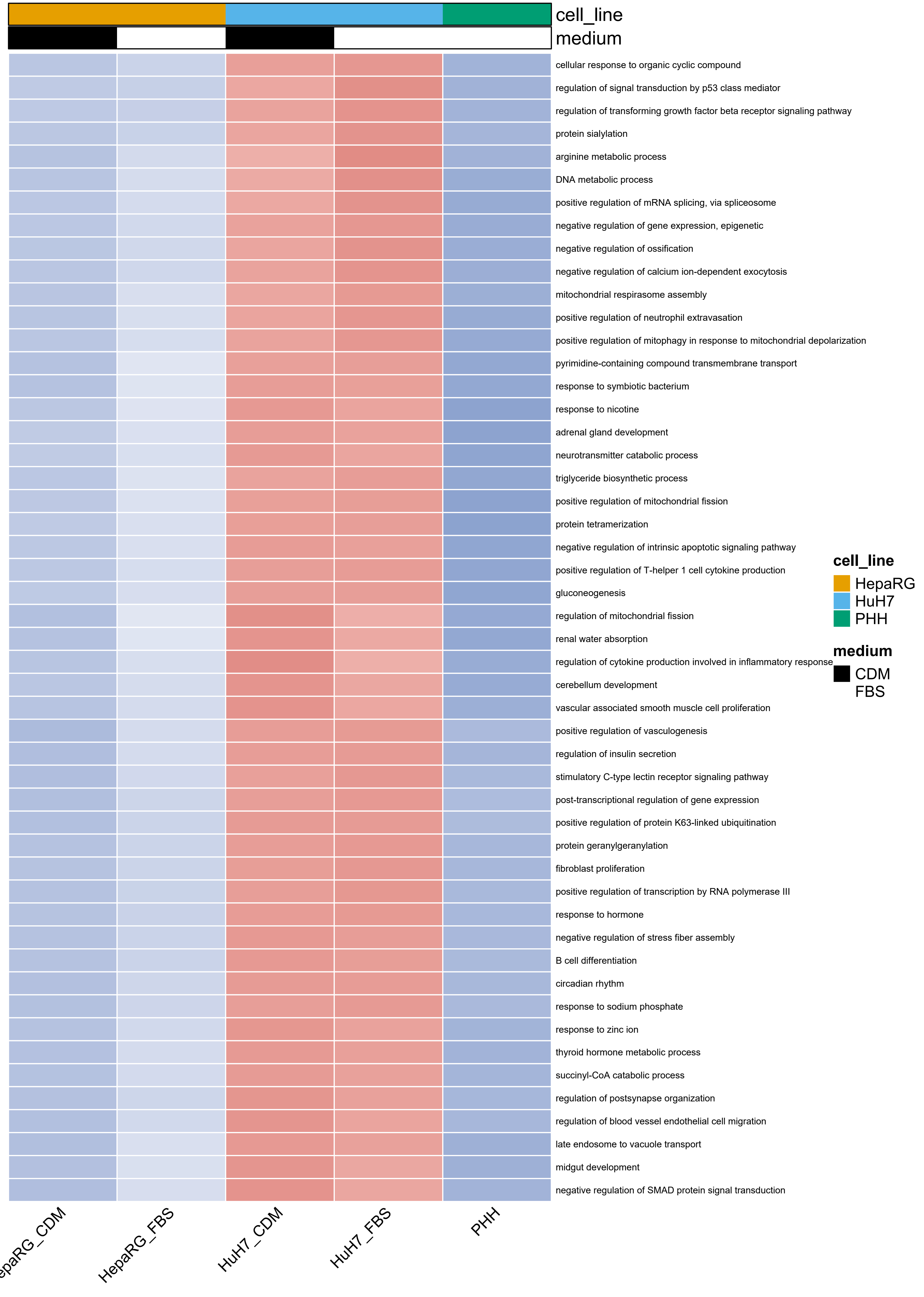
cell_line
medium
cellular response to organic cyclic compound
regulation of signal transduction by p53 class mediator
regulation of transforming growth factor beta receptor signaling pathway
protein sialylation
arginine metabolic process
DNA metabolic process
positive regulation of mRNA splicing, via spliceosome
negative regulation of gene expression, epigenetic
negative regulation of ossification
negative regulation of calcium ion-dependent exocytosis
mitochondrial respirasome assembly
positive regulation of neutrophil extravasation
positive regulation of mitophagy in response to mitochondrial depolarization
pyrimidine-containing compound transmembrane transport
response to symbiotic bacterium
response to nicotine
adrenal gland development
neurotransmitter catabolic process
triglyceride biosynthetic process
positive regulation of mitochondrial fission
protein tetramerization
negative regulation of intrinsic apoptotic signaling pathway
positive regulation of T-helper 1 cell cytokine production
gluconeogenesis
regulation of mitochondrial fission
renal water absorption
regulation of cytokine production involved in inflammatory response
cerebellum development
vascular associated smooth muscle cell proliferation
positive regulation of vasculogenesis
regulation of insulin secretion
stimulatory C-type lectin receptor signaling pathway
post-transcriptional regulation of gene expression
positive regulation of protein K63-linked ubiquitination
protein geranylgeranylation
fibroblast proliferation
positive regulation of transcription by RNA polymerase III
response to hormone
negative regulation of stress fiber assembly
B cell differentiation
circadian rhythm
response to sodium phosphate
response to zinc ion
thyroid hormone metabolic process
succinyl-CoA catabolic process
regulation of postsynapse organization
regulation of blood vessel endothelial cell migration
late endosome to vacuole transport
midgut development
negative regulation of SMAD protein signal transduction
cell_line
HepaRG
HuH7
PHH
medium
CDM
FBS
HepaRG_FBS
HuH7_CDM
HuH7_FBS
PHH

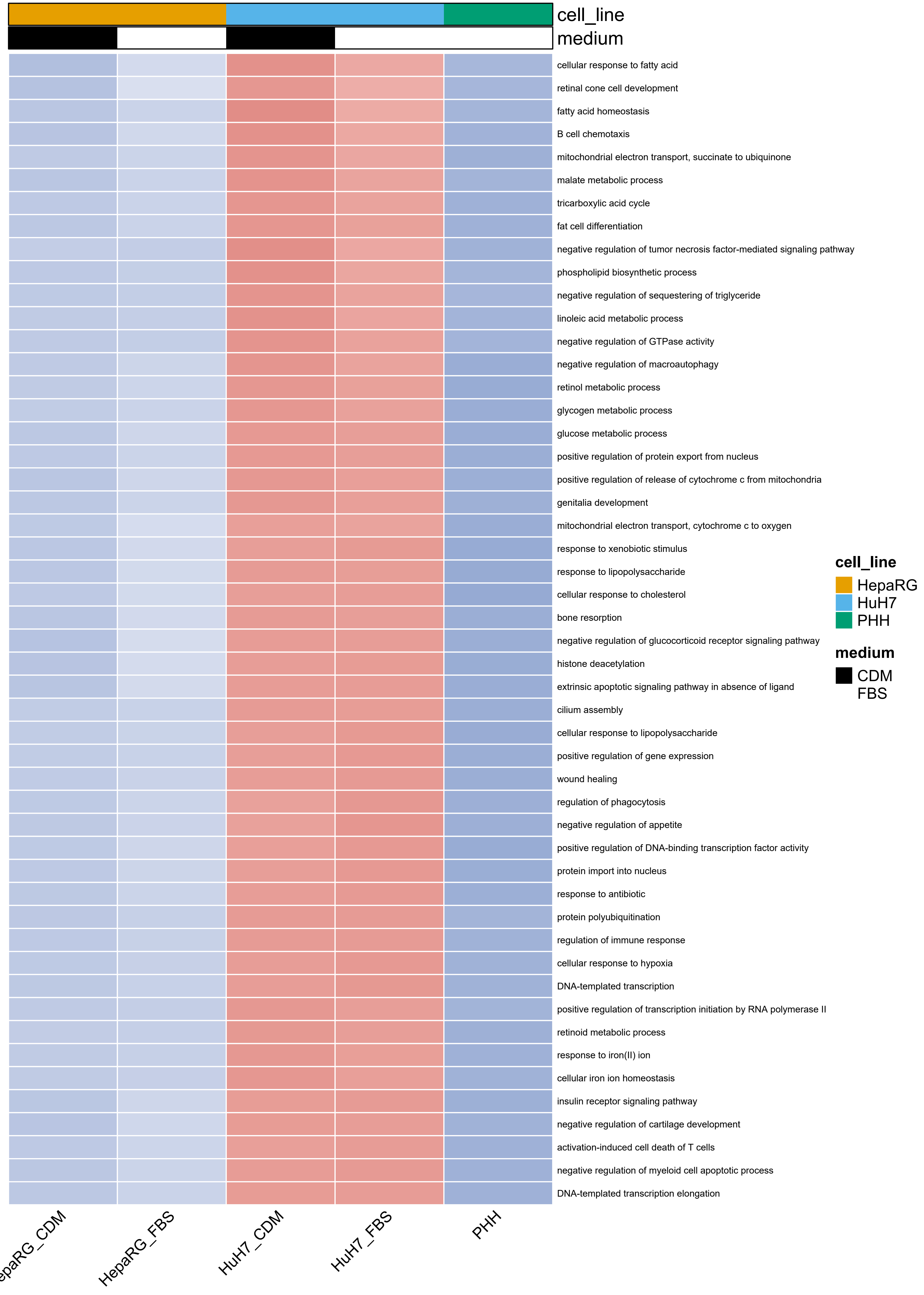
cell_line
medium
cellular response to fatty acid
retinal cone cell development
fatty acid homeostasis
B cell chemotaxis
mitochondrial electron transport, succinate to ubiquinone
malate metabolic process
tricarboxylic acid cycle
fat cell differentiation
negative regulation of tumor necrosis factor-mediated signaling pathway
phospholipid biosynthetic process
negative regulation of sequestering of triglyceride
linoleic acid metabolic process
negative regulation of GTPase activity
negative regulation of macroautophagy
retinol metabolic process
glycogen metabolic process
glucose metabolic process
positive regulation of protein export from nucleus
positive regulation of release of cytochrome c from mitochondria
genitalia development
mitochondrial electron transport, cytochrome c to oxygen
response to xenobiotic stimulus
response to lipopolysaccharide
cellular response to cholesterol
bone resorption
negative regulation of glucocorticoid receptor signaling pathway
histone deacetylation
extrinsic apoptotic signaling pathway in absence of ligand
cilium assembly
cellular response to lipopolysaccharide
positive regulation of gene expression
wound healing
regulation of phagocytosis
negative regulation of appetite
positive regulation of DNA-binding transcription factor activity
protein import into nucleus
response to antibiotic
protein polyubiquitination
regulation of immune response
cellular response to hypoxia
DNA-templated transcription
positive regulation of transcription initiation by RNA polymerase II
retinoid metabolic process
response to iron(II) ion
cellular iron ion homeostasis
insulin receptor signaling pathway
negative regulation of cartilage development
activation-induced cell death of T cells
negative regulation of myeloid cell apoptotic process
DNA-templated transcription elongation
cell_line
HepaRG
HuH7
PHH
medium
CDM
FBS
epaRG_CDM
HepaRG_FBS
HuH7_CDM
HuH7_FBS
PHH

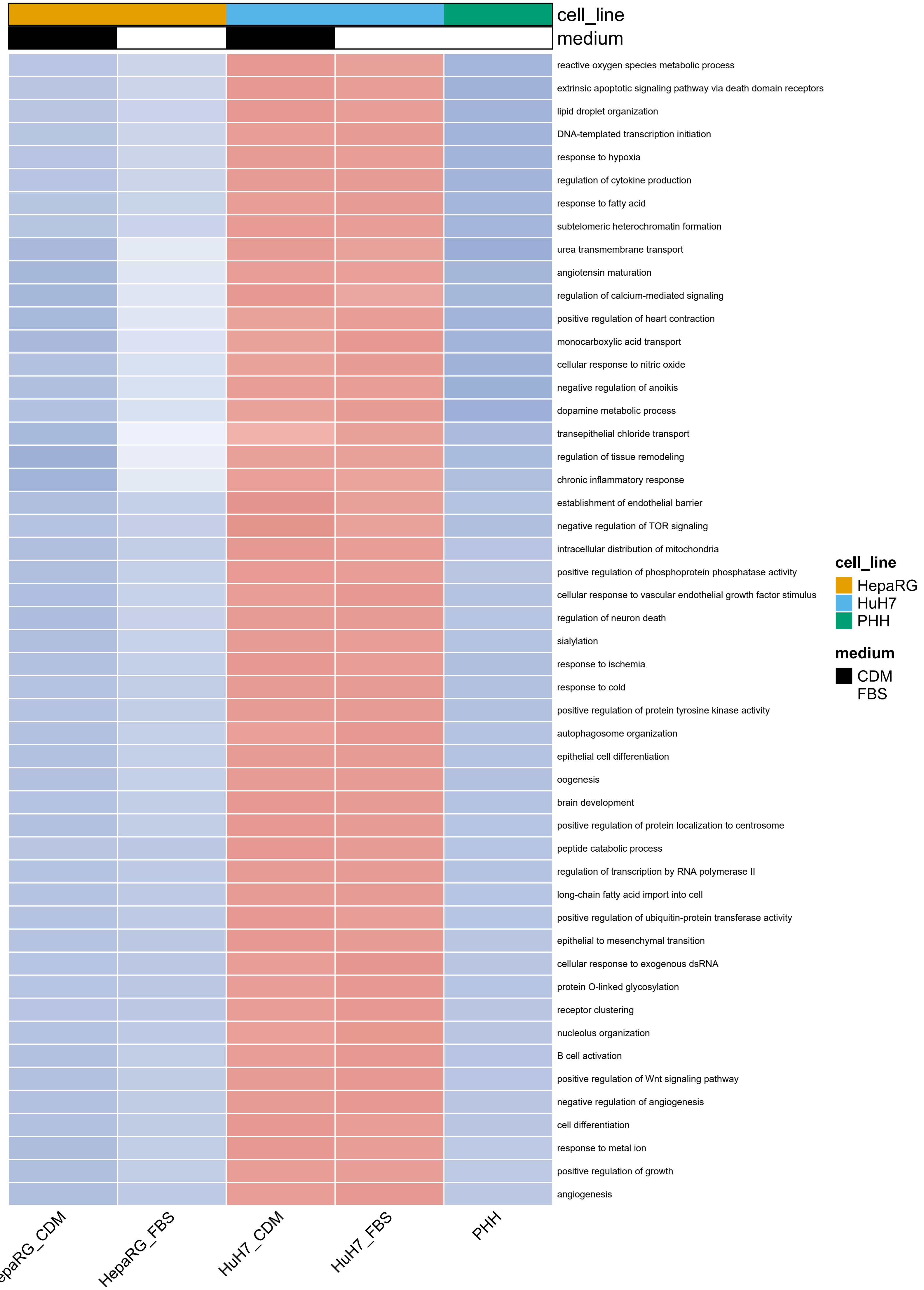

cell_line
medium
reactive oxygen species metabolic process
extrinsic apoptotic signaling pathway via death domain receptors
lipid droplet organization
DNA-templated transcription initiation
response to hypoxia
regulation of cytokine production
response to fatty acid
subtelomeric heterochromatin formation
urea transmembrane transport
angiotensin maturation
regulation of calcium-mediated signaling
positive regulation of heart contraction
monocarboxylic acid transport
cellular response to nitric oxide
negative regulation of anoikis
dopamine metabolic process
transepithelial chloride transport
regulation of tissue remodeling
chronic inflammatory response
establishment of endothelial barrier
negative regulation of TOR signaling
intracellular distribution of mitochondria
positive regulation of phosphoprotein phosphatase activity
cellular response to vascular endothelial growth factor stimulus
regulation of neuron death
sialylation
response to ischemia
response to cold
positive regulation of protein tyrosine kinase activity
autophagosome organization
epithelial cell differentiation
oogenesis
brain development
positive regulation of protein localization to centrosome
peptide catabolic process
regulation of transcription by RNA polymerase II
long-chain fatty acid import into cell
positive regulation of ubiquitin-protein transferase activity
epithelial to mesenchymal transition
cellular response to exogenous dsRNA
protein O-linked glycosylation
receptor clustering
nucleolus organization
B cell activation
positive regulation of Wnt signaling pathway
negative regulation of angiogenesis
cell differentiation
response to metal ion
positive regulation of growth
angiogenesis
cell_line
HepaRG
HuH7
PHH
medium
CDM
FBS
HepaRG_CDM
HepaRG_FBS
HuH7_CDM
HuH7_FBS
PHH

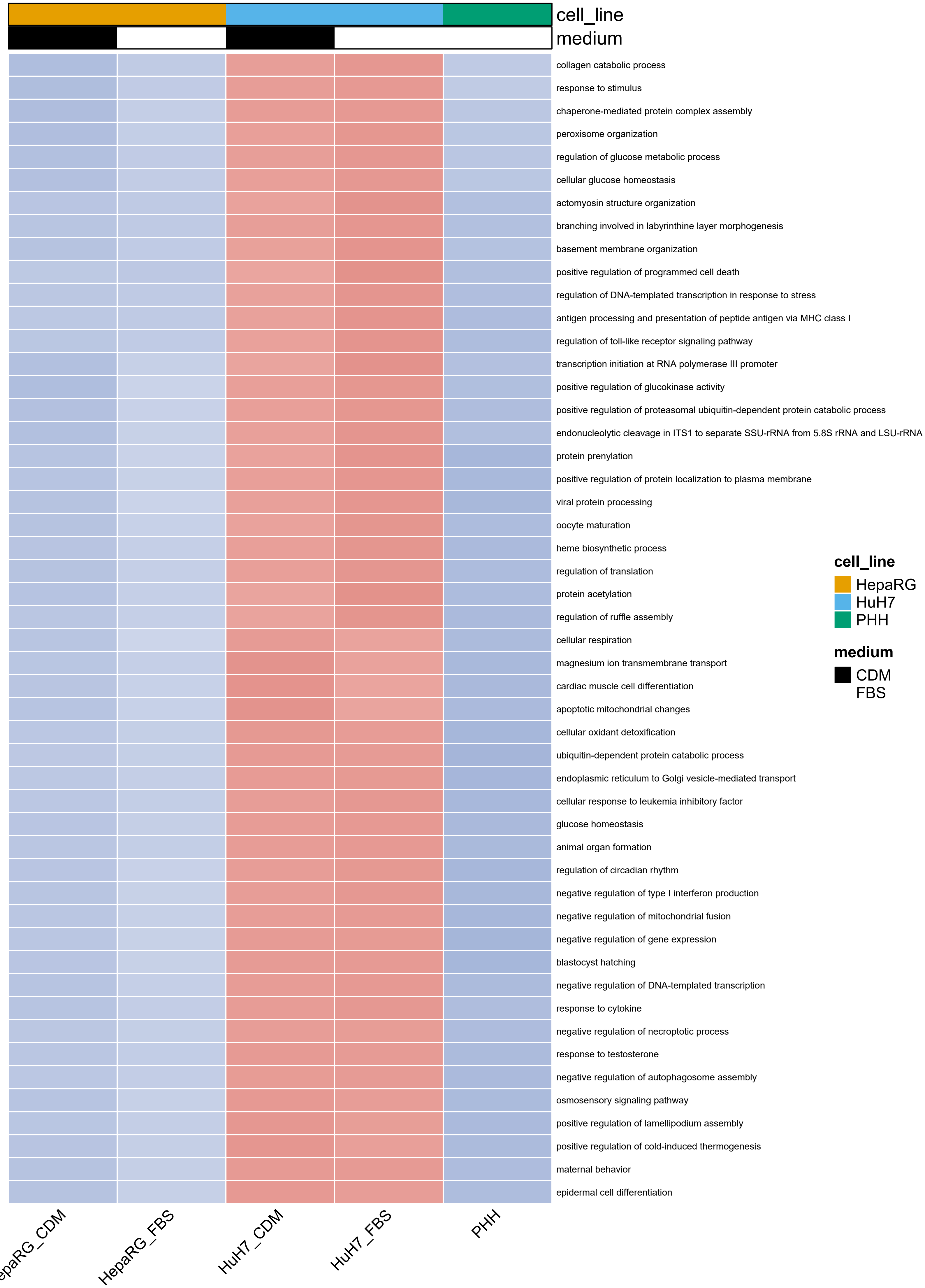
cell_line
medium
collagen catabolic process
response to stimulus
chaperone-mediated protein complex assembly
peroxisome organization
regulation of glucose metabolic process
cellular glucose homeostasis
actomyosin structure organization
branching involved in labyrinthine layer morphogenesis
basement membrane organization
positive regulation of programmed cell death
regulation of DNA-templated transcription in response to stress
antigen processing and presentation of peptide antigen via MHC class I
regulation of toll-like receptor signaling pathway
transcription initiation at RNA polymerase III promoter
positive regulation of glucokinase activity
positive regulation of proteasomal ubiquitin-dependent protein catabolic process
endonucleolytic cleavage in ITS1 to separate SSU-rRNA from 5.8S rRNA and LSU-rRNA
protein prenylation
positive regulation of protein localization to plasma membrane
viral protein processing
oocyte maturation
heme biosynthetic process
regulation of translation
protein acetylation
regulation of ruffle assembly
cellular respiration
magnesium ion transmembrane transport
cardiac muscle cell differentiation
apoptotic mitochondrial changes
cellular oxidant detoxification
ubiquitin-dependent protein catabolic process
endoplasmic reticulum to Golgi vesicle-mediated transport
cellular response to leukemia inhibitory factor
glucose homeostasis
animal organ formation
regulation of circadian rhythm
negative regulation of type I interferon production
negative regulation of mitochondrial fusion
negative regulation of gene expression
blastocyst hatching
negative regulation of DNA-templated transcription
response to cytokine
negative regulation of necroptotic process
response to testosterone
negative regulation of autophagosome assembly
osmosensory signaling pathway
positive regulation of lamellipodium assembly
positive regulation of cold-induced thermogenesis
maternal behavior
epidermal cell differentiation
cell_line
HepaRG
HuH7
PHH
medium
CDM
FBS
HepaRG_CDM
HepaRG_FBS
HuH7_CDM
HuH7_FBS
PHH

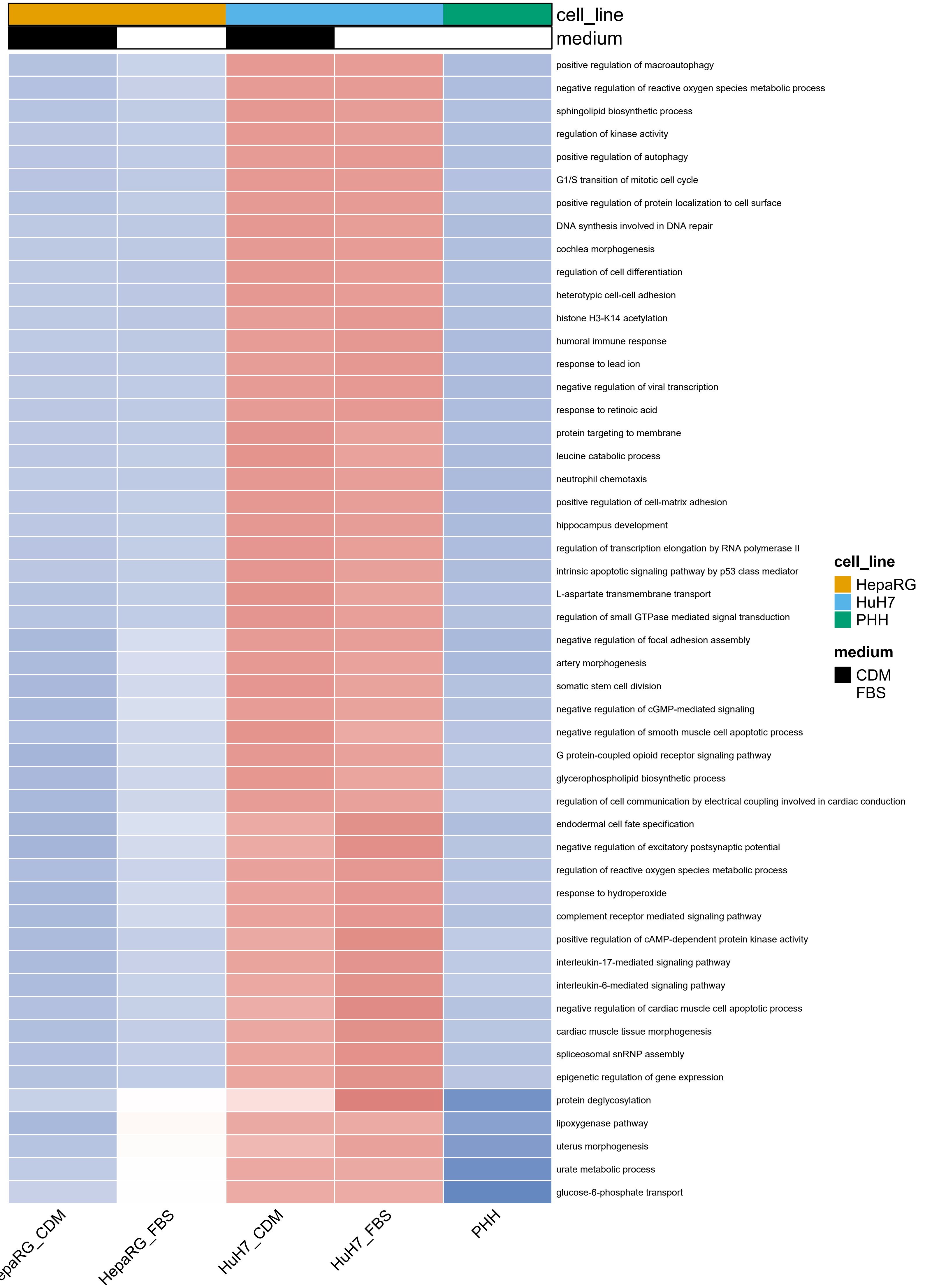
cell_line
medium
positive regulation of macroautophagy
negative regulation of reactive oxygen species metabolic process
sphingolipid biosynthetic process
regulation of kinase activity
positive regulation of autophagy
G1/S transition of mitotic cell cycle
positive regulation of protein localization to cell surface
DNA synthesis involved in DNA repair
cochlea morphogenesis
regulation of cell differentiation
heterotypic cell-cell adhesion
histone H3-K14 acetylation
humoral immune response
response to lead ion
negative regulation of viral transcription
response to retinoic acid
protein targeting to membrane
leucine catabolic process
neutrophil chemotaxis
positive regulation of cell-matrix adhesion
hippocampus development
regulation of transcription elongation by RNA polymerase II
intrinsic apoptotic signaling pathway by p53 class mediator
L-aspartate transmembrane transport
regulation of small GTPase mediated signal transduction
negative regulation of focal adhesion assembly
artery morphogenesis
somatic stem cell division
negative regulation of cGMP-mediated signaling
negative regulation of smooth muscle cell apoptotic process
G protein-coupled opioid receptor signaling pathway
glycerophospholipid biosynthetic process
regulation of cell communication by electrical coupling involved in cardiac conduction
endodermal cell fate specification
negative regulation of excitatory postsynaptic potential
regulation of reactive oxygen species metabolic process
response to hydroperoxide
complement receptor mediated signaling pathway
positive regulation of cAMP-dependent protein kinase activity
interleukin-17-mediated signaling pathway
interleukin-6-mediated signaling pathway
negative regulation of cardiac muscle cell apoptotic process
cardiac muscle tissue morphogenesis
spliceosomal snRNP assembly
epigenetic regulation of gene expression
protein deglycosylation
lipoxygenase pathway
uterus morphogenesis
urate metabolic process
glucose-6-phosphate transport
cell_line
HepaRG
HuH7
PHH
medium
CDM
FBS
HepaRG_FBS
HuH7_CDM
HuH7_FBS
PHH

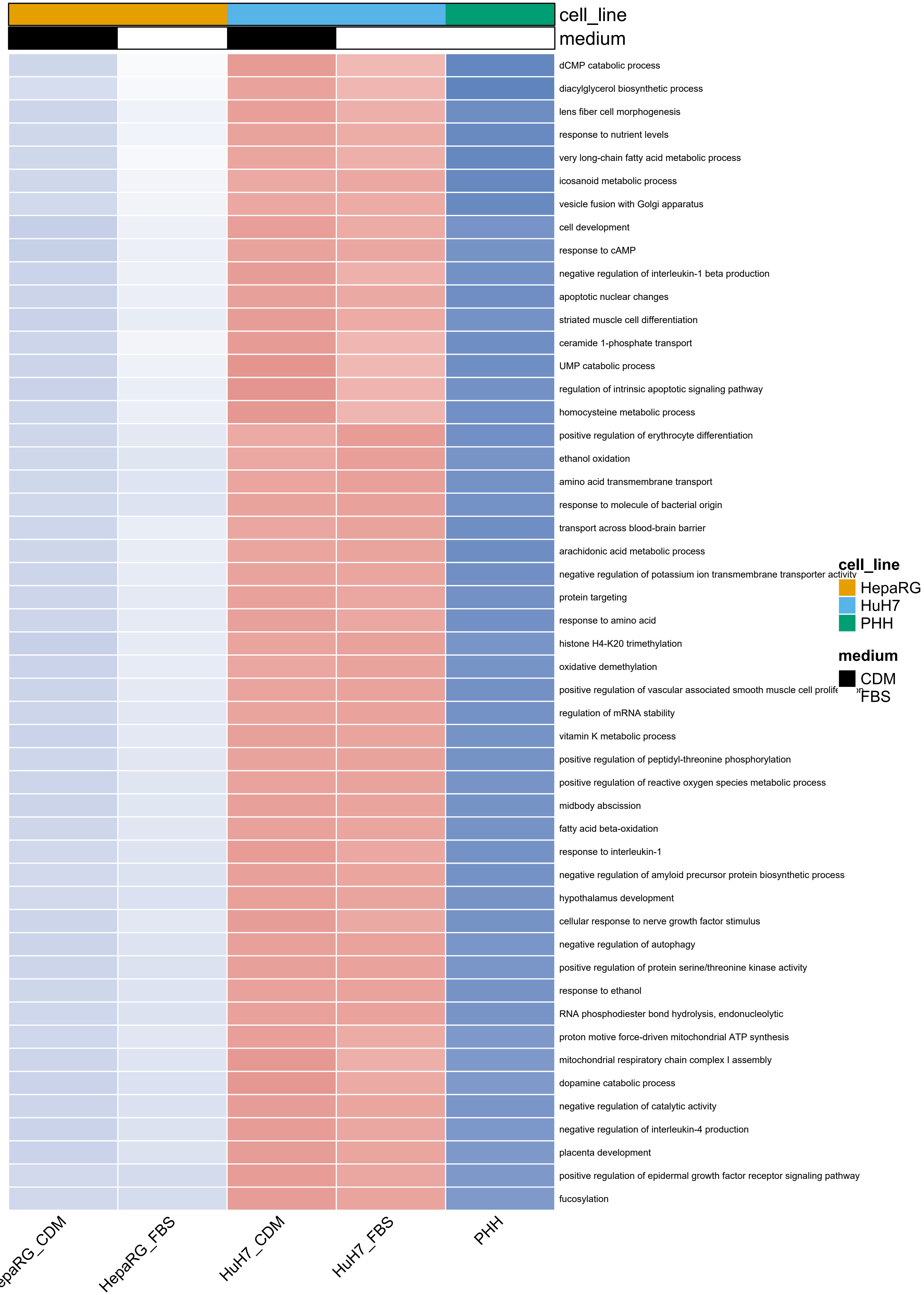

cell_line
medium
dCMP catabolic process
diacylglycerol biosynthetic process
lens fiber cell morphogenesis
response to nutrient levels
very long-chain fatty acid metabolic process
icosanoid metabolic process
vesicle fusion with Golgi apparatus
cell development
response to cAMP
negative regulation of interleukin-1 beta production
apoptotic nuclear changes
striated muscle cell differentiation
ceramide 1-phosphate transport
UMP catabolic process
regulation of intrinsic apoptotic signaling pathway
homocysteine metabolic process
positive regulation of erythrocyte differentiation
ethanol oxidation
amino acid transmembrane transport
response to molecule of bacterial origin
transport across blood-brain barrier
arachidonic acid metabolic process
negative regulation of potassium ion transmembrane transporter activity
protein targeting
response to amino acid
histone H4-K20 trimethylation
oxidative demethylation
positive regulation of vascular associated smooth muscle cell prolife
regulation of mRNA stability
vitamin K metabolic process
positive regulation of peptidyl-threonine phosphorylation
positive regulation of reactive oxygen species metabolic process
midbody abscission
fatty acid beta-oxidation
response to interleukin-1
negative regulation of amyloid precursor protein biosynthetic process
hypothalamus development
cellular response to nerve growth factor stimulus
negative regulation of autophagy
positive regulation of protein serine/threonine kinase activity
response to ethanol
RNA phosphodiester bond hydrolysis, endonucleolytic
proton motive force-driven mitochondrial ATP synthesis
mitochondrial respiratory chain complex I assembly
dopamine catabolic process
negative regulation of catalytic activity
negative regulation of interleukin-4 production
placenta development
positive regulation of epidermal growth factor receptor signaling pathway
fucosylation
cell_line
HepaRG
HuH7
PHH
medium
CDM
FBS
HepaRG_FBS
HuH7_CDM
HuH7_FBS
PHH

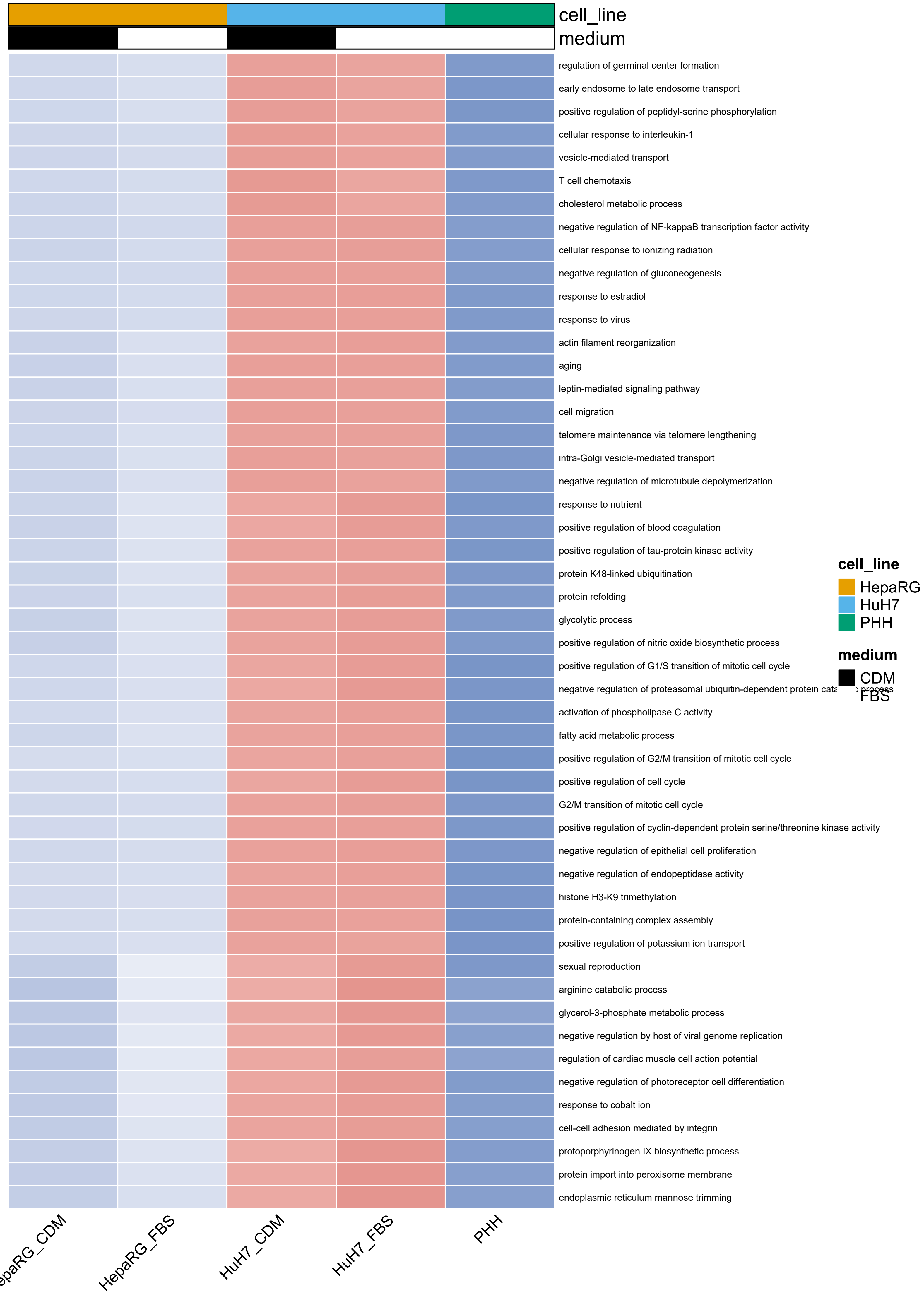
cell_line
medium
regulation of germinal center formation
early endosome to late endosome transport
positive regulation of peptidyl-serine phosphorylation
cellular response to interleukin-1
vesicle-mediated transport
T cell chemotaxis
cholesterol metabolic process
negative regulation of NF-kappaB transcription factor activity
cellular response to ionizing radiation
negative regulation of gluconeogenesis
response to estradiol
response to virus
actin filament reorganization
aging
leptin-mediated signaling pathway
cell migration
telomere maintenance via telomere lengthening
intra-Golgi vesicle-mediated transport
negative regulation of microtubule depolymerization
response to nutrient
positive regulation of blood coagulation
positive regulation of tau-protein kinase activity
protein K48-linked ubiquitination
protein refolding
glycolytic process
positive regulation of nitric oxide biosynthetic process
positive regulation of G1/S transition of mitotic cell cycle
negative regulation of proteasomal ubiquitin-dependent protein cata
activation of phospholipase C activity
fatty acid metabolic process
positive regulation of G2/M transition of mitotic cell cycle
positive regulation of cell cycle
G2/M transition of mitotic cell cycle
positive regulation of cyclin-dependent protein serine/threonine kinase activity
negative regulation of epithelial cell proliferation
negative regulation of endopeptidase activity
histone H3-K9 trimethylation
protein-containing complex assembly
positive regulation of potassium ion transport
sexual reproduction
arginine catabolic process
glycerol-3-phosphate metabolic process
negative regulation by host of viral genome replication
regulation of cardiac muscle cell action potential
negative regulation of photoreceptor cell differentiation
response to cobalt ion
cell-cell adhesion mediated by integrin
protoporphyrinogen IX biosynthetic process
protein import into peroxisome membrane
endoplasmic reticulum mannose trimming
cell_line
HepaRG
HuH7
PHH
medium
CDM
FBS
HepaRG_FBS
HuH7_CDM
HuH7_FBS
PHH

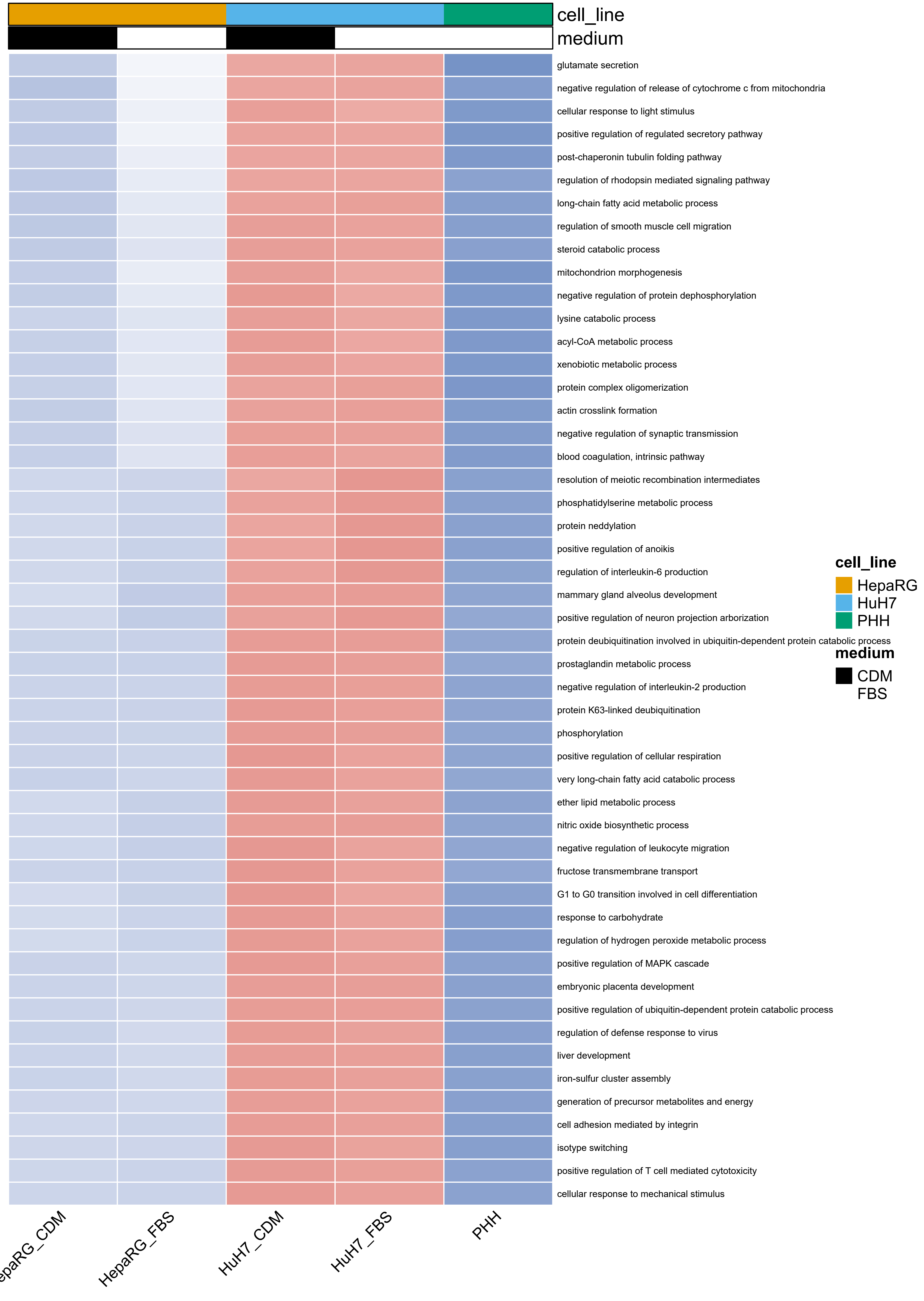

cell_line
medium
glutamate secretion
negative regulation of release of cytochrome c from mitochondria
cellular response to light stimulus
positive regulation of regulated secretory pathway
post-chaperonin tubulin folding pathway
regulation of rhodopsin mediated signaling pathway
long-chain fatty acid metabolic process
regulation of smooth muscle cell migration
steroid catabolic process
mitochondrion morphogenesis
negative regulation of protein dephosphorylation
lysine catabolic process
acyl-CoA metabolic process
xenobiotic metabolic process
protein complex oligomerization
actin crosslink formation
negative regulation of synaptic transmission
blood coagulation, intrinsic pathway
resolution of meiotic recombination intermediates
phosphatidylserine metabolic process
protein neddylation
positive regulation of anoikis
regulation of interleukin-6 production
mammary gland alveolus development
positive regulation of neuron projection arborization
protein deubiquitination involved in ubiquitin-dependent protein catabolic process
prostaglandin metabolic process
negative regulation of interleukin-2 production
protein K63-linked deubiquitination
phosphorylation
positive regulation of cellular respiration
very long-chain fatty acid catabolic process
ether lipid metabolic process
nitric oxide biosynthetic process
negative regulation of leukocyte migration
fructose transmembrane transport
G1 to G0 transition involved in cell differentiation
response to carbohydrate
regulation of hydrogen peroxide metabolic process
positive regulation of MAPK cascade
embryonic placenta development
positive regulation of ubiquitin-dependent protein catabolic process
regulation of defense response to virus
liver development
iron-sulfur cluster assembly
generation of precursor metabolites and energy
cell adhesion mediated by integrin
isotype switching
positive regulation of T cell mediated cytotoxicity
cellular response to mechanical stimulus
HepaRG_FBS
HuH7_CDM
HuH7_FBS
PHH
cell_line
HepaRG
HuH7
PHH
medium
CDM
FBS

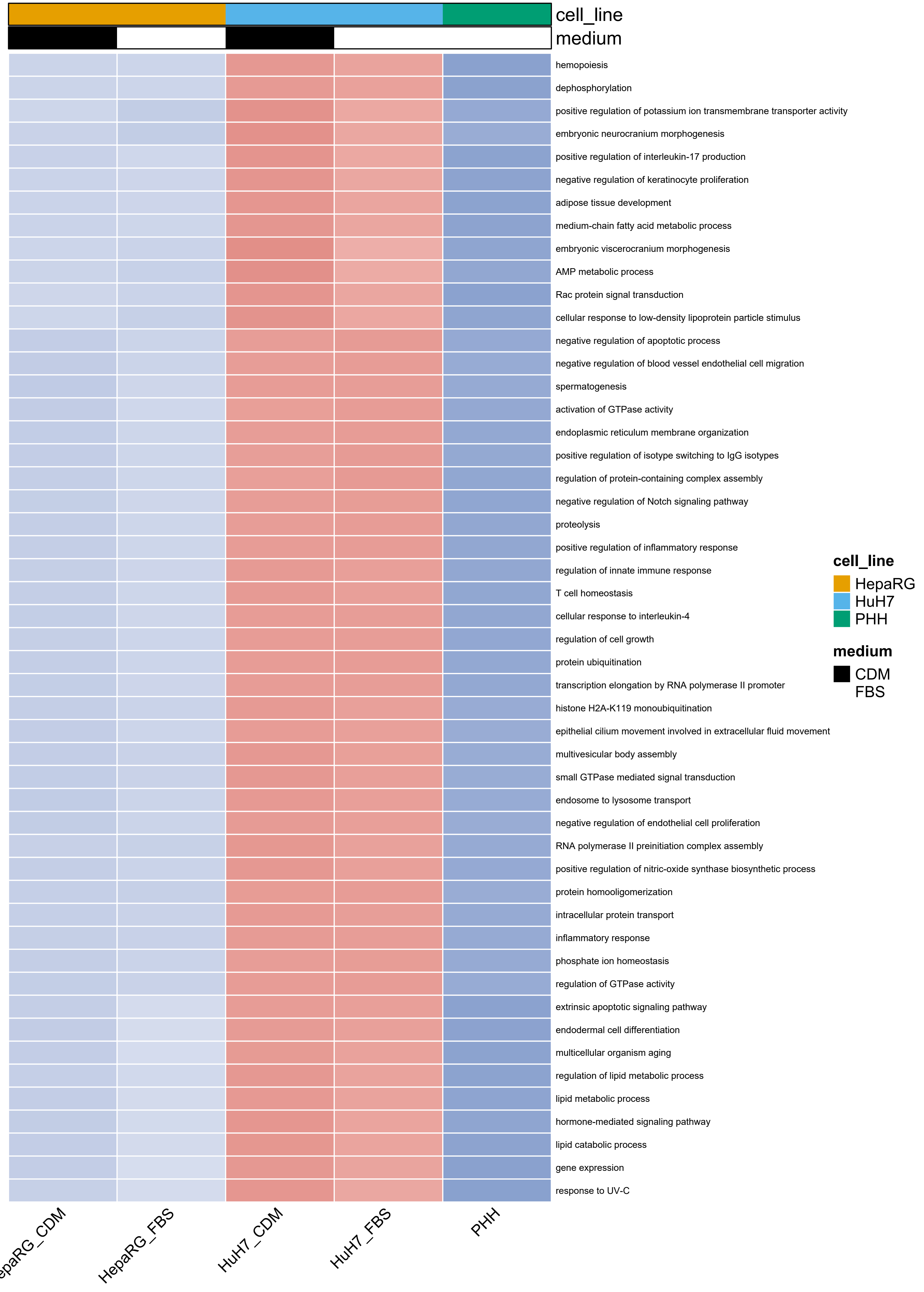

cell_line
medium
hemopoiesis
dephosphorylation
positive regulation of potassium ion transmembrane transporter activity
embryonic neurocranium morphogenesis
positive regulation of interleukin-17 production
negative regulation of keratinocyte proliferation
adipose tissue development
medium-chain fatty acid metabolic process
embryonic viscerocranium morphogenesis
AMP metabolic process
Rac protein signal transduction
cellular response to low-density lipoprotein particle stimulus
negative regulation of apoptotic process
negative regulation of blood vessel endothelial cell migration
spermatogenesis
activation of GTPase activity
endoplasmic reticulum membrane organization
positive regulation of isotype switching to IgG isotypes
regulation of protein-containing complex assembly
negative regulation of Notch signaling pathway
proteolysis
positive regulation of inflammatory response
regulation of innate immune response
T cell homeostasis
cellular response to interleukin-4
regulation of cell growth
protein ubiquitination
transcription elongation by RNA polymerase II promoter
histone H2A-K119 monoubiquitination
epithelial cilium movement involved in extracellular fluid movement
multivesicular body assembly
small GTPase mediated signal transduction
endosome to lysosome transport
negative regulation of endothelial cell proliferation
RNA polymerase II preinitiation complex assembly
positive regulation of nitric-oxide synthase biosynthetic process
protein homooligomerization
intracellular protein transport
inflammatory response
phosphate ion homeostasis
regulation of GTPase activity
extrinsic apoptotic signaling pathway
endodermal cell differentiation
multicellular organism aging
regulation of lipid metabolic process
lipid metabolic process
hormone-mediated signaling pathway
lipid catabolic process
gene expression
response to UV-C
HepaRG_FBS
HuH7_CDM
HuH7_FBS
PHH
cell_line
HepaRG
HuH7
PHH
medium
CDM
FBS

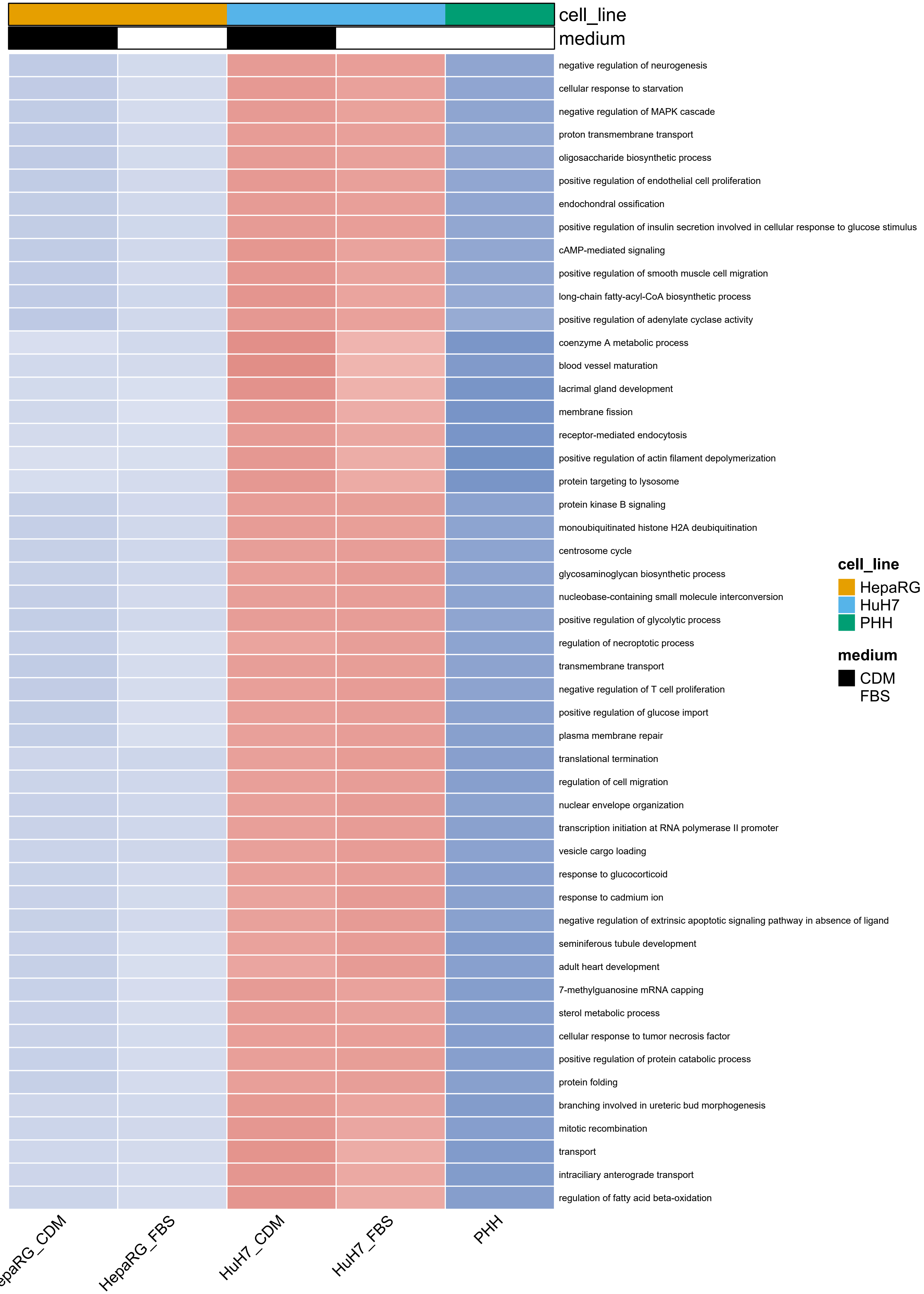
cell_line
medium
negative regulation of neurogenesis
cellular response to starvation
negative regulation of MAPK cascade
proton transmembrane transport
oligosaccharide biosynthetic process
positive regulation of endothelial cell proliferation
endochondral ossification
positive regulation of insulin secretion involved in cellular response to glucose stimulus
cAMP-mediated signaling
positive regulation of smooth muscle cell migration
long-chain fatty-acyl-CoA biosynthetic process
positive regulation of adenylate cyclase activity
coenzyme A metabolic process
blood vessel maturation
lacrimal gland development
membrane fission
receptor-mediated endocytosis
positive regulation of actin filament depolymerization
protein targeting to lysosome
protein kinase B signaling
monoubiquitinated histone H2A deubiquitination
centrosome cycle
glycosaminoglycan biosynthetic process
nucleobase-containing small molecule interconversion
positive regulation of glycolytic process
regulation of necroptotic process
transmembrane transport
negative regulation of T cell proliferation
positive regulation of glucose import
plasma membrane repair
translational termination
regulation of cell migration
nuclear envelope organization
transcription initiation at RNA polymerase II promoter
vesicle cargo loading
response to glucocorticoid
response to cadmium ion
negative regulation of extrinsic apoptotic signaling pathway in absence of ligand
seminiferous tubule development
adult heart development
7-methylguanosine mRNA capping
sterol metabolic process
cellular response to tumor necrosis factor
positive regulation of protein catabolic process
protein folding
branching involved in ureteric bud morphogenesis
mitotic recombination
transport
intraciliary anterograde transport
regulation of fatty acid beta-oxidation
HepaRG_CDM
HepaRG_FBS
HuH7_CDM
HuH7_FBS
PHH
cell_line
HepaRG
HuH7
PHH
medium
CDM
FBS

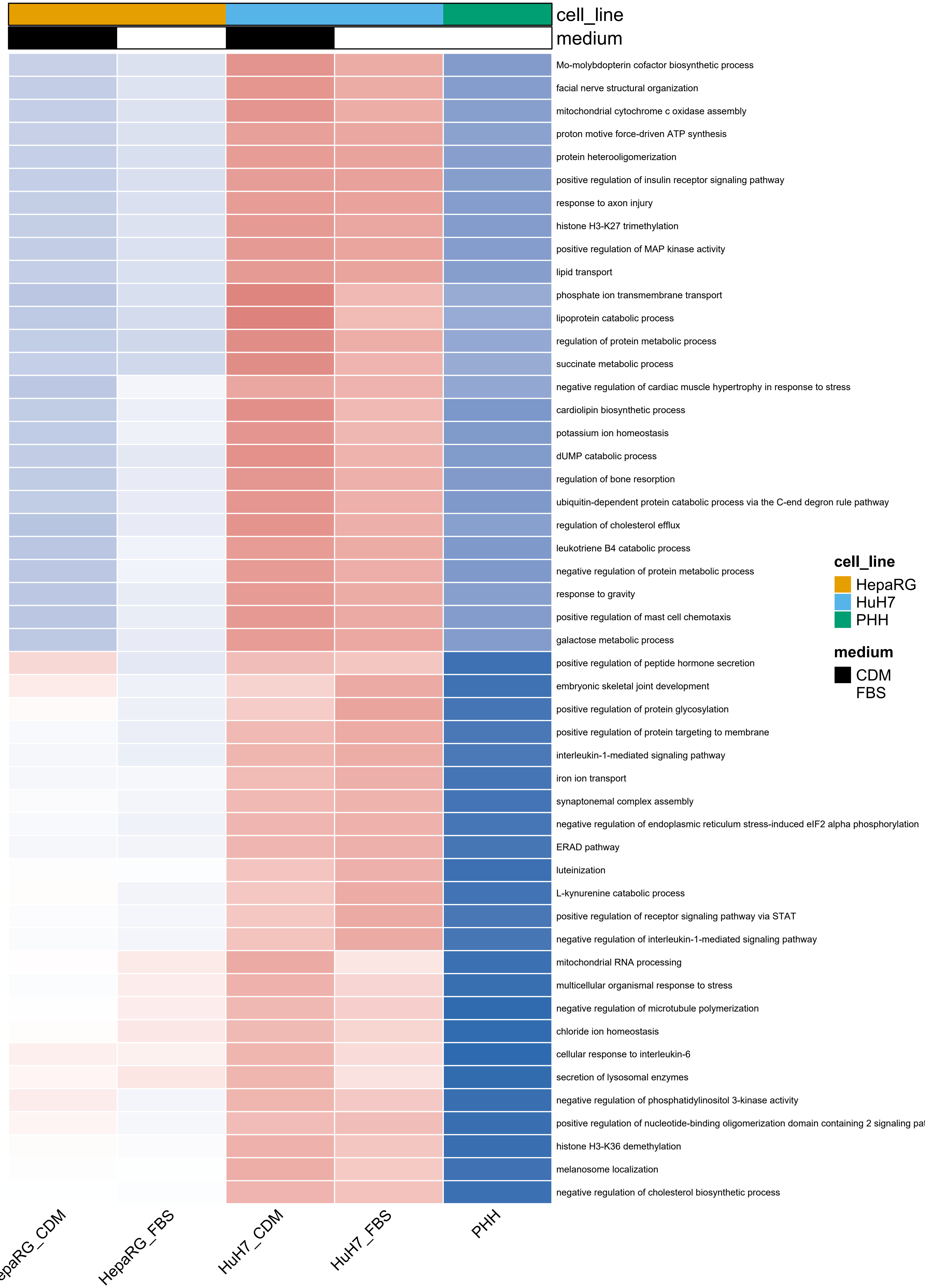
cell_line
medium
Mo-molybdopterin cofactor biosynthetic process
facial nerve structural organization
mitochondrial cytochrome c oxidase assembly
proton motive force-driven ATP synthesis
protein heterooligomerization
positive regulation of insulin receptor signaling pathway
response to axon injury
histone H3-K27 trimethylation
positive regulation of MAP kinase activity
lipid transport
phosphate ion transmembrane transport
lipoprotein catabolic process
regulation of protein metabolic process
succinate metabolic process
negative regulation of cardiac muscle hypertrophy in response to stress
cardiolipin biosynthetic process
potassium ion homeostasis
dUMP catabolic process
regulation of bone resorption
ubiquitin-dependent protein catabolic process via the C-end degron rule pathway
regulation of cholesterol efflux
leukotriene B4 catabolic process
negative regulation of protein metabolic process
response to gravity
positive regulation of mast cell chemotaxis
galactose metabolic process
positive regulation of peptide hormone secretion
embryonic skeletal joint development
positive regulation of protein glycosylation
positive regulation of protein targeting to membrane
interleukin-1-mediated signaling pathway
iron ion transport
synaptonemal complex assembly
negative regulation of endoplasmic reticulum stress-induced eIF2 alpha phosphorylation
ERAD pathway
luteinization
L-kynurenine catabolic process
positive regulation of receptor signaling pathway via STAT
negative regulation of interleukin-1-mediated signaling pathway
mitochondrial RNA processing
multicellular organismal response to stress
negative regulation of microtubule polymerization
chloride ion homeostasis
cellular response to interleukin-6
secretion of lysosomal enzymes
negative regulation of phosphatidylinositol 3-kinase activity
positive regulation of nucleotide-binding oligomerization domain containing 2 signaling pa
histone H3-K36 demethylation
melanosome localization
negative regulation of cholesterol biosynthetic process
cell_line
HepaRG
HuH7
PHH
medium
CDM
FBS
epaRG_CDM
HepaRG_FBS
HuH7_CDM
HuH7_FBS
PHH

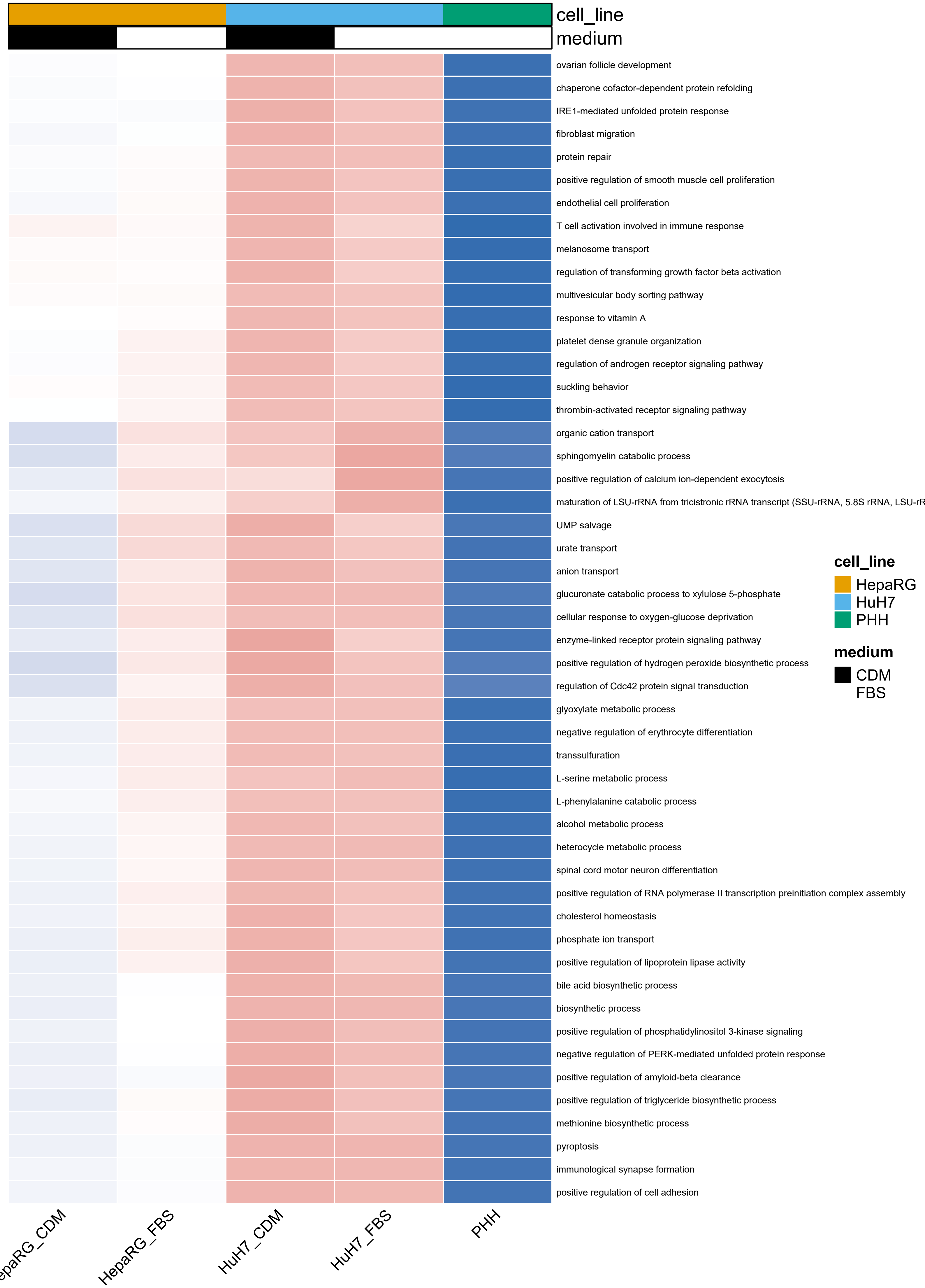

cell_line
medium
ovarian follicle development
chaperone cofactor-dependent protein refolding
IRE1-mediated unfolded protein response
fibroblast migration
protein repair
positive regulation of smooth muscle cell proliferation
endothelial cell proliferation
T cell activation involved in immune response
melanosome transport
regulation of transforming growth factor beta activation
multivesicular body sorting pathway
response to vitamin A
platelet dense granule organization
regulation of androgen receptor signaling pathway
suckling behavior
thrombin-activated receptor signaling pathway
organic cation transport
sphingomyelin catabolic process
positive regulation of calcium ion-dependent exocytosis
maturation of LSU-rRNA from tricistronic rRNA transcript (SSU-rRNA, 5.8S rRNA, LSU-rR
UMP salvage
urate transport
anion transport
glucuronate catabolic process to xylulose 5-phosphate
cellular response to oxygen-glucose deprivation
enzyme-linked receptor protein signaling pathway
positive regulation of hydrogen peroxide biosynthetic process
regulation of Cdc42 protein signal transduction
glyoxylate metabolic process
negative regulation of erythrocyte differentiation
transsulfuration
L-serine metabolic process
L-phenylalanine catabolic process
alcohol metabolic process
heterocycle metabolic process
spinal cord motor neuron differentiation
positive regulation of RNA polymerase II transcription preinitiation complex assembly
cholesterol homeostasis
phosphate ion transport
positive regulation of lipoprotein lipase activity
bile acid biosynthetic process
biosynthetic process
positive regulation of phosphatidylinositol 3-kinase signaling
negative regulation of PERK-mediated unfolded protein response
positive regulation of amyloid-beta clearance
positive regulation of triglyceride biosynthetic process
methionine biosynthetic process
pyroptosis
immunological synapse formation
positive regulation of cell adhesion
cell_line
HepaRG
HuH7
PHH
medium
CDM
FBS
epaRG_CDM
HepaRG_FBS
HuH7_CDM
HuH7_FBS
PHH

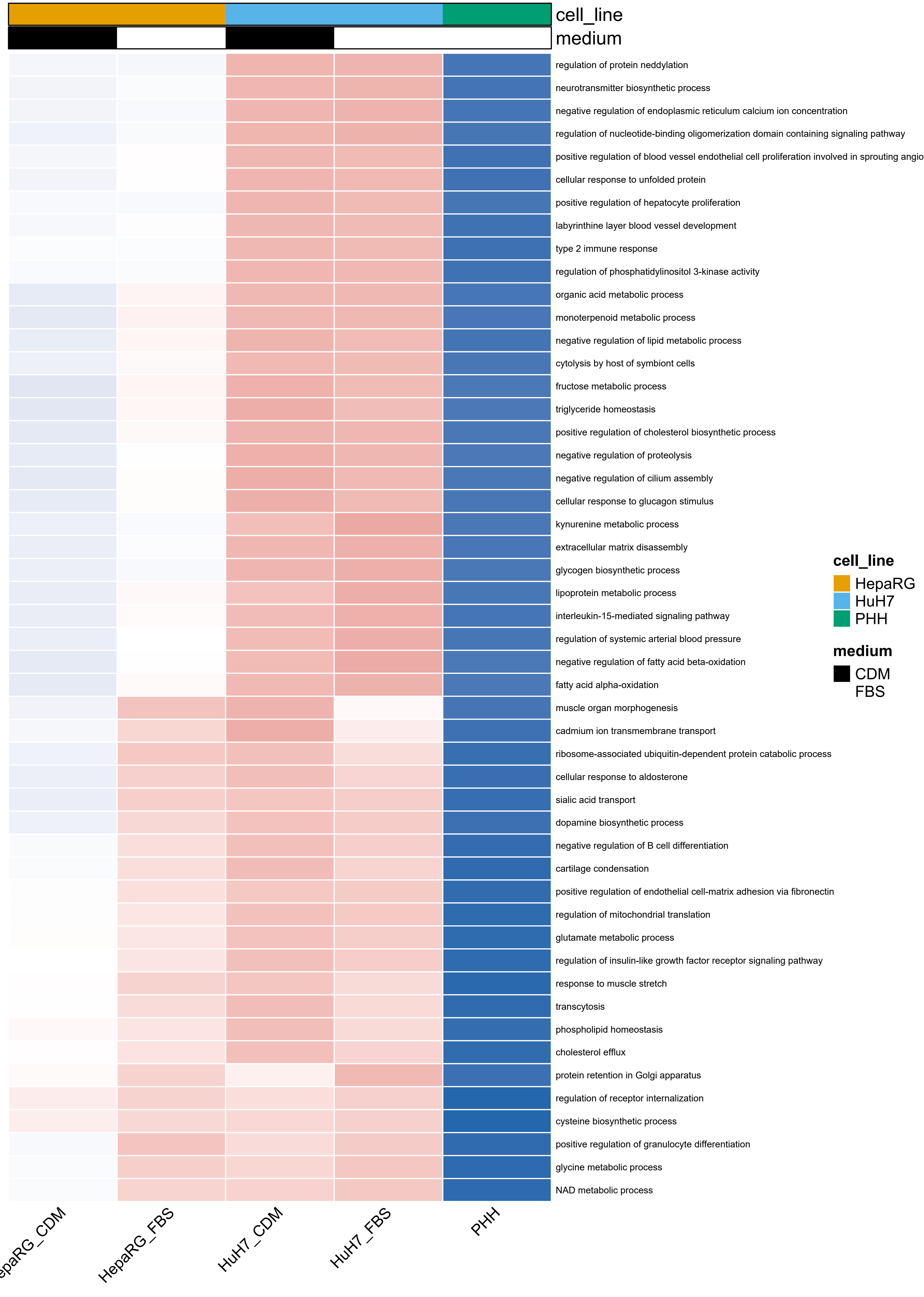
cell_line
medium
regulation of protein neddylation
neurotransmitter biosynthetic process
negative regulation of endoplasmic reticulum calcium ion concentration
regulation of nucleotide-binding oligomerization domain containing signaling pathway
positive regulation of blood vessel endothelial cell proliferation involved in sprouting angio
cellular response to unfolded protein
positive regulation of hepatocyte proliferation
labyrinthine layer blood vessel development
type 2 immune response
regulation of phosphatidylinositol 3-kinase activity
organic acid metabolic process
monoterpenoid metabolic process
negative regulation of lipid metabolic process
cytolysis by host of symbiont cells
fructose metabolic process
triglyceride homeostasis
positive regulation of cholesterol biosynthetic process
negative regulation of proteolysis
negative regulation of cilium assembly
cellular response to glucagon stimulus
kynurenine metabolic process
extracellular matrix disassembly
glycogen biosynthetic process
lipoprotein metabolic process
interleukin-15-mediated signaling pathway
regulation of systemic arterial blood pressure
negative regulation of fatty acid beta-oxidation
fatty acid alpha-oxidation
muscle organ morphogenesis
cadmium ion transmembrane transport
ribosome-associated ubiquitin-dependent protein catabolic process
cellular response to aldosterone
sialic acid transport
dopamine biosynthetic process
negative regulation of B cell differentiation
cartilage condensation
positive regulation of endothelial cell-matrix adhesion via fibronectin
regulation of mitochondrial translation
glutamate metabolic process
regulation of insulin-like growth factor receptor signaling pathway
response to muscle stretch
transcytosis
phospholipid homeostasis
cholesterol efflux
protein retention in Golgi apparatus
regulation of receptor internalization
cysteine biosynthetic process
positive regulation of granulocyte differentiation
glycine metabolic process
NAD metabolic process
cell_line
HepaRG
HuH7
PHH
medium
CDM
FBS
epaRG_CDM
HepaRG_FBS
HuH7_CDM
HuH7_FBS
PHH

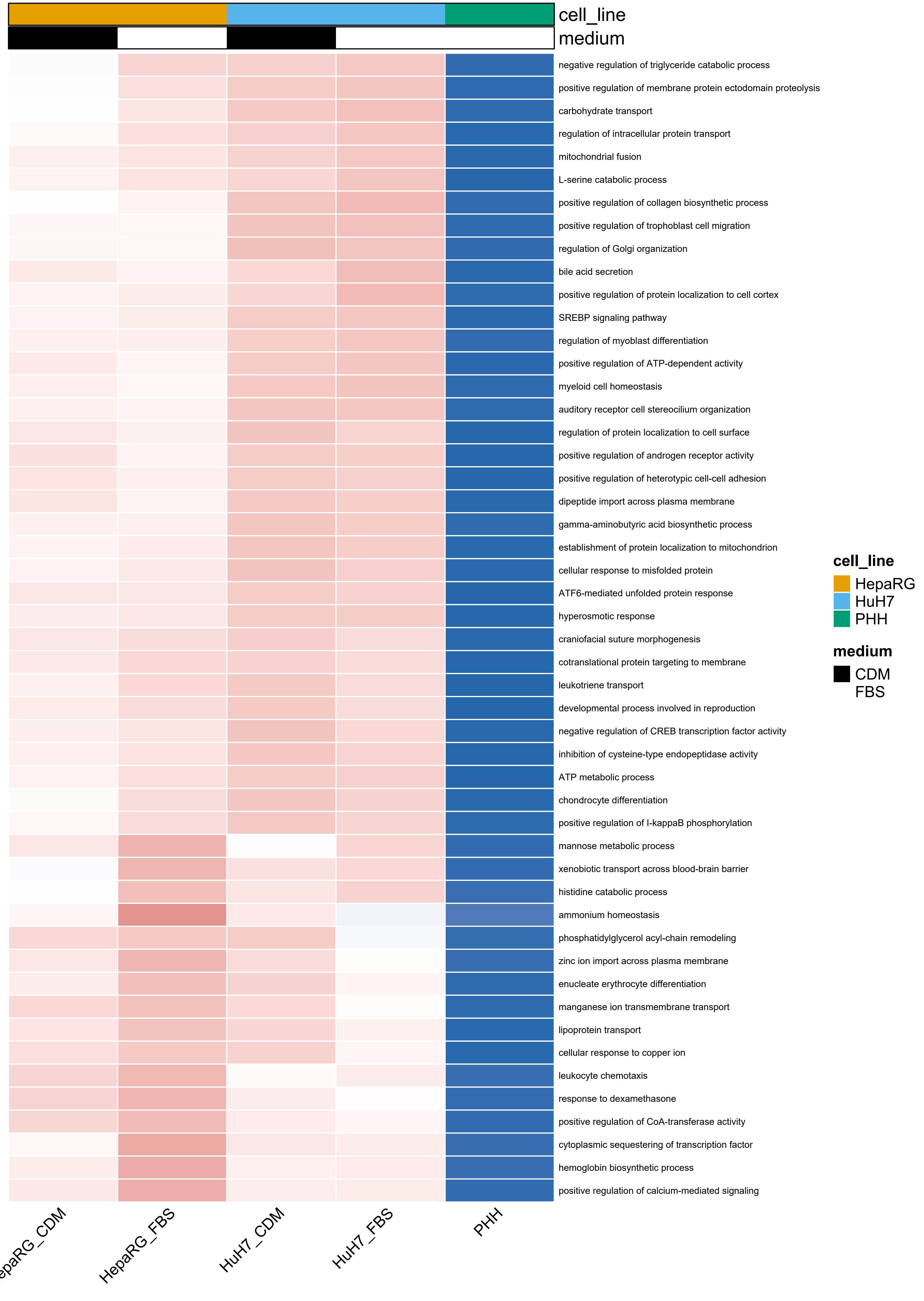
cell_line
medium
negative regulation of triglyceride catabolic process
positive regulation of membrane protein ectodomain proteolysis
carbohydrate transport
regulation of intracellular protein transport
mitochondrial fusion
L-serine catabolic process
positive regulation of collagen biosynthetic process
positive regulation of trophoblast cell migration
regulation of Golgi organization
bile acid secretion
positive regulation of protein localization to cell cortex
SREBP signaling pathway
regulation of myoblast differentiation
positive regulation of ATP-dependent activity
myeloid cell homeostasis
auditory receptor cell stereocilium organization
regulation of protein localization to cell surface
positive regulation of androgen receptor activity
positive regulation of heterotypic cell-cell adhesion
dipeptide import across plasma membrane
gamma-aminobutyric acid biosynthetic process
establishment of protein localization to mitochondrion
cellular response to misfolded protein
ATF6-mediated unfolded protein response
hyperosmotic response
craniofacial suture morphogenesis
cotranslational protein targeting to membrane
leukotriene transport
developmental process involved in reproduction
negative regulation of CREB transcription factor activity
inhibition of cysteine-type endopeptidase activity
ATP metabolic process
chondrocyte differentiation
positive regulation of I-kappaB phosphorylation
mannose metabolic process
xenobiotic transport across blood-brain barrier
histidine catabolic process
ammonium homeostasis
phosphatidylglycerol acyl-chain remodeling
zinc ion import across plasma membrane
enucleate erythrocyte differentiation
manganese ion transmembrane transport
lipoprotein transport
cellular response to copper ion
leukocyte chemotaxis
response to dexamethasone
positive regulation of CoA-transferase activity
cytoplasmic sequestering of transcription factor
hemoglobin biosynthetic process
positive regulation of calcium-mediated signaling
cell_line
HepaRG
HuH7
PHH
medium
CDM
FBS
HepaRG_FBS
HuH7_CDM
HuH7_FBS
PHH

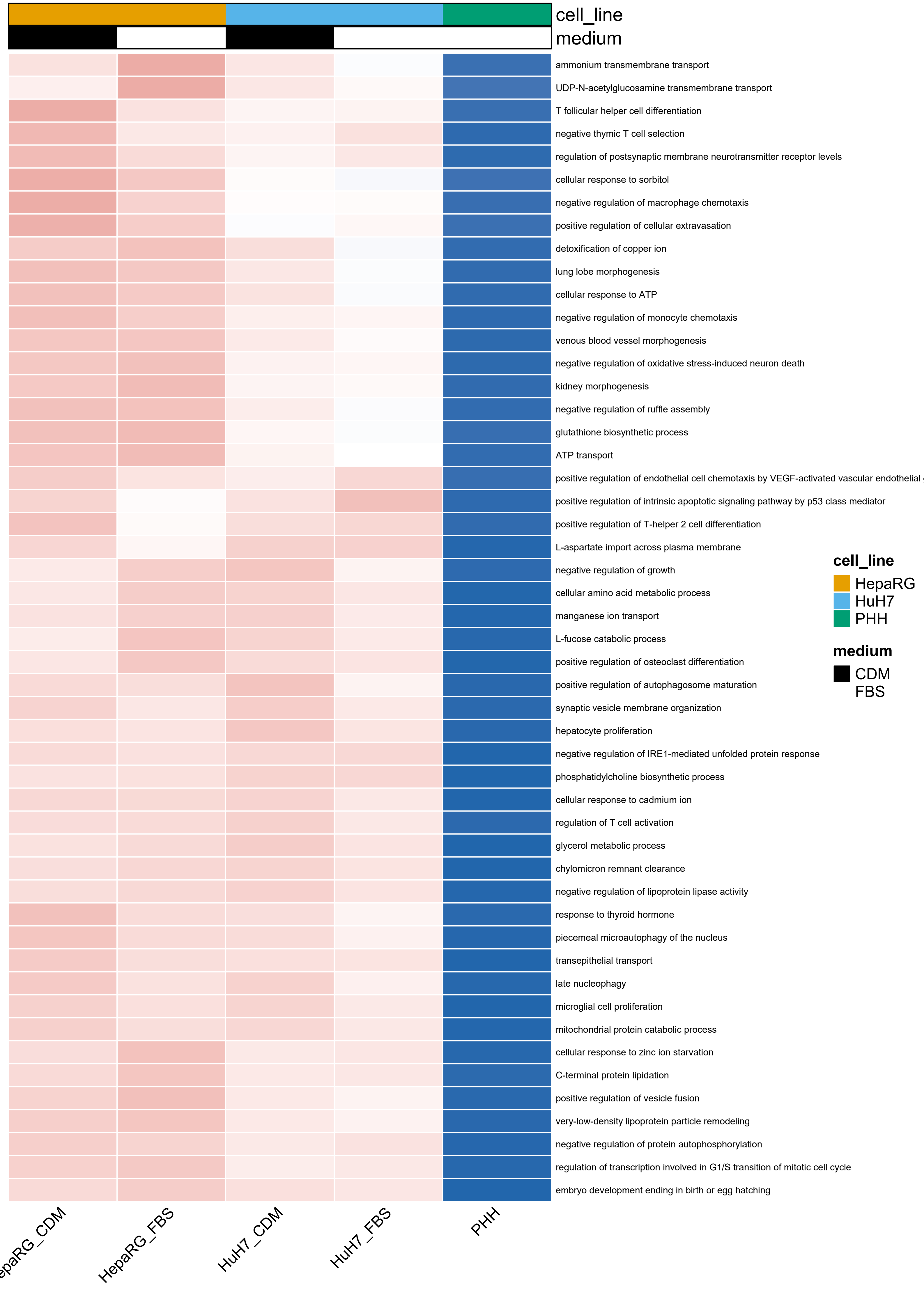
cell_line
medium
ammonium transmembrane transport
UDP-N-acetylglucosamine transmembrane transport
T follicular helper cell differentiation
negative thymic T cell selection
regulation of postsynaptic membrane neurotransmitter receptor levels
cellular response to sorbitol
negative regulation of macrophage chemotaxis
positive regulation of cellular extravasation
detoxification of copper ion
lung lobe morphogenesis
cellular response to ATP
negative regulation of monocyte chemotaxis
venous blood vessel morphogenesis
negative regulation of oxidative stress-induced neuron death
kidney morphogenesis
negative regulation of ruffle assembly
glutathione biosynthetic process
ATP transport
positive regulation of endothelial cell chemotaxis by VEGF-activated vascular endothelial
positive regulation of intrinsic apoptotic signaling pathway by p53 class mediator
positive regulation of T-helper 2 cell differentiation
L-aspartate import across plasma membrane
negative regulation of growth
cellular amino acid metabolic process
manganese ion transport
L-fucose catabolic process
positive regulation of osteoclast differentiation
positive regulation of autophagosome maturation
synaptic vesicle membrane organization
hepatocyte proliferation
negative regulation of IRE1-mediated unfolded protein response
phosphatidylcholine biosynthetic process
cellular response to cadmium ion
regulation of T cell activation
glycerol metabolic process
chylomicron remnant clearance
negative regulation of lipoprotein lipase activity
response to thyroid hormone
piecemeal microautophagy of the nucleus
transepithelial transport
late nucleophagy
microglial cell proliferation
mitochondrial protein catabolic process
cellular response to zinc ion starvation
C-terminal protein lipidation
positive regulation of vesicle fusion
very-low-density lipoprotein particle remodeling
negative regulation of protein autophosphorylation
regulation of transcription involved in G1/S transition of mitotic cell cycle
embryo development ending in birth or egg hatching
cell_line
HepaRG
HuH7
PHH
medium
CDM
FBS
HepaRG_FBS
HuH7_CDM
HuH7_FBS
PHH

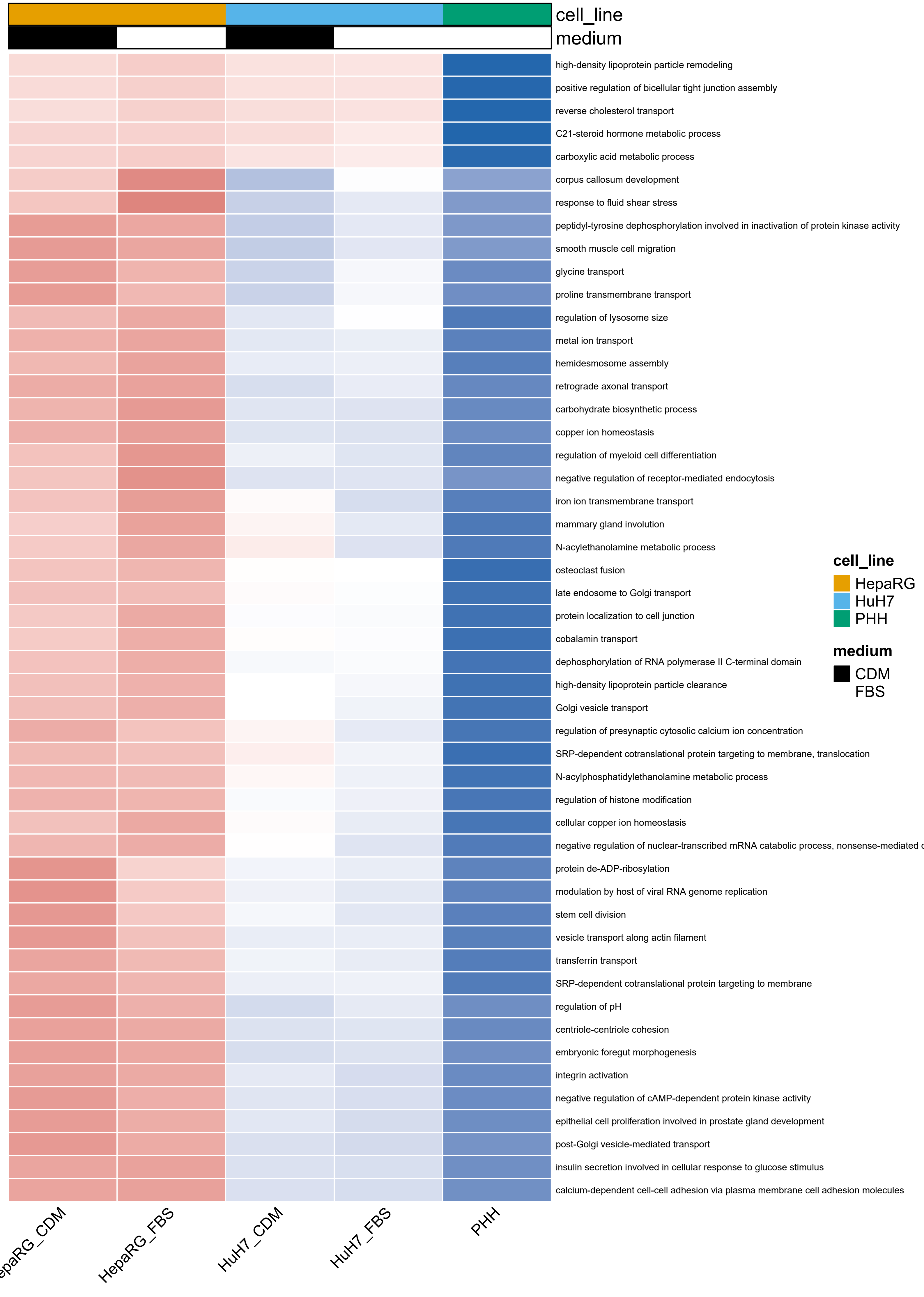
cell_line
medium
high-density lipoprotein particle remodeling
positive regulation of bicellular tight junction assembly
reverse cholesterol transport
C21-steroid hormone metabolic process
carboxylic acid metabolic process
corpus callosum development
response to fluid shear stress
peptidyl-tyrosine dephosphorylation involved in inactivation of protein kinase activity
smooth muscle cell migration
glycine transport
proline transmembrane transport
regulation of lysosome size
metal ion transport
hemidesmosome assembly
retrograde axonal transport
carbohydrate biosynthetic process
copper ion homeostasis
regulation of myeloid cell differentiation
negative regulation of receptor-mediated endocytosis
iron ion transmembrane transport
mammary gland involution
N-acylethanolamine metabolic process
osteoclast fusion
late endosome to Golgi transport
protein localization to cell junction
cobalamin transport
dephosphorylation of RNA polymerase II C-terminal domain
high-density lipoprotein particle clearance
Golgi vesicle transport
regulation of presynaptic cytosolic calcium ion concentration
SRP-dependent cotranslational protein targeting to membrane, translocation
N-acylphosphatidylethanolamine metabolic process
regulation of histone modification
cellular copper ion homeostasis
negative regulation of nuclear-transcribed mRNA catabolic process, nonsense-mediated
protein de-ADP-ribosylation
modulation by host of viral RNA genome replication
stem cell division
vesicle transport along actin filament
transferrin transport
SRP-dependent cotranslational protein targeting to membrane
regulation of pH
centriole-centriole cohesion
embryonic foregut morphogenesis
integrin activation
negative regulation of cAMP-dependent protein kinase activity
epithelial cell proliferation involved in prostate gland development
post-Golgi vesicle-mediated transport
insulin secretion involved in cellular response to glucose stimulus
calcium-dependent cell-cell adhesion via plasma membrane cell adhesion molecules
cell_line
HepaRG
HuH7
PHH
medium
CDM
FBS
HepaRG_FBS
HuH7_CDM
HuH7_FBS
PHH

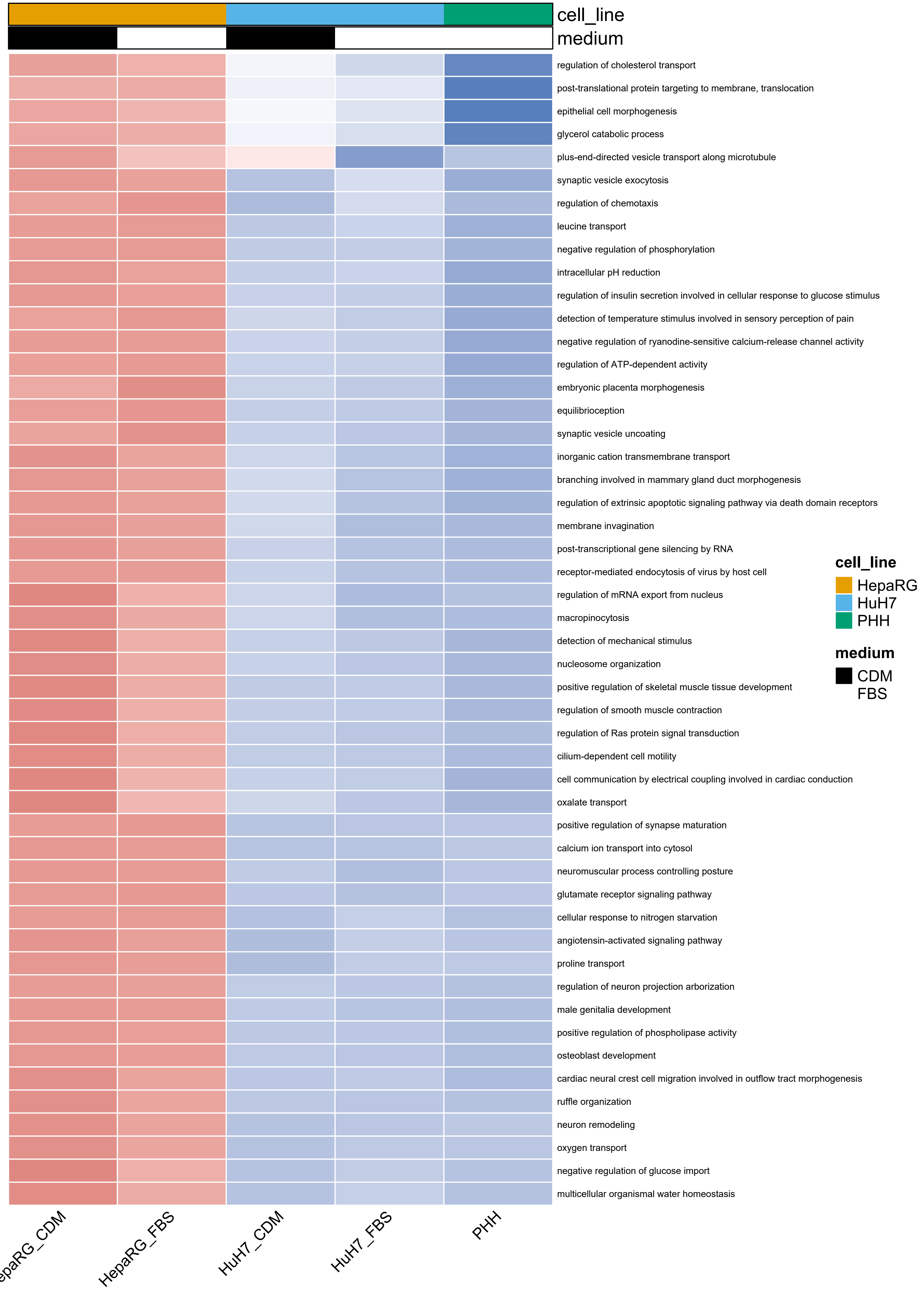

cell_line
medium
regulation of cholesterol transport
post-translational protein targeting to membrane, translocation
epithelial cell morphogenesis
glycerol catabolic process
plus-end-directed vesicle transport along microtubule
synaptic vesicle exocytosis
regulation of chemotaxis
leucine transport
negative regulation of phosphorylation
intracellular pH reduction
regulation of insulin secretion involved in cellular response to glucose stimulus
detection of temperature stimulus involved in sensory perception of pain
negative regulation of ryanodine-sensitive calcium-release channel activity
regulation of ATP-dependent activity
embryonic placenta morphogenesis
equilibrioception
synaptic vesicle uncoating
inorganic cation transmembrane transport
branching involved in mammary gland duct morphogenesis
regulation of extrinsic apoptotic signaling pathway via death domain receptors
membrane invagination
post-transcriptional gene silencing by RNA
receptor-mediated endocytosis of virus by host cell
regulation of mRNA export from nucleus
macropinocytosis
detection of mechanical stimulus
nucleosome organization
positive regulation of skeletal muscle tissue development
regulation of smooth muscle contraction
regulation of Ras protein signal transduction
cilium-dependent cell motility
cell communication by electrical coupling involved in cardiac conduction
oxalate transport
positive regulation of synapse maturation
calcium ion transport into cytosol
neuromuscular process controlling posture
glutamate receptor signaling pathway
cellular response to nitrogen starvation
angiotensin-activated signaling pathway
proline transport
regulation of neuron projection arborization
male genitalia development
positive regulation of phospholipase activity
osteoblast development
cardiac neural crest cell migration involved in outflow tract morphogenesis
ruffle organization
neuron remodeling
oxygen transport
negative regulation of glucose import
multicellular organismal water homeostasis
cell_line
HepaRG
HuH7
PHH
medium
CDM
FBS
HepaRG_FBS
HuH7_CDM
HuH7_FBS
PHH

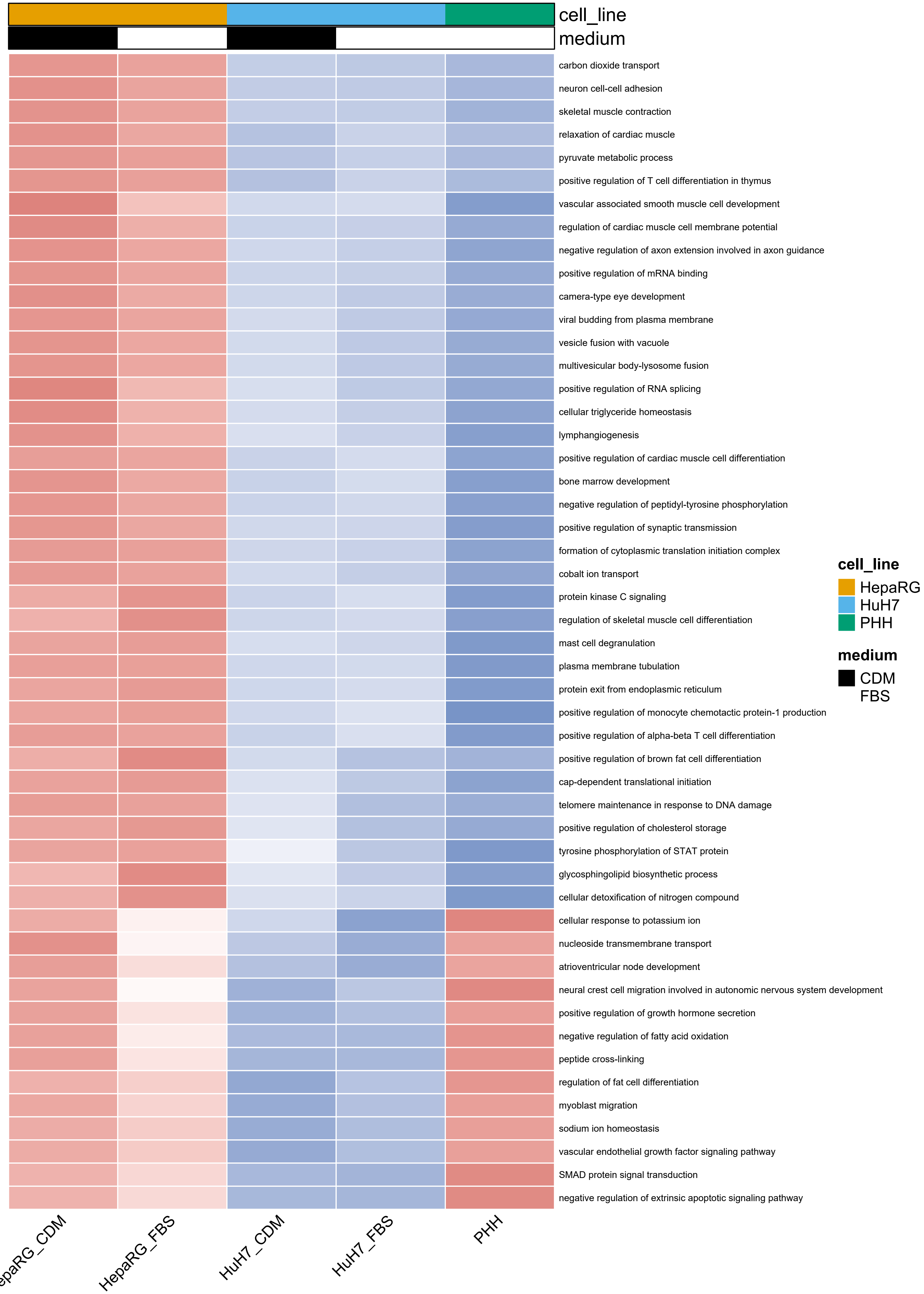
cell_line
medium
carbon dioxide transport
neuron cell-cell adhesion
skeletal muscle contraction
relaxation of cardiac muscle
pyruvate metabolic process
positive regulation of T cell differentiation in thymus
vascular associated smooth muscle cell development
regulation of cardiac muscle cell membrane potential
negative regulation of axon extension involved in axon guidance
positive regulation of mRNA binding
camera-type eye development
viral budding from plasma membrane
vesicle fusion with vacuole
multivesicular body-lysosome fusion
positive regulation of RNA splicing
cellular triglyceride homeostasis
lymphangiogenesis
positive regulation of cardiac muscle cell differentiation
bone marrow development
negative regulation of peptidyl-tyrosine phosphorylation
positive regulation of synaptic transmission
formation of cytoplasmic translation initiation complex
cobalt ion transport
protein kinase C signaling
regulation of skeletal muscle cell differentiation
mast cell degranulation
plasma membrane tubulation
protein exit from endoplasmic reticulum
positive regulation of monocyte chemotactic protein-1 production
positive regulation of alpha-beta T cell differentiation
positive regulation of brown fat cell differentiation
cap-dependent translational initiation
telomere maintenance in response to DNA damage
positive regulation of cholesterol storage
tyrosine phosphorylation of STAT protein
glycosphingolipid biosynthetic process
cellular detoxification of nitrogen compound
cellular response to potassium ion
nucleoside transmembrane transport
atrioventricular node development
neural crest cell migration involved in autonomic nervous system development
positive regulation of growth hormone secretion
negative regulation of fatty acid oxidation
peptide cross-linking
regulation of fat cell differentiation
myoblast migration
sodium ion homeostasis
vascular endothelial growth factor signaling pathway
SMAD protein signal transduction
negative regulation of extrinsic apoptotic signaling pathway
cell_line
HepaRG
HuH7
PHH
medium
CDM
FBS
HepaRG_FBS
HuH7_CDM
HuH7_FBS
PHH

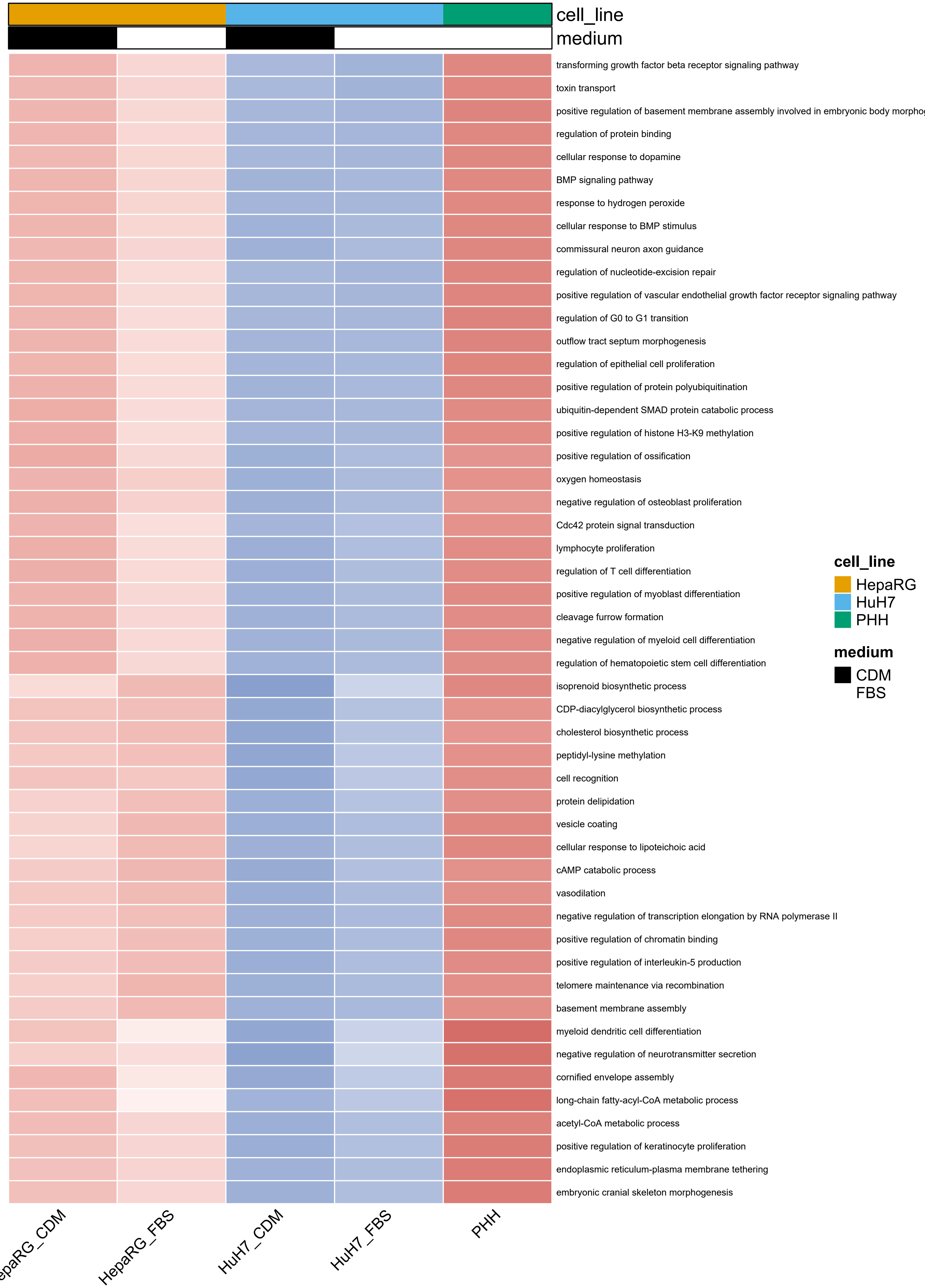
cell_line
medium
transforming growth factor beta receptor signaling pathway
toxin transport
positive regulation of basement membrane assembly involved in embryonic body morpho
regulation of protein binding
cellular response to dopamine
BMP signaling pathway
response to hydrogen peroxide
cellular response to BMP stimulus
commissural neuron axon guidance
regulation of nucleotide-excision repair
positive regulation of vascular endothelial growth factor receptor signaling pathway
regulation of G0 to G1 transition
outflow tract septum morphogenesis
regulation of epithelial cell proliferation
positive regulation of protein polyubiquitination
ubiquitin-dependent SMAD protein catabolic process
positive regulation of histone H3-K9 methylation
positive regulation of ossification
oxygen homeostasis
negative regulation of osteoblast proliferation
Cdc42 protein signal transduction
lymphocyte proliferation
regulation of T cell differentiation
positive regulation of myoblast differentiation
cleavage furrow formation
negative regulation of myeloid cell differentiation
regulation of hematopoietic stem cell differentiation
isoprenoid biosynthetic process
CDP-diacylglycerol biosynthetic process
cholesterol biosynthetic process
peptidyl-lysine methylation
cell recognition
protein delipidation
vesicle coating
cellular response to lipoteichoic acid
cAMP catabolic process
vasodilation
negative regulation of transcription elongation by RNA polymerase II
positive regulation of chromatin binding
positive regulation of interleukin-5 production
telomere maintenance via recombination
basement membrane assembly
myeloid dendritic cell differentiation
negative regulation of neurotransmitter secretion
cornified envelope assembly
long-chain fatty-acyl-CoA metabolic process
acetyl-CoA metabolic process
positive regulation of keratinocyte proliferation
endoplasmic reticulum-plasma membrane tethering
embryonic cranial skeleton morphogenesis
cell_line
HepaRG
HuH7
PHH
medium
CDM
FBS
epaRG_CDM
HepaRG_FBS
HuH7_CDM
HuH7_FBS
PHH

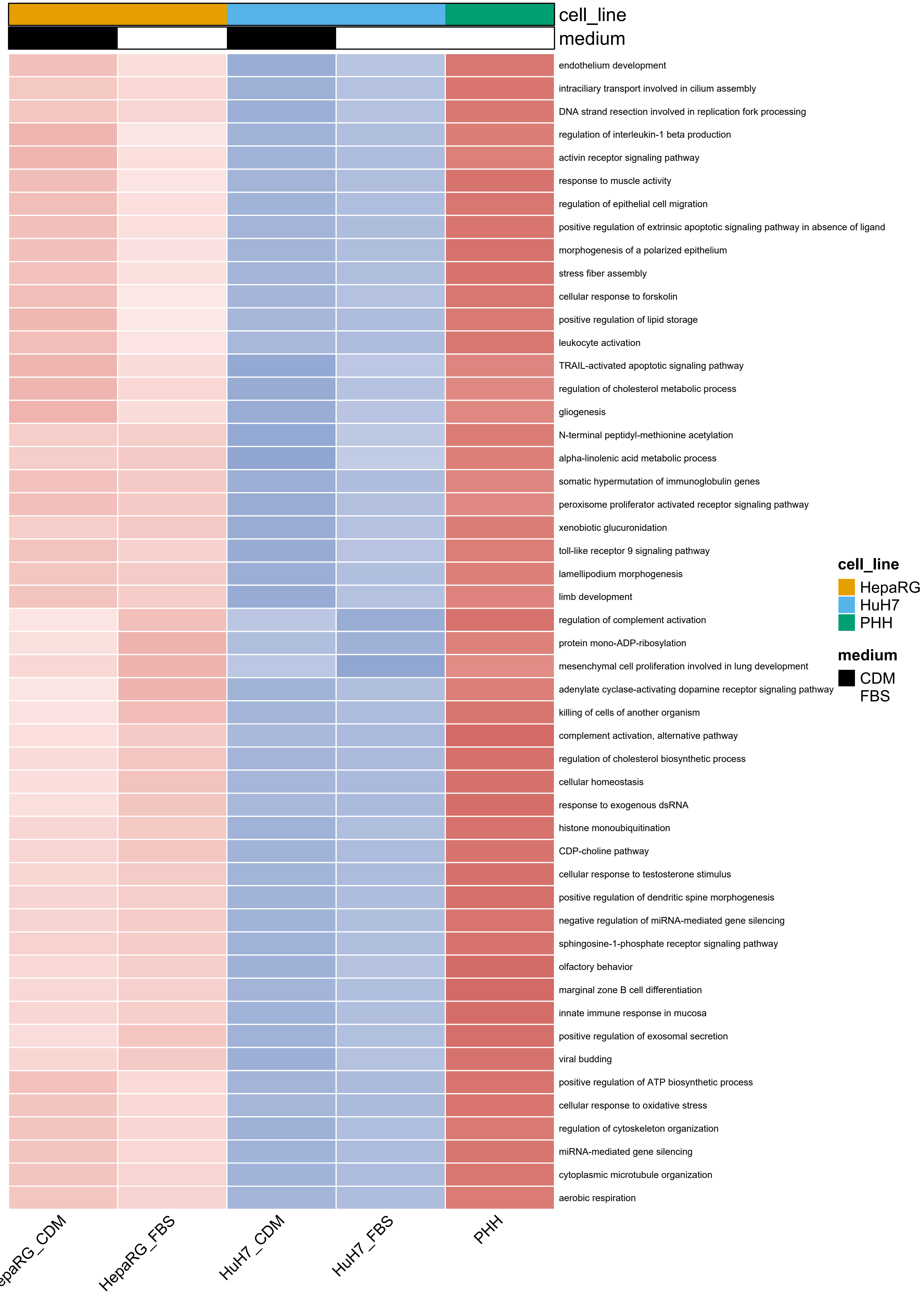

cell_line
medium
endothelium development
intraciliary transport involved in cilium assembly
DNA strand resection involved in replication fork processing
regulation of interleukin-1 beta production
activin receptor signaling pathway
response to muscle activity
regulation of epithelial cell migration
positive regulation of extrinsic apoptotic signaling pathway in absence of ligand
morphogenesis of a polarized epithelium
stress fiber assembly
cellular response to forskolin
positive regulation of lipid storage
leukocyte activation
TRAIL-activated apoptotic signaling pathway
regulation of cholesterol metabolic process
gliogenesis
N-terminal peptidyl-methionine acetylation
alpha-linolenic acid metabolic process
somatic hypermutation of immunoglobulin genes
peroxisome proliferator activated receptor signaling pathway
xenobiotic glucuronidation
toll-like receptor 9 signaling pathway
lamellipodium morphogenesis
limb development
regulation of complement activation
protein mono-ADP-ribosylation
mesenchymal cell proliferation involved in lung development
adenylate cyclase-activating dopamine receptor signaling pathway
killing of cells of another organism
complement activation, alternative pathway
regulation of cholesterol biosynthetic process
cellular homeostasis
response to exogenous dsRNA
histone monoubiquitination
CDP-choline pathway
cellular response to testosterone stimulus
positive regulation of dendritic spine morphogenesis
negative regulation of miRNA-mediated gene silencing
sphingosine-1-phosphate receptor signaling pathway
olfactory behavior
marginal zone B cell differentiation
innate immune response in mucosa
positive regulation of exosomal secretion
viral budding
positive regulation of ATP biosynthetic process
cellular response to oxidative stress
regulation of cytoskeleton organization
miRNA-mediated gene silencing
cytoplasmic microtubule organization
aerobic respiration
cell_line
HepaRG
HuH7
PHH
medium
CDM
FBS
HepaRG_FBS
HuH7_CDM
HuH7_FBS
PHH

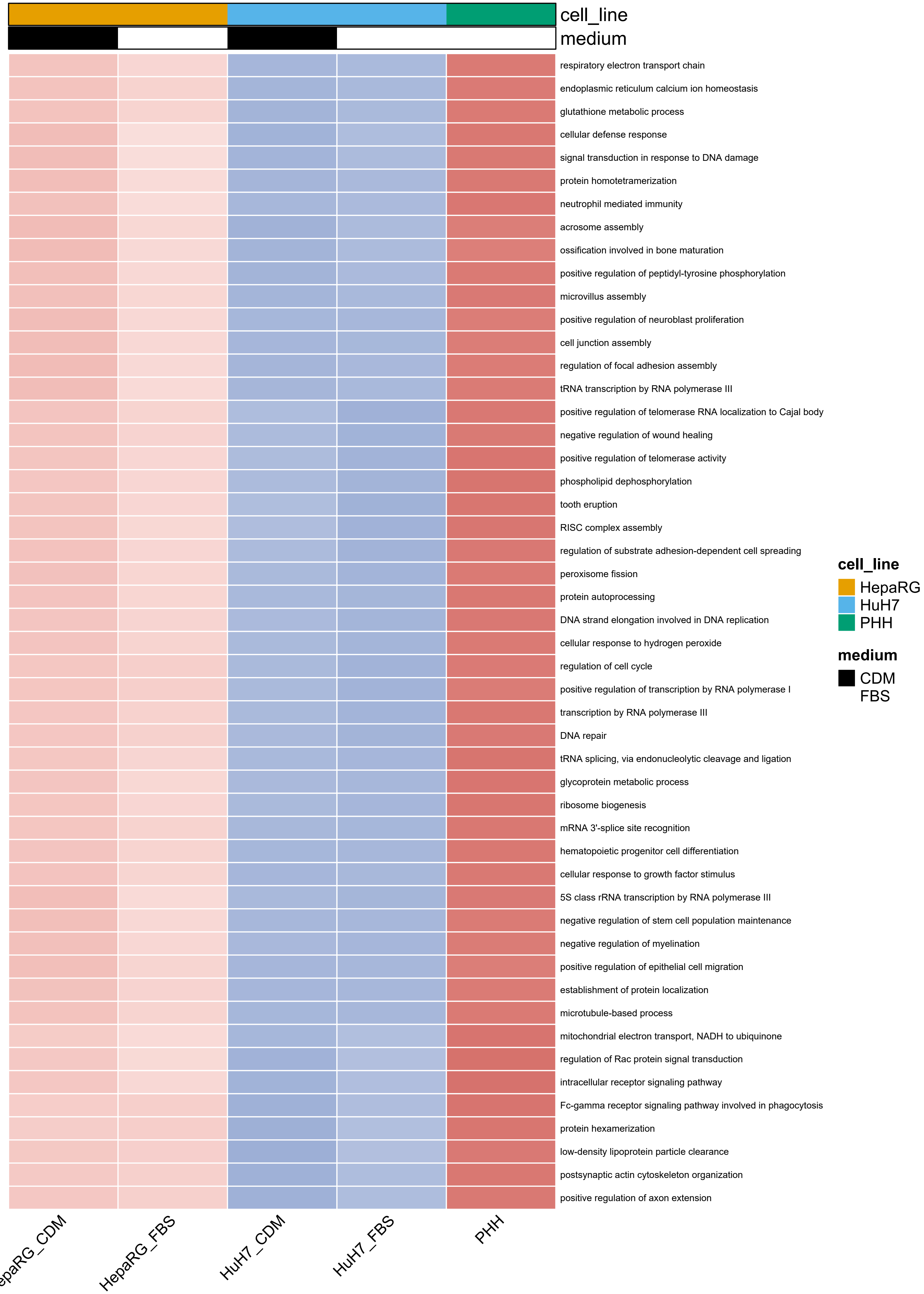
cell_line
medium
respiratory electron transport chain
endoplasmic reticulum calcium ion homeostasis
glutathione metabolic process
cellular defense response
signal transduction in response to DNA damage
protein homotetramerization
neutrophil mediated immunity
acrosome assembly
ossification involved in bone maturation
positive regulation of peptidyl-tyrosine phosphorylation
microvillus assembly
positive regulation of neuroblast proliferation
cell junction assembly
regulation of focal adhesion assembly
tRNA transcription by RNA polymerase III
positive regulation of telomerase RNA localization to Cajal body
negative regulation of wound healing
positive regulation of telomerase activity
phospholipid dephosphorylation
tooth eruption
RISC complex assembly
regulation of substrate adhesion-dependent cell spreading
peroxisome fission
protein autoprocessing
DNA strand elongation involved in DNA replication
cellular response to hydrogen peroxide
regulation of cell cycle
positive regulation of transcription by RNA polymerase I
transcription by RNA polymerase III
DNA repair
tRNA splicing, via endonucleolytic cleavage and ligation
glycoprotein metabolic process
ribosome biogenesis
mRNA 3'-splice site recognition
hematopoietic progenitor cell differentiation
cellular response to growth factor stimulus
5S class rRNA transcription by RNA polymerase III
negative regulation of stem cell population maintenance
negative regulation of myelination
positive regulation of epithelial cell migration
establishment of protein localization
microtubule-based process
mitochondrial electron transport, NADH to ubiquinone
regulation of Rac protein signal transduction
intracellular receptor signaling pathway
Fc-gamma receptor signaling pathway involved in phagocytosis
protein hexamerization
low-density lipoprotein particle clearance
postsynaptic actin cytoskeleton organization
positive regulation of axon extension
HepaRG_CDM
HepaRG_FBS
HuH7_CDM
HuH7_FBS
PHH
cell_line
HepaRG
HuH7
PHH
medium
CDM
FBS

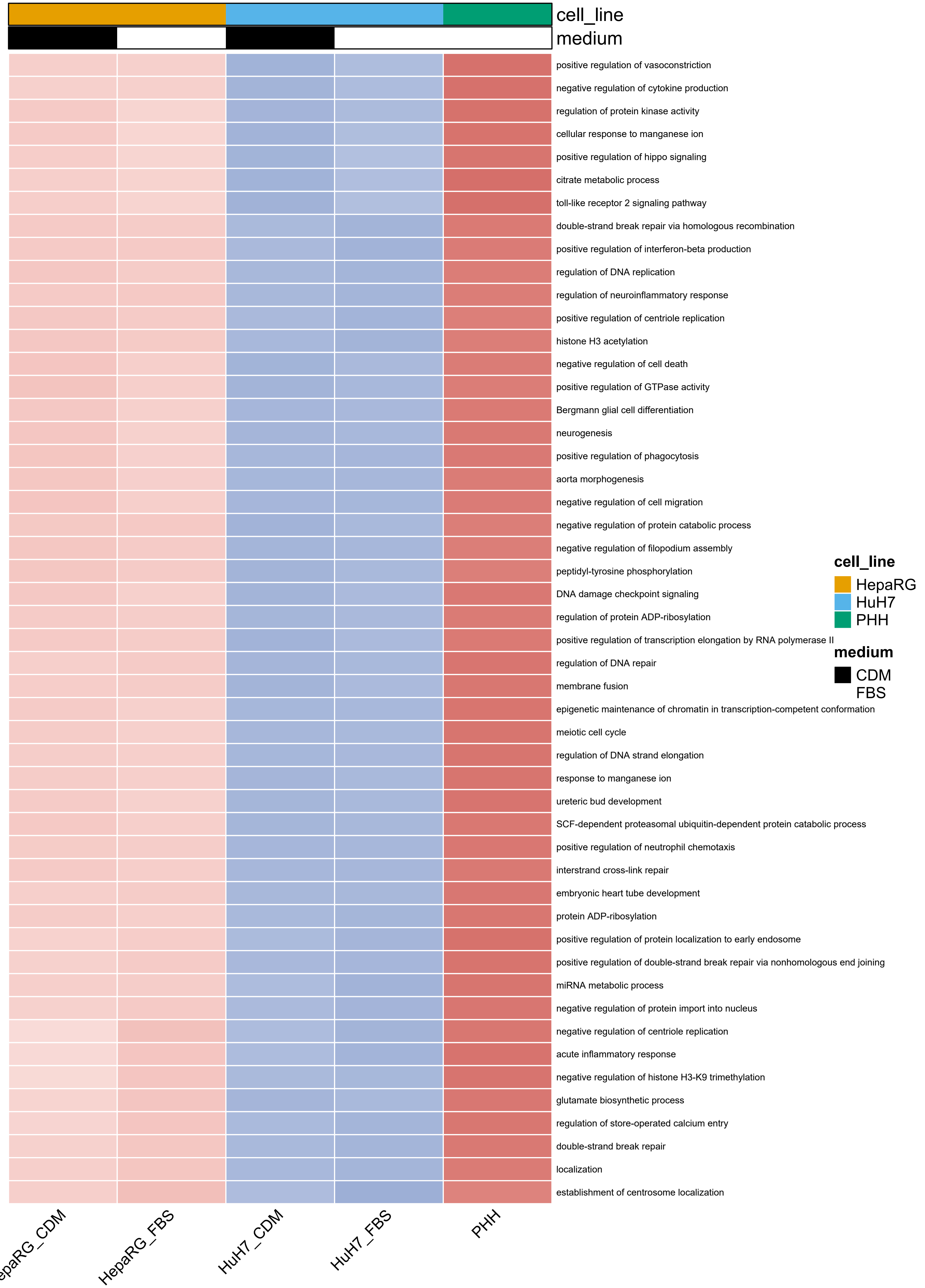
cell_line
medium
positive regulation of vasoconstriction
negative regulation of cytokine production
regulation of protein kinase activity
cellular response to manganese ion
positive regulation of hippo signaling
citrate metabolic process
toll-like receptor 2 signaling pathway
double-strand break repair via homologous recombination
positive regulation of interferon-beta production
regulation of DNA replication
regulation of neuroinflammatory response
positive regulation of centriole replication
histone H3 acetylation
negative regulation of cell death
positive regulation of GTPase activity
Bergmann glial cell differentiation
neurogenesis
positive regulation of phagocytosis
aorta morphogenesis
negative regulation of cell migration
negative regulation of protein catabolic process
negative regulation of filopodium assembly
peptidyl-tyrosine phosphorylation
DNA damage checkpoint signaling
regulation of protein ADP-ribosylation
positive regulation of transcription elongation by RNA polymerase II
regulation of DNA repair
membrane fusion
epigenetic maintenance of chromatin in transcription-competent conformation
meiotic cell cycle
regulation of DNA strand elongation
response to manganese ion
ureteric bud development
SCF-dependent proteasomal ubiquitin-dependent protein catabolic process
positive regulation of neutrophil chemotaxis
interstrand cross-link repair
embryonic heart tube development
protein ADP-ribosylation
positive regulation of protein localization to early endosome
positive regulation of double-strand break repair via nonhomologous end joining
miRNA metabolic process
negative regulation of protein import into nucleus
negative regulation of centriole replication
acute inflammatory response
negative regulation of histone H3-K9 trimethylation
glutamate biosynthetic process
regulation of store-operated calcium entry
double-strand break repair
localization
establishment of centrosome localization
cell_line
HepaRG
HuH7
PHH
medium
CDM
FBS
HepaRG_FBS
HuH7_CDM
HuH7_FBS
PHH

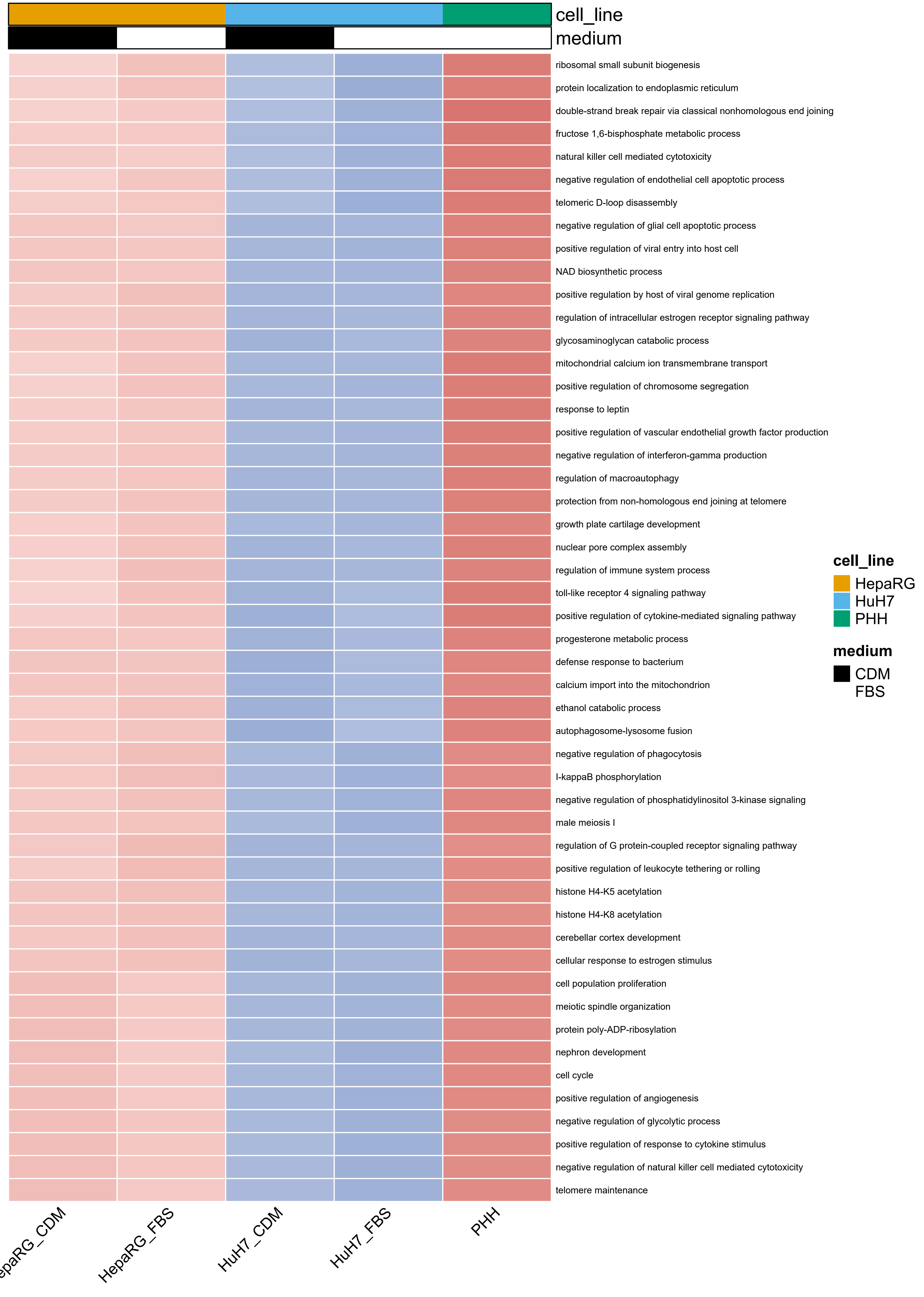
cell_line
medium
ribosomal small subunit biogenesis
protein localization to endoplasmic reticulum
double-strand break repair via classical nonhomologous end joining
fructose 1,6-bisphosphate metabolic process
natural killer cell mediated cytotoxicity
negative regulation of endothelial cell apoptotic process
telomeric D-loop disassembly
negative regulation of glial cell apoptotic process
positive regulation of viral entry into host cell
NAD biosynthetic process
positive regulation by host of viral genome replication
regulation of intracellular estrogen receptor signaling pathway
glycosaminoglycan catabolic process
mitochondrial calcium ion transmembrane transport
positive regulation of chromosome segregation
response to leptin
positive regulation of vascular endothelial growth factor production
negative regulation of interferon-gamma production
regulation of macroautophagy
protection from non-homologous end joining at telomere
growth plate cartilage development
nuclear pore complex assembly
regulation of immune system process
toll-like receptor 4 signaling pathway
positive regulation of cytokine-mediated signaling pathway
progesterone metabolic process
defense response to bacterium
calcium import into the mitochondrion
ethanol catabolic process
autophagosome-lysosome fusion
negative regulation of phagocytosis
I-kappaB phosphorylation
negative regulation of phosphatidylinositol 3-kinase signaling
male meiosis I
regulation of G protein-coupled receptor signaling pathway
positive regulation of leukocyte tethering or rolling
histone H4-K5 acetylation
histone H4-K8 acetylation
cerebellar cortex development
cellular response to estrogen stimulus
cell population proliferation
meiotic spindle organization
protein poly-ADP-ribosylation
nephron development
cell cycle
positive regulation of angiogenesis
negative regulation of glycolytic process
positive regulation of response to cytokine stimulus
negative regulation of natural killer cell mediated cytotoxicity
telomere maintenance
epaRG_CDM
HepaRG_FBS
HuH7_CDM
HuH7_FBS
PHH
cell_line
HepaRG
HuH7
PHH
medium
CDM
FBS

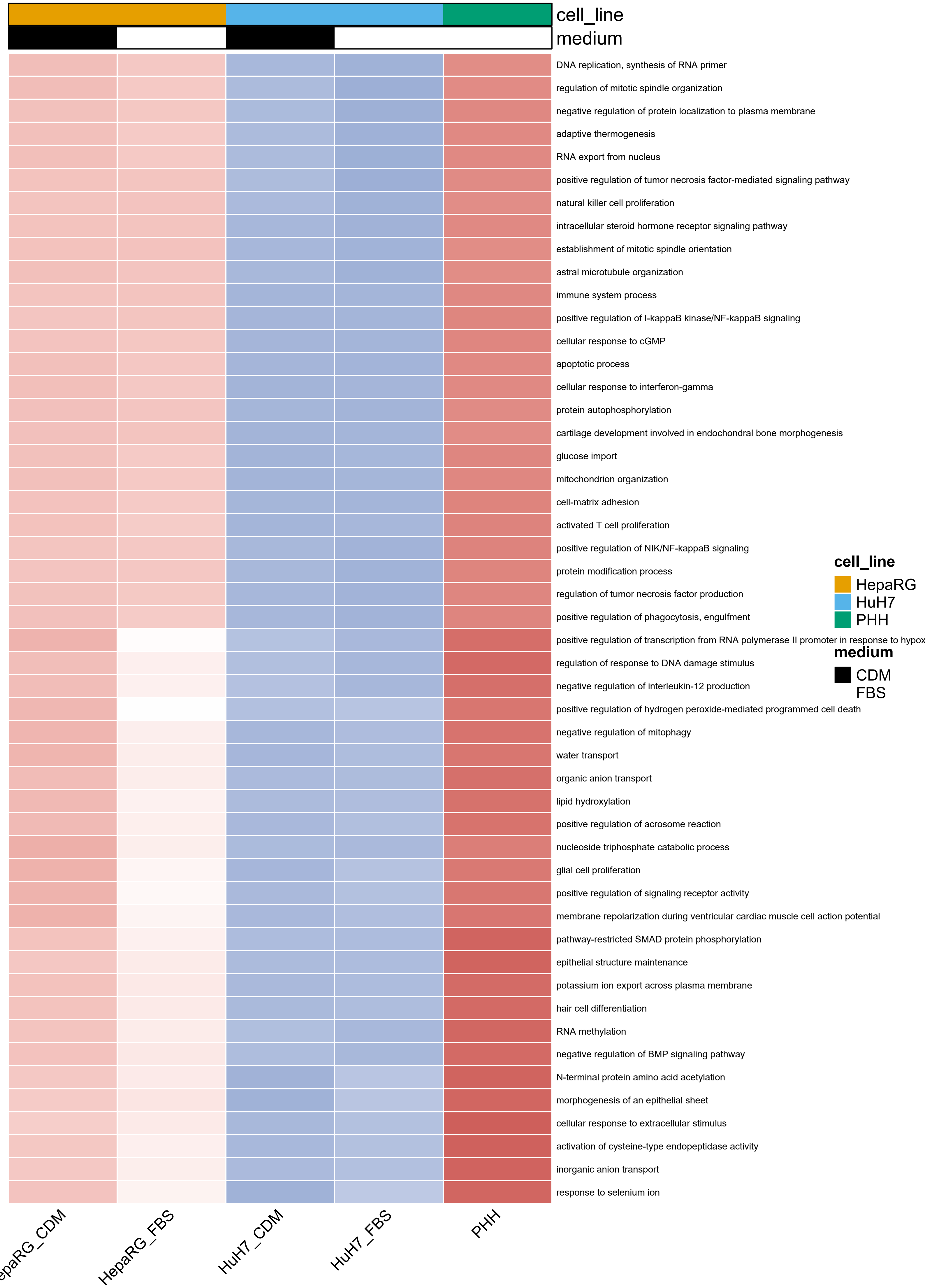
cell_line
medium
DNA replication, synthesis of RNA primer
regulation of mitotic spindle organization
negative regulation of protein localization to plasma membrane
adaptive thermogenesis
RNA export from nucleus
positive regulation of tumor necrosis factor-mediated signaling pathway
natural killer cell proliferation
intracellular steroid hormone receptor signaling pathway
establishment of mitotic spindle orientation
astral microtubule organization
immune system process
positive regulation of I-kappaB kinase/NF-kappaB signaling
cellular response to cGMP
apoptotic process
cellular response to interferon-gamma
protein autophosphorylation
cartilage development involved in endochondral bone morphogenesis
glucose import
mitochondrion organization
cell-matrix adhesion
activated T cell proliferation
positive regulation of NIK/NF-kappaB signaling
protein modification process
regulation of tumor necrosis factor production
positive regulation of phagocytosis, engulfment
positive regulation of transcription from RNA polymerase II promoter in response to hypox
regulation of response to DNA damage stimulus
negative regulation of interleukin-12 production
positive regulation of hydrogen peroxide-mediated programmed cell death
negative regulation of mitophagy
water transport
organic anion transport
lipid hydroxylation
positive regulation of acrosome reaction
nucleoside triphosphate catabolic process
glial cell proliferation
positive regulation of signaling receptor activity
membrane repolarization during ventricular cardiac muscle cell action potential
pathway-restricted SMAD protein phosphorylation
epithelial structure maintenance
potassium ion export across plasma membrane
hair cell differentiation
RNA methylation
negative regulation of BMP signaling pathway
N-terminal protein amino acid acetylation
morphogenesis of an epithelial sheet
cellular response to extracellular stimulus
activation of cysteine-type endopeptidase activity
inorganic anion transport
response to selenium ion
cell_line
HepaRG
HuH7
PHH
medium
CDM
FBS
epaRG_CDM
HepaRG_FBS
HuH7_CDM
HuH7_FBS
PHH

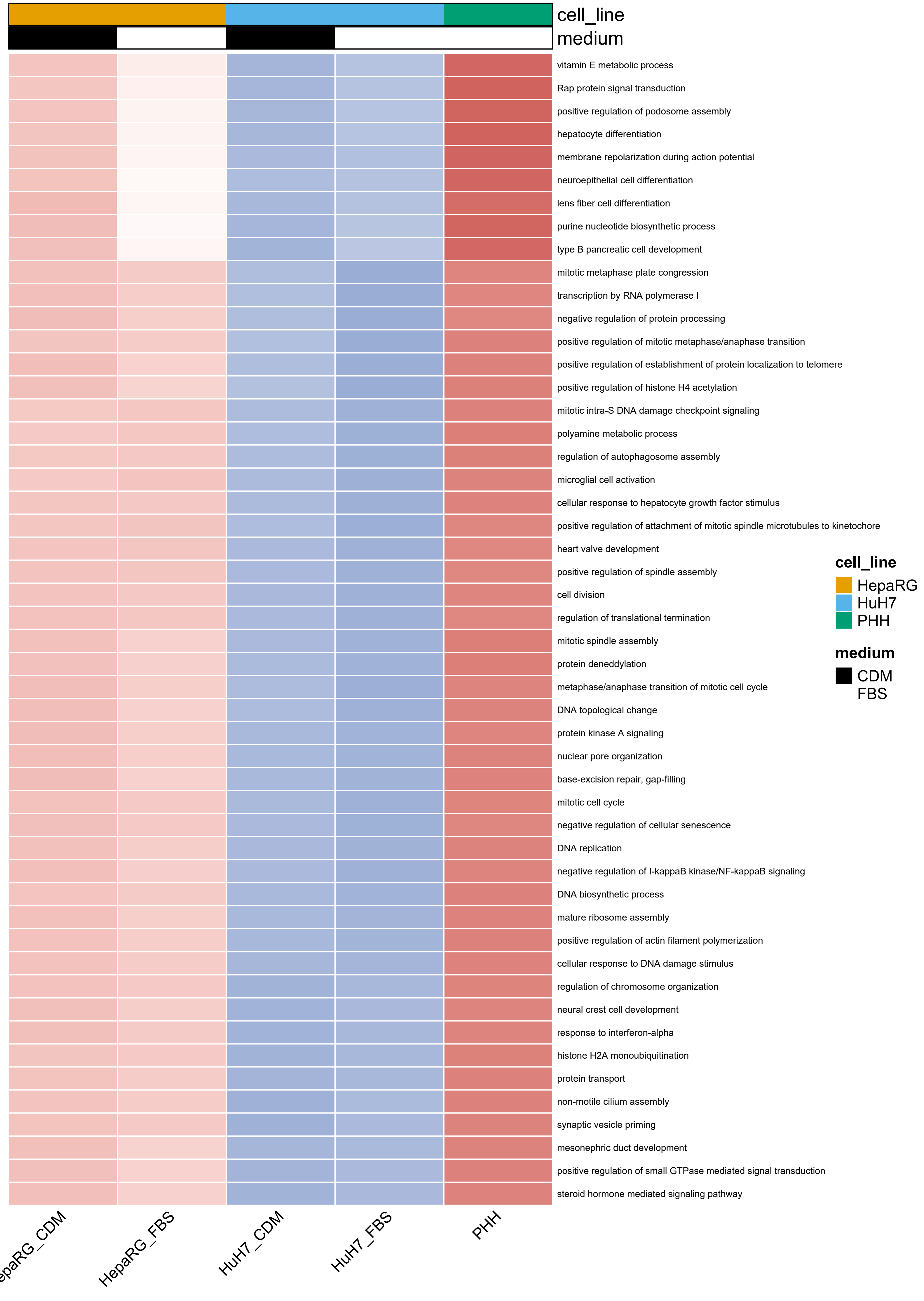
cell_line
medium
vitamin E metabolic process
Rap protein signal transduction
positive regulation of podosome assembly
hepatocyte differentiation
membrane repolarization during action potential
neuroepithelial cell differentiation
lens fiber cell differentiation
purine nucleotide biosynthetic process
type B pancreatic cell development
mitotic metaphase plate congression
transcription by RNA polymerase I
negative regulation of protein processing
positive regulation of mitotic metaphase/anaphase transition
positive regulation of establishment of protein localization to telomere
positive regulation of histone H4 acetylation
mitotic intra-S DNA damage checkpoint signaling
polyamine metabolic process
regulation of autophagosome assembly
microglial cell activation
cellular response to hepatocyte growth factor stimulus
positive regulation of attachment of mitotic spindle microtubules to kinetochore
heart valve development
positive regulation of spindle assembly
cell division
regulation of translational termination
mitotic spindle assembly
protein deneddylation
metaphase/anaphase transition of mitotic cell cycle
DNA topological change
protein kinase A signaling
nuclear pore organization
base-excision repair, gap-filling
mitotic cell cycle
negative regulation of cellular senescence
DNA replication
negative regulation of I-kappaB kinase/NF-kappaB signaling
DNA biosynthetic process
mature ribosome assembly
positive regulation of actin filament polymerization
cellular response to DNA damage stimulus
regulation of chromosome organization
neural crest cell development
response to interferon-alpha
histone H2A monoubiquitination
protein transport
non-motile cilium assembly
synaptic vesicle priming
mesonephric duct development
positive regulation of small GTPase mediated signal transduction
steroid hormone mediated signaling pathway
cell_line
HepaRG
HuH7
PHH
medium
CDM
FBS
HepaRG_FBS
HuH7_CDM
HuH7_FBS
PHH

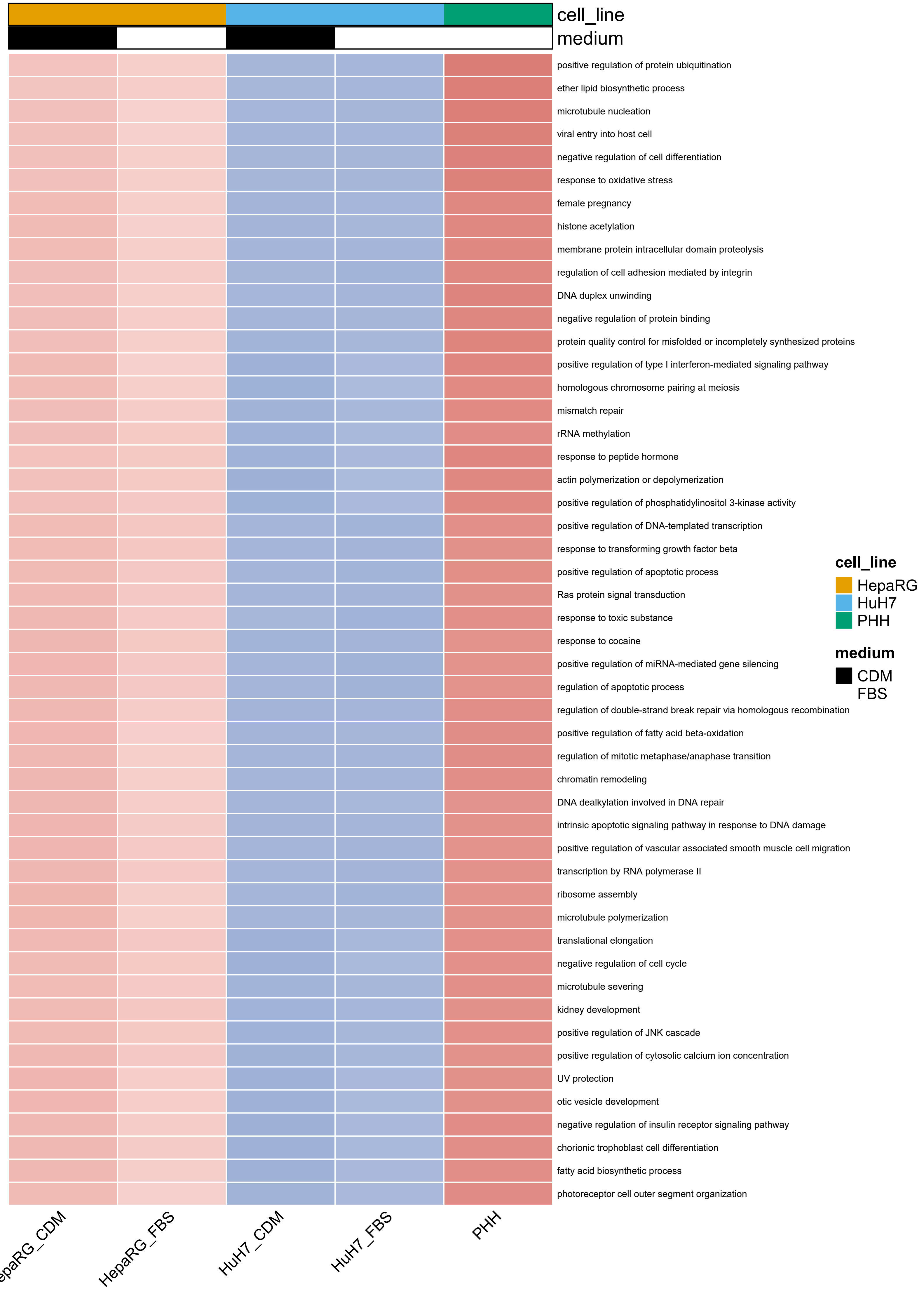
cell_line
medium
positive regulation of protein ubiquitination
ether lipid biosynthetic process
microtubule nucleation
viral entry into host cell
negative regulation of cell differentiation
response to oxidative stress
female pregnancy
histone acetylation
membrane protein intracellular domain proteolysis
regulation of cell adhesion mediated by integrin
DNA duplex unwinding
negative regulation of protein binding
protein quality control for misfolded or incompletely synthesized proteins
positive regulation of type I interferon-mediated signaling pathway
homologous chromosome pairing at meiosis
mismatch repair
rRNA methylation
response to peptide hormone
actin polymerization or depolymerization
positive regulation of phosphatidylinositol 3-kinase activity
positive regulation of DNA-templated transcription
response to transforming growth factor beta
positive regulation of apoptotic process
Ras protein signal transduction
response to toxic substance
response to cocaine
positive regulation of miRNA-mediated gene silencing
regulation of apoptotic process
regulation of double-strand break repair via homologous recombination
positive regulation of fatty acid beta-oxidation
regulation of mitotic metaphase/anaphase transition
chromatin remodeling
DNA dealkylation involved in DNA repair
intrinsic apoptotic signaling pathway in response to DNA damage
positive regulation of vascular associated smooth muscle cell migration
transcription by RNA polymerase II
ribosome assembly
microtubule polymerization
translational elongation
negative regulation of cell cycle
microtubule severing
kidney development
positive regulation of JNK cascade
positive regulation of cytosolic calcium ion concentration
UV protection
otic vesicle development
negative regulation of insulin receptor signaling pathway
chorionic trophoblast cell differentiation
fatty acid biosynthetic process
photoreceptor cell outer segment organization
cell_line
HepaRG
HuH7
PHH
medium
CDM
FBS
HepaRG_FBS
HuH7_CDM
HuH7_FBS
PHH

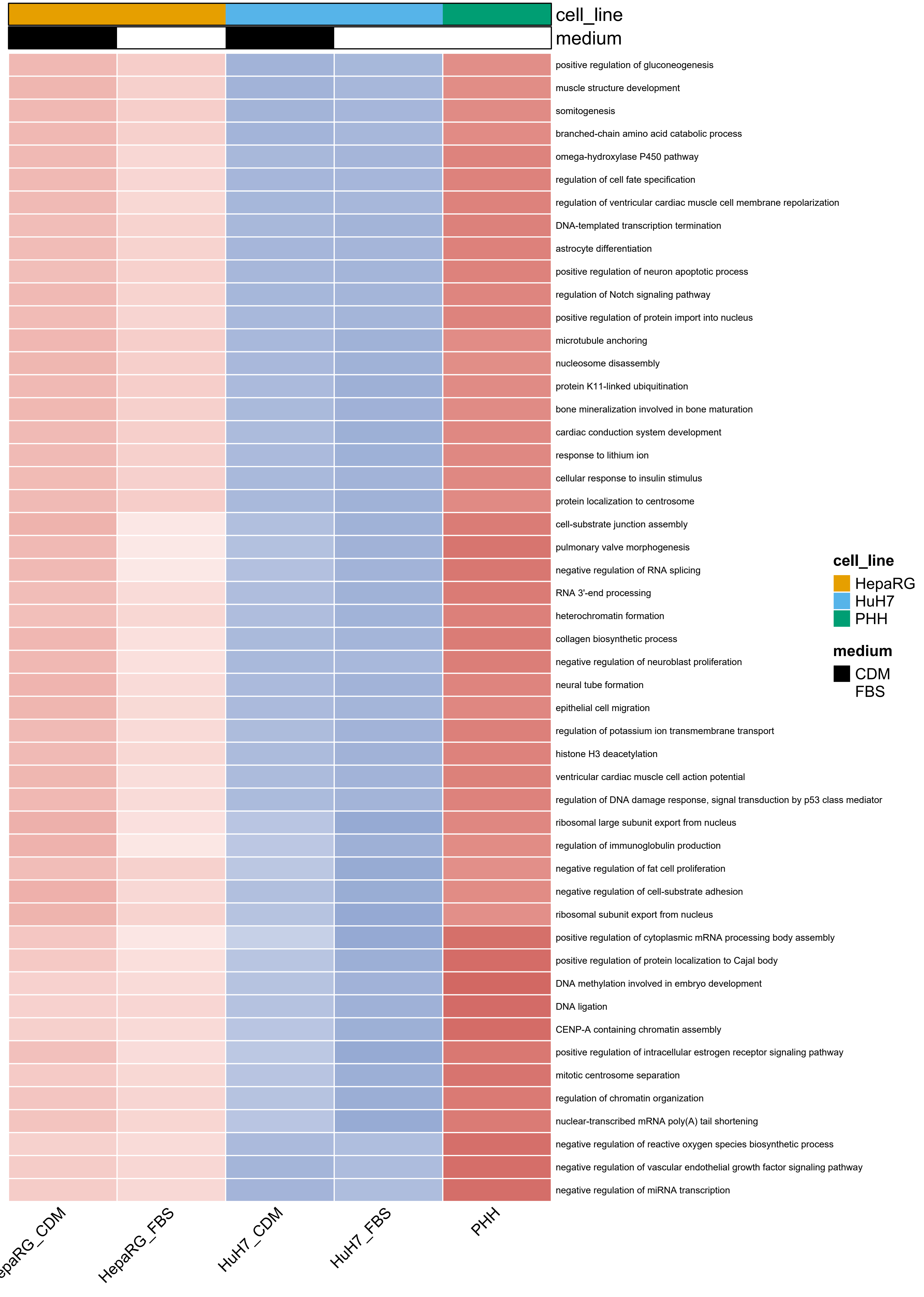
cell_line
medium
positive regulation of gluconeogenesis
muscle structure development
somitogenesis
branched-chain amino acid catabolic process
omega-hydroxylase P450 pathway
regulation of cell fate specification
regulation of ventricular cardiac muscle cell membrane repolarization
DNA-templated transcription termination
astrocyte differentiation
positive regulation of neuron apoptotic process
regulation of Notch signaling pathway
positive regulation of protein import into nucleus
microtubule anchoring
nucleosome disassembly
protein K11-linked ubiquitination
bone mineralization involved in bone maturation
cardiac conduction system development
response to lithium ion
cellular response to insulin stimulus
protein localization to centrosome
cell-substrate junction assembly
pulmonary valve morphogenesis
negative regulation of RNA splicing
RNA 3'-end processing
heterochromatin formation
collagen biosynthetic process
negative regulation of neuroblast proliferation
neural tube formation
epithelial cell migration
regulation of potassium ion transmembrane transport
histone H3 deacetylation
ventricular cardiac muscle cell action potential
regulation of DNA damage response, signal transduction by p53 class mediator
ribosomal large subunit export from nucleus
regulation of immunoglobulin production
negative regulation of fat cell proliferation
negative regulation of cell-substrate adhesion
ribosomal subunit export from nucleus
positive regulation of cytoplasmic mRNA processing body assembly
positive regulation of protein localization to Cajal body
DNA methylation involved in embryo development
DNA ligation
CENP-A containing chromatin assembly
positive regulation of intracellular estrogen receptor signaling pathway
mitotic centrosome separation
regulation of chromatin organization
nuclear-transcribed mRNA poly(A) tail shortening
negative regulation of reactive oxygen species biosynthetic process
negative regulation of vascular endothelial growth factor signaling pathway
negative regulation of miRNA transcription
HepaRG_FBS
HuH7_CDM
HuH7_FBS
PHH
cell_line
HepaRG
HuH7
PHH
medium
CDM
FBS

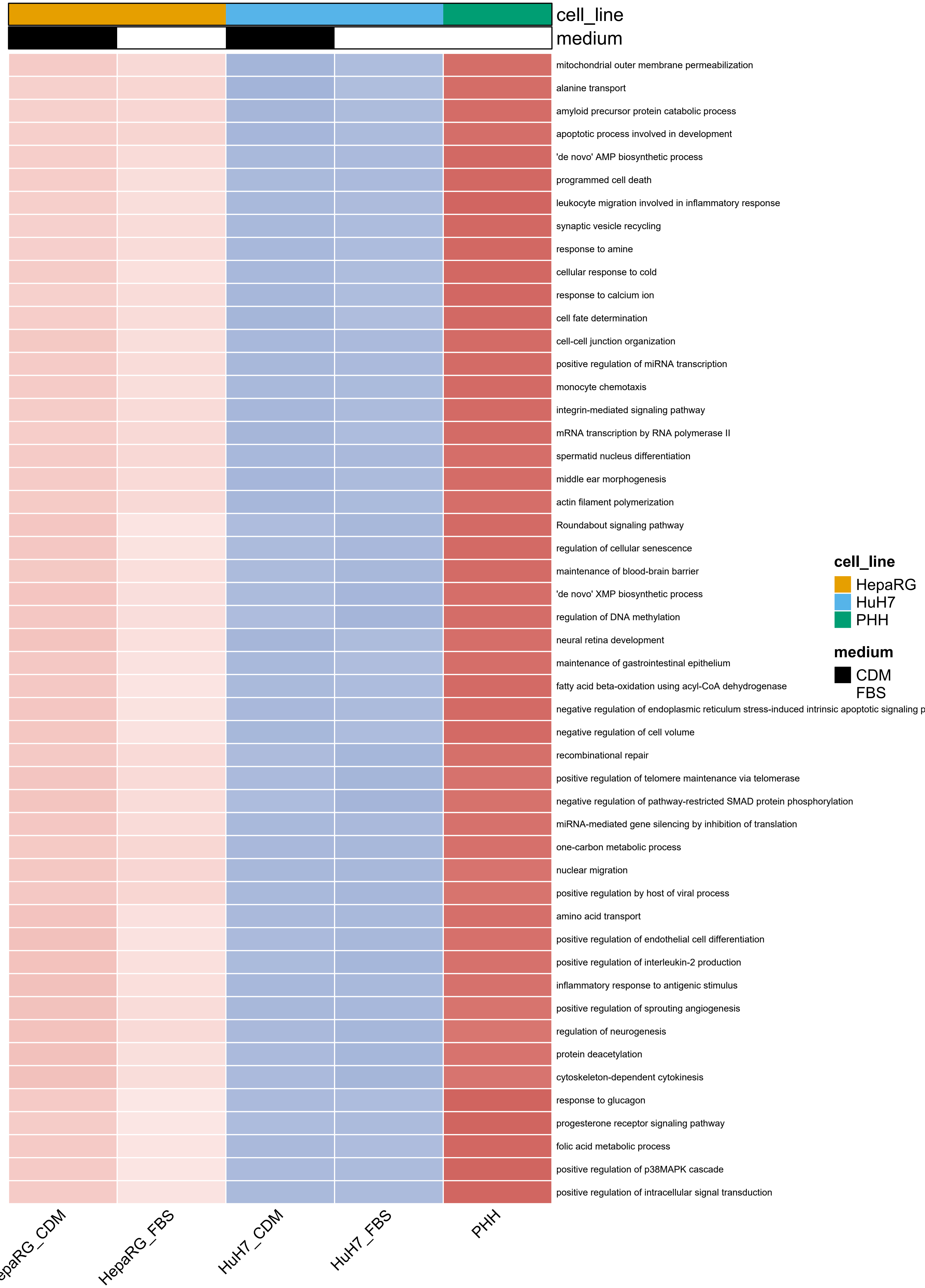
cell_line
medium
mitochondrial outer membrane permeabilization
alanine transport
amyloid precursor protein catabolic process
apoptotic process involved in development
'de novo' AMP biosynthetic process
programmed cell death
leukocyte migration involved in inflammatory response
synaptic vesicle recycling
response to amine
cellular response to cold
response to calcium ion
cell fate determination
cell-cell junction organization
positive regulation of miRNA transcription
monocyte chemotaxis
integrin-mediated signaling pathway
mRNA transcription by RNA polymerase II
spermatid nucleus differentiation
middle ear morphogenesis
actin filament polymerization
Roundabout signaling pathway
regulation of cellular senescence
maintenance of blood-brain barrier
'de novo' XMP biosynthetic process
regulation of DNA methylation
neural retina development
maintenance of gastrointestinal epithelium
fatty acid beta-oxidation using acyl-CoA dehydrogenase
negative regulation of endoplasmic reticulum stress-induced intrinsic apoptotic signaling p
negative regulation of cell volume
recombinational repair
positive regulation of telomere maintenance via telomerase
negative regulation of pathway-restricted SMAD protein phosphorylation
miRNA-mediated gene silencing by inhibition of translation
one-carbon metabolic process
nuclear migration
positive regulation by host of viral process
amino acid transport
positive regulation of endothelial cell differentiation
positive regulation of interleukin-2 production
inflammatory response to antigenic stimulus
positive regulation of sprouting angiogenesis
regulation of neurogenesis
protein deacetylation
cytoskeleton-dependent cytokinesis
response to glucagon
progesterone receptor signaling pathway
folic acid metabolic process
positive regulation of p38MAPK cascade
positive regulation of intracellular signal transduction
epaRG_CDM
HepaRG_FBS
HuH7_CDM
HuH7_FBS
PHH
cell_line
HepaRG
HuH7
PHH
medium
CDM
FBS

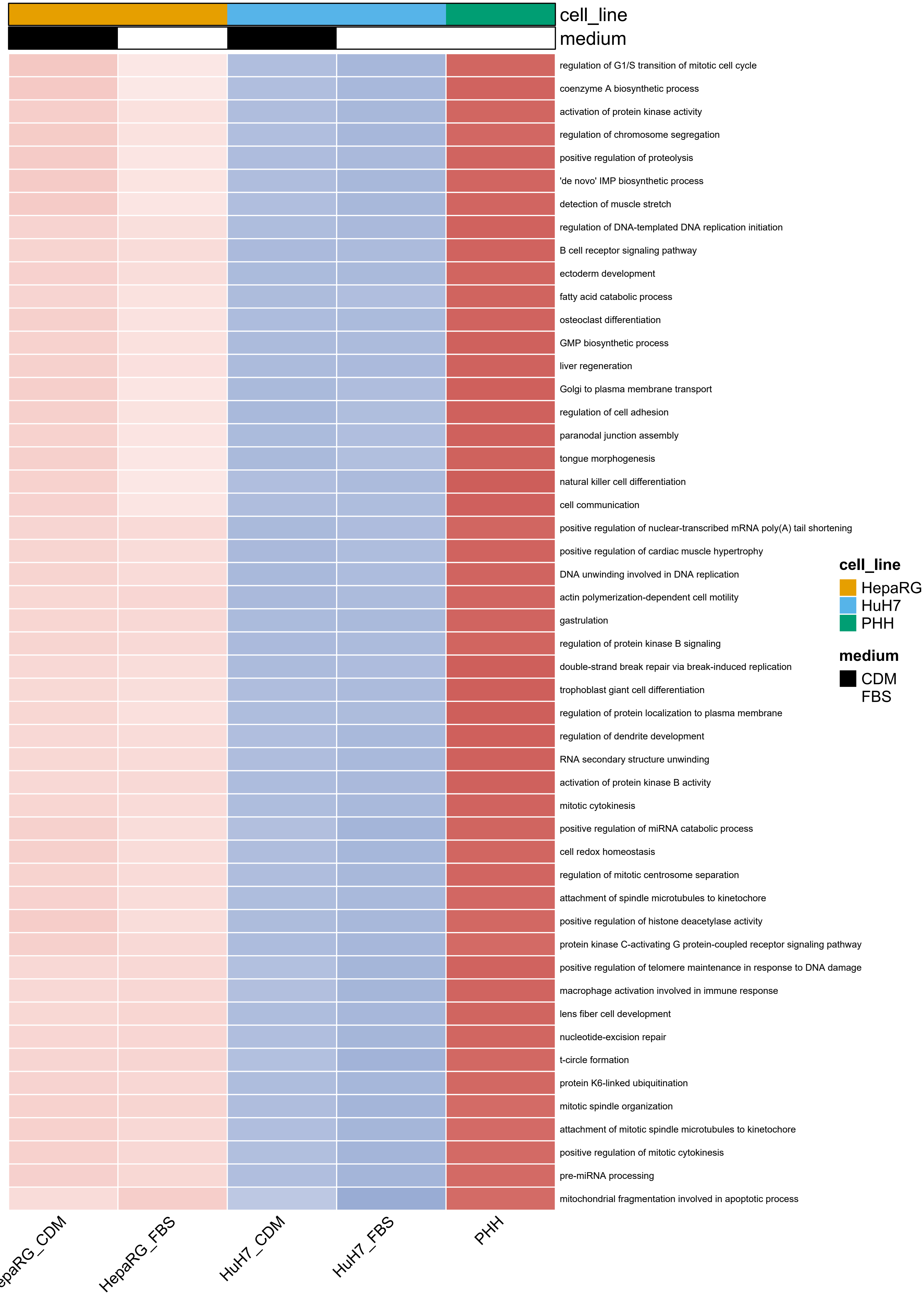
cell_line
medium
regulation of G1/S transition of mitotic cell cycle
coenzyme A biosynthetic process
activation of protein kinase activity
regulation of chromosome segregation
positive regulation of proteolysis
'de novo' IMP biosynthetic process
detection of muscle stretch
regulation of DNA-templated DNA replication initiation
B cell receptor signaling pathway
ectoderm development
fatty acid catabolic process
osteoclast differentiation
GMP biosynthetic process
liver regeneration
Golgi to plasma membrane transport
regulation of cell adhesion
paranodal junction assembly
tongue morphogenesis
natural killer cell differentiation
cell communication
positive regulation of nuclear-transcribed mRNA poly(A) tail shortening
positive regulation of cardiac muscle hypertrophy
DNA unwinding involved in DNA replication
actin polymerization-dependent cell motility
gastrulation
regulation of protein kinase B signaling
double-strand break repair via break-induced replication
trophoblast giant cell differentiation
regulation of protein localization to plasma membrane
regulation of dendrite development
RNA secondary structure unwinding
activation of protein kinase B activity
mitotic cytokinesis
positive regulation of miRNA catabolic process
cell redox homeostasis
regulation of mitotic centrosome separation
attachment of spindle microtubules to kinetochore
positive regulation of histone deacetylase activity
protein kinase C-activating G protein-coupled receptor signaling pathway
positive regulation of telomere maintenance in response to DNA damage
macrophage activation involved in immune response
lens fiber cell development
nucleotide-excision repair
t-circle formation
protein K6-linked ubiquitination
mitotic spindle organization
attachment of mitotic spindle microtubules to kinetochore
positive regulation of mitotic cytokinesis
pre-miRNA processing
mitochondrial fragmentation involved in apoptotic process
cell_line
HepaRG
HuH7
PHH
medium
CDM
FBS
HepaRG_FBS
HuH7_CDM
HuH7_FBS
PHH

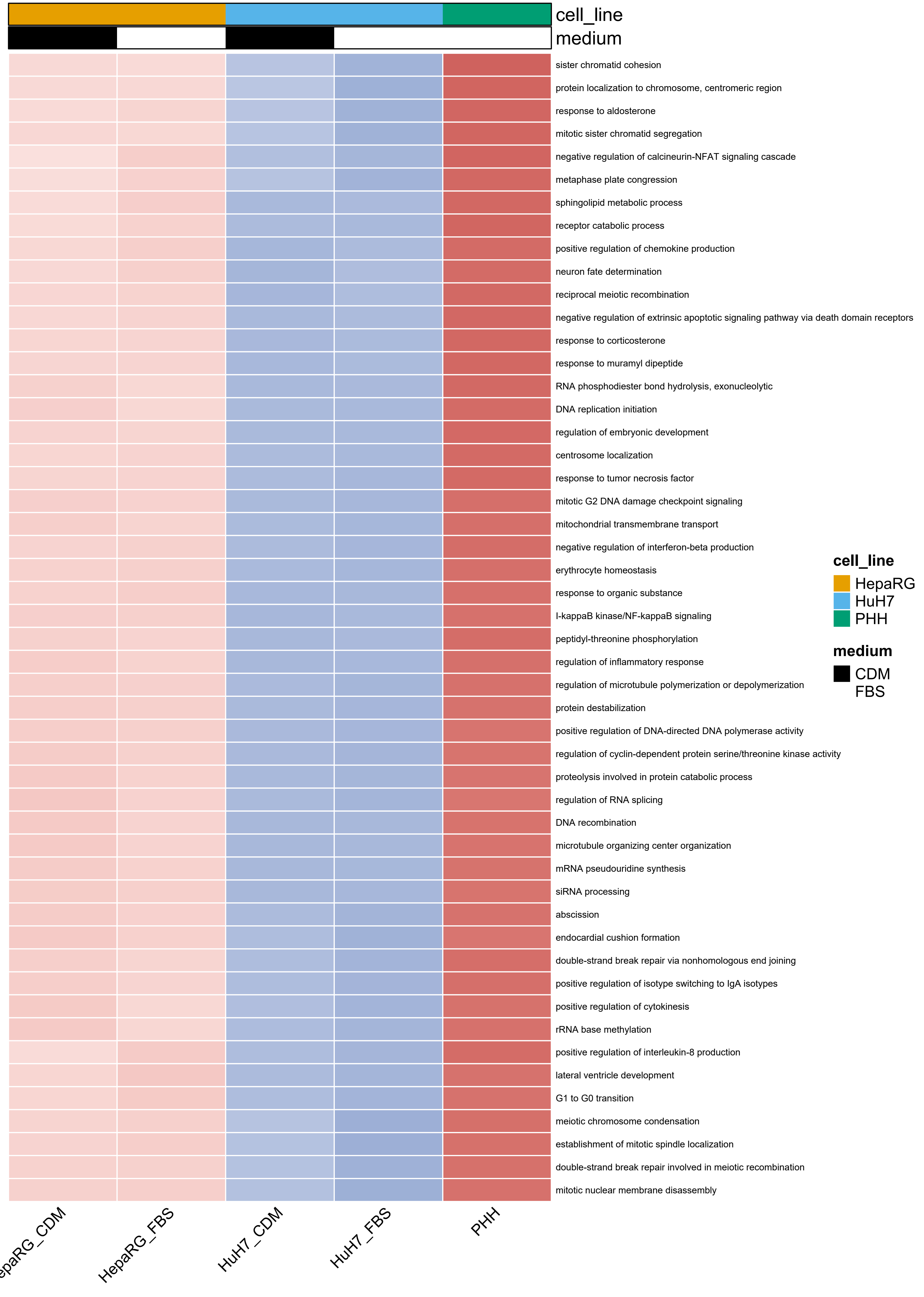
cell_line
medium
sister chromatid cohesion
protein localization to chromosome, centromeric region
response to aldosterone
mitotic sister chromatid segregation
negative regulation of calcineurin-NFAT signaling cascade
metaphase plate congression
sphingolipid metabolic process
receptor catabolic process
positive regulation of chemokine production
neuron fate determination
reciprocal meiotic recombination
negative regulation of extrinsic apoptotic signaling pathway via death domain receptors
response to corticosterone
response to muramyl dipeptide
RNA phosphodiester bond hydrolysis, exonucleolytic
DNA replication initiation
regulation of embryonic development
centrosome localization
response to tumor necrosis factor
mitotic G2 DNA damage checkpoint signaling
mitochondrial transmembrane transport
negative regulation of interferon-beta production
erythrocyte homeostasis
response to organic substance
I-kappaB kinase/NF-kappaB signaling
peptidyl-threonine phosphorylation
regulation of inflammatory response
regulation of microtubule polymerization or depolymerization
protein destabilization
positive regulation of DNA-directed DNA polymerase activity
regulation of cyclin-dependent protein serine/threonine kinase activity
proteolysis involved in protein catabolic process
regulation of RNA splicing
DNA recombination
microtubule organizing center organization
mRNA pseudouridine synthesis
siRNA processing
abscission
endocardial cushion formation
double-strand break repair via nonhomologous end joining
positive regulation of isotype switching to IgA isotypes
positive regulation of cytokinesis
rRNA base methylation
positive regulation of interleukin-8 production
lateral ventricle development
G1 to G0 transition
meiotic chromosome condensation
establishment of mitotic spindle localization
double-strand break repair involved in meiotic recombination
mitotic nuclear membrane disassembly
cell_line
HepaRG
HuH7
PHH
medium
CDM
FBS
HepaRG_FBS
HuH7_CDM
HuH7_FBS
PHH

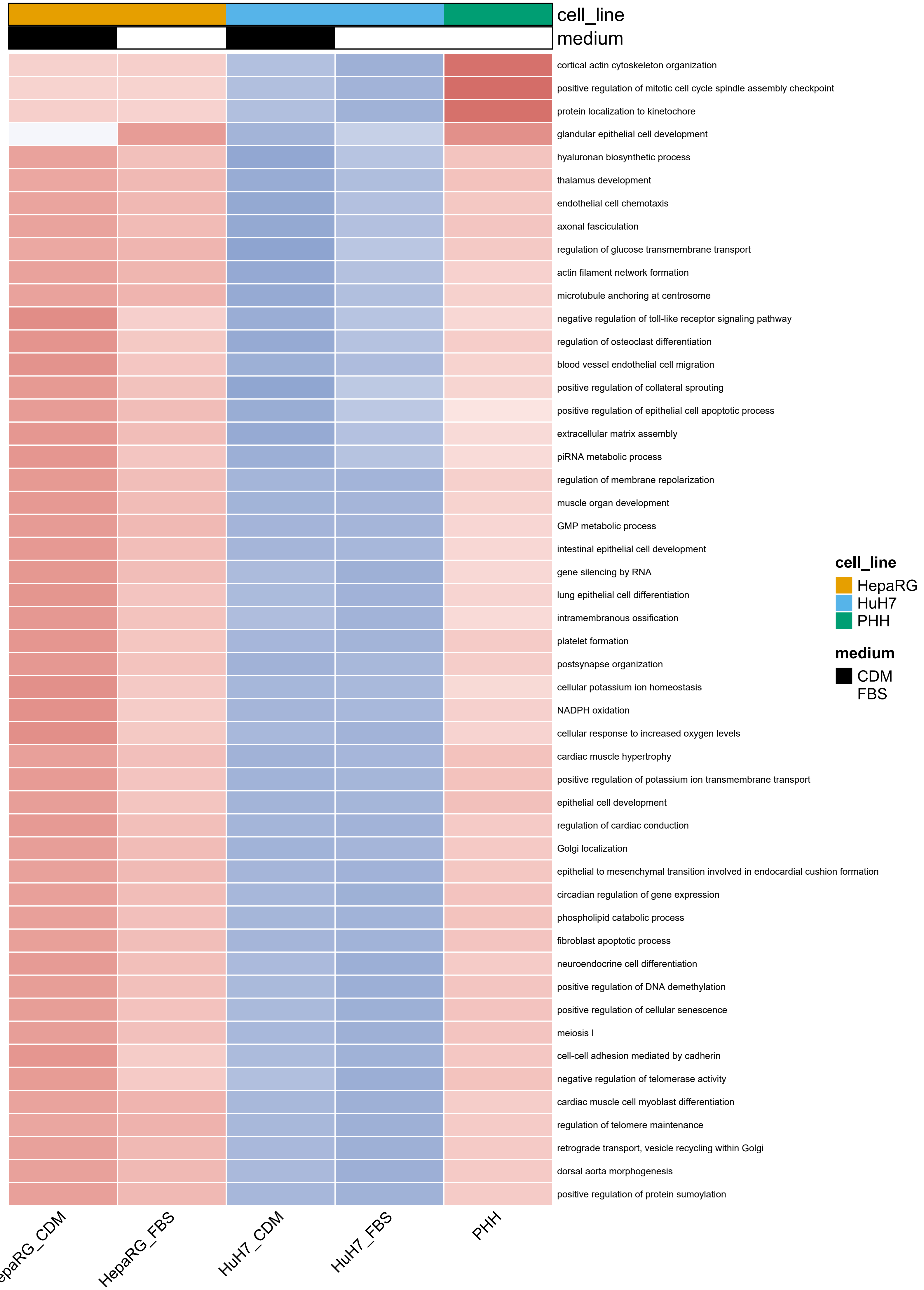
cell_line
medium
cortical actin cytoskeleton organization
positive regulation of mitotic cell cycle spindle assembly checkpoint
protein localization to kinetochore
glandular epithelial cell development
hyaluronan biosynthetic process
thalamus development
endothelial cell chemotaxis
axonal fasciculation
regulation of glucose transmembrane transport
actin filament network formation
microtubule anchoring at centrosome
negative regulation of toll-like receptor signaling pathway
regulation of osteoclast differentiation
blood vessel endothelial cell migration
positive regulation of collateral sprouting
positive regulation of epithelial cell apoptotic process
extracellular matrix assembly
piRNA metabolic process
regulation of membrane repolarization
muscle organ development
GMP metabolic process
intestinal epithelial cell development
gene silencing by RNA
lung epithelial cell differentiation
intramembranous ossification
platelet formation
postsynapse organization
cellular potassium ion homeostasis
NADPH oxidation
cellular response to increased oxygen levels
cardiac muscle hypertrophy
positive regulation of potassium ion transmembrane transport
epithelial cell development
regulation of cardiac conduction
Golgi localization
epithelial to mesenchymal transition involved in endocardial cushion formation
circadian regulation of gene expression
phospholipid catabolic process
fibroblast apoptotic process
neuroendocrine cell differentiation
positive regulation of DNA demethylation
positive regulation of cellular senescence
meiosis I
cell-cell adhesion mediated by cadherin
negative regulation of telomerase activity
cardiac muscle cell myoblast differentiation
regulation of telomere maintenance
retrograde transport, vesicle recycling within Golgi
dorsal aorta morphogenesis
positive regulation of protein sumoylation
HepaRG_FBS
HuH7_CDM
HuH7_FBS
PHH
cell_line
HepaRG
HuH7
PHH
medium
CDM
FBS

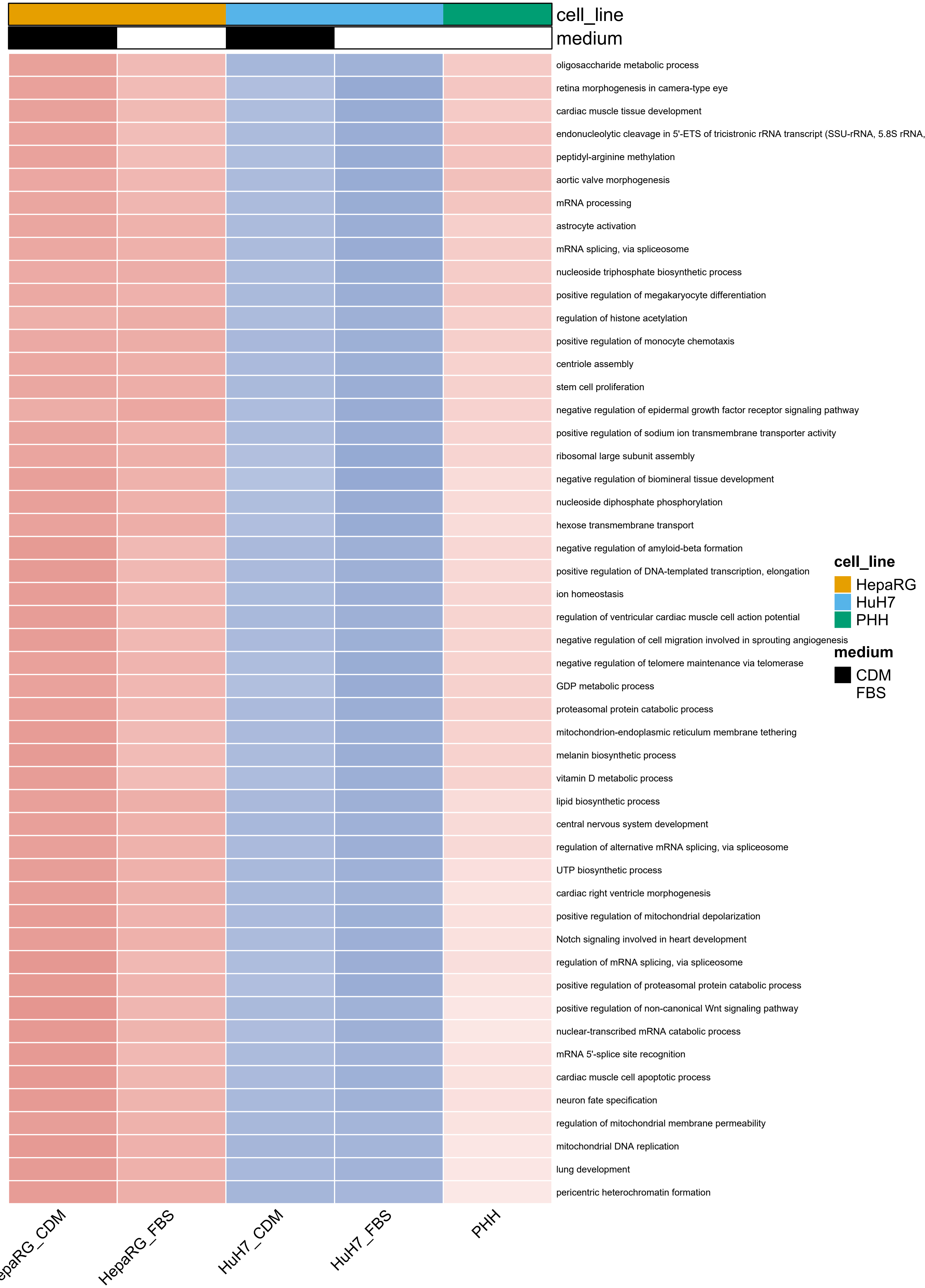
cell_line
medium
oligosaccharide metabolic process
retina morphogenesis in camera-type eye
cardiac muscle tissue development
endonucleolytic cleavage in 5'-ETS of tricistronic rRNA transcript (SSU-rRNA, 5.8S rRNA,
peptidyl-arginine methylation
aortic valve morphogenesis
mRNA processing
astrocyte activation
mRNA splicing, via spliceosome
nucleoside triphosphate biosynthetic process
positive regulation of megakaryocyte differentiation
regulation of histone acetylation
positive regulation of monocyte chemotaxis
centriole assembly
stem cell proliferation
negative regulation of epidermal growth factor receptor signaling pathway
positive regulation of sodium ion transmembrane transporter activity
ribosomal large subunit assembly
negative regulation of biomineral tissue development
nucleoside diphosphate phosphorylation
hexose transmembrane transport
negative regulation of amyloid-beta formation
positive regulation of DNA-templated transcription, elongation
ion homeostasis
regulation of ventricular cardiac muscle cell action potential
negative regulation of cell migration involved in sprouting angiogenesis
negative regulation of telomere maintenance via telomerase
GDP metabolic process
proteasomal protein catabolic process
mitochondrion-endoplasmic reticulum membrane tethering
melanin biosynthetic process
vitamin D metabolic process
lipid biosynthetic process
central nervous system development
regulation of alternative mRNA splicing, via spliceosome
UTP biosynthetic process
cardiac right ventricle morphogenesis
positive regulation of mitochondrial depolarization
Notch signaling involved in heart development
regulation of mRNA splicing, via spliceosome
positive regulation of proteasomal protein catabolic process
positive regulation of non-canonical Wnt signaling pathway
nuclear-transcribed mRNA catabolic process
mRNA 5'-splice site recognition
cardiac muscle cell apoptotic process
neuron fate specification
regulation of mitochondrial membrane permeability
mitochondrial DNA replication
lung development
pericentric heterochromatin formation
cell_line
HepaRG
HuH7
PHH
medium
CDM
FBS
epaRG_CDM
HepaRG_FBS
HuH7_CDM
HuH7_FBS
PHH

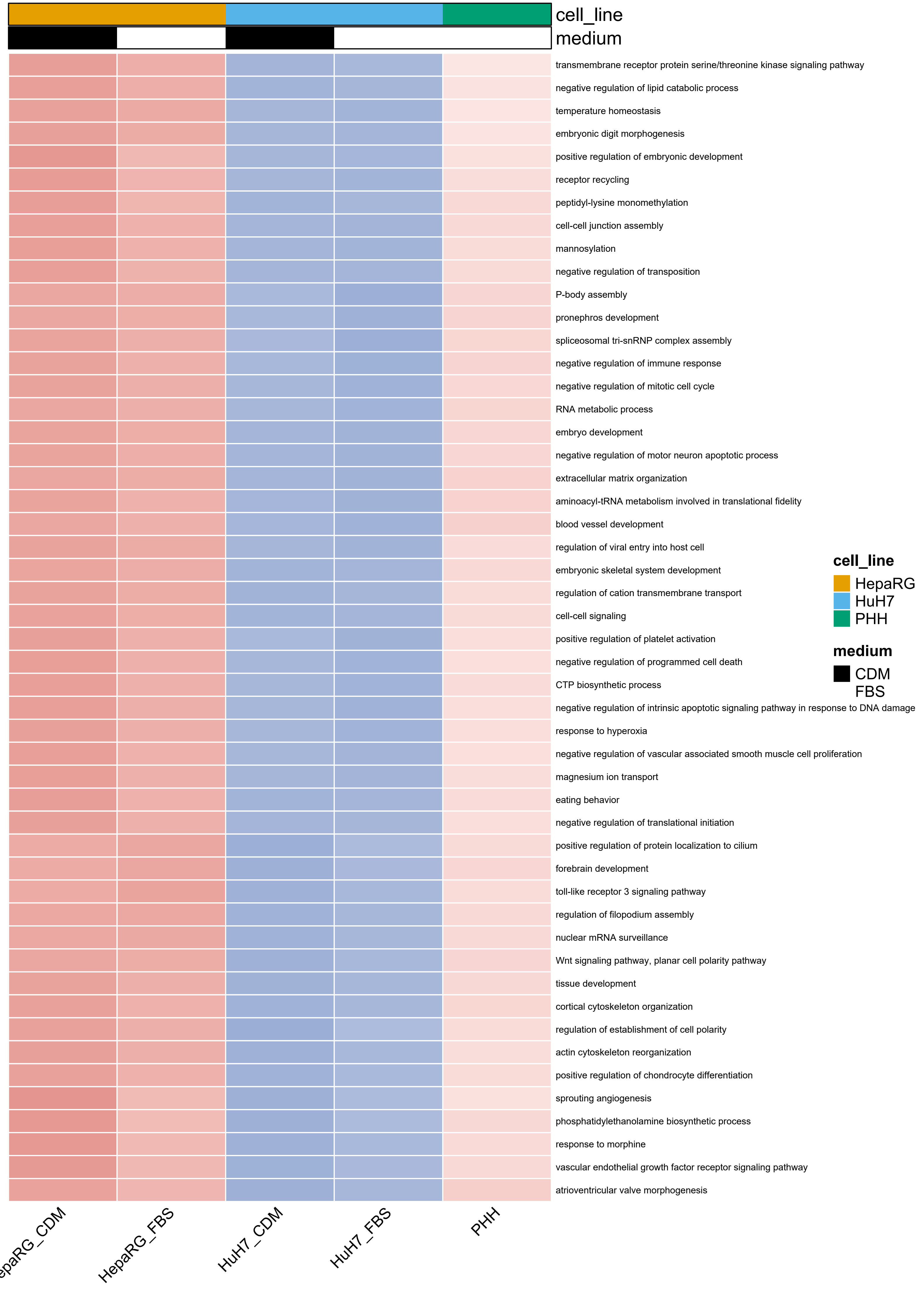
cell_line
medium
transmembrane receptor protein serine/threonine kinase signaling pathway
negative regulation of lipid catabolic process
temperature homeostasis
embryonic digit morphogenesis
positive regulation of embryonic development
receptor recycling
peptidyl-lysine monomethylation
cell-cell junction assembly
mannosylation
negative regulation of transposition
P-body assembly
pronephros development
spliceosomal tri-snRNP complex assembly
negative regulation of immune response
negative regulation of mitotic cell cycle
RNA metabolic process
embryo development
negative regulation of motor neuron apoptotic process
extracellular matrix organization
aminoacyl-tRNA metabolism involved in translational fidelity
blood vessel development
regulation of viral entry into host cell
embryonic skeletal system development
regulation of cation transmembrane transport
cell-cell signaling
positive regulation of platelet activation
negative regulation of programmed cell death
CTP biosynthetic process
negative regulation of intrinsic apoptotic signaling pathway in response to DNA damage
response to hyperoxia
negative regulation of vascular associated smooth muscle cell proliferation
magnesium ion transport
eating behavior
negative regulation of translational initiation
positive regulation of protein localization to cilium
forebrain development
toll-like receptor 3 signaling pathway
regulation of filopodium assembly
nuclear mRNA surveillance
Wnt signaling pathway, planar cell polarity pathway
tissue development
cortical cytoskeleton organization
regulation of establishment of cell polarity
actin cytoskeleton reorganization
positive regulation of chondrocyte differentiation
sprouting angiogenesis
phosphatidylethanolamine biosynthetic process
response to morphine
vascular endothelial growth factor receptor signaling pathway
atrioventricular valve morphogenesis
cell_line
HepaRG
HuH7
PHH
medium
CDM
FBS
HepaRG_FBS
HuH7_CDM
HuH7_FBS
PHH

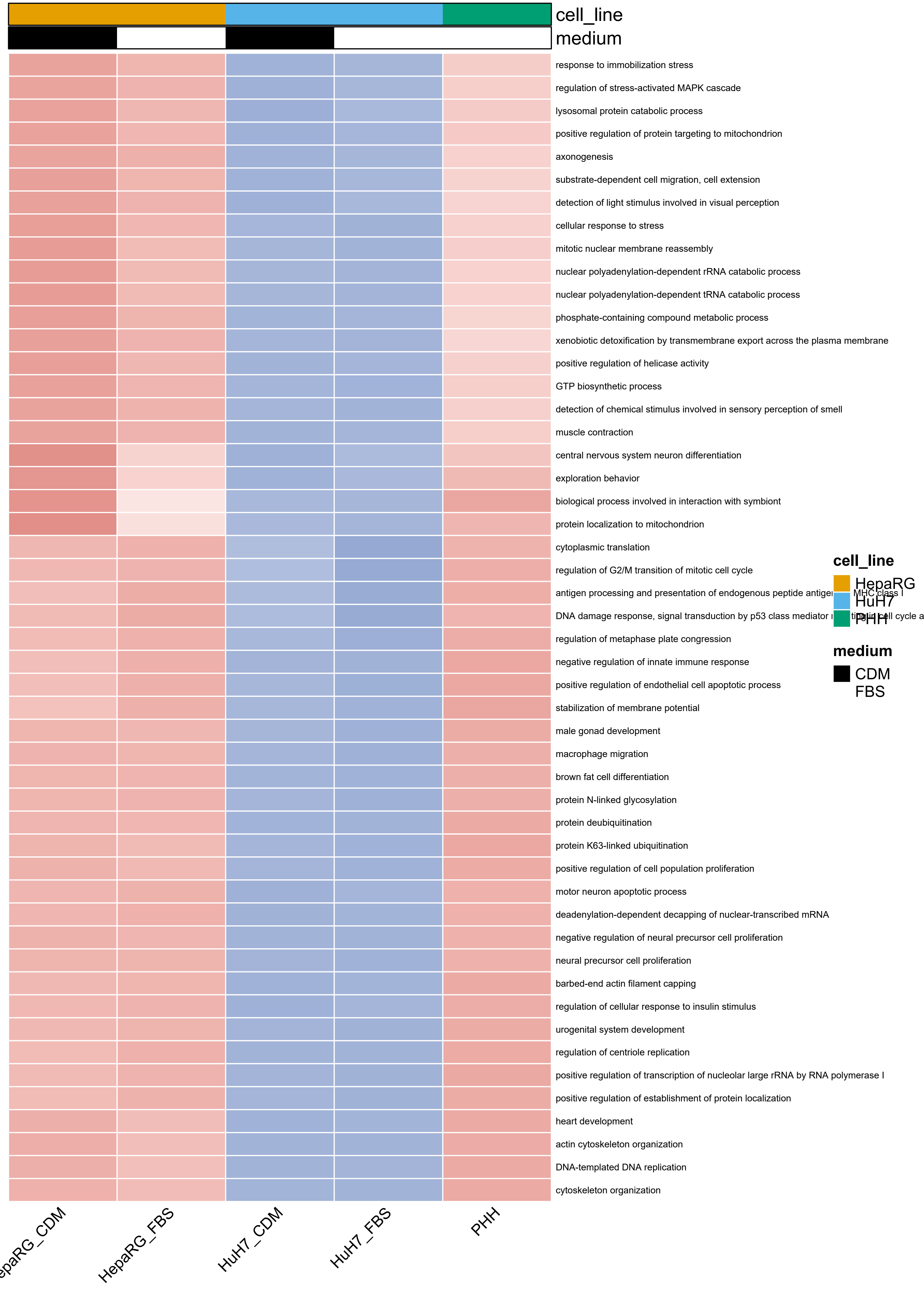

cell_line
medium
response to immobilization stress
regulation of stress-activated MAPK cascade
lysosomal protein catabolic process
positive regulation of protein targeting to mitochondrion
axonogenesis
substrate-dependent cell migration, cell extension
detection of light stimulus involved in visual perception
cellular response to stress
mitotic nuclear membrane reassembly
nuclear polyadenylation-dependent rRNA catabolic process
nuclear polyadenylation-dependent tRNA catabolic process
phosphate-containing compound metabolic process
xenobiotic detoxification by transmembrane export across the plasma membrane
positive regulation of helicase activity
GTP biosynthetic process
detection of chemical stimulus involved in sensory perception of smell
muscle contraction
central nervous system neuron differentiation
exploration behavior
biological process involved in interaction with symbiont
protein localization to mitochondrion
cytoplasmic translation
regulation of G2/M transition of mitotic cell cycle
antigen processing and presentation of endogenous peptide antige
MHC class I
DNA damage response, signal transduction by p53 class mediator
regulation of metaphase plate congression
negative regulation of innate immune response
positive regulation of endothelial cell apoptotic process
stabilization of membrane potential
male gonad development
macrophage migration
brown fat cell differentiation
protein N-linked glycosylation
protein deubiquitination
protein K63-linked ubiquitination
positive regulation of cell population proliferation
motor neuron apoptotic process
deadenylation-dependent decapping of nuclear-transcribed mRNA
negative regulation of neural precursor cell proliferation
neural precursor cell proliferation
barbed-end actin filament capping
regulation of cellular response to insulin stimulus
urogenital system development
regulation of centriole replication
positive regulation of transcription of nucleolar large rRNA by RNA polymerase I
positive regulation of establishment of protein localization
heart development
actin cytoskeleton organization
DNA-templated DNA replication
cytoskeleton organization
epaRG_CDM
HepaRG_FBS
HuH7_CDM
HuH7_FBS
PHH
cell_line
HepaRG
HuH7
PHH
medium
CDM
FBS

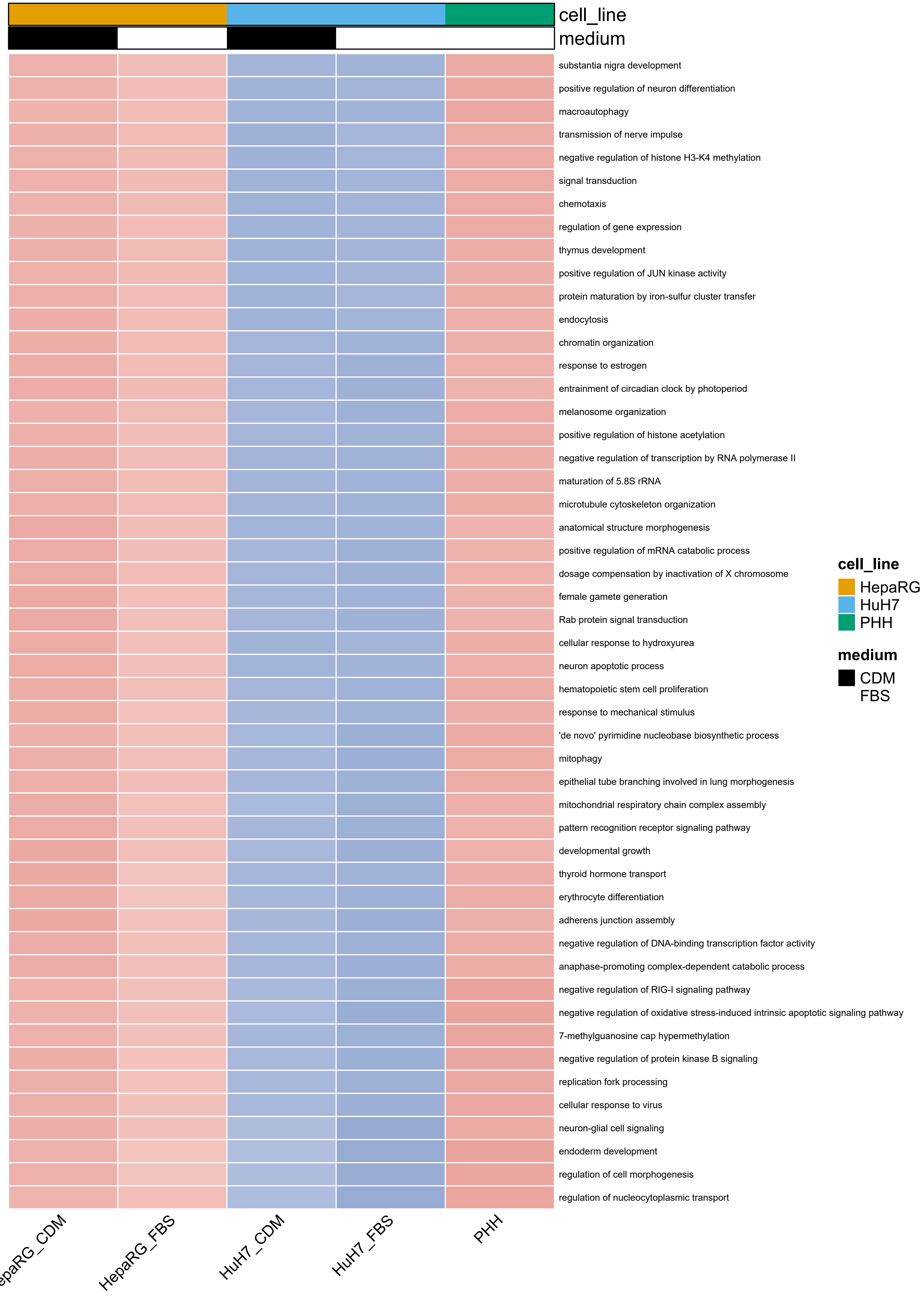
cell_line
medium
substantia nigra development
positive regulation of neuron differentiation
macroautophagy
transmission of nerve impulse
negative regulation of histone H3-K4 methylation
signal transduction
chemotaxis
regulation of gene expression
thymus development
positive regulation of JUN kinase activity
protein maturation by iron-sulfur cluster transfer
endocytosis
chromatin organization
response to estrogen
entrainment of circadian clock by photoperiod
melanosome organization
positive regulation of histone acetylation
negative regulation of transcription by RNA polymerase II
maturation of 5.8S rRNA
microtubule cytoskeleton organization
anatomical structure morphogenesis
positive regulation of mRNA catabolic process
dosage compensation by inactivation of X chromosome
female gamete generation
Rab protein signal transduction
cellular response to hydroxyurea
neuron apoptotic process
hematopoietic stem cell proliferation
response to mechanical stimulus
'de novo' pyrimidine nucleobase biosynthetic process
mitophagy
epithelial tube branching involved in lung morphogenesis
mitochondrial respiratory chain complex assembly
pattern recognition receptor signaling pathway
developmental growth
thyroid hormone transport
erythrocyte differentiation
adherens junction assembly
negative regulation of DNA-binding transcription factor activity
anaphase-promoting complex-dependent catabolic process
negative regulation of RIG-I signaling pathway
negative regulation of oxidative stress-induced intrinsic apoptotic signaling pathway
7-methylguanosine cap hypermethylation
negative regulation of protein kinase B signaling
replication fork processing
cellular response to virus
neuron-glial cell signaling
endoderm development
regulation of cell morphogenesis
regulation of nucleocytoplasmic transport
epaRG_CDM
HepaRG_FBS
HuH7_CDM
HuH7_FBS
PHH
cell_line
HepaRG
HuH7
PHH
medium
CDM
FBS

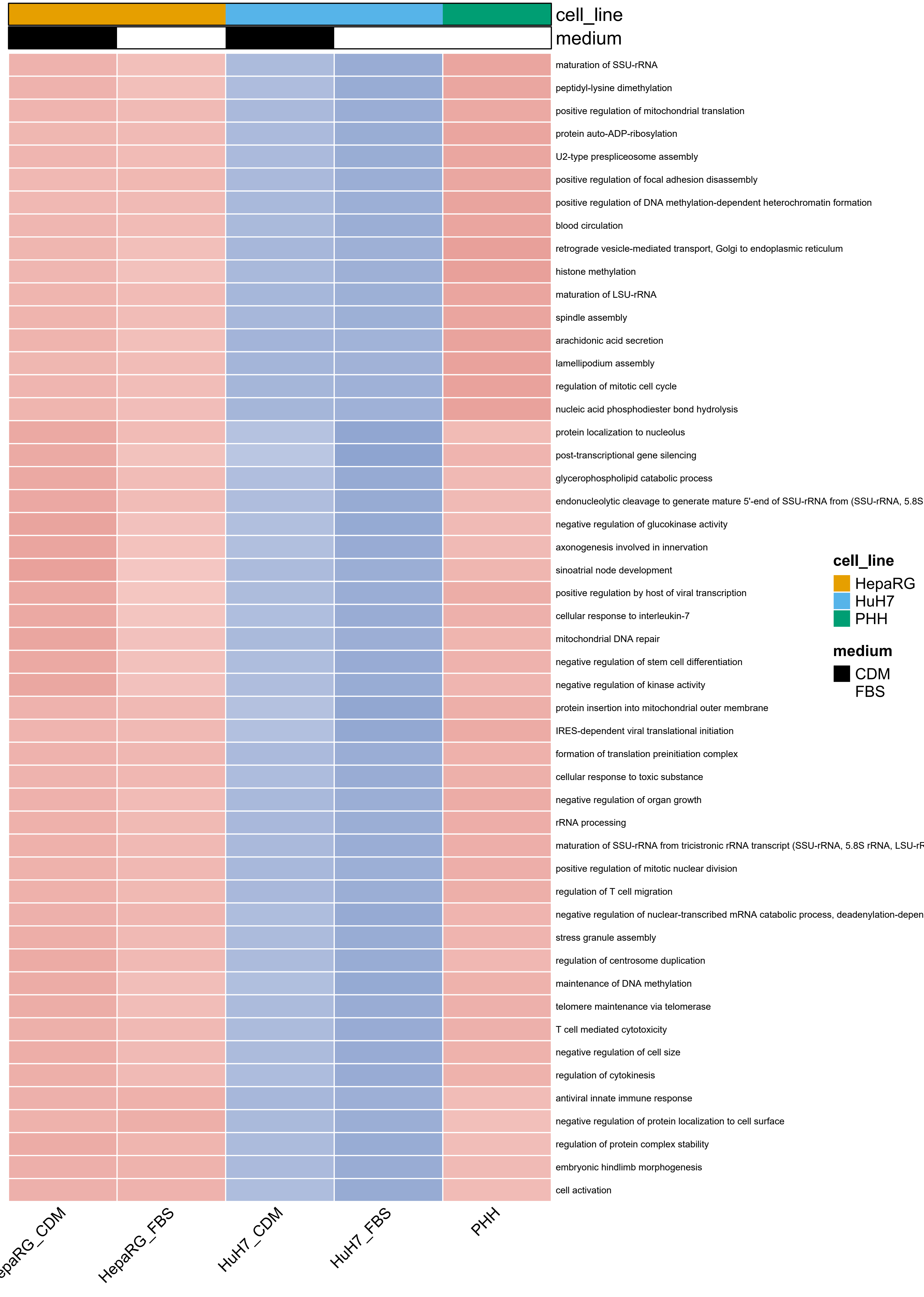
cell_line
medium
maturation of SSU-rRNA
peptidyl-lysine dimethylation
positive regulation of mitochondrial translation
protein auto-ADP-ribosylation
U2-type prespliceosome assembly
positive regulation of focal adhesion disassembly
positive regulation of DNA methylation-dependent heterochromatin formation
blood circulation
retrograde vesicle-mediated transport, Golgi to endoplasmic reticulum
histone methylation
maturation of LSU-rRNA
spindle assembly
arachidonic acid secretion
lamellipodium assembly
regulation of mitotic cell cycle
nucleic acid phosphodiester bond hydrolysis
protein localization to nucleolus
post-transcriptional gene silencing
glycerophospholipid catabolic process
endonucleolytic cleavage to generate mature 5'-end of SSU-rRNA from (SSU-rRNA, 5.8S
negative regulation of glucokinase activity
axonogenesis involved in innervation
sinoatrial node development
positive regulation by host of viral transcription
cellular response to interleukin-7
mitochondrial DNA repair
negative regulation of stem cell differentiation
negative regulation of kinase activity
protein insertion into mitochondrial outer membrane
IRES-dependent viral translational initiation
formation of translation preinitiation complex
cellular response to toxic substance
negative regulation of organ growth
rRNA processing
maturation of SSU-rRNA from tricistronic rRNA transcript (SSU-rRNA, 5.8S rRNA, LSU-rF
positive regulation of mitotic nuclear division
regulation of T cell migration
negative regulation of nuclear-transcribed mRNA catabolic process, deadenylation-depen
stress granule assembly
regulation of centrosome duplication
maintenance of DNA methylation
telomere maintenance via telomerase
T cell mediated cytotoxicity
negative regulation of cell size
regulation of cytokinesis
antiviral innate immune response
negative regulation of protein localization to cell surface
regulation of protein complex stability
embryonic hindlimb morphogenesis
cell activation
cell_line
HepaRG
HuH7
PHH
medium
CDM
FBS
epaRG_CDM
HepaRG_FBS
HuH7_CDM
HuH7_FBS
PHH

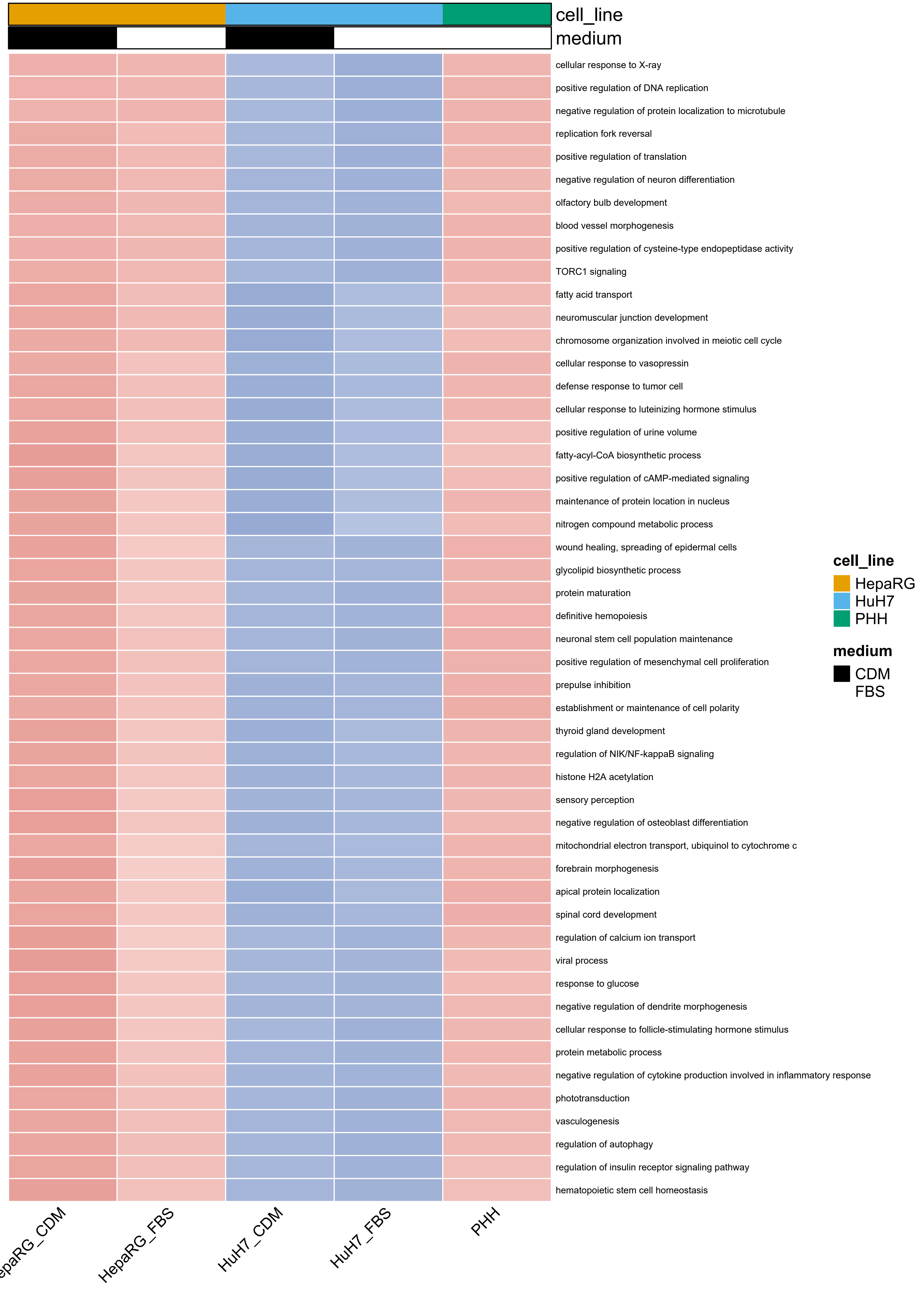
cell_line
medium
cellular response to X-ray
positive regulation of DNA replication
negative regulation of protein localization to microtubule
replication fork reversal
positive regulation of translation
negative regulation of neuron differentiation
olfactory bulb development
blood vessel morphogenesis
positive regulation of cysteine-type endopeptidase activity
TORC1 signaling
fatty acid transport
neuromuscular junction development
chromosome organization involved in meiotic cell cycle
cellular response to vasopressin
defense response to tumor cell
cellular response to luteinizing hormone stimulus
positive regulation of urine volume
fatty-acyl-CoA biosynthetic process
positive regulation of cAMP-mediated signaling
maintenance of protein location in nucleus
nitrogen compound metabolic process
wound healing, spreading of epidermal cells
glycolipid biosynthetic process
protein maturation
definitive hemopoiesis
neuronal stem cell population maintenance
positive regulation of mesenchymal cell proliferation
prepulse inhibition
establishment or maintenance of cell polarity
thyroid gland development
regulation of NIK/NF-kappaB signaling
histone H2A acetylation
sensory perception
negative regulation of osteoblast differentiation
mitochondrial electron transport, ubiquinol to cytochrome c
forebrain morphogenesis
apical protein localization
spinal cord development
regulation of calcium ion transport
viral process
response to glucose
negative regulation of dendrite morphogenesis
cellular response to follicle-stimulating hormone stimulus
protein metabolic process
negative regulation of cytokine production involved in inflammatory response
phototransduction
vasculogenesis
regulation of autophagy
regulation of insulin receptor signaling pathway
hematopoietic stem cell homeostasis
cell_line
HepaRG
HuH7
PHH
medium
CDM
FBS
HepaRG_FBS
HuH7_CDM
HuH7_FBS
PHH

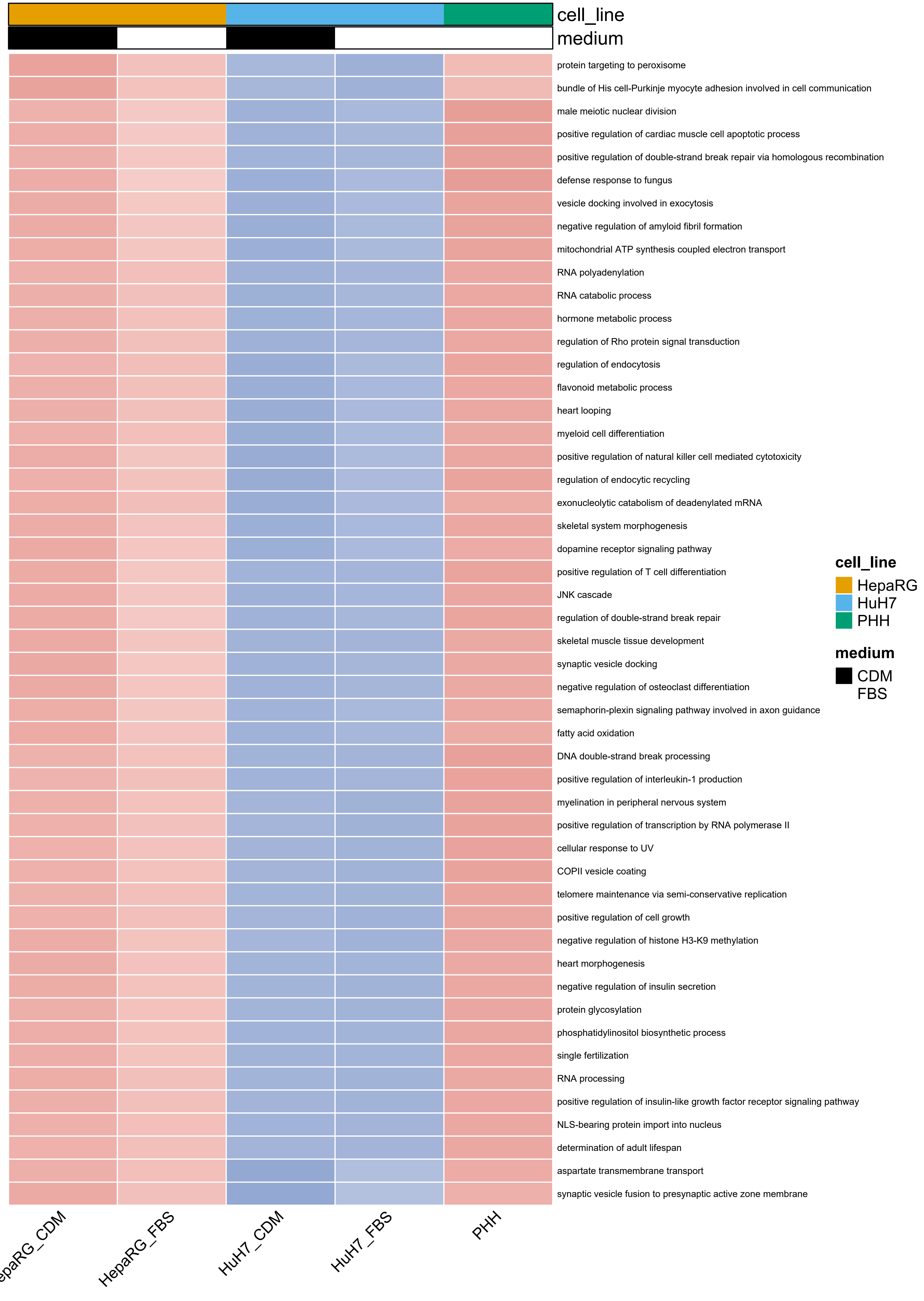
cell_line
medium
protein targeting to peroxisome
bundle of His cell-Purkinje myocyte adhesion involved in cell communication
male meiotic nuclear division
positive regulation of cardiac muscle cell apoptotic process
positive regulation of double-strand break repair via homologous recombination
defense response to fungus
vesicle docking involved in exocytosis
negative regulation of amyloid fibril formation
mitochondrial ATP synthesis coupled electron transport
RNA polyadenylation
RNA catabolic process
hormone metabolic process
regulation of Rho protein signal transduction
regulation of endocytosis
flavonoid metabolic process
heart looping
myeloid cell differentiation
positive regulation of natural killer cell mediated cytotoxicity
regulation of endocytic recycling
exonucleolytic catabolism of deadenylated mRNA
skeletal system morphogenesis
dopamine receptor signaling pathway
positive regulation of T cell differentiation
JNK cascade
regulation of double-strand break repair
skeletal muscle tissue development
synaptic vesicle docking
negative regulation of osteoclast differentiation
semaphorin-plexin signaling pathway involved in axon guidance
fatty acid oxidation
DNA double-strand break processing
positive regulation of interleukin-1 production
myelination in peripheral nervous system
positive regulation of transcription by RNA polymerase II
cellular response to UV
COPII vesicle coating
telomere maintenance via semi-conservative replication
positive regulation of cell growth
negative regulation of histone H3-K9 methylation
heart morphogenesis
negative regulation of insulin secretion
protein glycosylation
phosphatidylinositol biosynthetic process
single fertilization
RNA processing
positive regulation of insulin-like growth factor receptor signaling pathway
NLS-bearing protein import into nucleus
determination of adult lifespan
aspartate transmembrane transport
synaptic vesicle fusion to presynaptic active zone membrane
HepaRG_FBS
HuH7_CDM
HuH7_FBS
PHH
cell_line
HepaRG
HuH7
PHH
medium
CDM
FBS

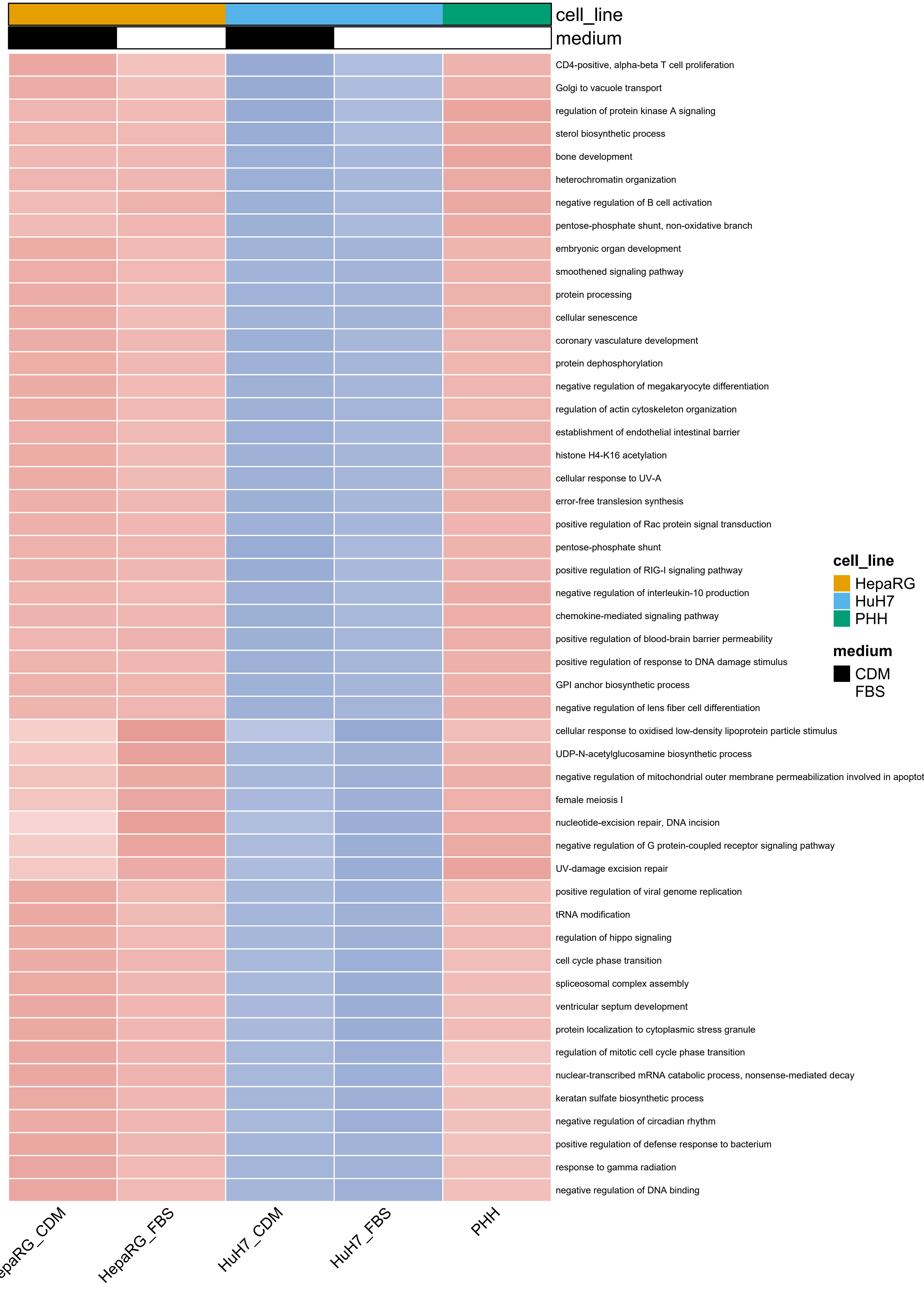

cell_line
medium
CD4-positive, alpha-beta T cell proliferation
Golgi to vacuole transport
regulation of protein kinase A signaling
sterol biosynthetic process
bone development
heterochromatin organization
negative regulation of B cell activation
pentose-phosphate shunt, non-oxidative branch
embryonic organ development
smoothened signaling pathway
protein processing
cellular senescence
coronary vasculature development
protein dephosphorylation
negative regulation of megakaryocyte differentiation
regulation of actin cytoskeleton organization
establishment of endothelial intestinal barrier
histone H4-K16 acetylation
cellular response to UV-A
error-free translesion synthesis
positive regulation of Rac protein signal transduction
pentose-phosphate shunt
positive regulation of RIG-I signaling pathway
negative regulation of interleukin-10 production
chemokine-mediated signaling pathway
positive regulation of blood-brain barrier permeability
positive regulation of response to DNA damage stimulus
GPI anchor biosynthetic process
negative regulation of lens fiber cell differentiation
cellular response to oxidised low-density lipoprotein particle stimulus
UDP-N-acetylglucosamine biosynthetic process
negative regulation of mitochondrial outer membrane permeabilization involved in apoptot
female meiosis I
nucleotide-excision repair, DNA incision
negative regulation of G protein-coupled receptor signaling pathway
UV-damage excision repair
positive regulation of viral genome replication
tRNA modification
regulation of hippo signaling
cell cycle phase transition
spliceosomal complex assembly
ventricular septum development
protein localization to cytoplasmic stress granule
regulation of mitotic cell cycle phase transition
nuclear-transcribed mRNA catabolic process, nonsense-mediated decay
keratan sulfate biosynthetic process
negative regulation of circadian rhythm
positive regulation of defense response to bacterium
response to gamma radiation
negative regulation of DNA binding
epaRG_CDM
HepaRG_FBS
HuH7_CDM
HuH7_FBS
PHH
cell_line
HepaRG
HuH7
PHH
medium
CDM
FBS

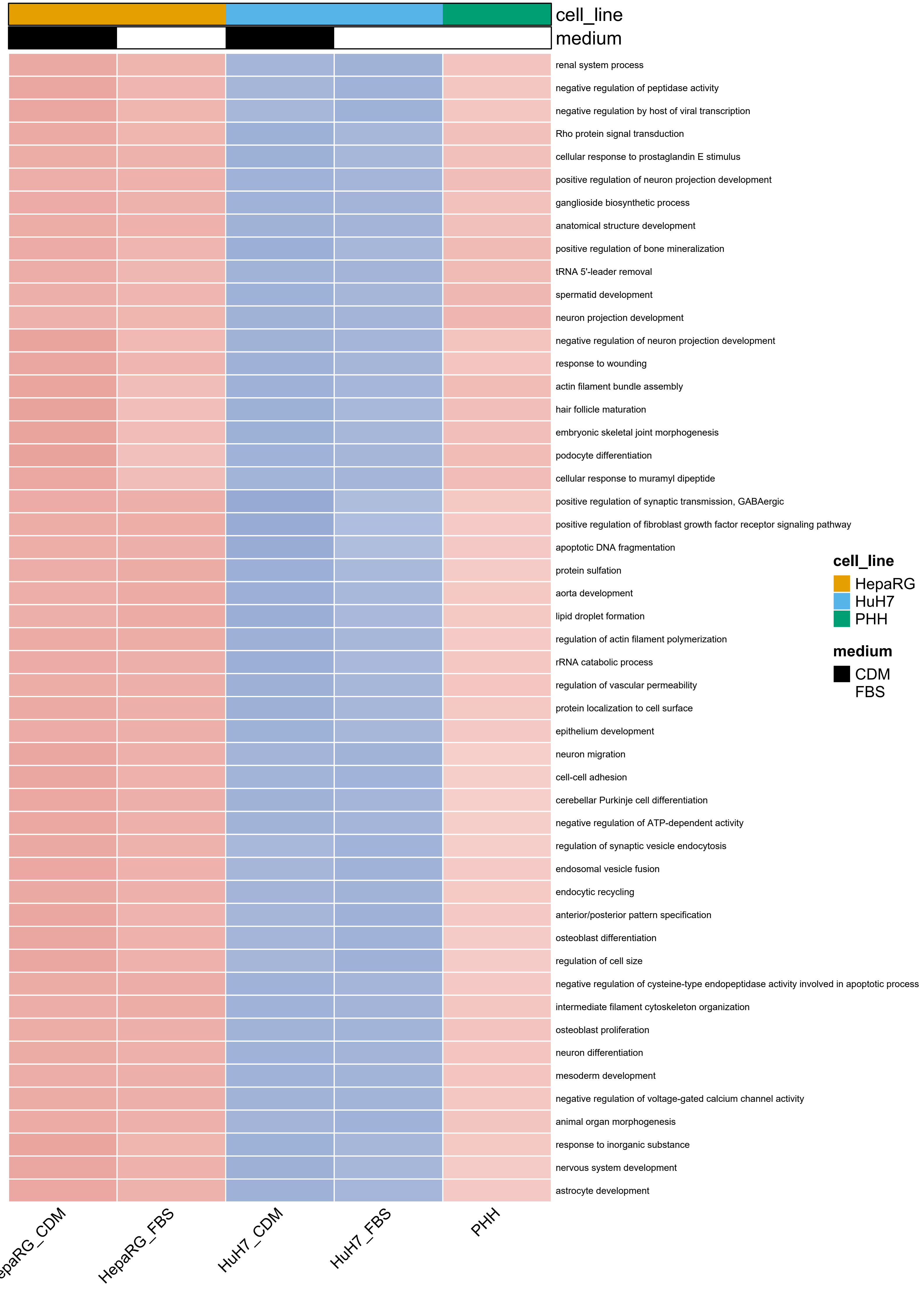
cell_line
medium
renal system process
negative regulation of peptidase activity
negative regulation by host of viral transcription
Rho protein signal transduction
cellular response to prostaglandin E stimulus
positive regulation of neuron projection development
ganglioside biosynthetic process
anatomical structure development
positive regulation of bone mineralization
tRNA 5'-leader removal
spermatid development
neuron projection development
negative regulation of neuron projection development
response to wounding
actin filament bundle assembly
hair follicle maturation
embryonic skeletal joint morphogenesis
podocyte differentiation
cellular response to muramyl dipeptide
positive regulation of synaptic transmission, GABAergic
positive regulation of fibroblast growth factor receptor signaling pathway
apoptotic DNA fragmentation
protein sulfation
aorta development
lipid droplet formation
regulation of actin filament polymerization
rRNA catabolic process
regulation of vascular permeability
protein localization to cell surface
epithelium development
neuron migration
cell-cell adhesion
cerebellar Purkinje cell differentiation
negative regulation of ATP-dependent activity
regulation of synaptic vesicle endocytosis
endosomal vesicle fusion
endocytic recycling
anterior/posterior pattern specification
osteoblast differentiation
regulation of cell size
negative regulation of cysteine-type endopeptidase activity involved in apoptotic process
intermediate filament cytoskeleton organization
osteoblast proliferation
neuron differentiation
mesoderm development
negative regulation of voltage-gated calcium channel activity
animal organ morphogenesis
response to inorganic substance
nervous system development
astrocyte development
epaRG_CDM
HepaRG_FBS
HuH7_CDM
HuH7_FBS
PHH
cell_line
HepaRG
HuH7
PHH
medium
CDM
FBS

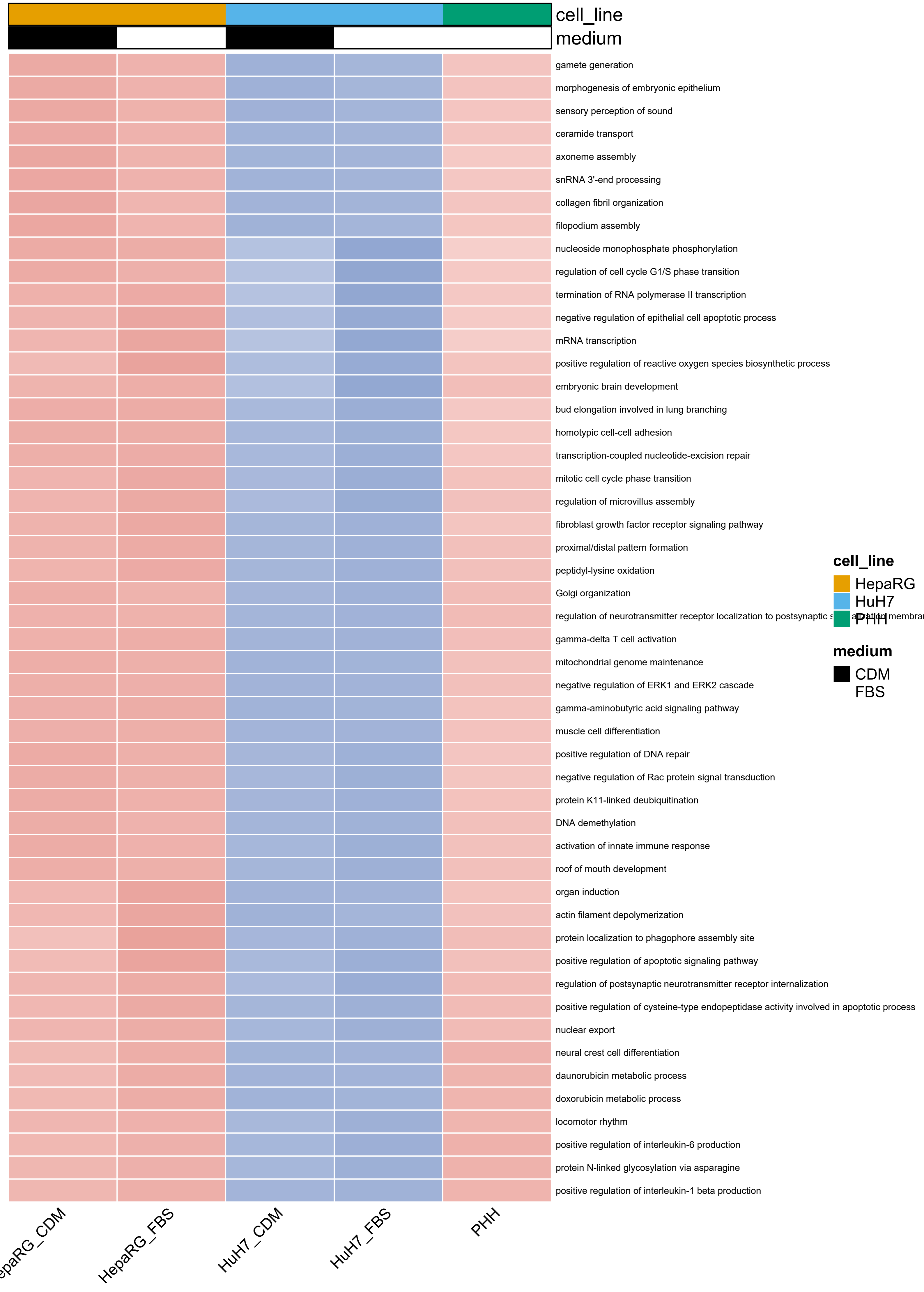
cell_line
medium
gamete generation
morphogenesis of embryonic epithelium
sensory perception of sound
ceramide transport
axoneme assembly
snRNA 3'-end processing
collagen fibril organization
filopodium assembly
nucleoside monophosphate phosphorylation
regulation of cell cycle G1/S phase transition
termination of RNA polymerase II transcription
negative regulation of epithelial cell apoptotic process
mRNA transcription
positive regulation of reactive oxygen species biosynthetic process
embryonic brain development
bud elongation involved in lung branching
homotypic cell-cell adhesion
transcription-coupled nucleotide-excision repair
mitotic cell cycle phase transition
regulation of microvillus assembly
fibroblast growth factor receptor signaling pathway
proximal/distal pattern formation
peptidyl-lysine oxidation
Golgi organization
regulation of neurotransmitter receptor localization to postsynaptic s
gamma-delta T cell activation
mitochondrial genome maintenance
negative regulation of ERK1 and ERK2 cascade
gamma-aminobutyric acid signaling pathway
muscle cell differentiation
positive regulation of DNA repair
negative regulation of Rac protein signal transduction
protein K11-linked deubiquitination
DNA demethylation
activation of innate immune response
roof of mouth development
organ induction
actin filament depolymerization
protein localization to phagophore assembly site
positive regulation of apoptotic signaling pathway
regulation of postsynaptic neurotransmitter receptor internalization
positive regulation of cysteine-type endopeptidase activity involved in apoptotic process
nuclear export
neural crest cell differentiation
daunorubicin metabolic process
doxorubicin metabolic process
locomotor rhythm
positive regulation of interleukin-6 production
protein N-linked glycosylation via asparagine
positive regulation of interleukin-1 beta production
epaRG_CDM
HepaRG_FBS
HuH7_CDM
HuH7_FBS
PHH
cell_line
HepaRG
HuH7
PHH
medium
CDM
FBS

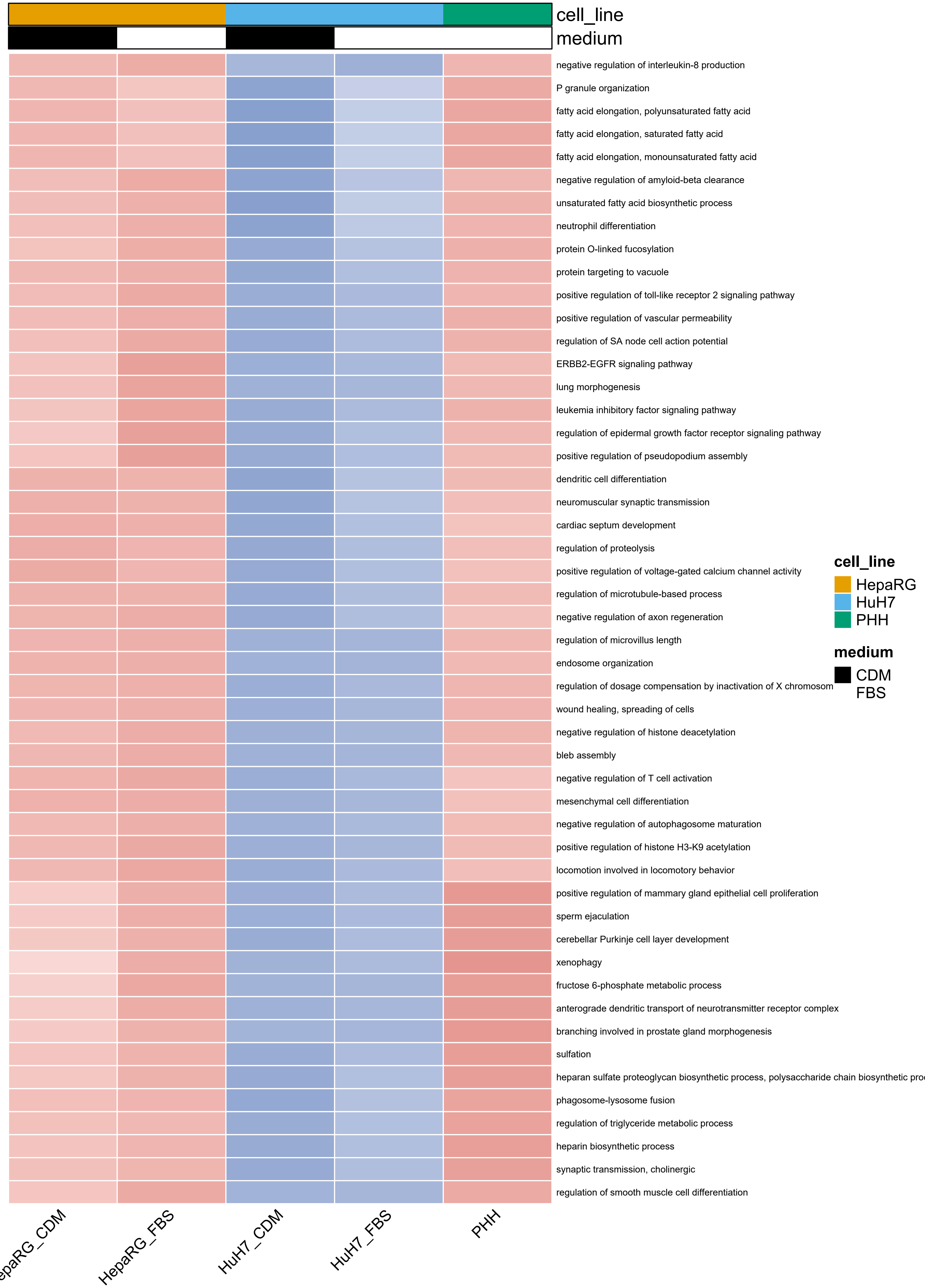
cell_line
medium
negative regulation of interleukin-8 production
P granule organization
fatty acid elongation, polyunsaturated fatty acid
fatty acid elongation, saturated fatty acid
fatty acid elongation, monounsaturated fatty acid
negative regulation of amyloid-beta clearance
unsaturated fatty acid biosynthetic process
neutrophil differentiation
protein O-linked fucosylation
protein targeting to vacuole
positive regulation of toll-like receptor 2 signaling pathway
positive regulation of vascular permeability
regulation of SA node cell action potential
ERBB2-EGFR signaling pathway
lung morphogenesis
leukemia inhibitory factor signaling pathway
regulation of epidermal growth factor receptor signaling pathway
positive regulation of pseudopodium assembly
dendritic cell differentiation
neuromuscular synaptic transmission
cardiac septum development
regulation of proteolysis
positive regulation of voltage-gated calcium channel activity
regulation of microtubule-based process
negative regulation of axon regeneration
regulation of microvillus length
endosome organization
regulation of dosage compensation by inactivation of X chromosom
wound healing, spreading of cells
negative regulation of histone deacetylation
bleb assembly
negative regulation of T cell activation
mesenchymal cell differentiation
negative regulation of autophagosome maturation
positive regulation of histone H3-K9 acetylation
locomotion involved in locomotory behavior
positive regulation of mammary gland epithelial cell proliferation
sperm ejaculation
cerebellar Purkinje cell layer development
xenophagy
fructose 6-phosphate metabolic process
anterograde dendritic transport of neurotransmitter receptor complex
branching involved in prostate gland morphogenesis
sulfation
heparan sulfate proteoglycan biosynthetic process, polysaccharide chain biosynthetic pro
phagosome-lysosome fusion
regulation of triglyceride metabolic process
heparin biosynthetic process
synaptic transmission, cholinergic
regulation of smooth muscle cell differentiation
cell_line
HepaRG
HuH7
PHH
medium
CDM
FBS
epaRG_CDM
HepaRG_FBS
HuH7_CDM
HuH7_FBS
PHH

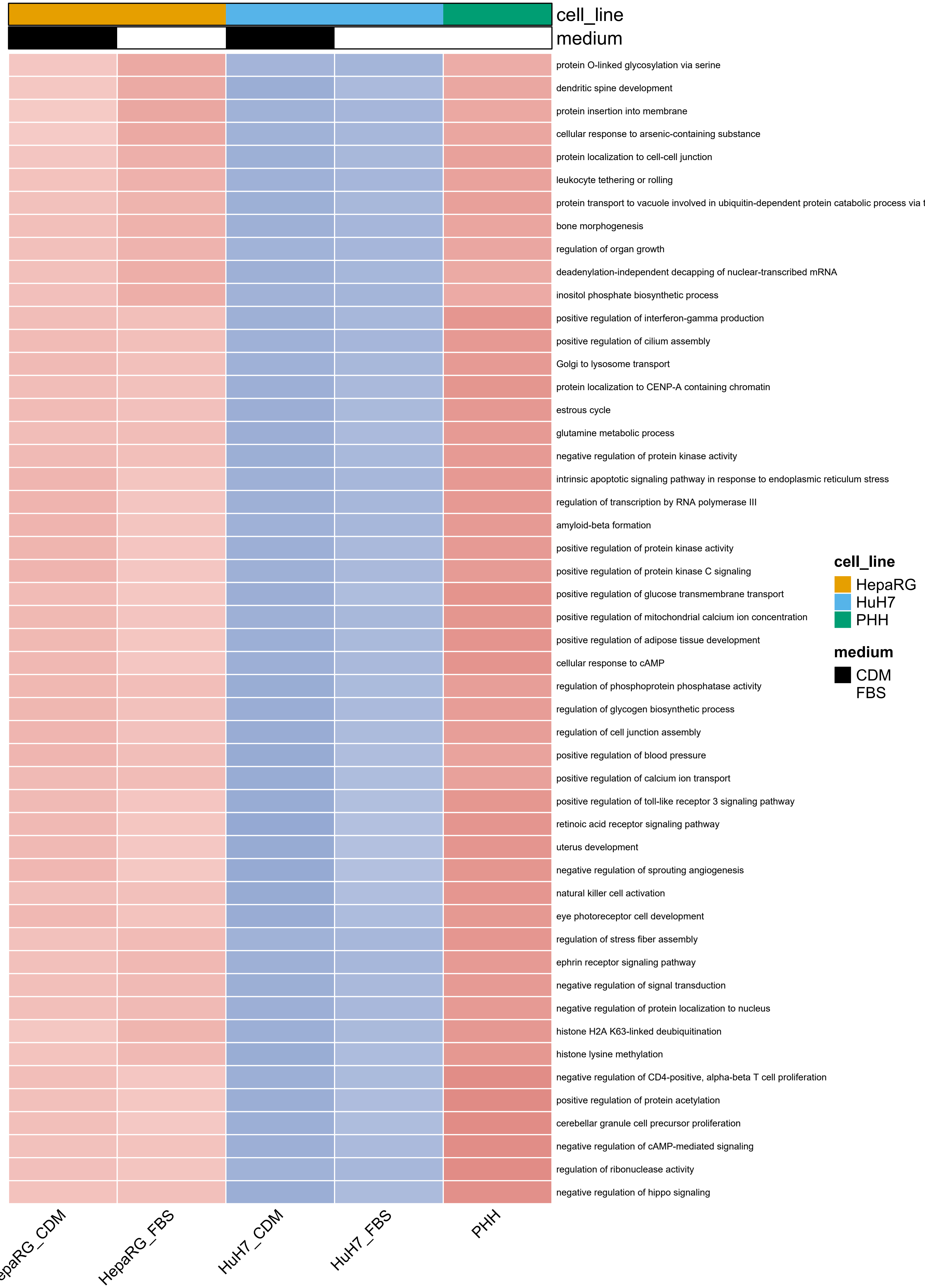
cell_line
medium
protein O-linked glycosylation via serine
dendritic spine development
protein insertion into membrane
cellular response to arsenic-containing substance
protein localization to cell-cell junction
leukocyte tethering or rolling
protein transport to vacuole involved in ubiquitin-dependent protein catabolic process via t
bone morphogenesis
regulation of organ growth
deadenylation-independent decapping of nuclear-transcribed mRNA
inositol phosphate biosynthetic process
positive regulation of interferon-gamma production
positive regulation of cilium assembly
Golgi to lysosome transport
protein localization to CENP-A containing chromatin
estrous cycle
glutamine metabolic process
negative regulation of protein kinase activity
intrinsic apoptotic signaling pathway in response to endoplasmic reticulum stress
regulation of transcription by RNA polymerase III
amyloid-beta formation
positive regulation of protein kinase activity
positive regulation of protein kinase C signaling
positive regulation of glucose transmembrane transport
positive regulation of mitochondrial calcium ion concentration
positive regulation of adipose tissue development
cellular response to cAMP
regulation of phosphoprotein phosphatase activity
regulation of glycogen biosynthetic process
regulation of cell junction assembly
positive regulation of blood pressure
positive regulation of calcium ion transport
positive regulation of toll-like receptor 3 signaling pathway
retinoic acid receptor signaling pathway
uterus development
negative regulation of sprouting angiogenesis
natural killer cell activation
eye photoreceptor cell development
regulation of stress fiber assembly
ephrin receptor signaling pathway
negative regulation of signal transduction
negative regulation of protein localization to nucleus
histone H2A K63-linked deubiquitination
histone lysine methylation
negative regulation of CD4-positive, alpha-beta T cell proliferation
positive regulation of protein acetylation
cerebellar granule cell precursor proliferation
negative regulation of cAMP-mediated signaling
regulation of ribonuclease activity
negative regulation of hippo signaling
cell_line
HepaRG
HuH7
PHH
medium
CDM
FBS
epaRG_CDM
HepaRG_FBS
HuH7_CDM
HuH7_FBS
PHH

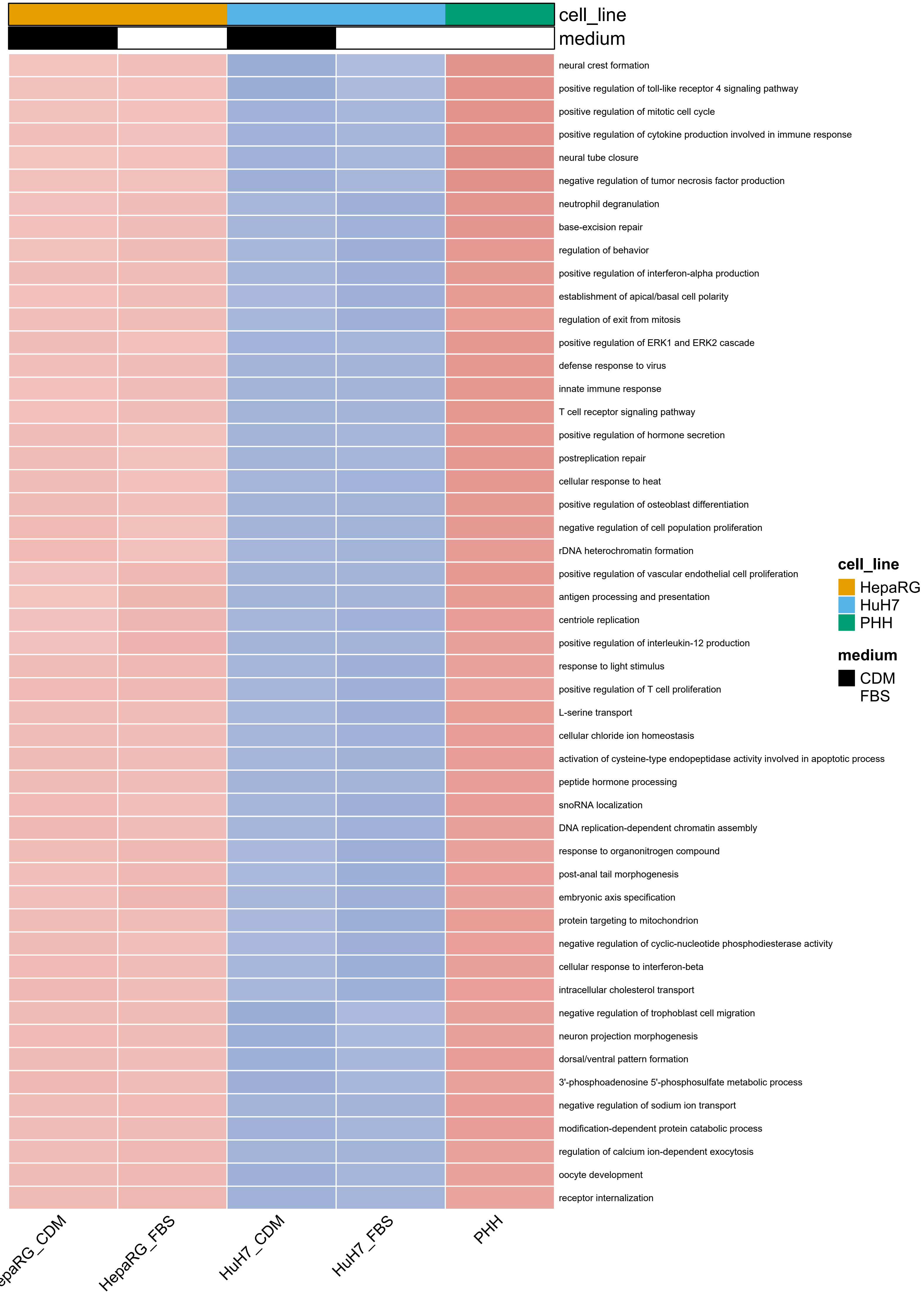
cell_line
medium
neural crest formation
positive regulation of toll-like receptor 4 signaling pathway
positive regulation of mitotic cell cycle
positive regulation of cytokine production involved in immune response
neural tube closure
negative regulation of tumor necrosis factor production
neutrophil degranulation
base-excision repair
regulation of behavior
positive regulation of interferon-alpha production
establishment of apical/basal cell polarity
regulation of exit from mitosis
positive regulation of ERK1 and ERK2 cascade
defense response to virus
innate immune response
T cell receptor signaling pathway
positive regulation of hormone secretion
postreplication repair
cellular response to heat
positive regulation of osteoblast differentiation
negative regulation of cell population proliferation
rDNA heterochromatin formation
positive regulation of vascular endothelial cell proliferation
antigen processing and presentation
centriole replication
positive regulation of interleukin-12 production
response to light stimulus
positive regulation of T cell proliferation
L-serine transport
cellular chloride ion homeostasis
activation of cysteine-type endopeptidase activity involved in apoptotic process
peptide hormone processing
snoRNA localization
DNA replication-dependent chromatin assembly
response to organonitrogen compound
post-anal tail morphogenesis
embryonic axis specification
protein targeting to mitochondrion
negative regulation of cyclic-nucleotide phosphodiesterase activity
cellular response to interferon-beta
intracellular cholesterol transport
negative regulation of trophoblast cell migration
neuron projection morphogenesis
dorsal/ventral pattern formation
3'-phosphoadenosine 5'-phosphosulfate metabolic process
negative regulation of sodium ion transport
modification-dependent protein catabolic process
regulation of calcium ion-dependent exocytosis
oocyte development
receptor internalization
cell_line
HepaRG
HuH7
PHH
medium
CDM
FBS
HepaRG_FBS
HuH7_CDM
HuH7_FBS
PHH

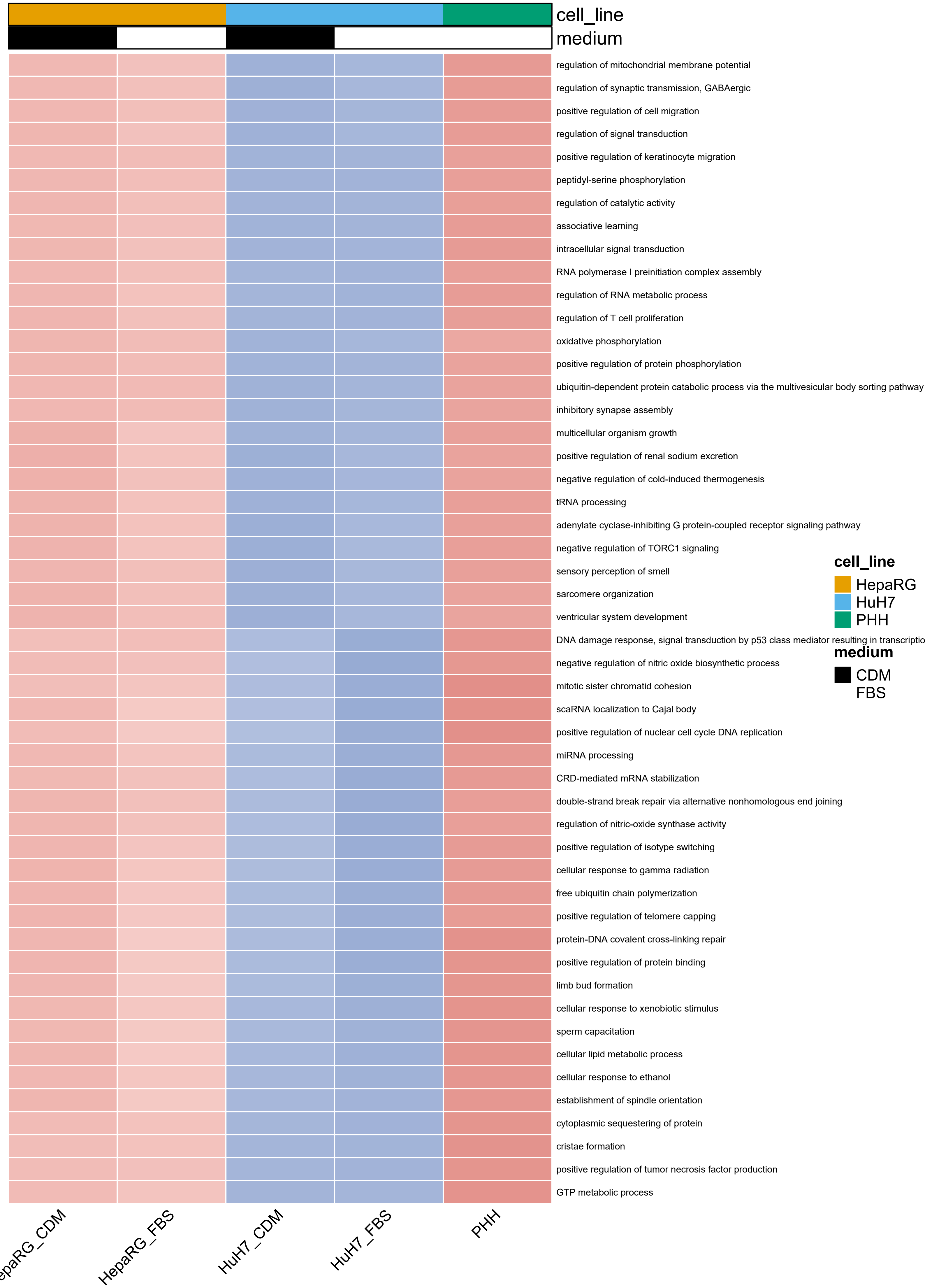

cell_line
medium
regulation of mitochondrial membrane potential
regulation of synaptic transmission, GABAergic
positive regulation of cell migration
regulation of signal transduction
positive regulation of keratinocyte migration
peptidyl-serine phosphorylation
regulation of catalytic activity
associative learning
intracellular signal transduction
RNA polymerase I preinitiation complex assembly
regulation of RNA metabolic process
regulation of T cell proliferation
oxidative phosphorylation
positive regulation of protein phosphorylation
ubiquitin-dependent protein catabolic process via the multivesicular body sorting pathway
inhibitory synapse assembly
multicellular organism growth
positive regulation of renal sodium excretion
negative regulation of cold-induced thermogenesis
tRNA processing
adenylate cyclase-inhibiting G protein-coupled receptor signaling pathway
negative regulation of TORC1 signaling
sensory perception of smell
sarcomere organization
ventricular system development
DNA damage response, signal transduction by p53 class mediator resulting in transcriptio
negative regulation of nitric oxide biosynthetic process
mitotic sister chromatid cohesion
scaRNA localization to Cajal body
positive regulation of nuclear cell cycle DNA replication
miRNA processing
CRD-mediated mRNA stabilization
double-strand break repair via alternative nonhomologous end joining
regulation of nitric-oxide synthase activity
positive regulation of isotype switching
cellular response to gamma radiation
free ubiquitin chain polymerization
positive regulation of telomere capping
protein-DNA covalent cross-linking repair
positive regulation of protein binding
limb bud formation
cellular response to xenobiotic stimulus
sperm capacitation
cellular lipid metabolic process
cellular response to ethanol
establishment of spindle orientation
cytoplasmic sequestering of protein
cristae formation
positive regulation of tumor necrosis factor production
GTP metabolic process
cell_line
HepaRG
HuH7
PHH
medium
CDM
FBS
epaRG_CDM
HepaRG_FBS
HuH7_CDM
HuH7_FBS
PHH

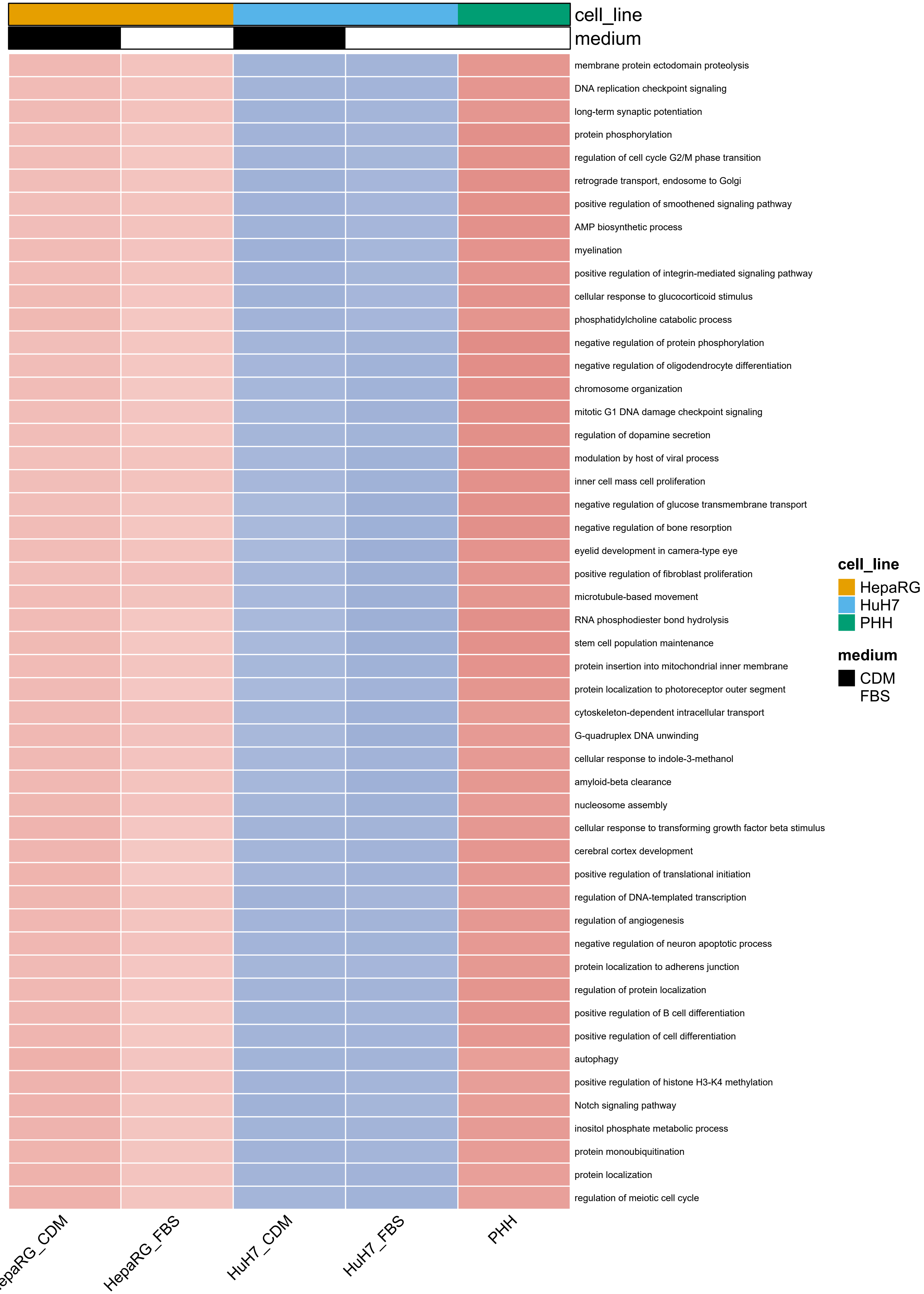

cell_line
medium
membrane protein ectodomain proteolysis
DNA replication checkpoint signaling
long-term synaptic potentiation
protein phosphorylation
regulation of cell cycle G2/M phase transition
retrograde transport, endosome to Golgi
positive regulation of smoothened signaling pathway
AMP biosynthetic process
myelination
positive regulation of integrin-mediated signaling pathway
cellular response to glucocorticoid stimulus
phosphatidylcholine catabolic process
negative regulation of protein phosphorylation
negative regulation of oligodendrocyte differentiation
chromosome organization
mitotic G1 DNA damage checkpoint signaling
regulation of dopamine secretion
modulation by host of viral process
inner cell mass cell proliferation
negative regulation of glucose transmembrane transport
negative regulation of bone resorption
eyelid development in camera-type eye
positive regulation of fibroblast proliferation
microtubule-based movement
RNA phosphodiester bond hydrolysis
stem cell population maintenance
protein insertion into mitochondrial inner membrane
protein localization to photoreceptor outer segment
cytoskeleton-dependent intracellular transport
G-quadruplex DNA unwinding
cellular response to indole-3-methanol
amyloid-beta clearance
nucleosome assembly
cellular response to transforming growth factor beta stimulus
cerebral cortex development
positive regulation of translational initiation
regulation of DNA-templated transcription
regulation of angiogenesis
negative regulation of neuron apoptotic process
protein localization to adherens junction
regulation of protein localization
positive regulation of B cell differentiation
positive regulation of cell differentiation
autophagy
positive regulation of histone H3-K4 methylation
Notch signaling pathway
inositol phosphate metabolic process
protein monoubiquitination
protein localization
regulation of meiotic cell cycle
HepaRG_FBS
HuH7_CDM
HuH7_FBS
PHH
cell_line
HepaRG
HuH7
PHH
medium
CDM
FBS

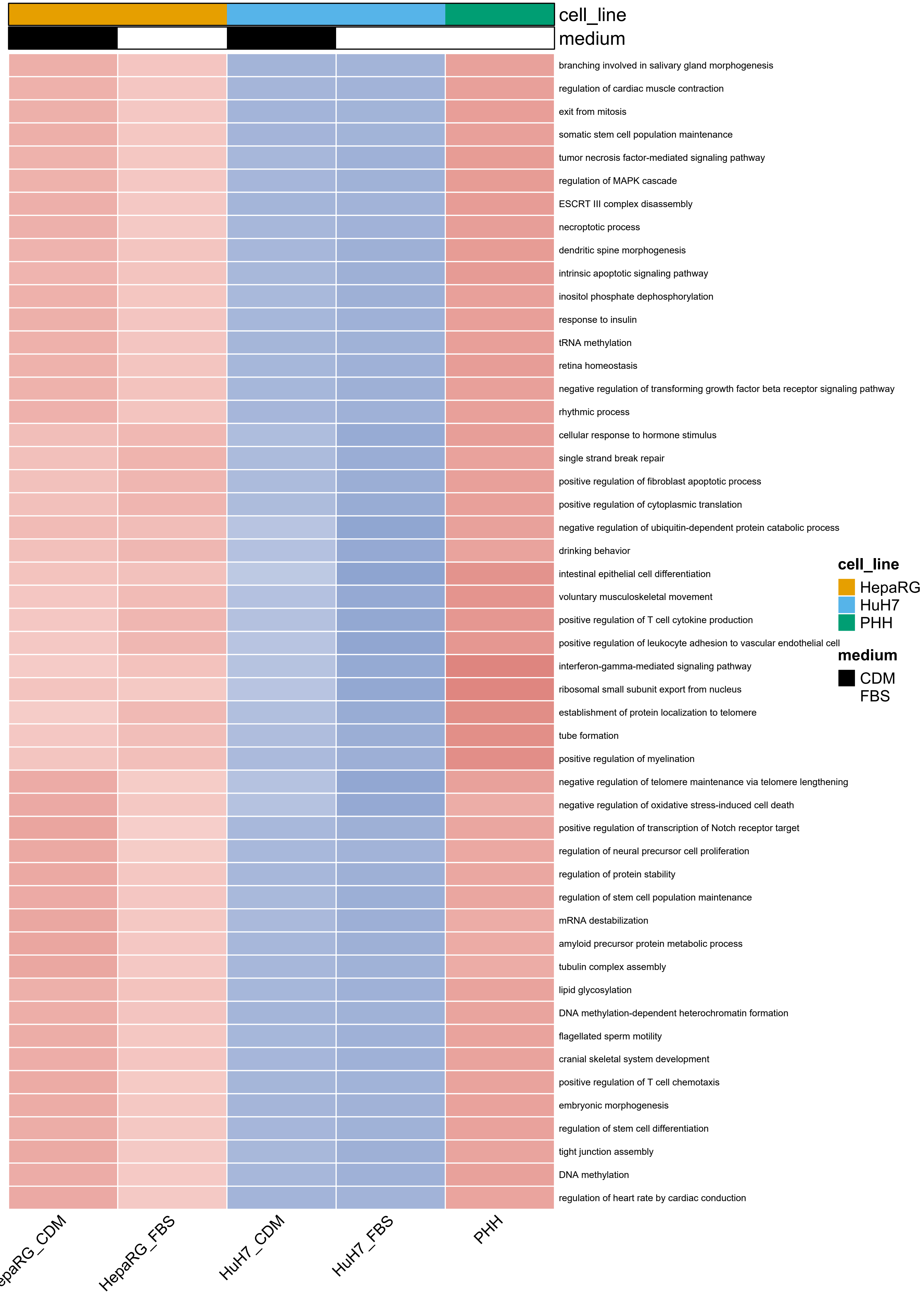
cell_line
medium
branching involved in salivary gland morphogenesis
regulation of cardiac muscle contraction
exit from mitosis
somatic stem cell population maintenance
tumor necrosis factor-mediated signaling pathway
regulation of MAPK cascade
ESCRT III complex disassembly
necroptotic process
dendritic spine morphogenesis
intrinsic apoptotic signaling pathway
inositol phosphate dephosphorylation
response to insulin
tRNA methylation
retina homeostasis
negative regulation of transforming growth factor beta receptor signaling pathway
rhythmic process
cellular response to hormone stimulus
single strand break repair
positive regulation of fibroblast apoptotic process
positive regulation of cytoplasmic translation
negative regulation of ubiquitin-dependent protein catabolic process
drinking behavior
intestinal epithelial cell differentiation
voluntary musculoskeletal movement
positive regulation of T cell cytokine production
positive regulation of leukocyte adhesion to vascular endothelial cell
interferon-gamma-mediated signaling pathway
ribosomal small subunit export from nucleus
establishment of protein localization to telomere
tube formation
positive regulation of myelination
negative regulation of telomere maintenance via telomere lengthening
negative regulation of oxidative stress-induced cell death
positive regulation of transcription of Notch receptor target
regulation of neural precursor cell proliferation
regulation of protein stability
regulation of stem cell population maintenance
mRNA destabilization
amyloid precursor protein metabolic process
tubulin complex assembly
lipid glycosylation
DNA methylation-dependent heterochromatin formation
flagellated sperm motility
cranial skeletal system development
positive regulation of T cell chemotaxis
embryonic morphogenesis
regulation of stem cell differentiation
tight junction assembly
DNA methylation
regulation of heart rate by cardiac conduction
cell_line
HepaRG
HuH7
PHH
medium
CDM
FBS
HepaRG_FBS
HuH7_CDM
HuH7_FBS
PHH

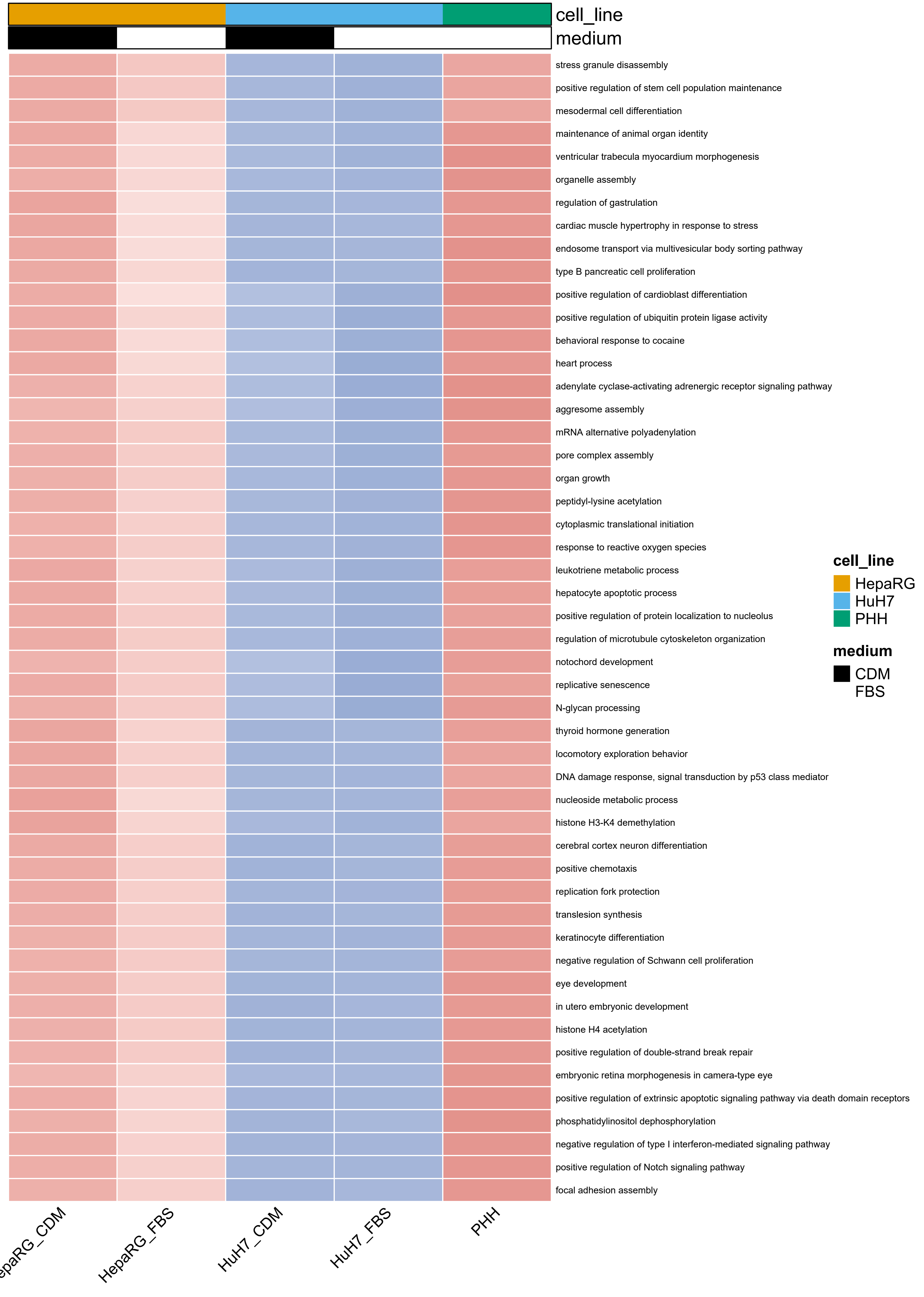
cell_line
medium
stress granule disassembly
positive regulation of stem cell population maintenance
mesodermal cell differentiation
maintenance of animal organ identity
ventricular trabecula myocardium morphogenesis
organelle assembly
regulation of gastrulation
cardiac muscle hypertrophy in response to stress
endosome transport via multivesicular body sorting pathway
type B pancreatic cell proliferation
positive regulation of cardioblast differentiation
positive regulation of ubiquitin protein ligase activity
behavioral response to cocaine
heart process
adenylate cyclase-activating adrenergic receptor signaling pathway
aggresome assembly
mRNA alternative polyadenylation
pore complex assembly
organ growth
peptidyl-lysine acetylation
cytoplasmic translational initiation
response to reactive oxygen species
leukotriene metabolic process
hepatocyte apoptotic process
positive regulation of protein localization to nucleolus
regulation of microtubule cytoskeleton organization
notochord development
replicative senescence
N-glycan processing
thyroid hormone generation
locomotory exploration behavior
DNA damage response, signal transduction by p53 class mediator
nucleoside metabolic process
histone H3-K4 demethylation
cerebral cortex neuron differentiation
positive chemotaxis
replication fork protection
translesion synthesis
keratinocyte differentiation
negative regulation of Schwann cell proliferation
eye development
in utero embryonic development
histone H4 acetylation
positive regulation of double-strand break repair
embryonic retina morphogenesis in camera-type eye
positive regulation of extrinsic apoptotic signaling pathway via death domain receptors
phosphatidylinositol dephosphorylation
negative regulation of type I interferon-mediated signaling pathway
positive regulation of Notch signaling pathway
focal adhesion assembly
cell_line
HepaRG
HuH7
PHH
medium
CDM
FBS
HepaRG_FBS
HuH7_CDM
HuH7_FBS
PHH

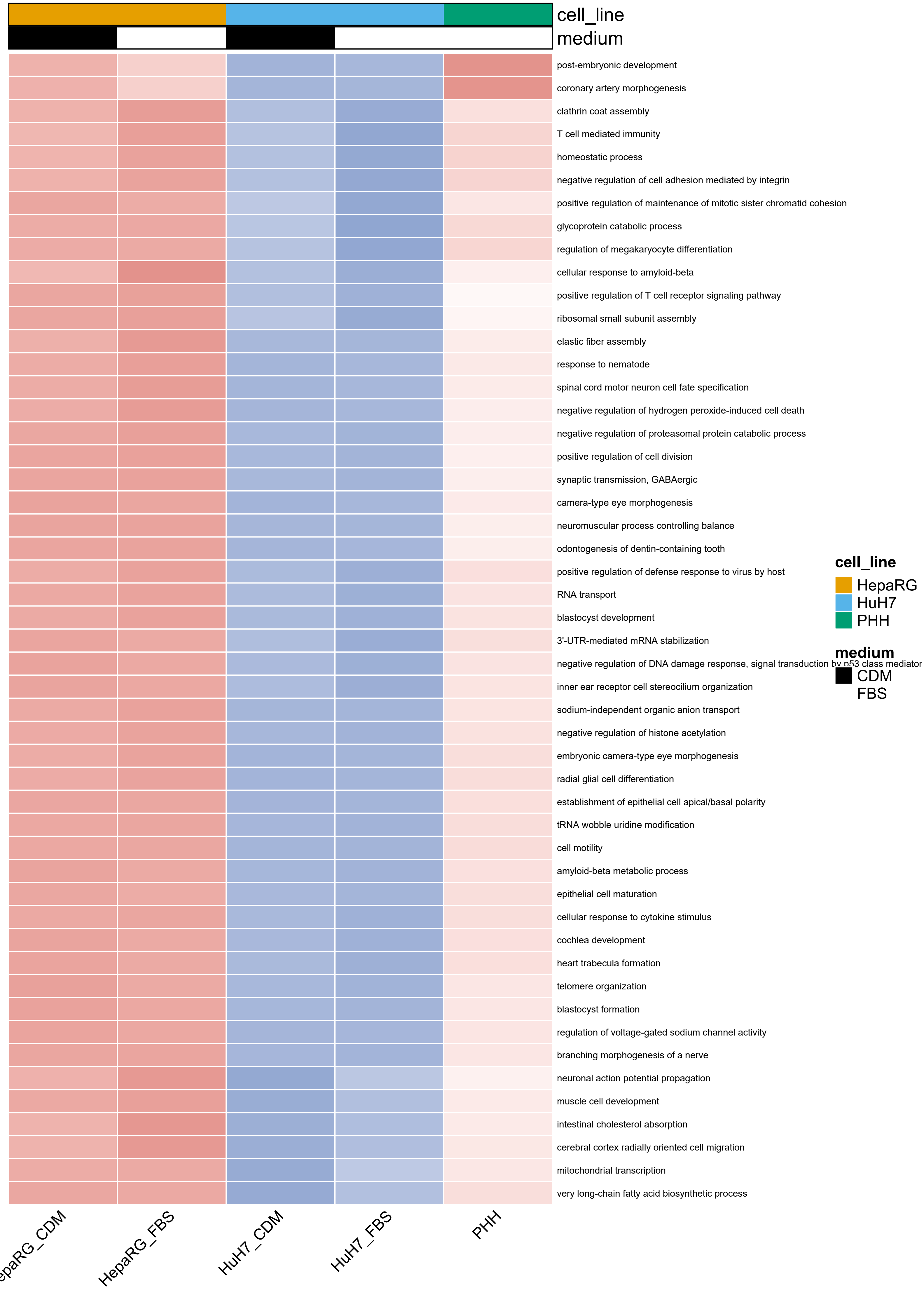
cell_line
medium
post-embryonic development
coronary artery morphogenesis
clathrin coat assembly
T cell mediated immunity
homeostatic process
negative regulation of cell adhesion mediated by integrin
positive regulation of maintenance of mitotic sister chromatid cohesion
glycoprotein catabolic process
regulation of megakaryocyte differentiation
cellular response to amyloid-beta
positive regulation of T cell receptor signaling pathway
ribosomal small subunit assembly
elastic fiber assembly
response to nematode
spinal cord motor neuron cell fate specification
negative regulation of hydrogen peroxide-induced cell death
negative regulation of proteasomal protein catabolic process
positive regulation of cell division
synaptic transmission, GABAergic
camera-type eye morphogenesis
neuromuscular process controlling balance
odontogenesis of dentin-containing tooth
positive regulation of defense response to virus by host
RNA transport
blastocyst development
3'-UTR-mediated mRNA stabilization
negative regulation of DNA damage response, signal transduction by p53 class mediator
inner ear receptor cell stereocilium organization
sodium-independent organic anion transport
negative regulation of histone acetylation
embryonic camera-type eye morphogenesis
radial glial cell differentiation
establishment of epithelial cell apical/basal polarity
tRNA wobble uridine modification
cell motility
amyloid-beta metabolic process
epithelial cell maturation
cellular response to cytokine stimulus
cochlea development
heart trabecula formation
telomere organization
blastocyst formation
regulation of voltage-gated sodium channel activity
branching morphogenesis of a nerve
neuronal action potential propagation
muscle cell development
intestinal cholesterol absorption
cerebral cortex radially oriented cell migration
mitochondrial transcription
very long-chain fatty acid biosynthetic process
HepaRG_CDM
HepaRG_FBS
HuH7_CDM
HuH7_FBS
PHH
cell_line
HepaRG
HuH7
PHH
medium
CDM
FBS

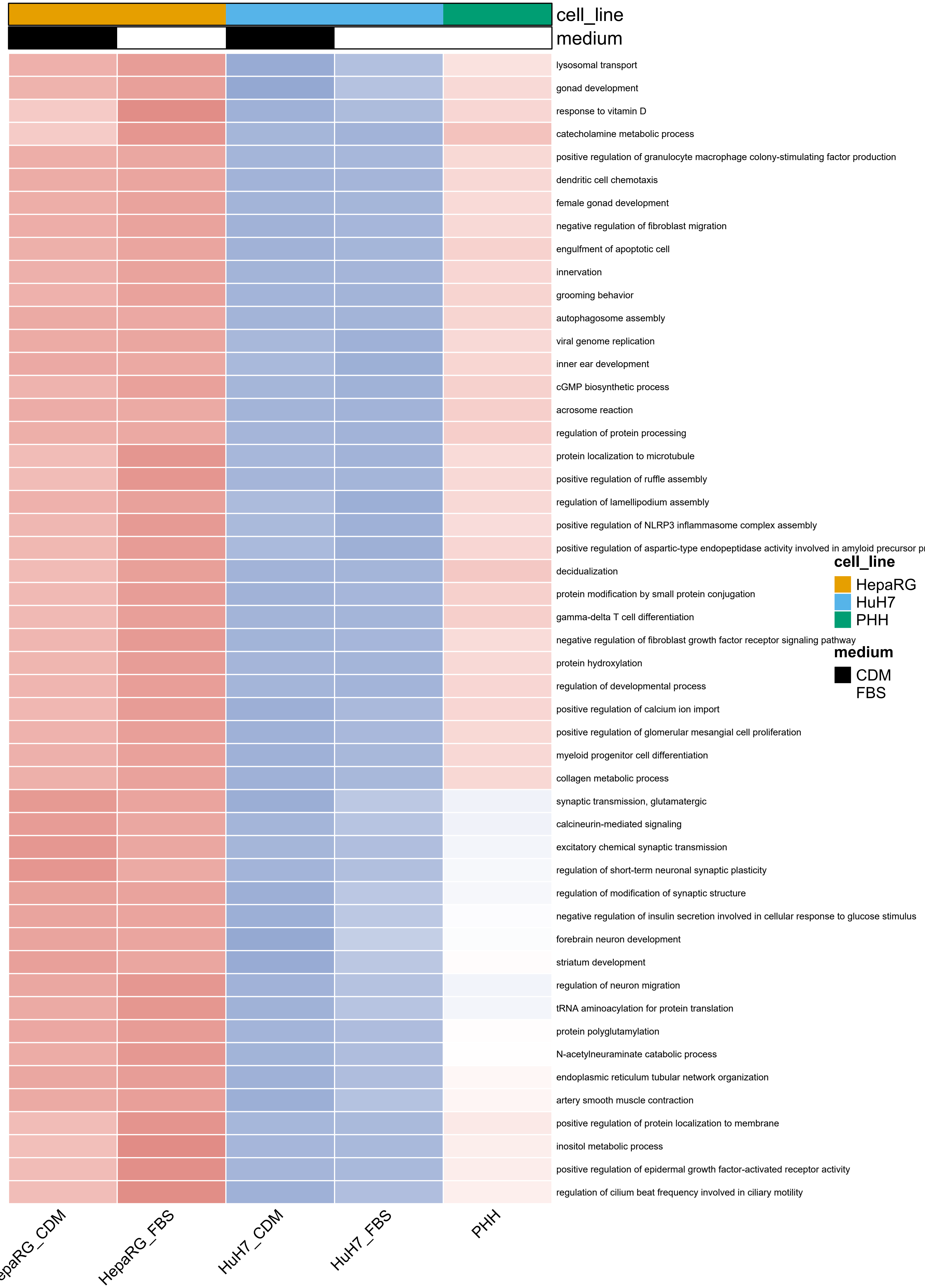
cell_line
medium
lysosomal transport
gonad development
response to vitamin D
catecholamine metabolic process
positive regulation of granulocyte macrophage colony-stimulating factor production
dendritic cell chemotaxis
female gonad development
negative regulation of fibroblast migration
engulfment of apoptotic cell
innervation
grooming behavior
autophagosome assembly
viral genome replication
inner ear development
cGMP biosynthetic process
acrosome reaction
regulation of protein processing
protein localization to microtubule
positive regulation of ruffle assembly
regulation of lamellipodium assembly
positive regulation of NLRP3 inflammasome complex assembly
positive regulation of aspartic-type endopeptidase activity involved in amyloid precursor p
decidualization
protein modification by small protein conjugation
gamma-delta T cell differentiation
negative regulation of fibroblast growth factor receptor signaling pathway
protein hydroxylation
regulation of developmental process
positive regulation of calcium ion import
positive regulation of glomerular mesangial cell proliferation
myeloid progenitor cell differentiation
collagen metabolic process
synaptic transmission, glutamatergic
calcineurin-mediated signaling
excitatory chemical synaptic transmission
regulation of short-term neuronal synaptic plasticity
regulation of modification of synaptic structure
negative regulation of insulin secretion involved in cellular response to glucose stimulus
forebrain neuron development
striatum development
regulation of neuron migration
tRNA aminoacylation for protein translation
protein polyglutamylation
N-acetylneuraminate catabolic process
endoplasmic reticulum tubular network organization
artery smooth muscle contraction
positive regulation of protein localization to membrane
inositol metabolic process
positive regulation of epidermal growth factor-activated receptor activity
regulation of cilium beat frequency involved in ciliary motility
cell_line
HepaRG
HuH7
PHH
medium
CDM
FBS
epaRG_CDM
HepaRG_FBS
HuH7_CDM
HuH7_FBS
PHH

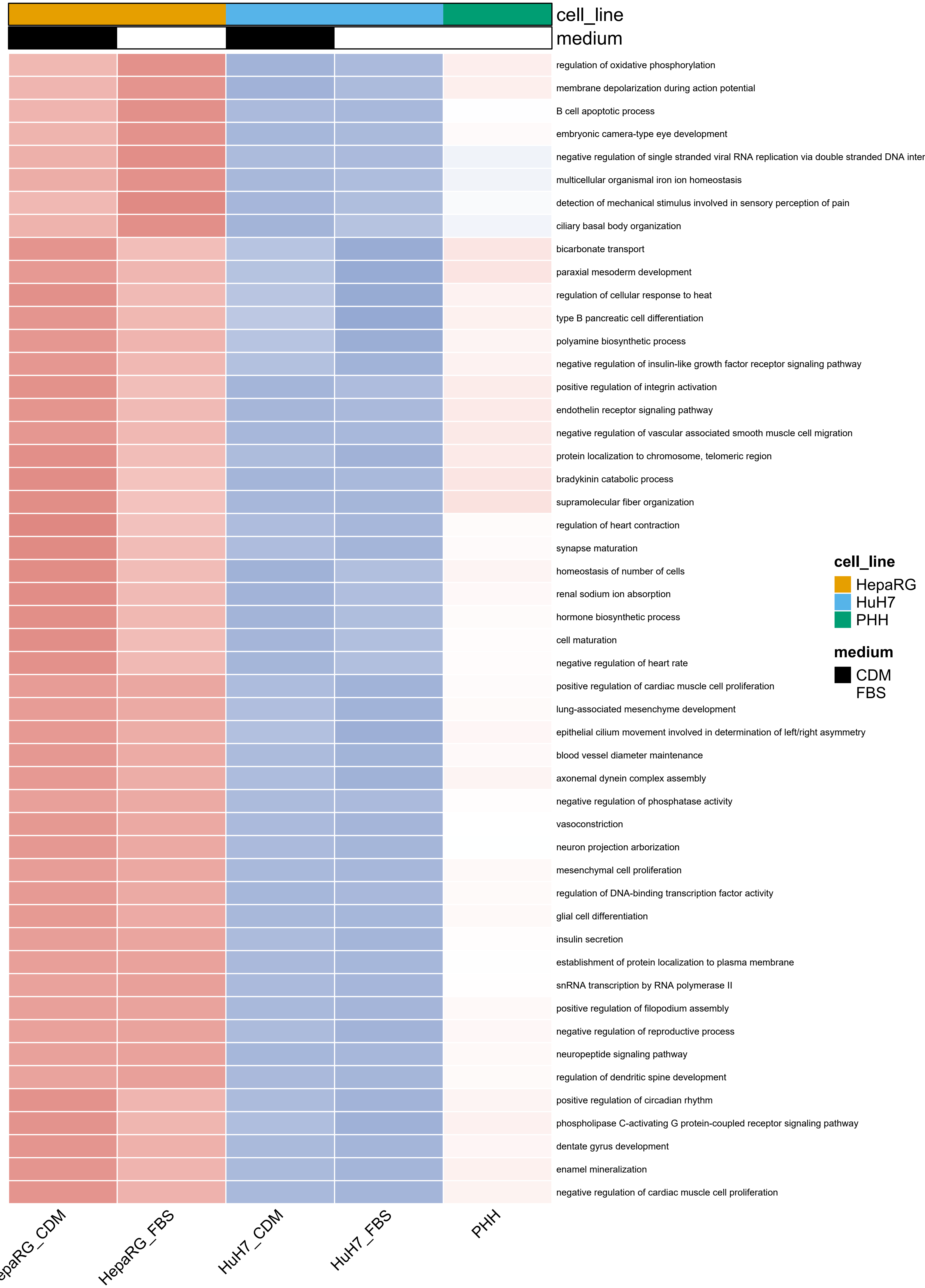
cell_line
medium
regulation of oxidative phosphorylation
membrane depolarization during action potential
B cell apoptotic process
embryonic camera-type eye development
negative regulation of single stranded viral RNA replication via double stranded DNA inter
multicellular organismal iron ion homeostasis
detection of mechanical stimulus involved in sensory perception of pain
ciliary basal body organization
bicarbonate transport
paraxial mesoderm development
regulation of cellular response to heat
type B pancreatic cell differentiation
polyamine biosynthetic process
negative regulation of insulin-like growth factor receptor signaling pathway
positive regulation of integrin activation
endothelin receptor signaling pathway
negative regulation of vascular associated smooth muscle cell migration
protein localization to chromosome, telomeric region
bradykinin catabolic process
supramolecular fiber organization
regulation of heart contraction
synapse maturation
homeostasis of number of cells
renal sodium ion absorption
hormone biosynthetic process
cell maturation
negative regulation of heart rate
positive regulation of cardiac muscle cell proliferation
lung-associated mesenchyme development
epithelial cilium movement involved in determination of left/right asymmetry
blood vessel diameter maintenance
axonemal dynein complex assembly
negative regulation of phosphatase activity
vasoconstriction
neuron projection arborization
mesenchymal cell proliferation
regulation of DNA-binding transcription factor activity
glial cell differentiation
insulin secretion
establishment of protein localization to plasma membrane
snRNA transcription by RNA polymerase II
positive regulation of filopodium assembly
negative regulation of reproductive process
neuropeptide signaling pathway
regulation of dendritic spine development
positive regulation of circadian rhythm
phospholipase C-activating G protein-coupled receptor signaling pathway
dentate gyrus development
enamel mineralization
negative regulation of cardiac muscle cell proliferation
epaRG_CDM
HepaRG_FBS
HuH7_CDM
HuH7_FBS
PHH
cell_line
HepaRG
HuH7
PHH
medium
CDM
FBS

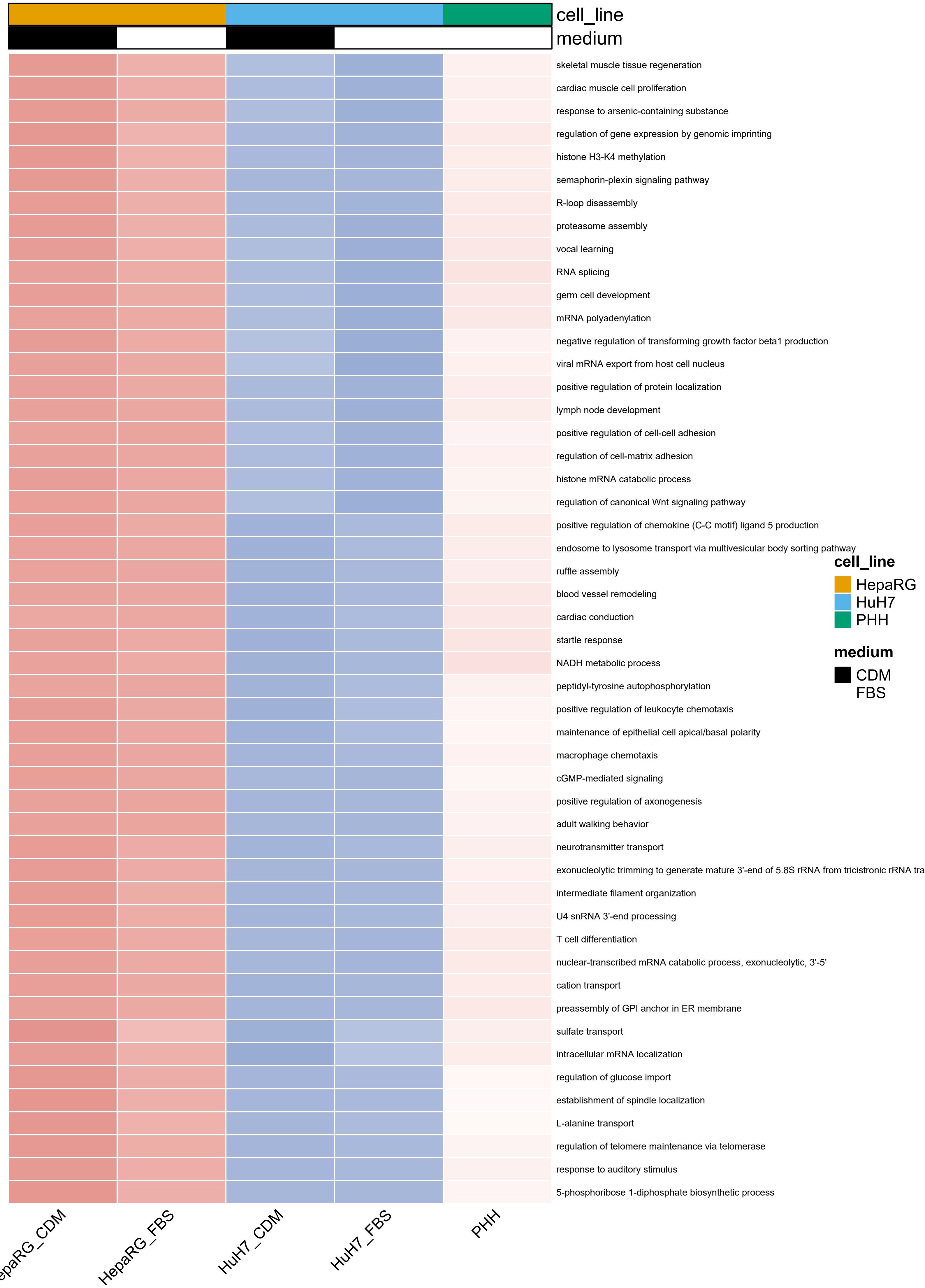
cell_line
medium
skeletal muscle tissue regeneration
cardiac muscle cell proliferation
response to arsenic-containing substance
regulation of gene expression by genomic imprinting
histone H3-K4 methylation
semaphorin-plexin signaling pathway
R-loop disassembly
proteasome assembly
vocal learning
RNA splicing
germ cell development
mRNA polyadenylation
negative regulation of transforming growth factor beta1 production
viral mRNA export from host cell nucleus
positive regulation of protein localization
lymph node development
positive regulation of cell-cell adhesion
regulation of cell-matrix adhesion
histone mRNA catabolic process
regulation of canonical Wnt signaling pathway
positive regulation of chemokine (C-C motif) ligand 5 production
endosome to lysosome transport via multivesicular body sorting pathway
ruffle assembly
blood vessel remodeling
cardiac conduction
startle response
NADH metabolic process
peptidyl-tyrosine autophosphorylation
positive regulation of leukocyte chemotaxis
maintenance of epithelial cell apical/basal polarity
macrophage chemotaxis
cGMP-mediated signaling
positive regulation of axonogenesis
adult walking behavior
neurotransmitter transport
exonucleolytic trimming to generate mature 3'-end of 5.8S rRNA from tricistronic rRNA tra
intermediate filament organization
U4 snRNA 3'-end processing
T cell differentiation
nuclear-transcribed mRNA catabolic process, exonucleolytic, 3'-5'
cation transport
preassembly of GPI anchor in ER membrane
sulfate transport
intracellular mRNA localization
regulation of glucose import
establishment of spindle localization
L-alanine transport
regulation of telomere maintenance via telomerase
response to auditory stimulus
5-phosphoribose 1-diphosphate biosynthetic process
epaRG_CDM
HepaRG_FBS
HuH7_CDM
HuH7_FBS
PHH
cell_line
HepaRG
HuH7
PHH
medium
CDM
FBS

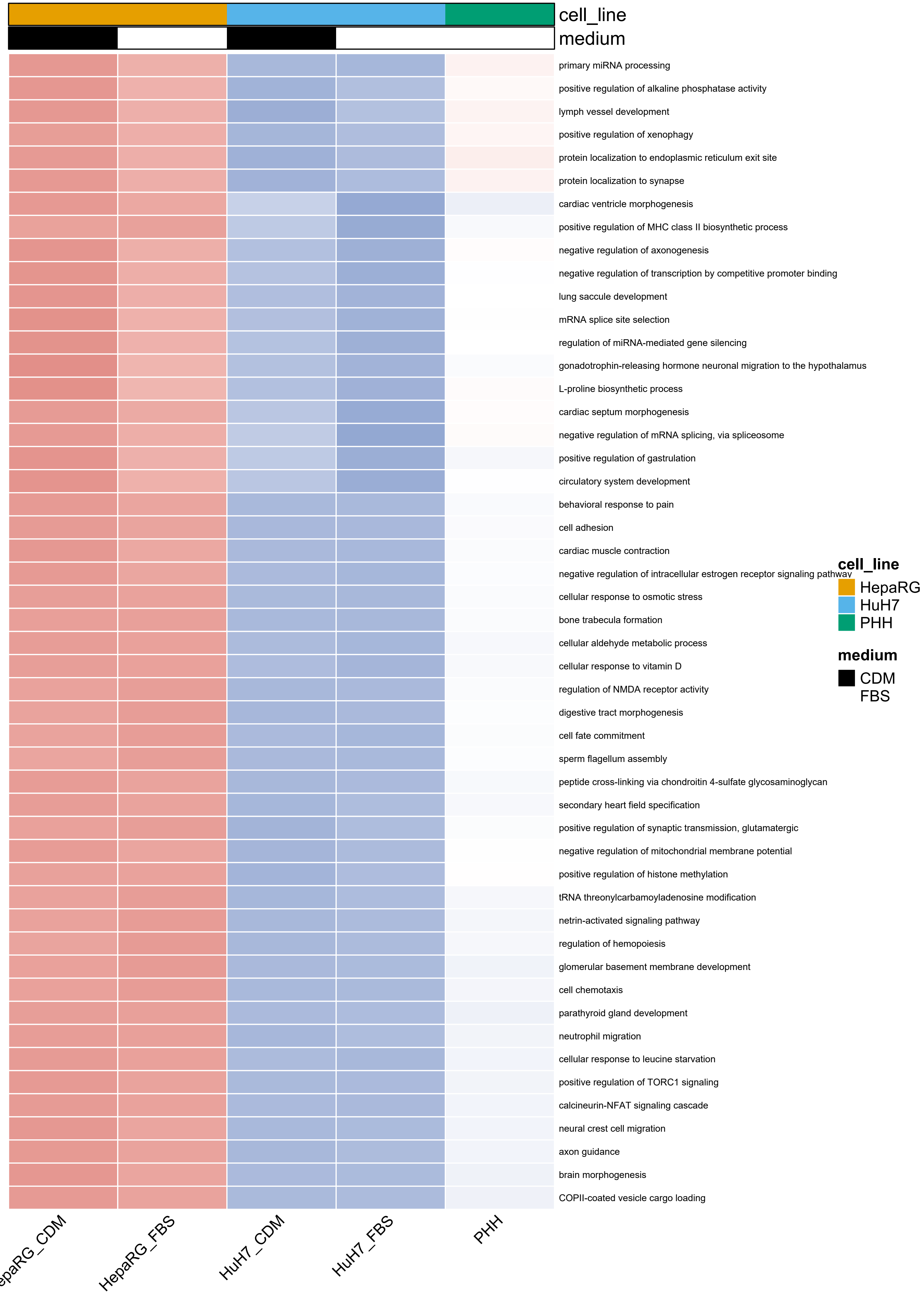
cell_line
medium
primary miRNA processing
positive regulation of alkaline phosphatase activity
lymph vessel development
positive regulation of xenophagy
protein localization to endoplasmic reticulum exit site
protein localization to synapse
cardiac ventricle morphogenesis
positive regulation of MHC class II biosynthetic process
negative regulation of axonogenesis
negative regulation of transcription by competitive promoter binding
lung saccule development
mRNA splice site selection
regulation of miRNA-mediated gene silencing
gonadotrophin-releasing hormone neuronal migration to the hypothalamus
L-proline biosynthetic process
cardiac septum morphogenesis
negative regulation of mRNA splicing, via spliceosome
positive regulation of gastrulation
circulatory system development
behavioral response to pain
cell adhesion
cardiac muscle contraction
negative regulation of intracellular estrogen receptor signaling pathway
cellular response to osmotic stress
bone trabecula formation
cellular aldehyde metabolic process
cellular response to vitamin D
regulation of NMDA receptor activity
digestive tract morphogenesis
cell fate commitment
sperm flagellum assembly
peptide cross-linking via chondroitin 4-sulfate glycosaminoglycan
secondary heart field specification
positive regulation of synaptic transmission, glutamatergic
negative regulation of mitochondrial membrane potential
positive regulation of histone methylation
tRNA threonylcarbamoyladenosine modification
netrin-activated signaling pathway
regulation of hemopoiesis
glomerular basement membrane development
cell chemotaxis
parathyroid gland development
neutrophil migration
cellular response to leucine starvation
positive regulation of TORC1 signaling
calcineurin-NFAT signaling cascade
neural crest cell migration
axon guidance
brain morphogenesis
COPII-coated vesicle cargo loading
HepaRG_CDM
HepaRG_FBS
HuH7_CDM
HuH7_FBS
PHH
cell_line
HepaRG
HuH7
PHH
medium
CDM
FBS

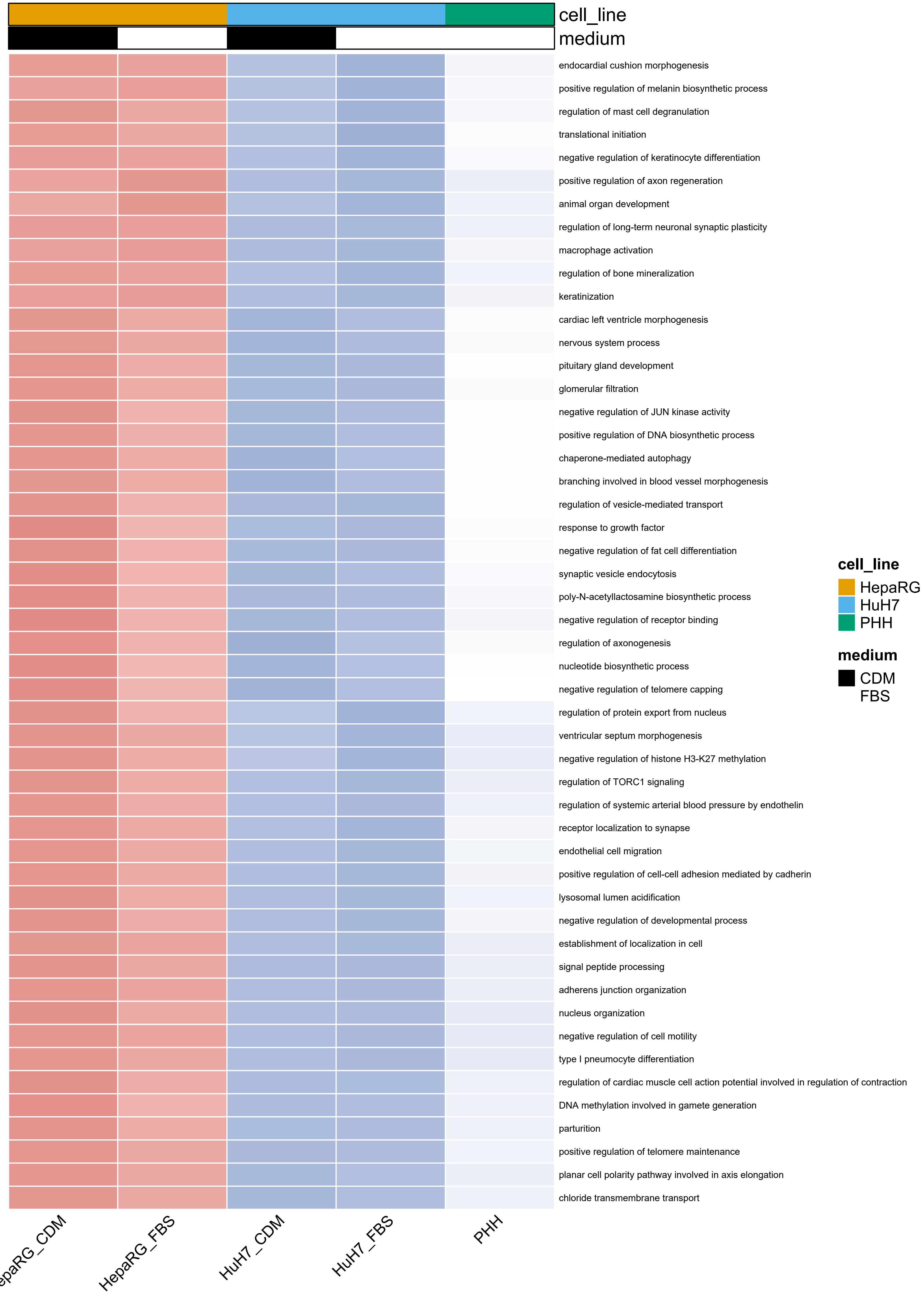
cell_line
medium
endocardial cushion morphogenesis
positive regulation of melanin biosynthetic process
regulation of mast cell degranulation
translational initiation
negative regulation of keratinocyte differentiation
positive regulation of axon regeneration
animal organ development
regulation of long-term neuronal synaptic plasticity
macrophage activation
regulation of bone mineralization
keratinization
cardiac left ventricle morphogenesis
nervous system process
pituitary gland development
glomerular filtration
negative regulation of JUN kinase activity
positive regulation of DNA biosynthetic process
chaperone-mediated autophagy
branching involved in blood vessel morphogenesis
regulation of vesicle-mediated transport
response to growth factor
negative regulation of fat cell differentiation
synaptic vesicle endocytosis
poly-N-acetyllactosamine biosynthetic process
negative regulation of receptor binding
regulation of axonogenesis
nucleotide biosynthetic process
negative regulation of telomere capping
regulation of protein export from nucleus
ventricular septum morphogenesis
negative regulation of histone H3-K27 methylation
regulation of TORC1 signaling
regulation of systemic arterial blood pressure by endothelin
receptor localization to synapse
endothelial cell migration
positive regulation of cell-cell adhesion mediated by cadherin
lysosomal lumen acidification
negative regulation of developmental process
establishment of localization in cell
signal peptide processing
adherens junction organization
nucleus organization
negative regulation of cell motility
type I pneumocyte differentiation
regulation of cardiac muscle cell action potential involved in regulation of contraction
DNA methylation involved in gamete generation
parturition
positive regulation of telomere maintenance
planar cell polarity pathway involved in axis elongation
chloride transmembrane transport
epaRG_CDM
HepaRG_FBS
HuH7_CDM
HuH7_FBS
PHH
cell_line
HepaRG
HuH7
PHH
medium
CDM
FBS

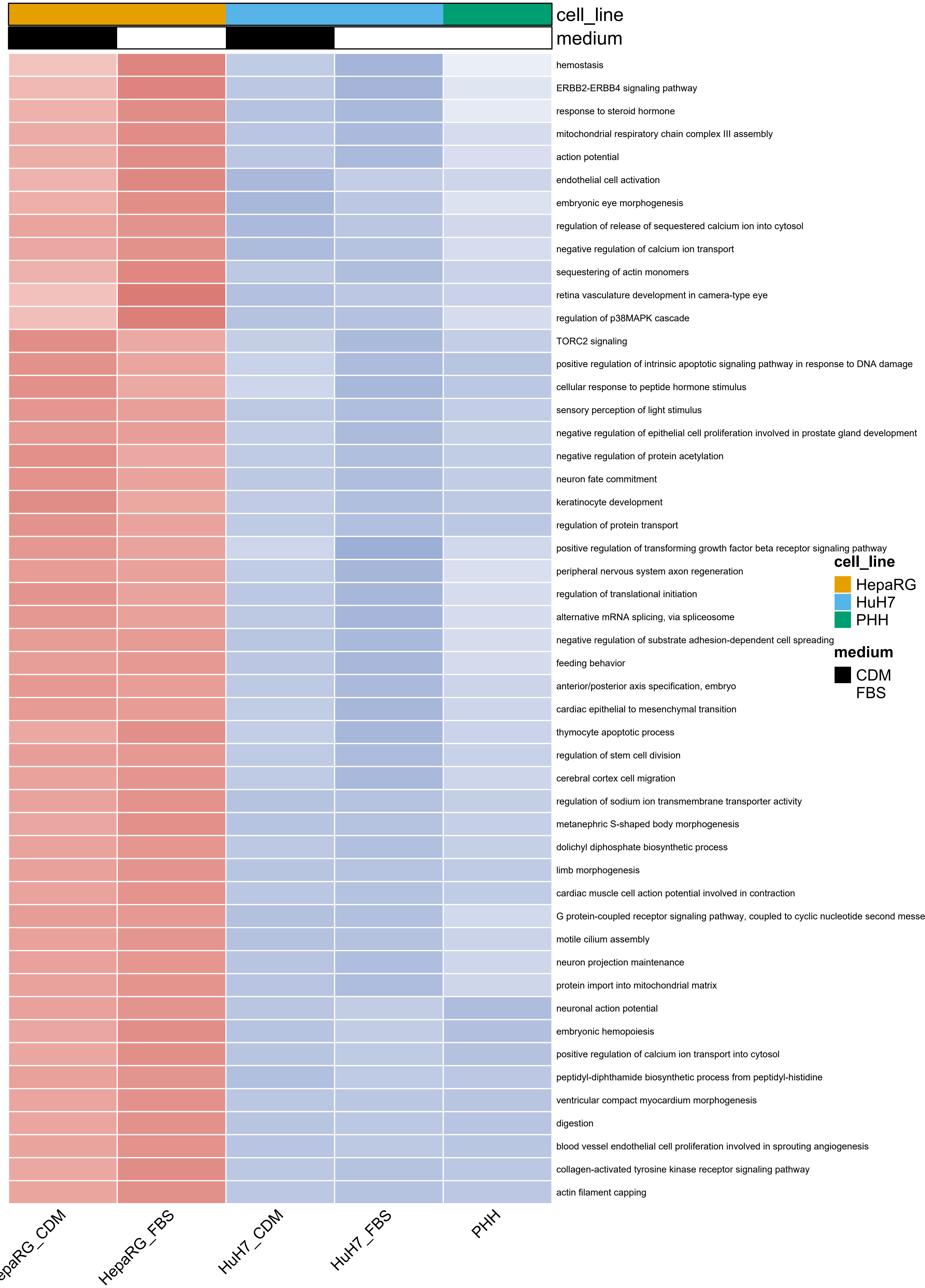

cell_line
medium
hemostasis
ERBB2-ERBB4 signaling pathway
response to steroid hormone
mitochondrial respiratory chain complex III assembly
action potential
endothelial cell activation
embryonic eye morphogenesis
regulation of release of sequestered calcium ion into cytosol
negative regulation of calcium ion transport
sequestering of actin monomers
retina vasculature development in camera-type eye
regulation of p38MAPK cascade
TORC2 signaling
positive regulation of intrinsic apoptotic signaling pathway in response to DNA damage
cellular response to peptide hormone stimulus
sensory perception of light stimulus
negative regulation of epithelial cell proliferation involved in prostate gland development
negative regulation of protein acetylation
neuron fate commitment
keratinocyte development
regulation of protein transport
positive regulation of transforming growth factor beta receptor signaling pathway
peripheral nervous system axon regeneration
regulation of translational initiation
alternative mRNA splicing, via spliceosome
negative regulation of substrate adhesion-dependent cell spreading
feeding behavior
anterior/posterior axis specification, embryo
cardiac epithelial to mesenchymal transition
thymocyte apoptotic process
regulation of stem cell division
cerebral cortex cell migration
regulation of sodium ion transmembrane transporter activity
metanephric S-shaped body morphogenesis
dolichyl diphosphate biosynthetic process
limb morphogenesis
cardiac muscle cell action potential involved in contraction
G protein-coupled receptor signaling pathway, coupled to cyclic nucleotide second messe
motile cilium assembly
neuron projection maintenance
protein import into mitochondrial matrix
neuronal action potential
embryonic hemopoiesis
positive regulation of calcium ion transport into cytosol
peptidyl-diphthamide biosynthetic process from peptidyl-histidine
ventricular compact myocardium morphogenesis
digestion
blood vessel endothelial cell proliferation involved in sprouting angiogenesis
collagen-activated tyrosine kinase receptor signaling pathway
actin filament capping
cell_line
HepaRG
HuH7
PHH
medium
CDM
FBS
epaRG_CDM
HepaRG_FBS
HuH7_CDM
HuH7_FBS
PHH

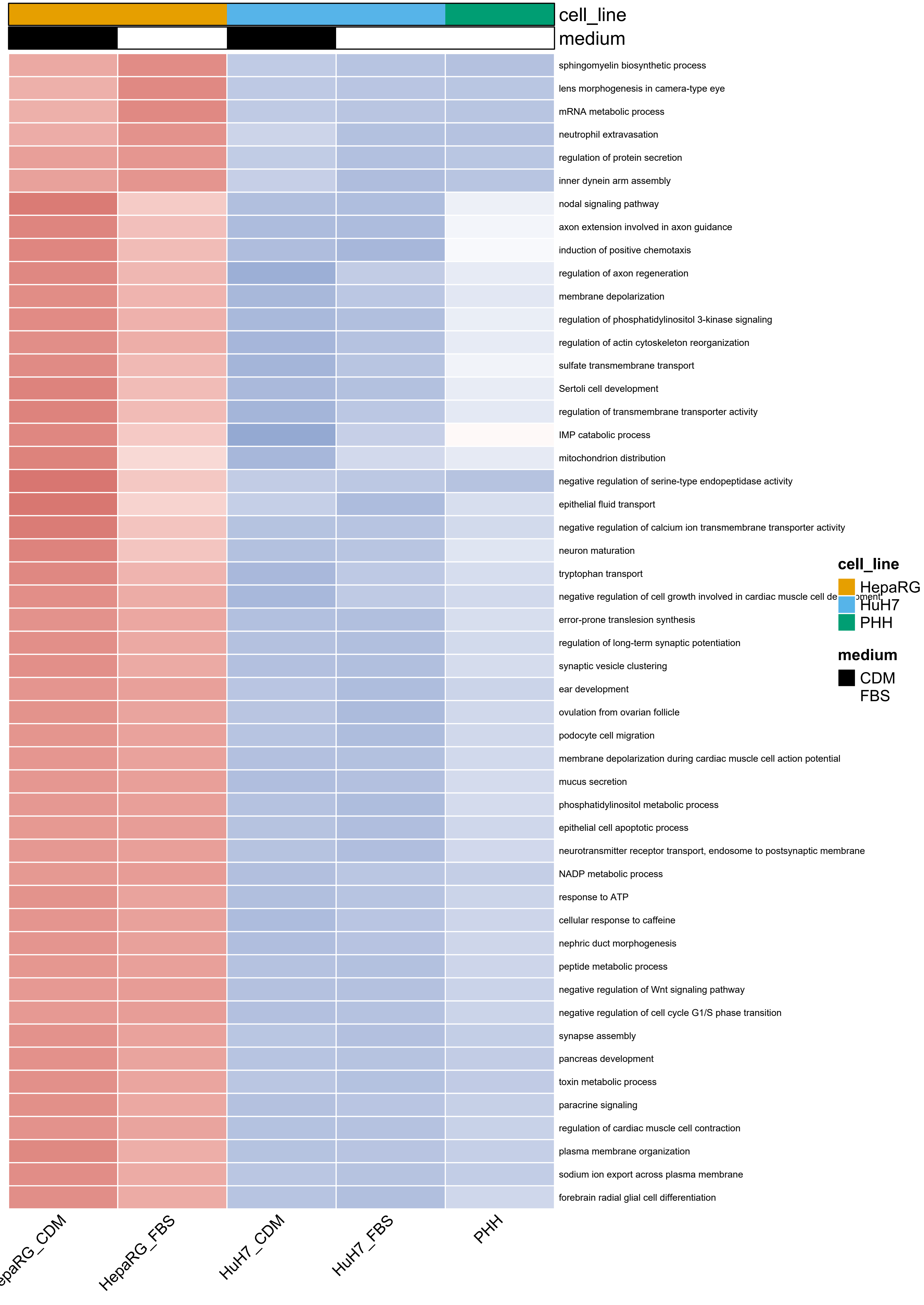

cell_line
medium
sphingomyelin biosynthetic process
lens morphogenesis in camera-type eye
mRNA metabolic process
neutrophil extravasation
regulation of protein secretion
inner dynein arm assembly
nodal signaling pathway
axon extension involved in axon guidance
induction of positive chemotaxis
regulation of axon regeneration
membrane depolarization
regulation of phosphatidylinositol 3-kinase signaling
regulation of actin cytoskeleton reorganization
sulfate transmembrane transport
Sertoli cell development
regulation of transmembrane transporter activity
IMP catabolic process
mitochondrion distribution
negative regulation of serine-type endopeptidase activity
epithelial fluid transport
negative regulation of calcium ion transmembrane transporter activity
neuron maturation
tryptophan transport
error-prone translesion synthesis
regulation of long-term synaptic potentiation
synaptic vesicle clustering
ear development
ovulation from ovarian follicle
podocyte cell migration
membrane depolarization during cardiac muscle cell action potential
mucus secretion
phosphatidylinositol metabolic process
epithelial cell apoptotic process
neurotransmitter receptor transport, endosome to postsynaptic membrane
NADP metabolic process
response to ATP
cellular response to caffeine
nephric duct morphogenesis
peptide metabolic process
negative regulation of Wnt signaling pathway
negative regulation of cell cycle G1/S phase transition
synapse assembly
pancreas development
toxin metabolic process
paracrine signaling
regulation of cardiac muscle cell contraction
plasma membrane organization
sodium ion export across plasma membrane
forebrain radial glial cell differentiation
cell_line
HepaRG
HuH7
PHH
medium
CDM
FBS
HepaRG_FBS
HuH7_CDM
HuH7_FBS
PHH

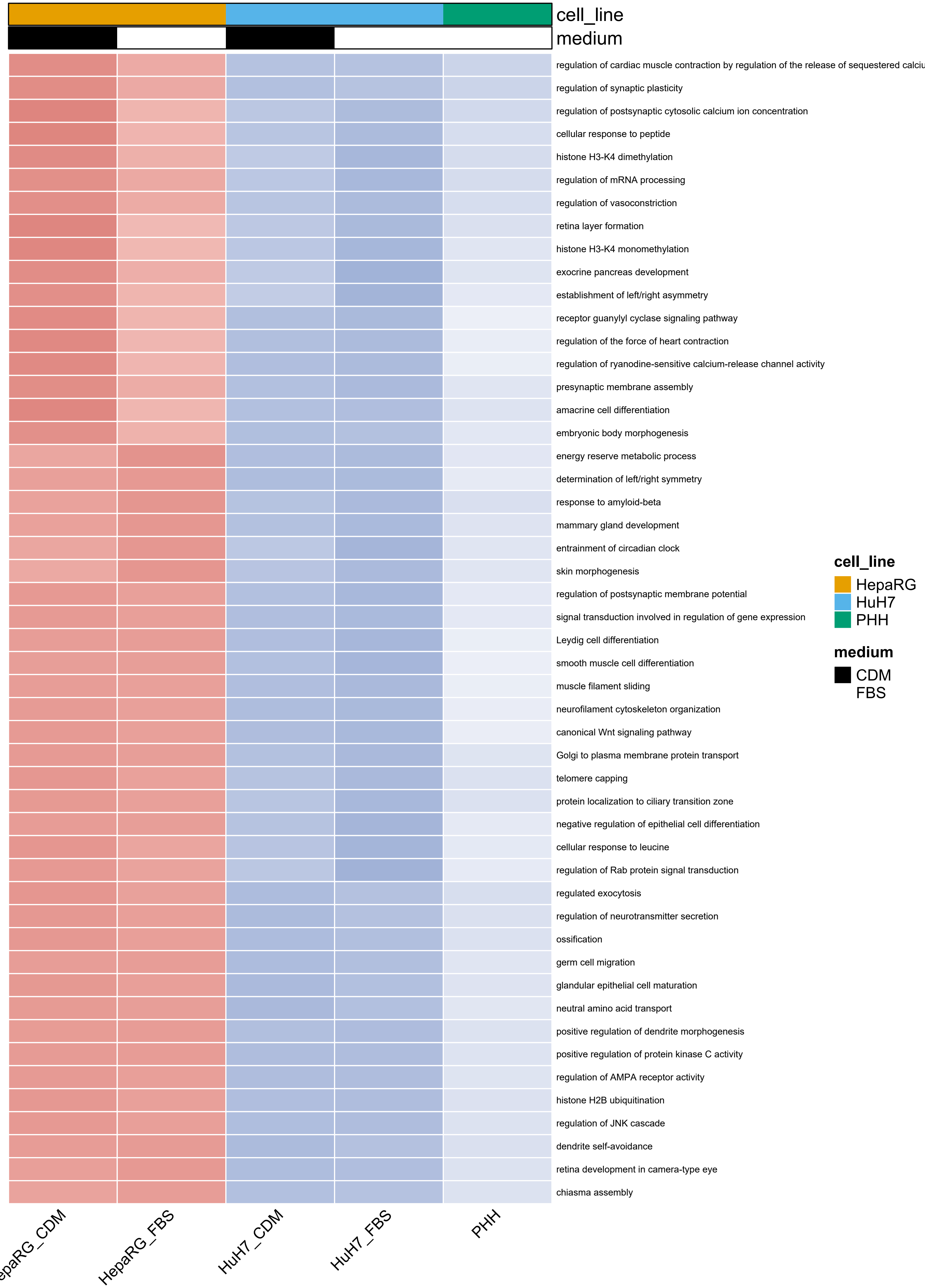
cell_line
medium
regulation of cardiac muscle contraction by regulation of the release of sequestered calciu
regulation of synaptic plasticity
regulation of postsynaptic cytosolic calcium ion concentration
cellular response to peptide
histone H3-K4 dimethylation
regulation of mRNA processing
regulation of vasoconstriction
retina layer formation
histone H3-K4 monomethylation
exocrine pancreas development
establishment of left/right asymmetry
receptor guanylyl cyclase signaling pathway
regulation of the force of heart contraction
regulation of ryanodine-sensitive calcium-release channel activity
presynaptic membrane assembly
amacrine cell differentiation
embryonic body morphogenesis
energy reserve metabolic process
determination of left/right symmetry
response to amyloid-beta
mammary gland development
entrainment of circadian clock
skin morphogenesis
regulation of postsynaptic membrane potential
signal transduction involved in regulation of gene expression
Leydig cell differentiation
smooth muscle cell differentiation
muscle filament sliding
neurofilament cytoskeleton organization
canonical Wnt signaling pathway
Golgi to plasma membrane protein transport
telomere capping
protein localization to ciliary transition zone
negative regulation of epithelial cell differentiation
cellular response to leucine
regulation of Rab protein signal transduction
regulated exocytosis
regulation of neurotransmitter secretion
ossification
germ cell migration
glandular epithelial cell maturation
neutral amino acid transport
positive regulation of dendrite morphogenesis
positive regulation of protein kinase C activity
regulation of AMPA receptor activity
histone H2B ubiquitination
regulation of JNK cascade
dendrite self-avoidance
retina development in camera-type eye
chiasma assembly
epaRG_CDM
HepaRG_FBS
HuH7_CDM
HuH7_FBS
PHH
cell_line
HepaRG
HuH7
PHH
medium
CDM
FBS

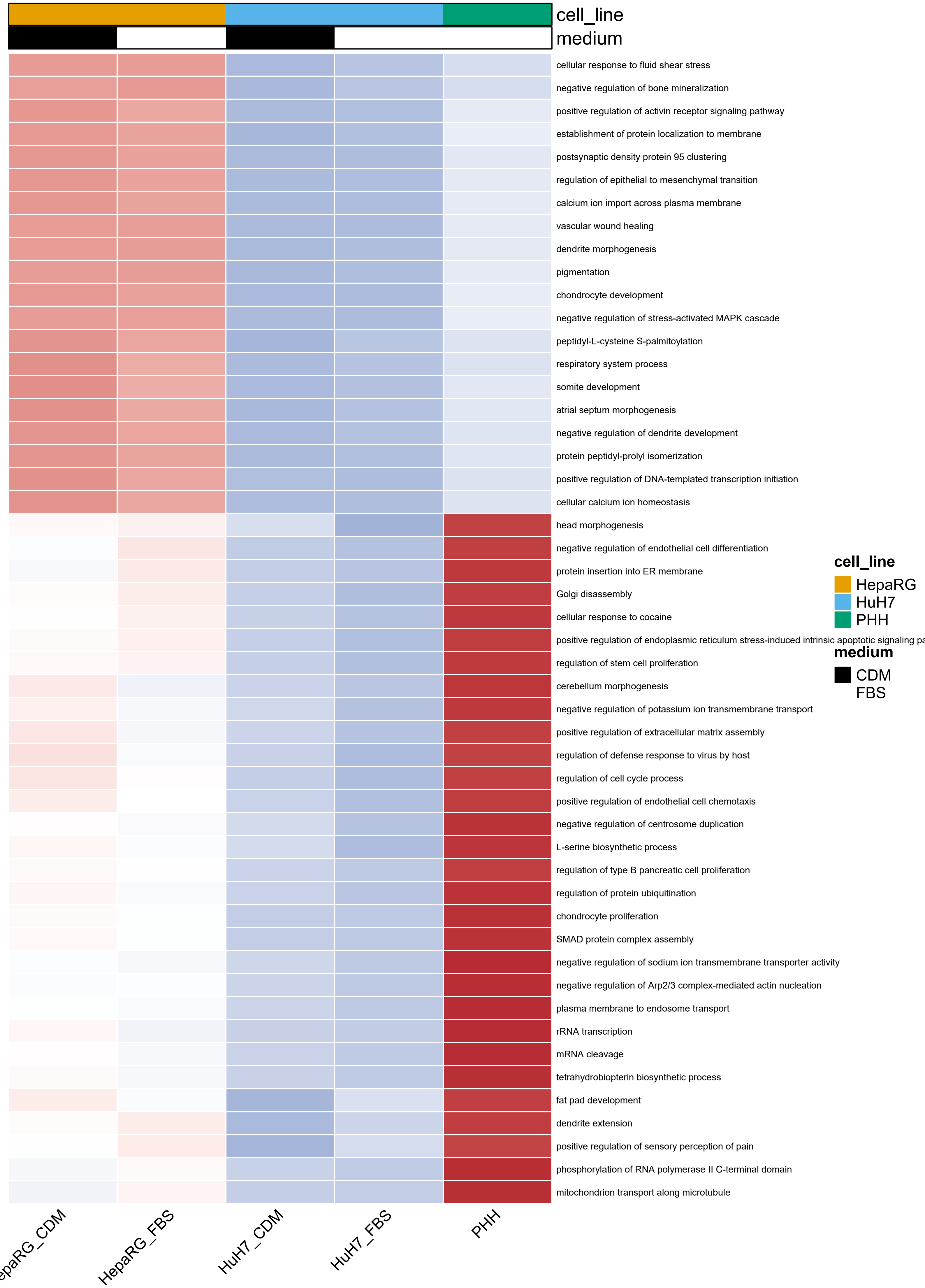
cell_line
medium
cellular response to fluid shear stress
negative regulation of bone mineralization
positive regulation of activin receptor signaling pathway
establishment of protein localization to membrane
postsynaptic density protein 95 clustering
regulation of epithelial to mesenchymal transition
calcium ion import across plasma membrane
vascular wound healing
dendrite morphogenesis
pigmentation
chondrocyte development
negative regulation of stress-activated MAPK cascade
peptidyl-L-cysteine S-palmitoylation
respiratory system process
somite development
atrial septum morphogenesis
negative regulation of dendrite development
protein peptidyl-prolyl isomerization
positive regulation of DNA-templated transcription initiation
cellular calcium ion homeostasis
head morphogenesis
negative regulation of endothelial cell differentiation
protein insertion into ER membrane
Golgi disassembly
cellular response to cocaine
positive regulation of endoplasmic reticulum stress-induced intrinsic apoptotic signaling pa
regulation of stem cell proliferation
cerebellum morphogenesis
negative regulation of potassium ion transmembrane transport
positive regulation of extracellular matrix assembly
regulation of defense response to virus by host
regulation of cell cycle process
positive regulation of endothelial cell chemotaxis
negative regulation of centrosome duplication
L-serine biosynthetic process
regulation of type B pancreatic cell proliferation
regulation of protein ubiquitination
chondrocyte proliferation
SMAD protein complex assembly
negative regulation of sodium ion transmembrane transporter activity
negative regulation of Arp2/3 complex-mediated actin nucleation
plasma membrane to endosome transport
rRNA transcription
mRNA cleavage
tetrahydrobiopterin biosynthetic process
fat pad development
dendrite extension
positive regulation of sensory perception of pain
phosphorylation of RNA polymerase II C-terminal domain
mitochondrion transport along microtubule
epaRG_CDM
HepaRG_FBS
HuH7_CDM
HuH7_FBS
PHH
cell_line
HepaRG
HuH7
PHH
medium
CDM
FBS

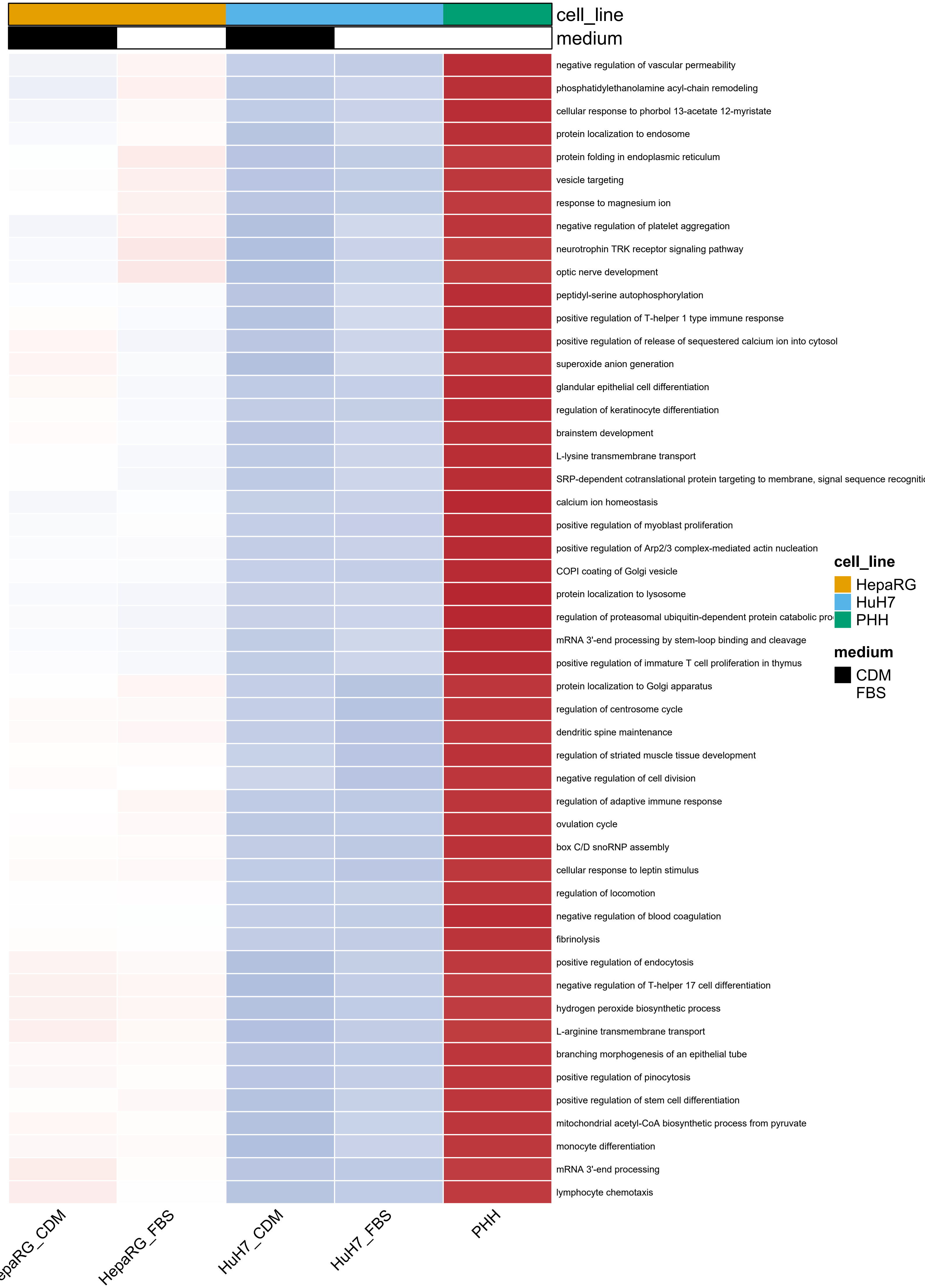

cell_line
medium
negative regulation of vascular permeability
phosphatidylethanolamine acyl-chain remodeling
cellular response to phorbol 13-acetate 12-myristate
protein localization to endosome
protein folding in endoplasmic reticulum
vesicle targeting
response to magnesium ion
negative regulation of platelet aggregation
neurotrophin TRK receptor signaling pathway
optic nerve development
peptidyl-serine autophosphorylation
positive regulation of T-helper 1 type immune response
positive regulation of release of sequestered calcium ion into cytosol
superoxide anion generation
glandular epithelial cell differentiation
regulation of keratinocyte differentiation
brainstem development
L-lysine transmembrane transport
SRP-dependent cotranslational protein targeting to membrane, signal sequence recogniti
calcium ion homeostasis
positive regulation of myoblast proliferation
positive regulation of Arp2/3 complex-mediated actin nucleation
COPI coating of Golgi vesicle
protein localization to lysosome
regulation of proteasomal ubiquitin-dependent protein catabolic pro
mRNA 3'-end processing by stem-loop binding and cleavage
positive regulation of immature T cell proliferation in thymus
protein localization to Golgi apparatus
regulation of centrosome cycle
dendritic spine maintenance
regulation of striated muscle tissue development
negative regulation of cell division
regulation of adaptive immune response
ovulation cycle
box C/D snoRNP assembly
cellular response to leptin stimulus
regulation of locomotion
negative regulation of blood coagulation
fibrinolysis
positive regulation of endocytosis
negative regulation of T-helper 17 cell differentiation
hydrogen peroxide biosynthetic process
L-arginine transmembrane transport
branching morphogenesis of an epithelial tube
positive regulation of pinocytosis
positive regulation of stem cell differentiation
mitochondrial acetyl-CoA biosynthetic process from pyruvate
monocyte differentiation
mRNA 3'-end processing
lymphocyte chemotaxis
cell_line
HepaRG
HuH7
PHH
medium
CDM
FBS
epaRG_CDM
HepaRG_FBS
HuH7_CDM
HuH7_FBS
PHH

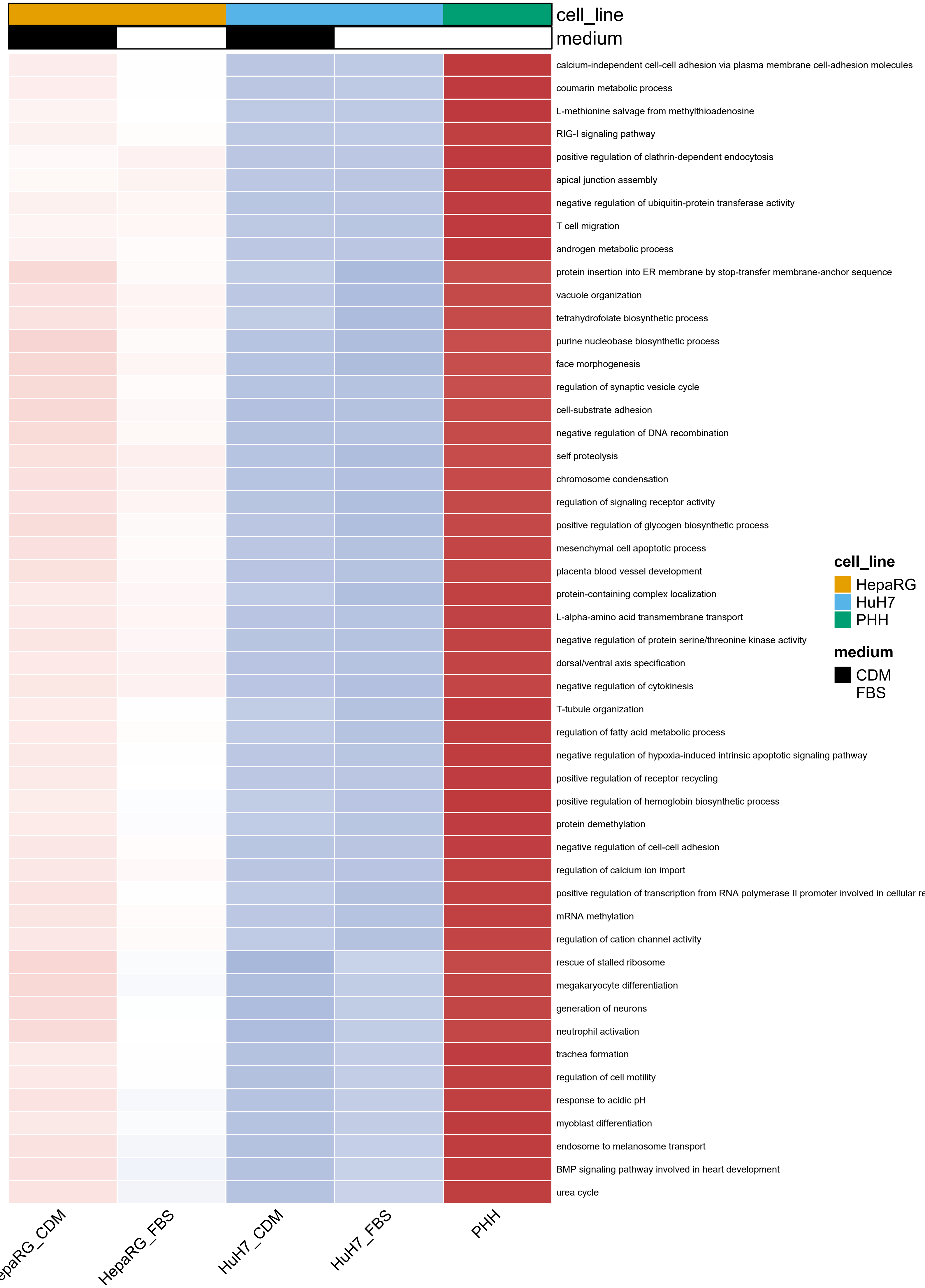
cell_line
medium
calcium-independent cell-cell adhesion via plasma membrane cell-adhesion molecules
coumarin metabolic process
L-methionine salvage from methylthioadenosine
RIG-I signaling pathway
positive regulation of clathrin-dependent endocytosis
apical junction assembly
negative regulation of ubiquitin-protein transferase activity
T cell migration
androgen metabolic process
protein insertion into ER membrane by stop-transfer membrane-anchor sequence
vacuole organization
tetrahydrofolate biosynthetic process
purine nucleobase biosynthetic process
face morphogenesis
regulation of synaptic vesicle cycle
cell-substrate adhesion
negative regulation of DNA recombination
self proteolysis
chromosome condensation
regulation of signaling receptor activity
positive regulation of glycogen biosynthetic process
mesenchymal cell apoptotic process
placenta blood vessel development
protein-containing complex localization
L-alpha-amino acid transmembrane transport
negative regulation of protein serine/threonine kinase activity
dorsal/ventral axis specification
negative regulation of cytokinesis
T-tubule organization
regulation of fatty acid metabolic process
negative regulation of hypoxia-induced intrinsic apoptotic signaling pathway
positive regulation of receptor recycling
positive regulation of hemoglobin biosynthetic process
protein demethylation
negative regulation of cell-cell adhesion
regulation of calcium ion import
positive regulation of transcription from RNA polymerase II promoter involved in cellular re
mRNA methylation
regulation of cation channel activity
rescue of stalled ribosome
megakaryocyte differentiation
generation of neurons
neutrophil activation
trachea formation
regulation of cell motility
response to acidic pH
myoblast differentiation
endosome to melanosome transport
BMP signaling pathway involved in heart development
urea cycle
cell_line
HepaRG
HuH7
PHH
medium
CDM
FBS
epaRG_CDM
HepaRG_FBS
HuH7_CDM
HuH7_FBS
PHH

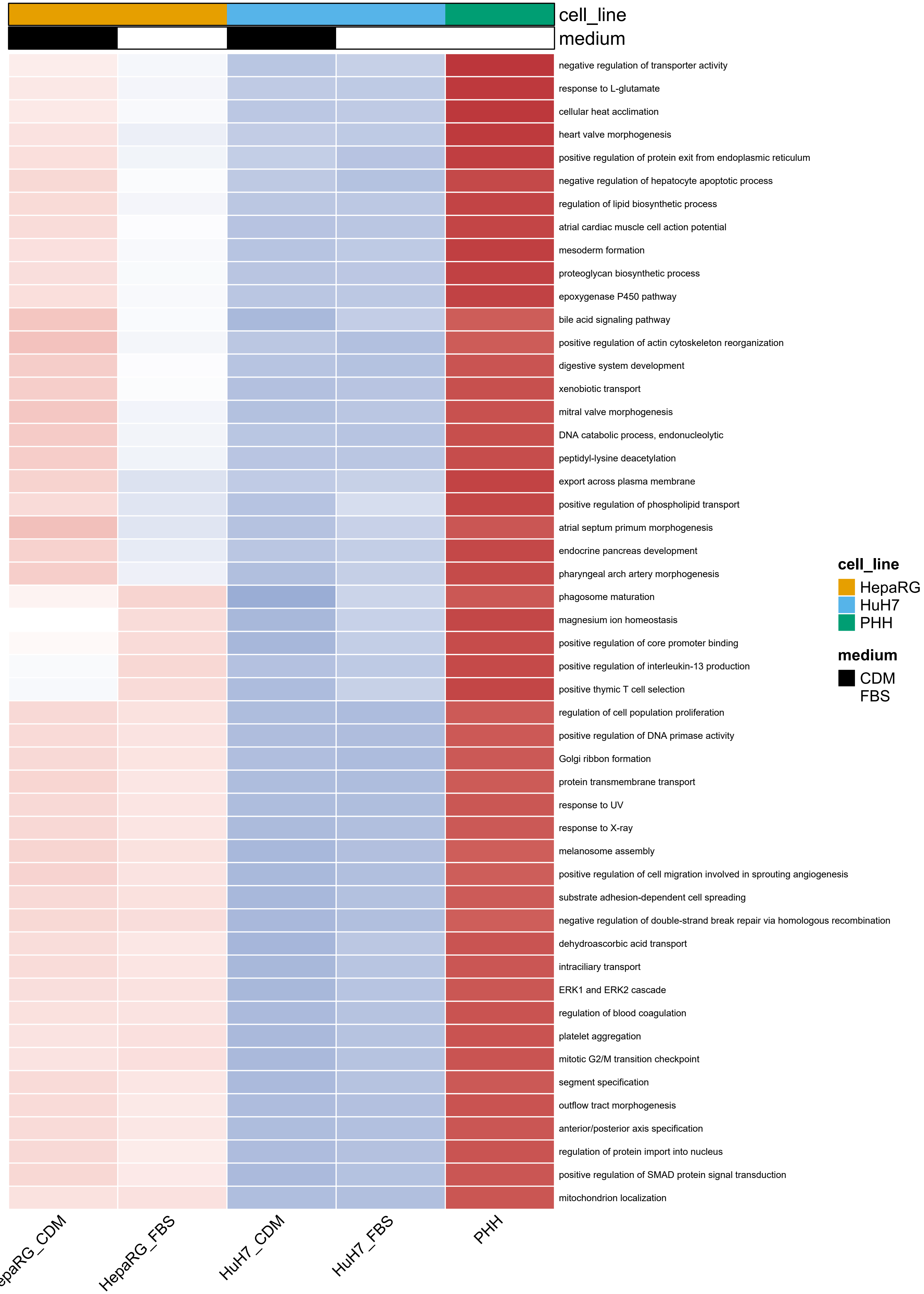
cell_line
medium
negative regulation of transporter activity
response to L-glutamate
cellular heat acclimation
heart valve morphogenesis
positive regulation of protein exit from endoplasmic reticulum
negative regulation of hepatocyte apoptotic process
regulation of lipid biosynthetic process
atrial cardiac muscle cell action potential
mesoderm formation
proteoglycan biosynthetic process
epoxygenase P450 pathway
bile acid signaling pathway
positive regulation of actin cytoskeleton reorganization
digestive system development
xenobiotic transport
mitral valve morphogenesis
DNA catabolic process, endonucleolytic
peptidyl-lysine deacetylation
export across plasma membrane
positive regulation of phospholipid transport
atrial septum primum morphogenesis
endocrine pancreas development
pharyngeal arch artery morphogenesis
phagosome maturation
magnesium ion homeostasis
positive regulation of core promoter binding
positive regulation of interleukin-13 production
positive thymic T cell selection
regulation of cell population proliferation
positive regulation of DNA primase activity
Golgi ribbon formation
protein transmembrane transport
response to UV
response to X-ray
melanosome assembly
positive regulation of cell migration involved in sprouting angiogenesis
substrate adhesion-dependent cell spreading
negative regulation of double-strand break repair via homologous recombination
dehydroascorbic acid transport
intraciliary transport
ERK1 and ERK2 cascade
regulation of blood coagulation
platelet aggregation
mitotic G2/M transition checkpoint
segment specification
outflow tract morphogenesis
anterior/posterior axis specification
regulation of protein import into nucleus
positive regulation of SMAD protein signal transduction
mitochondrion localization
cell_line
HepaRG
HuH7
PHH
medium
CDM
FBS
HepaRG_FBS
HuH7_CDM
HuH7_FBS
PHH

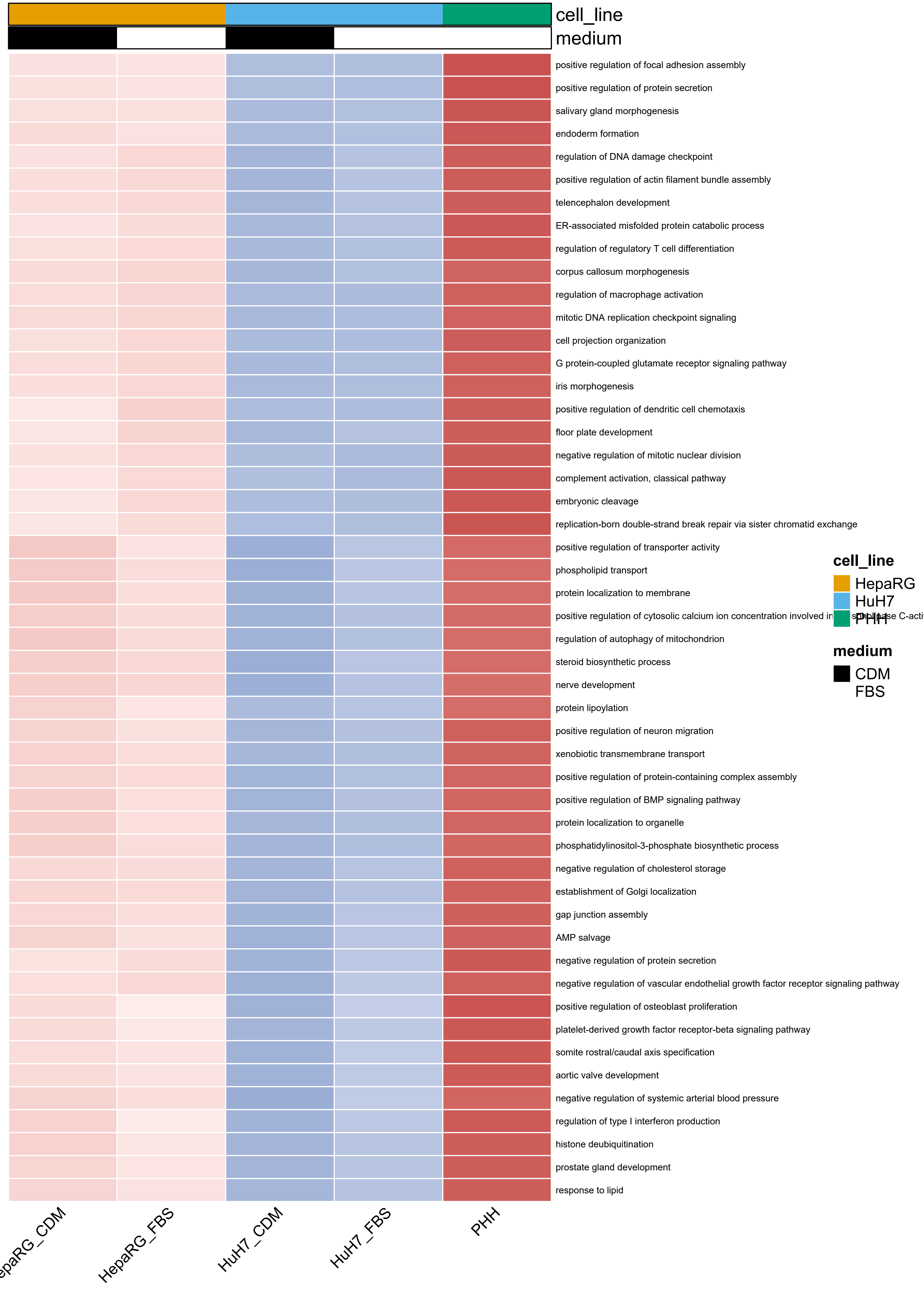
cell_line
medium
positive regulation of focal adhesion assembly
positive regulation of protein secretion
salivary gland morphogenesis
endoderm formation
regulation of DNA damage checkpoint
positive regulation of actin filament bundle assembly
telencephalon development
ER-associated misfolded protein catabolic process
regulation of regulatory T cell differentiation
corpus callosum morphogenesis
regulation of macrophage activation
mitotic DNA replication checkpoint signaling
cell projection organization
G protein-coupled glutamate receptor signaling pathway
iris morphogenesis
positive regulation of dendritic cell chemotaxis
floor plate development
negative regulation of mitotic nuclear division
complement activation, classical pathway
embryonic cleavage
replication-born double-strand break repair via sister chromatid exchange
positive regulation of transporter activity
phospholipid transport
protein localization to membrane
positive regulation of cytosolic calcium ion concentration involved in
regulation of autophagy of mitochondrion
steroid biosynthetic process
nerve development
protein lipoylation
positive regulation of neuron migration
xenobiotic transmembrane transport
positive regulation of protein-containing complex assembly
positive regulation of BMP signaling pathway
protein localization to organelle
phosphatidylinositol-3-phosphate biosynthetic process
negative regulation of cholesterol storage
establishment of Golgi localization
gap junction assembly
AMP salvage
negative regulation of protein secretion
negative regulation of vascular endothelial growth factor receptor signaling pathway
positive regulation of osteoblast proliferation
platelet-derived growth factor receptor-beta signaling pathway
somite rostral/caudal axis specification
aortic valve development
negative regulation of systemic arterial blood pressure
regulation of type I interferon production
histone deubiquitination
prostate gland development
response to lipid
cell_line
HepaRG
HuH7
PHH
medium
CDM
FBS
epaRG_CDM
HepaRG_FBS
HuH7_CDM
HuH7_FBS
PHH

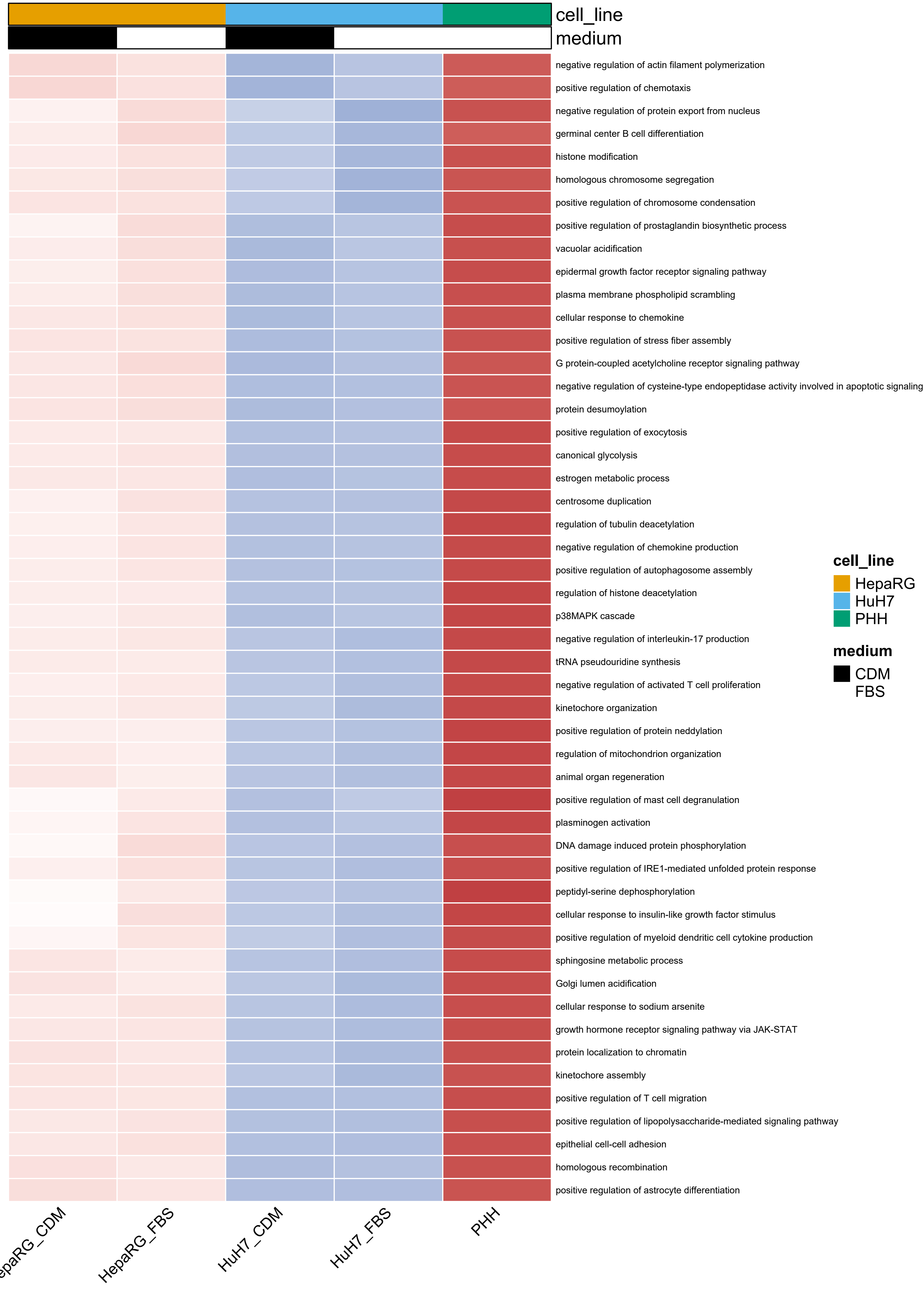
cell_line
medium
negative regulation of actin filament polymerization
positive regulation of chemotaxis
negative regulation of protein export from nucleus
germinal center B cell differentiation
histone modification
homologous chromosome segregation
positive regulation of chromosome condensation
positive regulation of prostaglandin biosynthetic process
vacuolar acidification
epidermal growth factor receptor signaling pathway
plasma membrane phospholipid scrambling
cellular response to chemokine
positive regulation of stress fiber assembly
G protein-coupled acetylcholine receptor signaling pathway
negative regulation of cysteine-type endopeptidase activity involved in apoptotic signaling
protein desumoylation
positive regulation of exocytosis
canonical glycolysis
estrogen metabolic process
centrosome duplication
regulation of tubulin deacetylation
negative regulation of chemokine production
positive regulation of autophagosome assembly
regulation of histone deacetylation
p38MAPK cascade
negative regulation of interleukin-17 production
tRNA pseudouridine synthesis
negative regulation of activated T cell proliferation
kinetochore organization
positive regulation of protein neddylation
regulation of mitochondrion organization
animal organ regeneration
positive regulation of mast cell degranulation
plasminogen activation
DNA damage induced protein phosphorylation
positive regulation of IRE1-mediated unfolded protein response
peptidyl-serine dephosphorylation
cellular response to insulin-like growth factor stimulus
positive regulation of myeloid dendritic cell cytokine production
sphingosine metabolic process
Golgi lumen acidification
cellular response to sodium arsenite
growth hormone receptor signaling pathway via JAK-STAT
protein localization to chromatin
kinetochore assembly
positive regulation of T cell migration
positive regulation of lipopolysaccharide-mediated signaling pathway
epithelial cell-cell adhesion
homologous recombination
positive regulation of astrocyte differentiation
epaRG_CDM
HepaRG_FBS
HuH7_CDM
HuH7_FBS
PHH
cell_line
HepaRG
HuH7
PHH
medium
CDM
FBS

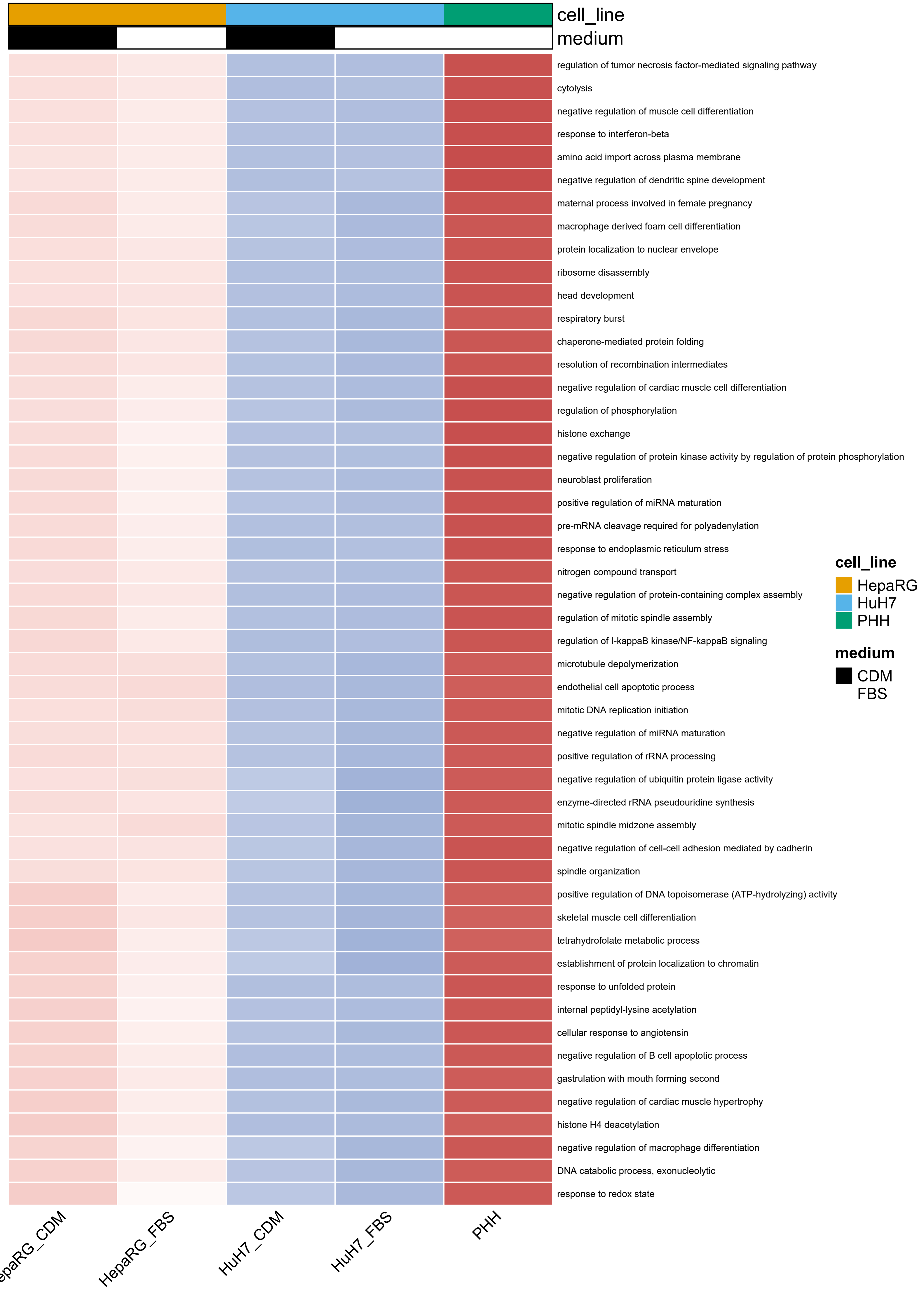

cell_line
medium
regulation of tumor necrosis factor-mediated signaling pathway
cytolysis
negative regulation of muscle cell differentiation
response to interferon-beta
amino acid import across plasma membrane
negative regulation of dendritic spine development
maternal process involved in female pregnancy
macrophage derived foam cell differentiation
protein localization to nuclear envelope
ribosome disassembly
head development
respiratory burst
chaperone-mediated protein folding
resolution of recombination intermediates
negative regulation of cardiac muscle cell differentiation
regulation of phosphorylation
histone exchange
negative regulation of protein kinase activity by regulation of protein phosphorylation
neuroblast proliferation
positive regulation of miRNA maturation
pre-mRNA cleavage required for polyadenylation
response to endoplasmic reticulum stress
nitrogen compound transport
negative regulation of protein-containing complex assembly
regulation of mitotic spindle assembly
regulation of I-kappaB kinase/NF-kappaB signaling
microtubule depolymerization
endothelial cell apoptotic process
mitotic DNA replication initiation
negative regulation of miRNA maturation
positive regulation of rRNA processing
negative regulation of ubiquitin protein ligase activity
enzyme-directed rRNA pseudouridine synthesis
mitotic spindle midzone assembly
negative regulation of cell-cell adhesion mediated by cadherin
spindle organization
positive regulation of DNA topoisomerase (ATP-hydrolyzing) activity
skeletal muscle cell differentiation
tetrahydrofolate metabolic process
establishment of protein localization to chromatin
response to unfolded protein
internal peptidyl-lysine acetylation
cellular response to angiotensin
negative regulation of B cell apoptotic process
gastrulation with mouth forming second
negative regulation of cardiac muscle hypertrophy
histone H4 deacetylation
negative regulation of macrophage differentiation
DNA catabolic process, exonucleolytic
response to redox state
cell_line
HepaRG
HuH7
PHH
medium
CDM
FBS
HepaRG_CDM
HepaRG_FBS
HuH7_CDM
HuH7_FBS
PHH

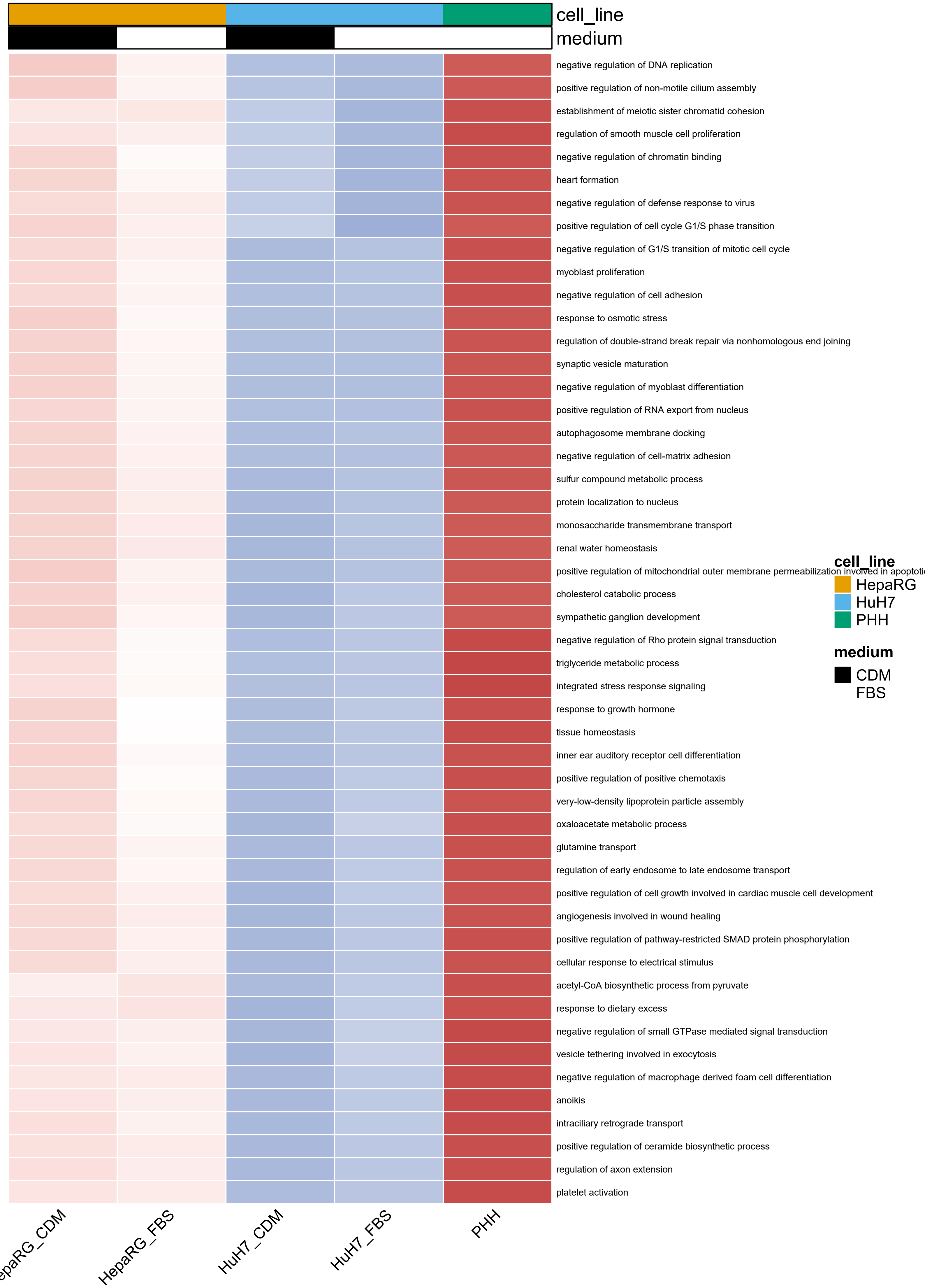
cell_line
medium
negative regulation of DNA replication
positive regulation of non-motile cilium assembly
establishment of meiotic sister chromatid cohesion
regulation of smooth muscle cell proliferation
negative regulation of chromatin binding
heart formation
negative regulation of defense response to virus
positive regulation of cell cycle G1/S phase transition
negative regulation of G1/S transition of mitotic cell cycle
myoblast proliferation
negative regulation of cell adhesion
response to osmotic stress
regulation of double-strand break repair via nonhomologous end joining
synaptic vesicle maturation
negative regulation of myoblast differentiation
positive regulation of RNA export from nucleus
autophagosome membrane docking
negative regulation of cell-matrix adhesion
sulfur compound metabolic process
protein localization to nucleus
monosaccharide transmembrane transport
renal water homeostasis
positive regulation of mitochondrial outer membrane permeabilization involved in apoptoti
cholesterol catabolic process
sympathetic ganglion development
negative regulation of Rho protein signal transduction
triglyceride metabolic process
integrated stress response signaling
response to growth hormone
tissue homeostasis
inner ear auditory receptor cell differentiation
positive regulation of positive chemotaxis
very-low-density lipoprotein particle assembly
oxaloacetate metabolic process
glutamine transport
regulation of early endosome to late endosome transport
positive regulation of cell growth involved in cardiac muscle cell development
angiogenesis involved in wound healing
positive regulation of pathway-restricted SMAD protein phosphorylation
cellular response to electrical stimulus
acetyl-CoA biosynthetic process from pyruvate
response to dietary excess
negative regulation of small GTPase mediated signal transduction
vesicle tethering involved in exocytosis
negative regulation of macrophage derived foam cell differentiation
anoikis
intraciliary retrograde transport
positive regulation of ceramide biosynthetic process
regulation of axon extension
platelet activation
cell_line
HepaRG
HuH7
PHH
medium
CDM
FBS
epaRG_CDM
HepaRG_FBS
HuH7_CDM
HuH7_FBS
PHH

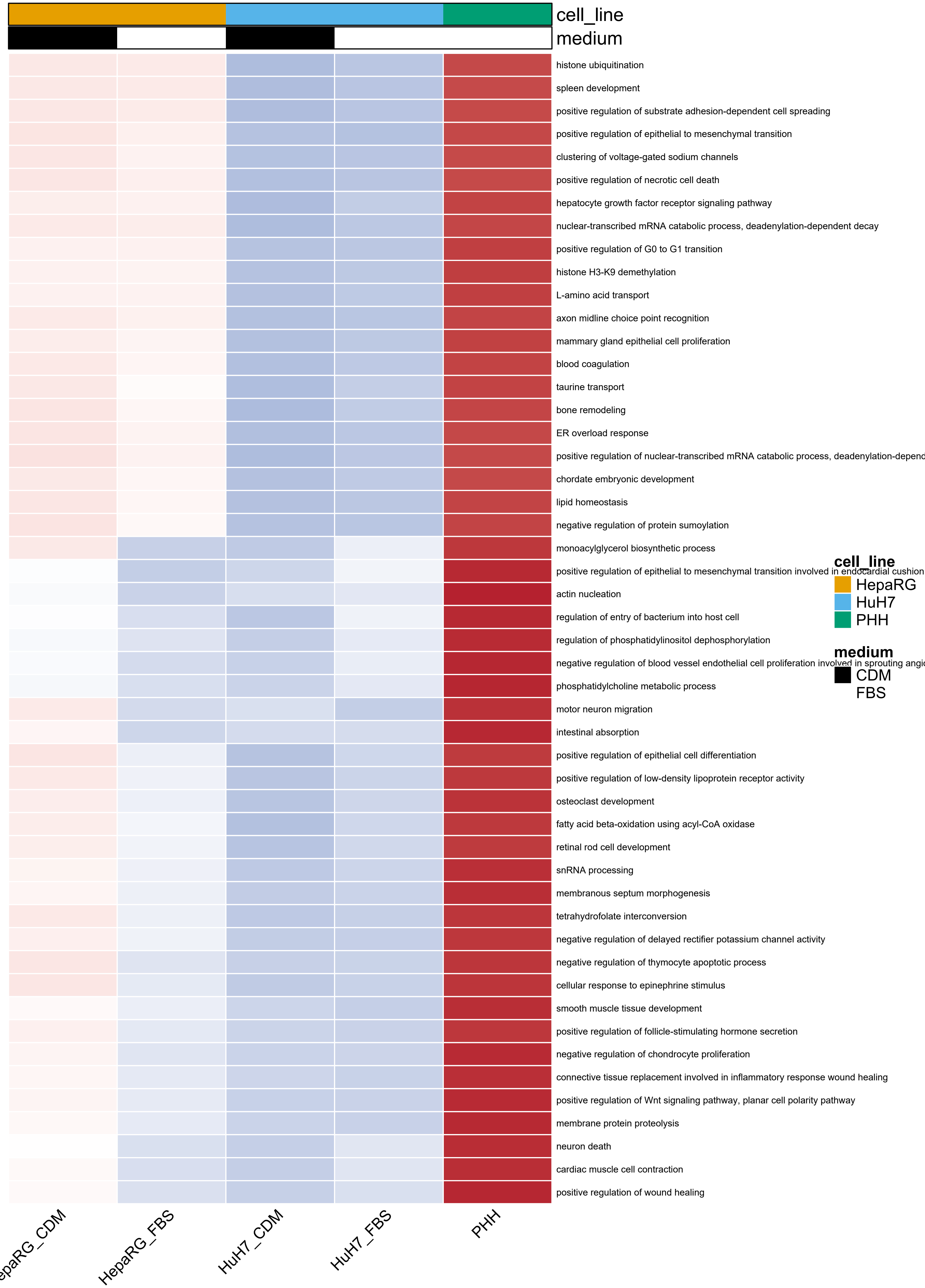
cell_line
medium
histone ubiquitination
spleen development
positive regulation of substrate adhesion-dependent cell spreading
positive regulation of epithelial to mesenchymal transition
clustering of voltage-gated sodium channels
positive regulation of necrotic cell death
hepatocyte growth factor receptor signaling pathway
nuclear-transcribed mRNA catabolic process, deadenylation-dependent decay
positive regulation of G0 to G1 transition
histone H3-K9 demethylation
L-amino acid transport
axon midline choice point recognition
mammary gland epithelial cell proliferation
blood coagulation
taurine transport
bone remodeling
ER overload response
positive regulation of nuclear-transcribed mRNA catabolic process, deadenylation-depend
chordate embryonic development
lipid homeostasis
negative regulation of protein sumoylation
monoacylglycerol biosynthetic process
positive regulation of epithelial to mesenchymal transition involved in endocardial cushion
actin nucleation
regulation of entry of bacterium into host cell
regulation of phosphatidylinositol dephosphorylation
negative regulation of blood vessel endothelial cell proliferation involved in sprouting angi
phosphatidylcholine metabolic process
motor neuron migration
intestinal absorption
positive regulation of epithelial cell differentiation
positive regulation of low-density lipoprotein receptor activity
osteoclast development
fatty acid beta-oxidation using acyl-CoA oxidase
retinal rod cell development
snRNA processing
membranous septum morphogenesis
tetrahydrofolate interconversion
negative regulation of delayed rectifier potassium channel activity
negative regulation of thymocyte apoptotic process
cellular response to epinephrine stimulus
smooth muscle tissue development
positive regulation of follicle-stimulating hormone secretion
negative regulation of chondrocyte proliferation
connective tissue replacement involved in inflammatory response wound healing
positive regulation of Wnt signaling pathway, planar cell polarity pathway
membrane protein proteolysis
neuron death
cardiac muscle cell contraction
positive regulation of wound healing
epaRG_CDM
HepaRG_FBS
HuH7_CDM
HuH7_FBS
PHH
cell_line
HepaRG
HuH7
PHH
medium
CDM
FBS

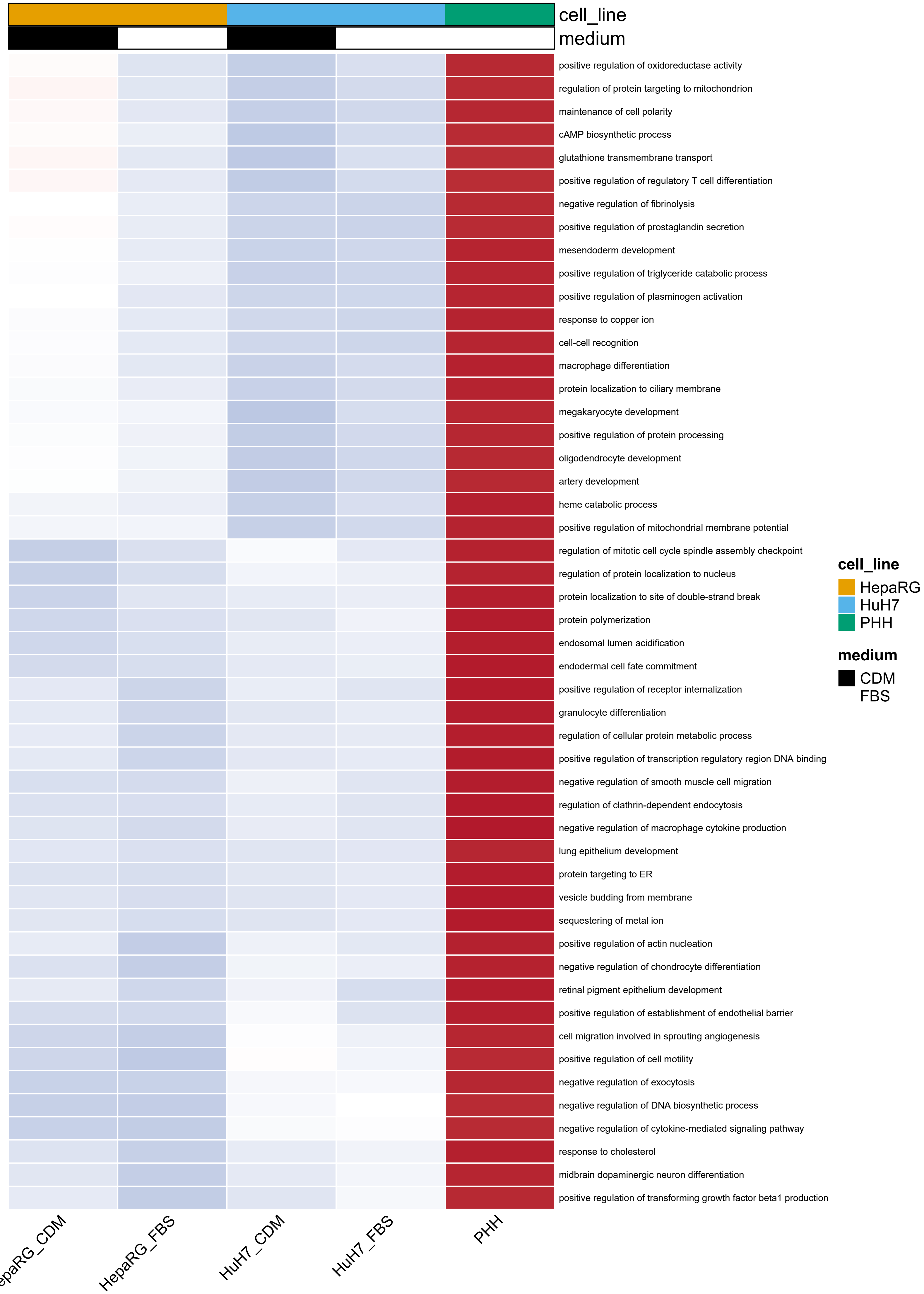
cell_line
medium
positive regulation of oxidoreductase activity
regulation of protein targeting to mitochondrion
maintenance of cell polarity
cAMP biosynthetic process
glutathione transmembrane transport
positive regulation of regulatory T cell differentiation
negative regulation of fibrinolysis
positive regulation of prostaglandin secretion
mesendoderm development
positive regulation of triglyceride catabolic process
positive regulation of plasminogen activation
response to copper ion
cell-cell recognition
macrophage differentiation
protein localization to ciliary membrane
megakaryocyte development
positive regulation of protein processing
oligodendrocyte development
artery development
heme catabolic process
positive regulation of mitochondrial membrane potential
regulation of mitotic cell cycle spindle assembly checkpoint
regulation of protein localization to nucleus
protein localization to site of double-strand break
protein polymerization
endosomal lumen acidification
endodermal cell fate commitment
positive regulation of receptor internalization
granulocyte differentiation
regulation of cellular protein metabolic process
positive regulation of transcription regulatory region DNA binding
negative regulation of smooth muscle cell migration
regulation of clathrin-dependent endocytosis
negative regulation of macrophage cytokine production
lung epithelium development
protein targeting to ER
vesicle budding from membrane
sequestering of metal ion
positive regulation of actin nucleation
negative regulation of chondrocyte differentiation
retinal pigment epithelium development
positive regulation of establishment of endothelial barrier
cell migration involved in sprouting angiogenesis
positive regulation of cell motility
negative regulation of exocytosis
negative regulation of DNA biosynthetic process
negative regulation of cytokine-mediated signaling pathway
response to cholesterol
midbrain dopaminergic neuron differentiation
positive regulation of transforming growth factor beta1 production
HepaRG_FBS
HuH7_CDM
HuH7_FBS
PHH
cell_line
HepaRG
HuH7
PHH
medium
CDM
FBS

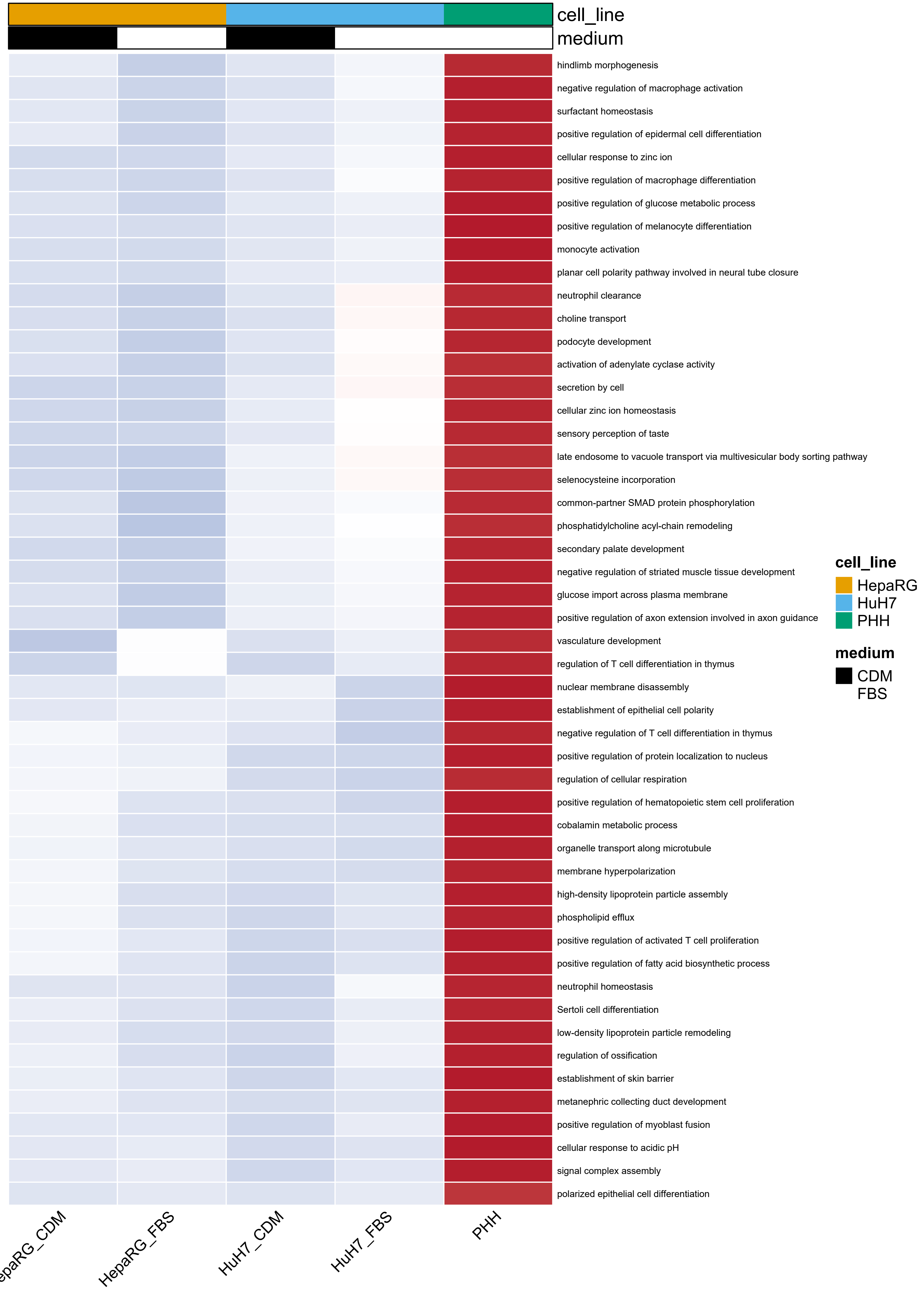
cell_line
medium
hindlimb morphogenesis
negative regulation of macrophage activation
surfactant homeostasis
positive regulation of epidermal cell differentiation
cellular response to zinc ion
positive regulation of macrophage differentiation
positive regulation of glucose metabolic process
positive regulation of melanocyte differentiation
monocyte activation
planar cell polarity pathway involved in neural tube closure
neutrophil clearance
choline transport
podocyte development
activation of adenylate cyclase activity
secretion by cell
cellular zinc ion homeostasis
sensory perception of taste
late endosome to vacuole transport via multivesicular body sorting pathway
selenocysteine incorporation
common-partner SMAD protein phosphorylation
phosphatidylcholine acyl-chain remodeling
secondary palate development
negative regulation of striated muscle tissue development
glucose import across plasma membrane
positive regulation of axon extension involved in axon guidance
vasculature development
regulation of T cell differentiation in thymus
nuclear membrane disassembly
establishment of epithelial cell polarity
negative regulation of T cell differentiation in thymus
positive regulation of protein localization to nucleus
regulation of cellular respiration
positive regulation of hematopoietic stem cell proliferation
cobalamin metabolic process
organelle transport along microtubule
membrane hyperpolarization
high-density lipoprotein particle assembly
phospholipid efflux
positive regulation of activated T cell proliferation
positive regulation of fatty acid biosynthetic process
neutrophil homeostasis
Sertoli cell differentiation
low-density lipoprotein particle remodeling
regulation of ossification
establishment of skin barrier
metanephric collecting duct development
positive regulation of myoblast fusion
cellular response to acidic pH
signal complex assembly
polarized epithelial cell differentiation
cell_line
HepaRG
HuH7
PHH
medium
CDM
FBS
HepaRG_FBS
HuH7_CDM
HuH7_FBS
PHH

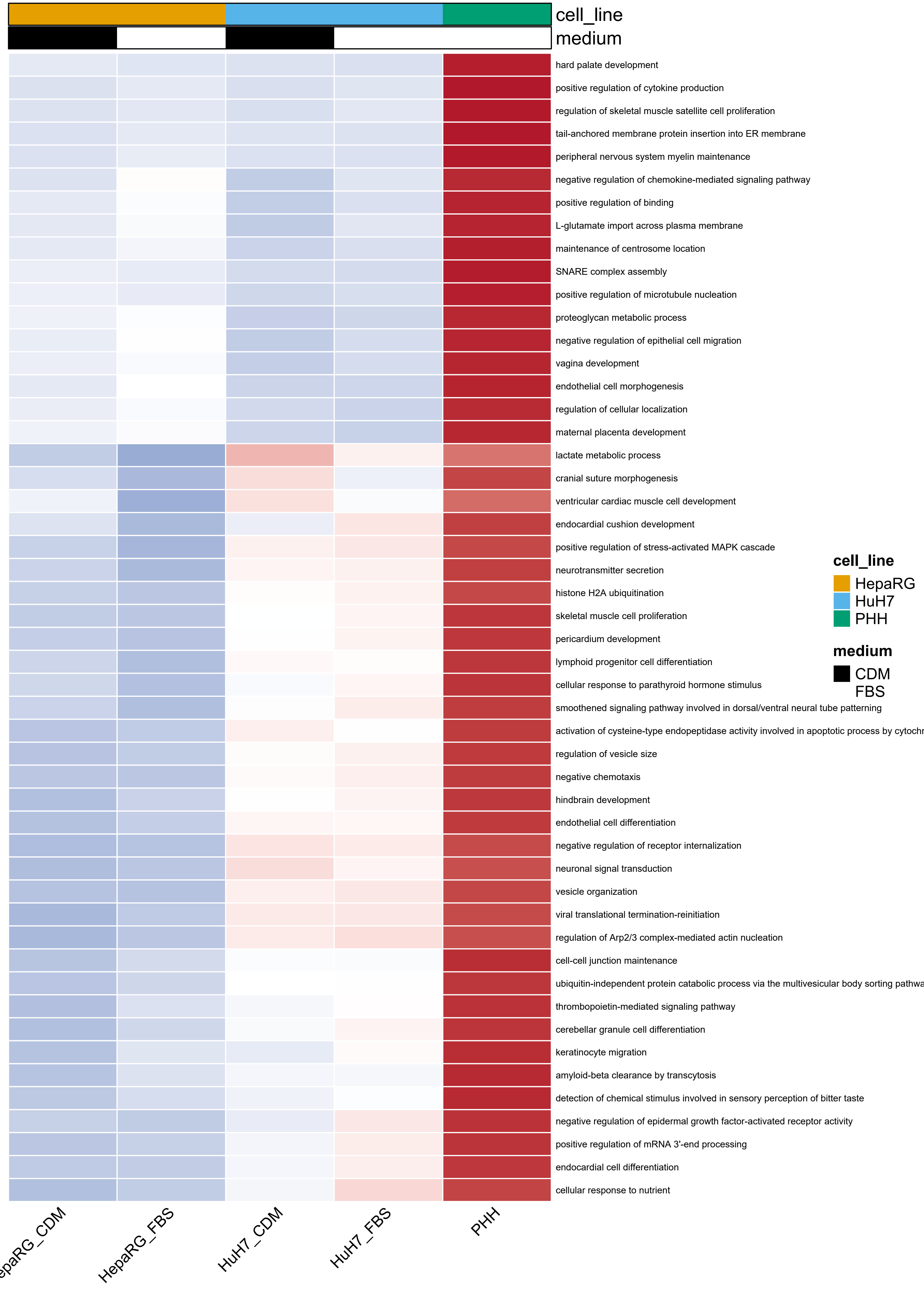
cell_line
medium
hard palate development
positive regulation of cytokine production
regulation of skeletal muscle satellite cell proliferation
tail-anchored membrane protein insertion into ER membrane
peripheral nervous system myelin maintenance
negative regulation of chemokine-mediated signaling pathway
positive regulation of binding
L-glutamate import across plasma membrane
maintenance of centrosome location
SNARE complex assembly
positive regulation of microtubule nucleation
proteoglycan metabolic process
negative regulation of epithelial cell migration
vagina development
endothelial cell morphogenesis
regulation of cellular localization
maternal placenta development
lactate metabolic process
cranial suture morphogenesis
ventricular cardiac muscle cell development
endocardial cushion development
positive regulation of stress-activated MAPK cascade
neurotransmitter secretion
histone H2A ubiquitination
skeletal muscle cell proliferation
pericardium development
lymphoid progenitor cell differentiation
cellular response to parathyroid hormone stimulus
smoothened signaling pathway involved in dorsal/ventral neural tube patterning
activation of cysteine-type endopeptidase activity involved in apoptotic process by cytochr
regulation of vesicle size
negative chemotaxis
hindbrain development
endothelial cell differentiation
negative regulation of receptor internalization
neuronal signal transduction
vesicle organization
viral translational termination-reinitiation
regulation of Arp2/3 complex-mediated actin nucleation
cell-cell junction maintenance
ubiquitin-independent protein catabolic process via the multivesicular body sorting pathwa
thrombopoietin-mediated signaling pathway
cerebellar granule cell differentiation
keratinocyte migration
amyloid-beta clearance by transcytosis
detection of chemical stimulus involved in sensory perception of bitter taste
negative regulation of epidermal growth factor-activated receptor activity
positive regulation of mRNA 3'-end processing
endocardial cell differentiation
cellular response to nutrient
cell_line
HepaRG
HuH7
PHH
medium
CDM
FBS
epaRG_CDM
HepaRG_FBS
HuH7_CDM
HuH7_FBS
PHH

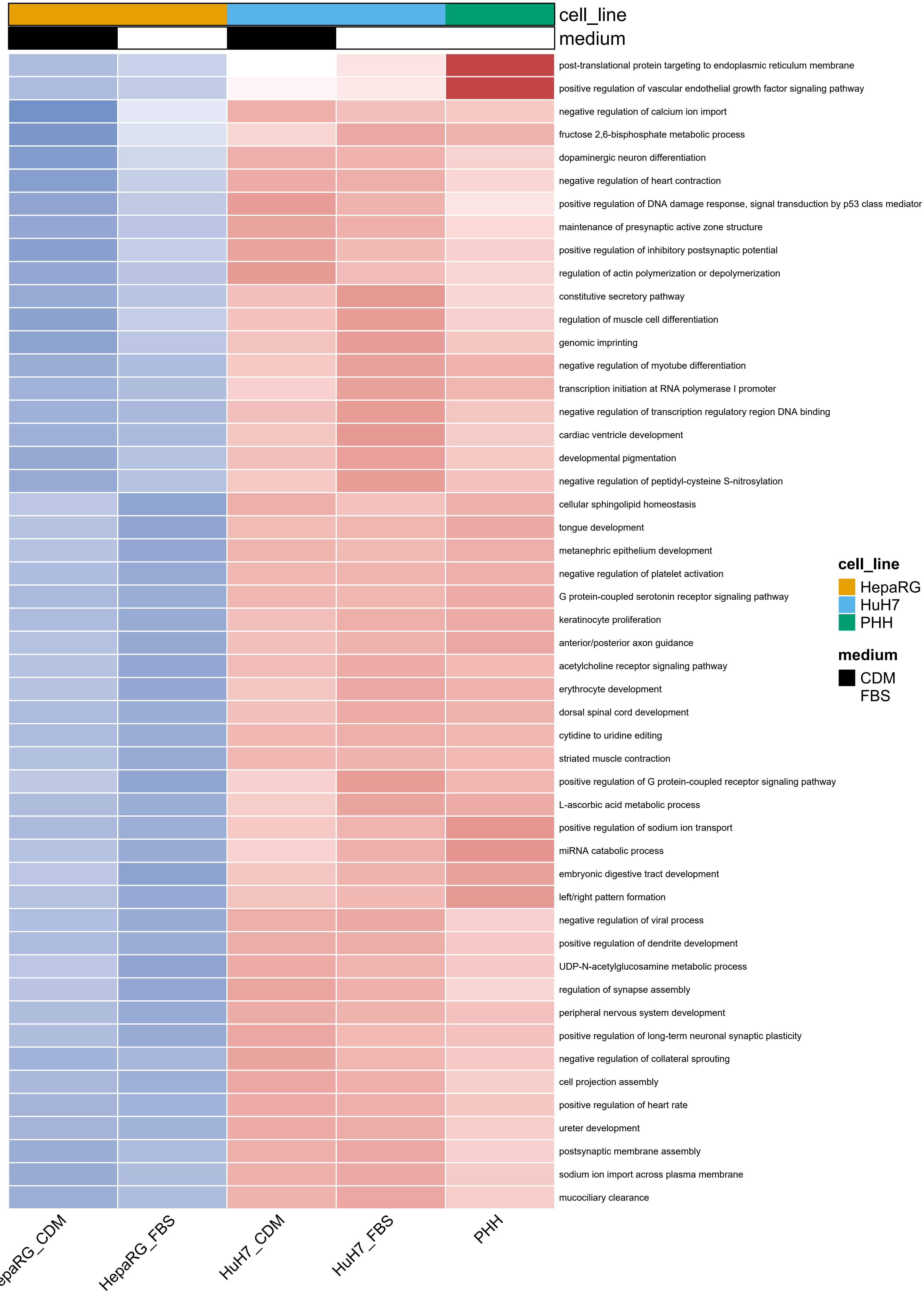

cell_line
medium
post-translational protein targeting to endoplasmic reticulum membrane
positive regulation of vascular endothelial growth factor signaling pathway
negative regulation of calcium ion import
fructose 2,6-bisphosphate metabolic process
dopaminergic neuron differentiation
negative regulation of heart contraction
positive regulation of DNA damage response, signal transduction by p53 class mediator
maintenance of presynaptic active zone structure
positive regulation of inhibitory postsynaptic potential
regulation of actin polymerization or depolymerization
constitutive secretory pathway
regulation of muscle cell differentiation
genomic imprinting
negative regulation of myotube differentiation
transcription initiation at RNA polymerase I promoter
negative regulation of transcription regulatory region DNA binding
cardiac ventricle development
developmental pigmentation
negative regulation of peptidyl-cysteine S-nitrosylation
cellular sphingolipid homeostasis
tongue development
metanephric epithelium development
negative regulation of platelet activation
G protein-coupled serotonin receptor signaling pathway
keratinocyte proliferation
anterior/posterior axon guidance
acetylcholine receptor signaling pathway
erythrocyte development
dorsal spinal cord development
cytidine to uridine editing
striated muscle contraction
positive regulation of G protein-coupled receptor signaling pathway
L-ascorbic acid metabolic process
positive regulation of sodium ion transport
miRNA catabolic process
embryonic digestive tract development
left/right pattern formation
negative regulation of viral process
positive regulation of dendrite development
UDP-N-acetylglucosamine metabolic process
regulation of synapse assembly
peripheral nervous system development
positive regulation of long-term neuronal synaptic plasticity
negative regulation of collateral sprouting
cell projection assembly
positive regulation of heart rate
ureter development
postsynaptic membrane assembly
sodium ion import across plasma membrane
mucociliary clearance
epaRG_CDM
HepaRG_FBS
HuH7_CDM
HuH7_FBS
PHH
cell_line
HepaRG
HuH7
PHH
medium
CDM
FBS

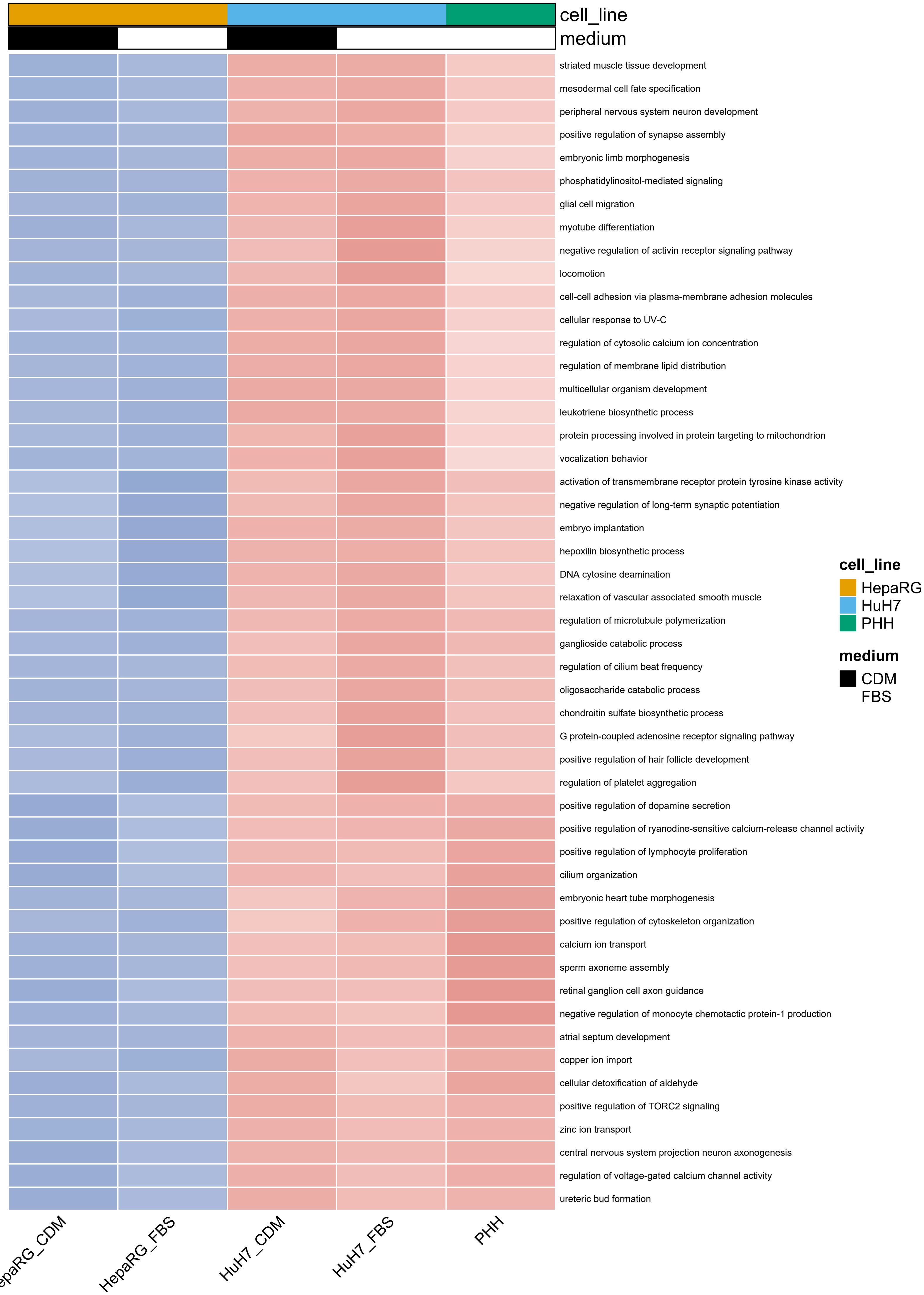

cell_line
medium
striated muscle tissue development
mesodermal cell fate specification
peripheral nervous system neuron development
positive regulation of synapse assembly
embryonic limb morphogenesis
phosphatidylinositol-mediated signaling
glial cell migration
myotube differentiation
negative regulation of activin receptor signaling pathway
locomotion
cell-cell adhesion via plasma-membrane adhesion molecules
cellular response to UV-C
regulation of cytosolic calcium ion concentration
regulation of membrane lipid distribution
multicellular organism development
leukotriene biosynthetic process
protein processing involved in protein targeting to mitochondrion
vocalization behavior
activation of transmembrane receptor protein tyrosine kinase activity
negative regulation of long-term synaptic potentiation
embryo implantation
hepoxilin biosynthetic process
DNA cytosine deamination
relaxation of vascular associated smooth muscle
regulation of microtubule polymerization
ganglioside catabolic process
regulation of cilium beat frequency
oligosaccharide catabolic process
chondroitin sulfate biosynthetic process
G protein-coupled adenosine receptor signaling pathway
positive regulation of hair follicle development
regulation of platelet aggregation
positive regulation of dopamine secretion
positive regulation of ryanodine-sensitive calcium-release channel activity
positive regulation of lymphocyte proliferation
cilium organization
embryonic heart tube morphogenesis
positive regulation of cytoskeleton organization
calcium ion transport
sperm axoneme assembly
retinal ganglion cell axon guidance
negative regulation of monocyte chemotactic protein-1 production
atrial septum development
copper ion import
cellular detoxification of aldehyde
positive regulation of TORC2 signaling
zinc ion transport
central nervous system projection neuron axonogenesis
regulation of voltage-gated calcium channel activity
ureteric bud formation
cell_line
HepaRG
HuH7
PHH
medium
CDM
FBS
HepaRG_FBS
HuH7_CDM
HuH7_FBS
PHH

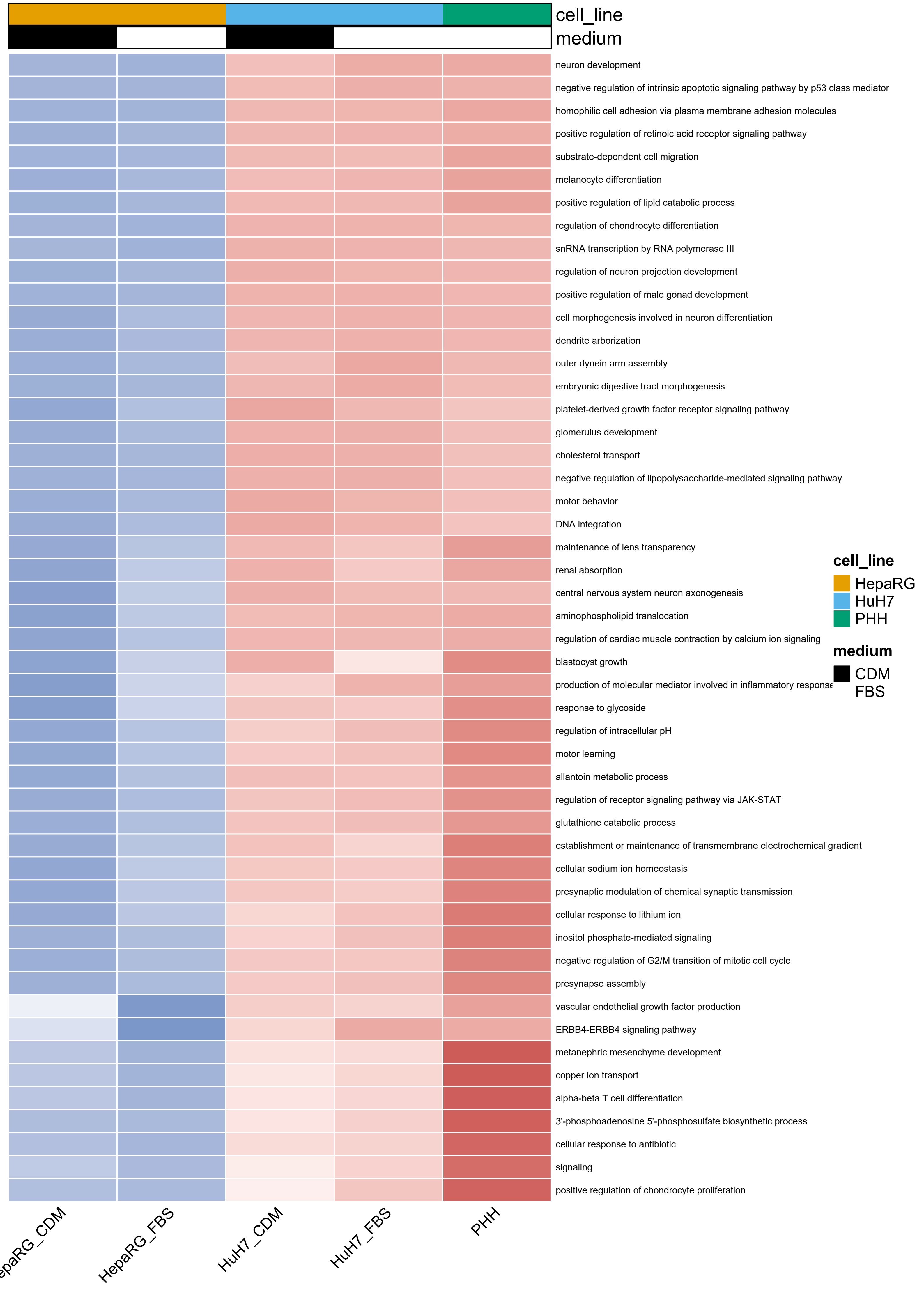
cell_line
medium
neuron development
negative regulation of intrinsic apoptotic signaling pathway by p53 class mediator
homophilic cell adhesion via plasma membrane adhesion molecules
positive regulation of retinoic acid receptor signaling pathway
substrate-dependent cell migration
melanocyte differentiation
positive regulation of lipid catabolic process
regulation of chondrocyte differentiation
snRNA transcription by RNA polymerase III
regulation of neuron projection development
positive regulation of male gonad development
cell morphogenesis involved in neuron differentiation
dendrite arborization
outer dynein arm assembly
embryonic digestive tract morphogenesis
platelet-derived growth factor receptor signaling pathway
glomerulus development
cholesterol transport
negative regulation of lipopolysaccharide-mediated signaling pathway
motor behavior
DNA integration
maintenance of lens transparency
renal absorption
central nervous system neuron axonogenesis
aminophospholipid translocation
regulation of cardiac muscle contraction by calcium ion signaling
blastocyst growth
production of molecular mediator involved in inflammatory response
response to glycoside
regulation of intracellular pH
motor learning
allantoin metabolic process
regulation of receptor signaling pathway via JAK-STAT
glutathione catabolic process
establishment or maintenance of transmembrane electrochemical gradient
cellular sodium ion homeostasis
presynaptic modulation of chemical synaptic transmission
cellular response to lithium ion
inositol phosphate-mediated signaling
negative regulation of G2/M transition of mitotic cell cycle
presynapse assembly
vascular endothelial growth factor production
ERBB4-ERBB4 signaling pathway
metanephric mesenchyme development
copper ion transport
alpha-beta T cell differentiation
3'-phosphoadenosine 5'-phosphosulfate biosynthetic process
cellular response to antibiotic
signaling
positive regulation of chondrocyte proliferation
HepaRG_FBS
HuH7_CDM
HuH7_FBS
PHH
cell_line
HepaRG
HuH7
PHH
medium
CDM
FBS

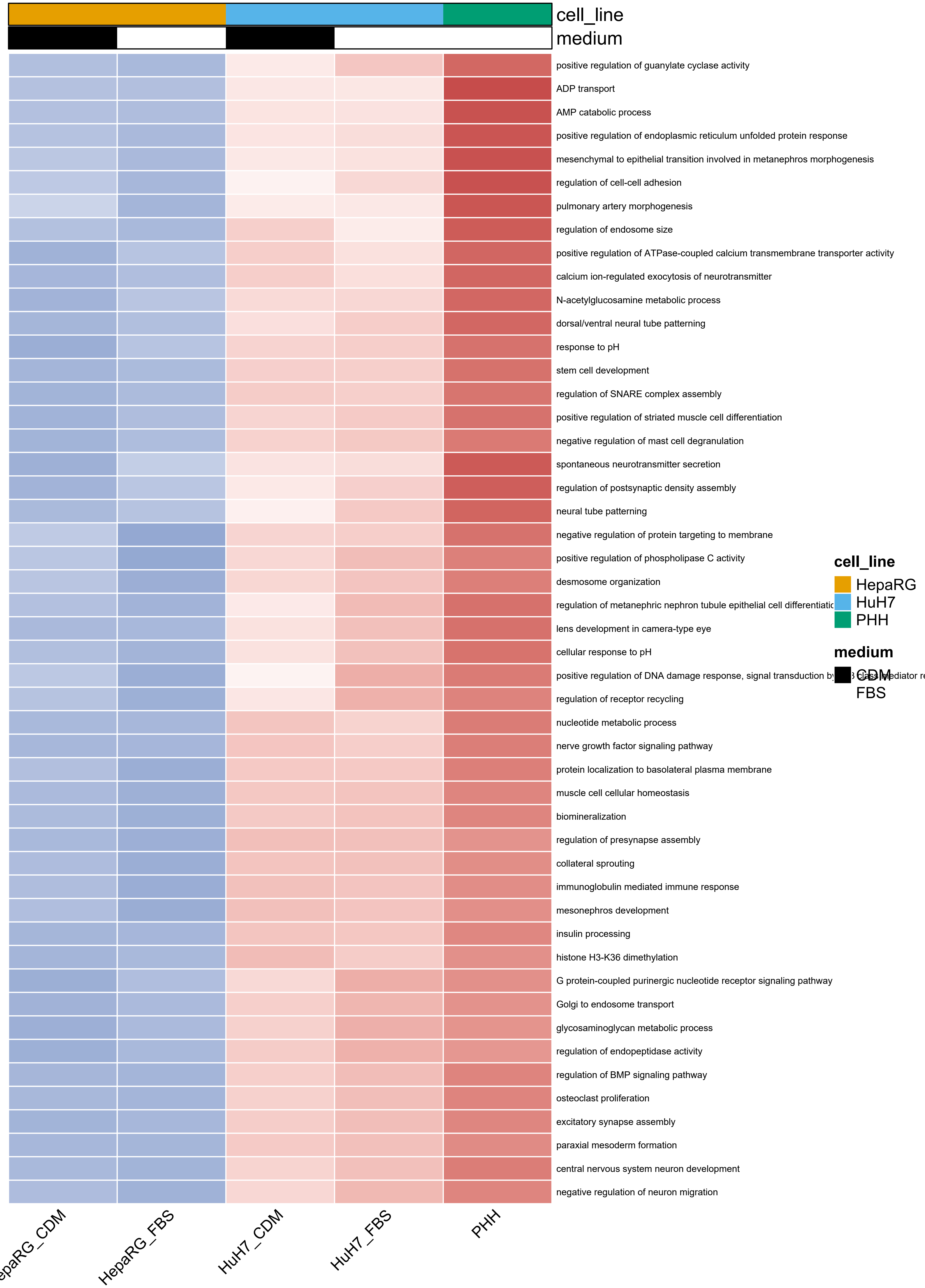
cell_line
medium
positive regulation of guanylate cyclase activity
ADP transport
AMP catabolic process
positive regulation of endoplasmic reticulum unfolded protein response
mesenchymal to epithelial transition involved in metanephros morphogenesis
regulation of cell-cell adhesion
pulmonary artery morphogenesis
regulation of endosome size
positive regulation of ATPase-coupled calcium transmembrane transporter activity
calcium ion-regulated exocytosis of neurotransmitter
N-acetylglucosamine metabolic process
dorsal/ventral neural tube patterning
response to pH
stem cell development
regulation of SNARE complex assembly
positive regulation of striated muscle cell differentiation
negative regulation of mast cell degranulation
spontaneous neurotransmitter secretion
regulation of postsynaptic density assembly
neural tube patterning
negative regulation of protein targeting to membrane
positive regulation of phospholipase C activity
desmosome organization
regulation of metanephric nephron tubule epithelial cell differentiatio
lens development in camera-type eye
cellular response to pH
positive regulation of DNA damage response, signal transduction by
regulation of receptor recycling
nucleotide metabolic process
nerve growth factor signaling pathway
protein localization to basolateral plasma membrane
muscle cell cellular homeostasis
biomineralization
regulation of presynapse assembly
collateral sprouting
immunoglobulin mediated immune response
mesonephros development
insulin processing
histone H3-K36 dimethylation
G protein-coupled purinergic nucleotide receptor signaling pathway
Golgi to endosome transport
glycosaminoglycan metabolic process
regulation of endopeptidase activity
regulation of BMP signaling pathway
osteoclast proliferation
excitatory synapse assembly
paraxial mesoderm formation
central nervous system neuron development
negative regulation of neuron migration
cell_line
HepaRG
HuH7
PHH
medium
CDM
FBS
HepaRG_FBS
HuH7_CDM
HuH7_FBS
PHH
