## Supplemental file 1 for "Cell line identity rather than medium composition determines transcriptomic profiles of HepaRG and HuH7 cells cultured in chemically defined or serum-based media: comparison with primary human hepatocytes"

Department of Applied Biochemistry, TIB 4/3-2

Gustav-Meyer-Allee 25, 13355 Berlin, Germany

**SM1. HeLa cell culture and RNA isolation**

HeLa cells were cultured under two parallel conditions (serum-supplemented medium versus chemically defined medium, CDM) prior to RNA isolation. For serum-supplemented culture, cells were thawed from a cryopreserved FBS-containing stock and maintained in DMEM (low glucose) (L0064, Biowest) supplemented with FBS (10%; S-10-L, c.c.pro GmbH), L-glutamine (2 mM; X0550, Biowest), and non-essential amino acids (NEAA; 1%; X0557, Biowest). For CDM culture, cells were cultured in CDM developed by Nessar et al. based on DMEM/F12 (L0090, Biowest) supplemented with NEAA (1%; X0557–100, Biowest), HEPES (15 mM; L0180, Biowest), D-glucose (0.1% w/v; G8769, Sigma-Aldrich), L-glutamine (2 mM; X0550, Biowest), insulin–transferrin–selenium (ITS; 1×; 41400045, Gibco), recombinant human epidermal growth factor (hEGF; 100 ng/mL; PHG0313, Sigma-Aldrich), and hydrocortisone (10 µg/mL; sc-250130, Santa Cruz Biotechnology) [1]. Cells were maintained for five consecutive passages in triplicate in T25 flasks (TPP) under standard culture conditions (37 °C, humidified atmosphere, 5% $CO_2$).

Total RNA was isolated using the same workflow described for HepaRG and HuH7 cells. Briefly, at ~90% confluence, monolayers were washed with phosphate-buffered saline (PBS) to remove residual medium and lysed directly in RLT buffer supplemented with β-mercaptoethanol (1% v/v), using 700 µL lysis buffer per T25 flask. RNA was purified using the QIAwave RNA Kit (50) (74534, Qiagen) according to the manufacturer's instructions, and RNA concentration and purity were assessed using a NanoDrop 2000 UV-Vis spectrophotometer (Thermo Scientific).

**Table S1**. ***Table_S1_detox_panels_similarity_and_matrices.xlsx***. *Detox panel benchmarking. Detox gene-panel expression matrices and PHH similarity scores (Phase I/II and transporters).*

**Table S2**. ***Table_S2_identity_panels_and_PHH_similarity.xlsx***. *Identity panel benchmarking. PHH-referenced similarity scores for hepatic identity gene panels across HepaRG and HuH7 in FBS-SM vs CDM.*

***Table S3****. Sequencing yield and primary read quality metrics per sample.*

| ID | Sample name | Total raw reads | Total non-rRNA reads | Total HQ reads | HQ bases (Q30) | GC content | rRNA reads (%) | HQ reads (%) |
|---|---|---|---|---|---|---|---|---|
| 1 | HuH7_CDM_1 | 66 M | 64.67 M | 63.96 M | 92.53% | 48.87% | 2.02% | 96.91% |
| 2 | HuH7_CDM_2 | 66 M | 65.65 M | 64.85 M | 96.10% | 49.04% | 0.54% | 98.25% |
| 3 | HuH7_CDM_3 | 66 M | 65.58 M | 64.84 M | 92.90% | 48.89% | 0.63% | 98.24% |
| 4 | HuH7_FBS-SM_1 | 66 M | 65.56 M | 64.81 M | 92.71% | 49.02% | 0.66% | 98.19% |
| 5 | HuH7_FBS-SM_2 | 66 M | 65.59 M | 64.77 M | 92.82% | 49.08% | 0.62% | 98.14% |
| 6 | HuH7_FBS-SM_3 | 66 M | 65.44 M | 64.70 M | 92.61% | 49.12% | 0.84% | 98.02% |
| 7 | PHH_1 | 66 M | 64.09 M | 63.34 M | 96.50% | 49.00% | 2.89% | 95.97% |
| 8 | PHH_2 | 66 M | 63.79 M | 63.05 M | 96.93% | 49.52% | 3.35% | 95.53% |
| 9 | HepaRG_CDM_1 | 66 M | 65.24 M | 64.54 M | 92.94% | 48.68% | 1.14% | 97.79% |
| 10 | HepaRG_CDM_2 | 66 M | 65.16 M | 64.41 M | 92.91% | 48.67% | 1.27% | 97.59% |
| 11 | HepaRG_CDM_3 | 66 M | 65.25 M | 64.54 M | 92.94% | 48.73% | 1.14% | 97.79% |
| 12 | HepaRG_FBS-SM_1 | 66 M | 65.04 M | 64.29 M | 92.24% | 49.88% | 1.46% | 97.40% |
| 13 | HepaRG_FBS-SM_2 | 66 M | 65.25 M | 64.47 M | 95.63% | 48.72% | 1.14% | 97.68% |
| 14 | HepaRG_FBS-SM_3 | 66 M | 65.25 M | 64.52 M | 92.79% | 48.60% | 1.14% | 97.76% |
| 15 | Hela_CDM_1 | 76 M | 75.55 M | 73.68 M | 95.92% | 49.73% | 0.59% | 96.95% |
| 16 | Hela_CDM_2 | 76 M | 75.54 M | 73.26 M | 95.93% | 49.79% | 0.61% | 96.40% |
| 17 | Hela_CDM_3 | 68.94 M | 68.76 M | 67.14 M | 97.09% | 49.70% | 0.26% | 97.40% |
| 18 | Hela_FBS-SM_1 | 75.46 M | 74.92 M | 73.01 M | 96.01% | 49.58% | 0.73% | 96.75% |
| 19 | Hela_FBS-SM_2 | 76 M | 75.56 M | 73.51 M | 96.01% | 49.75% | 0.58% | 96.72% |
| 20 | Hela_FBS-SM_3 | 74.55 M | 74.20 M | 72.82 M | 95.88% | 49.73% | 0.47% | 97.67% |

***Table S4****. Alignment and mapping metrics per sample.*

| ID | Sample | Total HQ reads | Mapped reads | Unmapped reads | Unique reads |
|---|---|---|---|---|---|
| 1 | HuH7_CDM_1 | 63.96 M | 63.43 M (99.17%) | 532.99 K (0.83%) | 62.33 M (97.45%) |
| 2 | HuH7_CDM_2 | 64.85 M | 64.23 M (99.05%) | 617.23 K (0.95%) | 63.07 M (97.26%) |
| 3 | HuH7_CDM_3 | 64.84 M | 64.34 M (99.24%) | 494.14 K (0.76%) | 63.12 M (97.36%) |
| 4 | HuH7_FBS-SM_1 | 64.81 M | 64.31 M (99.24%) | 494.68 K (0.76%) | 63.19 M (97.50%) |
| 5 | HuH7_FBS-SM_2 | 64.77 M | 64.27 M (99.22%) | 502.99 K (0.78%) | 63.06 M (97.36%) |
| 6 | HuH7_FBS-SM_3 | 64.70 M | 64.20 M (99.23%) | 498.86 K (0.77%) | 63.08 M (97.50%) |
| 7 | PHH_1 | 63.34 M | 61.86 M (97.67%) | 1.48 M (2.33%) | 58.94 M (93.06%) |
| 8 | PHH_2 | 63.05 M | 61.28 M (97.19%) | 1.77 M (2.81%) | 58.53 M (92.83%) |
| 9 | HepaRG_CDM_1 | 64.54 M | 64.00 M (99.16%) | 541.54 K (0.84%) | 62.52 M (96.87%) |
| 10 | HepaRG_CDM_2 | 64.41 M | 63.85 M (99.12%) | 565.48 K (0.88%) | 62.37 M (96.84%) |
| 11 | HepaRG_CDM_3 | 64.54 M | 63.99 M (99.15%) | 549.70 K (0.85%) | 62.46 M (96.78%) |
| 12 | HepaRG_FBS-SM_1 | 64.29 M | 63.71 M (99.10%) | 577.76 K (0.90%) | 62.32 M (96.94%) |
| 13 | HepaRG_FBS-SM_2 | 64.47 M | 63.83 M (99.02%) | 634.43 K (0.98%) | 62.34 M (96.70%) |
| 14 | HepaRG_FBS-SM_3 | 64.52 M | 63.95 M (99.11%) | 577.35 K (0.89%) | 62.52 M (96.90%) |
| 15 | Hela_CDM_1 | 73.68 M | 72.88 M (98.91%) | 803.44 K (1.09%) | 71.43 M (96.94%) |
| 16 | Hela_CDM_2 | 73.26 M | 72.41 M (98.84%) | 850.05 K (1.16%) | 71.02 M (96.94%) |
| 17 | Hela_CDM_3 | 67.14 M | 66.56 M (99.13%) | 586.98 K (0.87%) | 65.18 M (97.08%) |
| 18 | Hela_FBS-SM_1 | 73.01 M | 72.11 M (98.76%) | 906.93 K (1.24%) | 70.43 M (96.46%) |
| 19 | Hela_ FBS-SM_2 | 73.51 M | 72.65 M (98.84%) | 854.40 K (1.16%) | 70.89 M (96.44%) |
| 20 | Hela_FBS-SM_3 | 72.82 M | 71.44 M (98.11%) | 1.37 M (1.89%) | 69.83 M (95.90%) |

***Table S5****. Distribution of aligned reads across genomic features per sample.*

| ID | Sample name | exonic | intronic | intergenic | overlapping exon |
|---|---|---|---|---|---|
| 1 | HuH7_CDM_1 | 53,821,399 (90.17%) | 4,952,447 (8.3%) | 912,317 (1.53%) | 4,240,933 (7.11%) |
| 2 | HuH7_CDM_2 | 54,641,477 (90.48%) | 4,828,455 (8%) | 921,793 (1.53%) | 4,151,775 (6.87%) |
| 3 | HuH7_CDM_3 | 54,554,753 (90.22%) | 4,960,055 (8.2%) | 951,886 (1.57%) | 4,151,589 (6.87%) |
| 4 | HuH7_FBS-SM_1 | 54,706,606 (90.27%) | 5,015,947 (8.28%) | 879,585 (1.45%) | 4,293,205 (7.08%) |
| 5 | HuH7_FBS-SM_2 | 54,499,133 (90.28%) | 4,951,014 (8.2%) | 913,982 (1.51%) | 4,163,444 (6.9%) |
| 6 | HuH7_FBS-SM_3 | 54,490,965 (90.13%) | 5,066,622 (8.38%) | 902,161 (1.49%) | 4,291,037 (7.1%) |
| 7 | PHH_1 | 39,789,644 (87.43%) | 5,198,774 (11.42%) | 520,964 (1.14%) | 3,172,283 (6.97%) |
| 8 | PHH_2 | 40,687,603 (87.57%) | 5,255,539 (11.31%) | 518,573 (1.12%) | 3,261,425 (7.02%) |
| 9 | HepaRG_CDM_1 | 51,475,677 (90.19%) | 4,564,956 (8%) | 1,037,088 (1.82%) | 3,892,144 (6.82%) |
| 10 | HepaRG_CDM_2 | 51,236,611 (89.63%) | 4,824,769 (8.44%) | 1,101,735 (1.93%) | 3,906,622 (6.83%) |
| 11 | HepaRG_CDM_3 | 50,992,066 (89.88%) | 4,686,019 (8.26%) | 1,054,958 (1.86%) | 3,915,508 (6.9%) |
| 12 | HepaRG_FBS-SM_1 | 53,662,965 (90.39%) | 4,816,590 (8.11%) | 890,506 (1.5%) | 4,274,275 (7.2%) |
| 13 | HepaRG_FBS-SM_2 | 52,085,987 (90.04%) | 4,710,439 (8.14%) | 1,053,212 (1.82%) | 3,857,095 (6.67%) |
| 14 | HepaRG_FBS-SM_3 | 53,223,475 (90.42%) | 4,613,366 (7.84%) | 1,028,102 (1.75%) | 3,991,822 (6.78%) |
| 15 | Hela_CDM_1 | 60,795,068 (89.34%) | 6,526,743 (9.59%) | 724,360 (1.06%) | 4,782,472 (7.03%) |
| 16 | Hela_CDM_2 | 61,047,378 (90.25%) | 5,912,007 (8.74%) | 684,496 (1.01%) | 4,773,254 (7.06%) |
| 17 | Hela_CDM_3 | 55,814,633 (90.05%) | 5,531,112 (8.92%) | 637,680 (1.03%) | 4,386,690 (7.08%) |
| 18 | Hela_FBS-SM_1 | 61,881,496 (92.37%) | 4,413,817 (6.59%) | 700,239 (1.05%) | 4,524,895 (6.75%) |
| 19 | Hela_FBS-SM_2 | 62,105,513 (92.38%) | 4,411,033 (6.56%) | 712,770 (1.06%) | 4,495,801 (6.69%) |
| 20 | Hela_FBS-SM_3 | 60,203,596 (91.73%) | 4,695,051 (7.15%) | 734,978 (1.12%) | 4,410,376 (6.72%) |

**Table S6**. ***Table_S6_DESeq2_differential_expression_outputs.xlsx****. DESeq2 outputs. Full gene-level DESeq2 results.*

**Table S7**. ***Table_S7_QuickGO_ssGSEA_outputs.xlsx****. QuickGO ssGSEA scores. Per-sample ssGSEA matrices (GSVA/ssGSEA on DESeq2 VST) and condition summaries for the GO:BP term sets.*

**Table S8**. ***Table_S8_GO_BP_enrichment_outputs.xlsx****. GO:BP enrichment results. Complete clusterProfiler GO:BP over-representation outputs.*

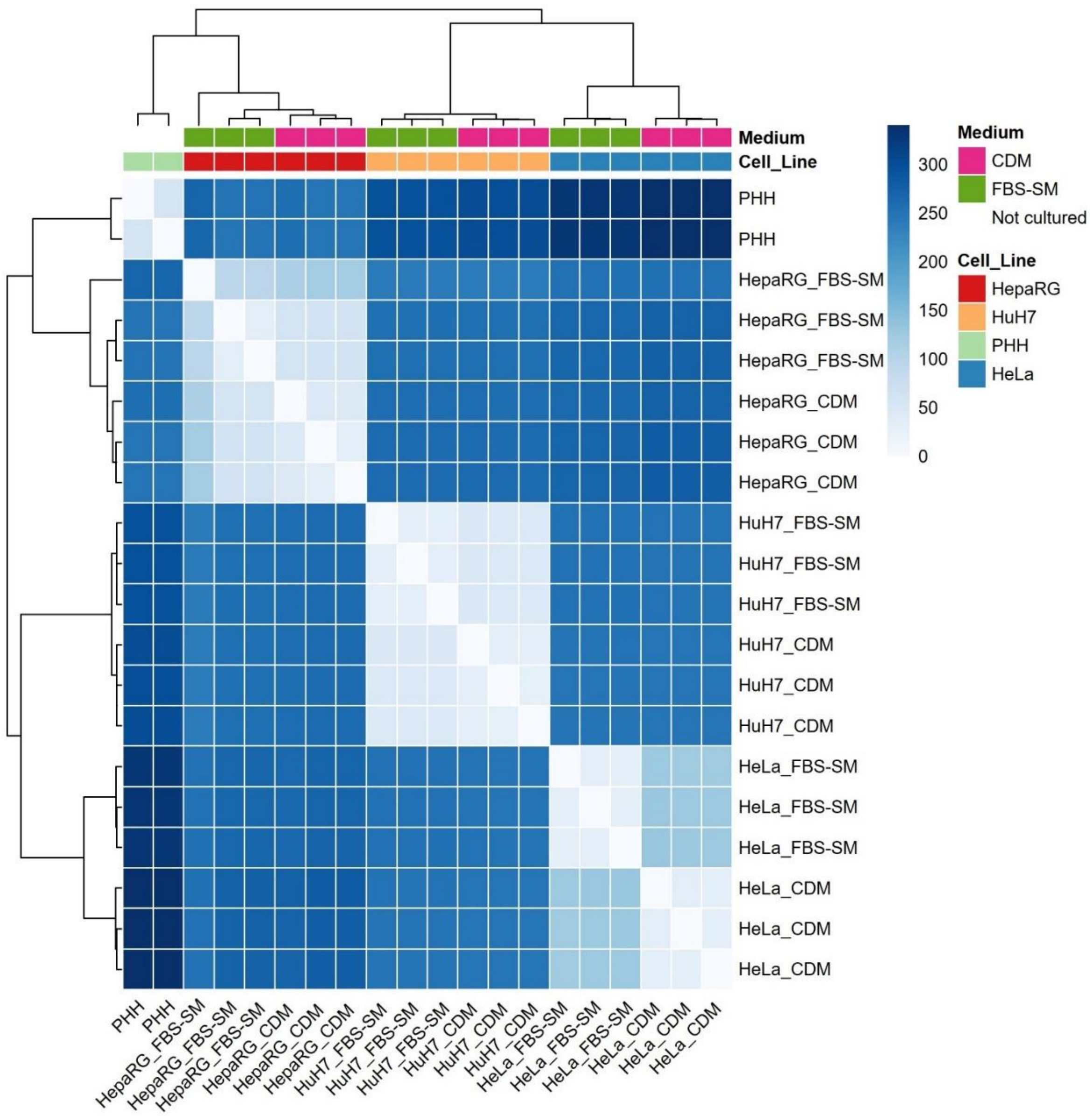


***Figure S1**. Sample-level transcriptomic distance structure. Sample-by-sample Euclidean distance heatmap computed from DESeq2 VST expression values, with annotation tracks indicating cell model and medium (CDM, FBS-SM, or PHH reference).*

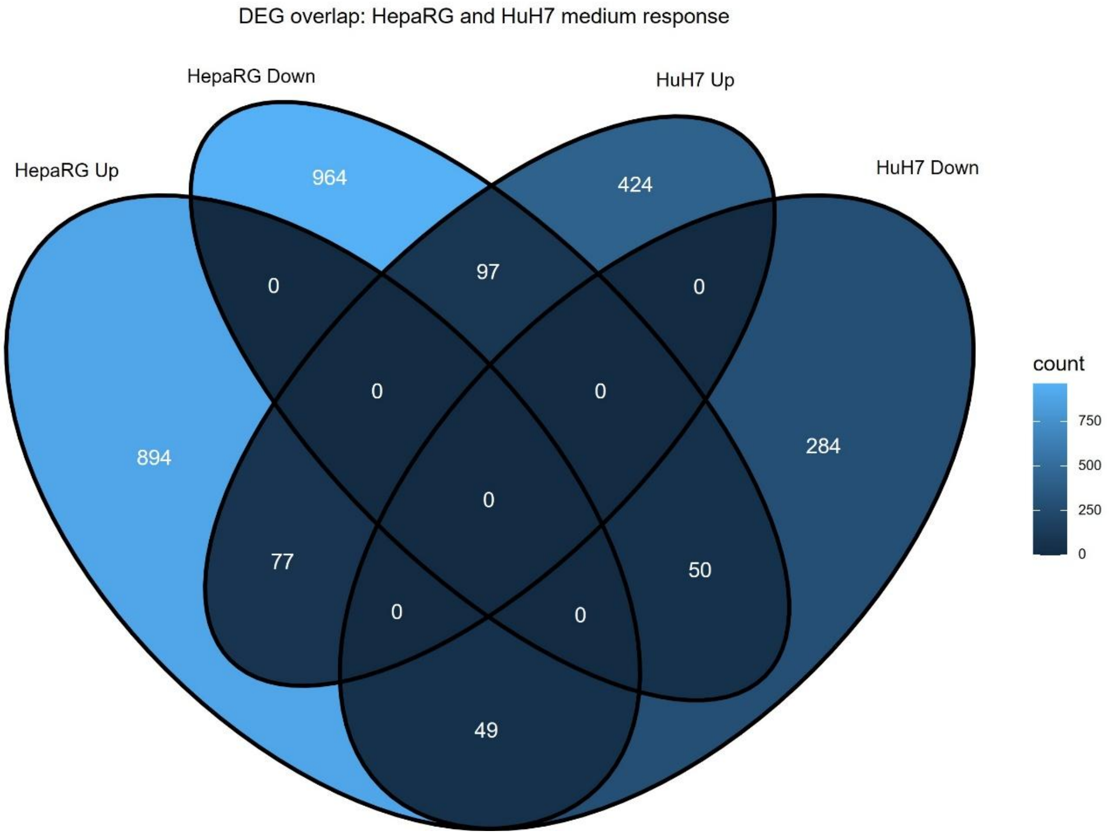


**Figure S2**. Venn diagram of direction-resolved DEG overlaps between HepaRG and HuH7 under CDM versus FBS-SM. The four sets represent genes upregulated or downregulated in CDM relative to FBS-SM in HepaRG and HuH7 cells, using DESeq2 significance criteria of FDR < 0.05 and |log2FC| > 0.5.

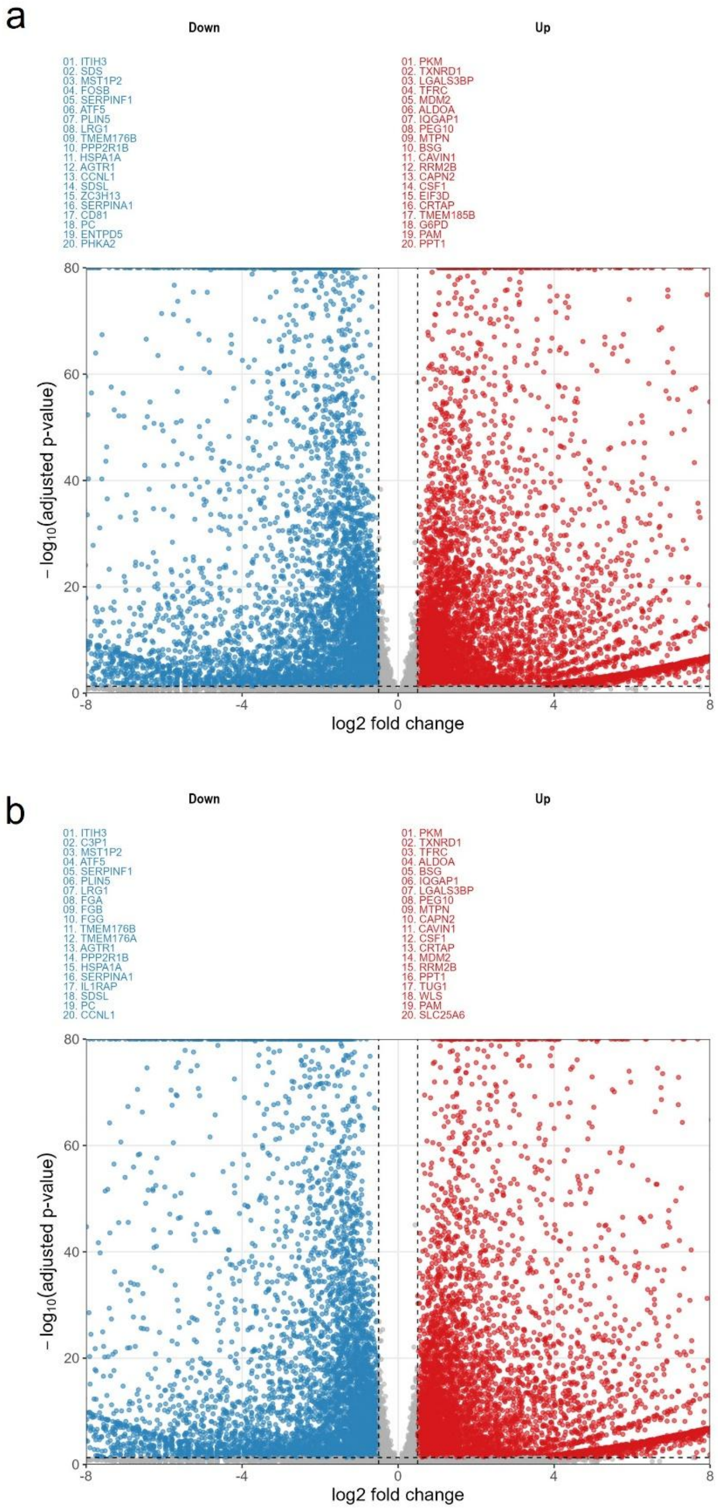


**Figure S3**. HepaRG benchmark differential expression relative to PHH. Full-size volcano plots showing differential expression of HepaRG cultured in (a) CDM and (b) FBS-SM relative to pooled PHH. Axes and thresholds match the main DESeq2 criteria (FDR < 0.05 and |log2FC| > 0.5), with representative high-effect and/or highly significant genes annotated.

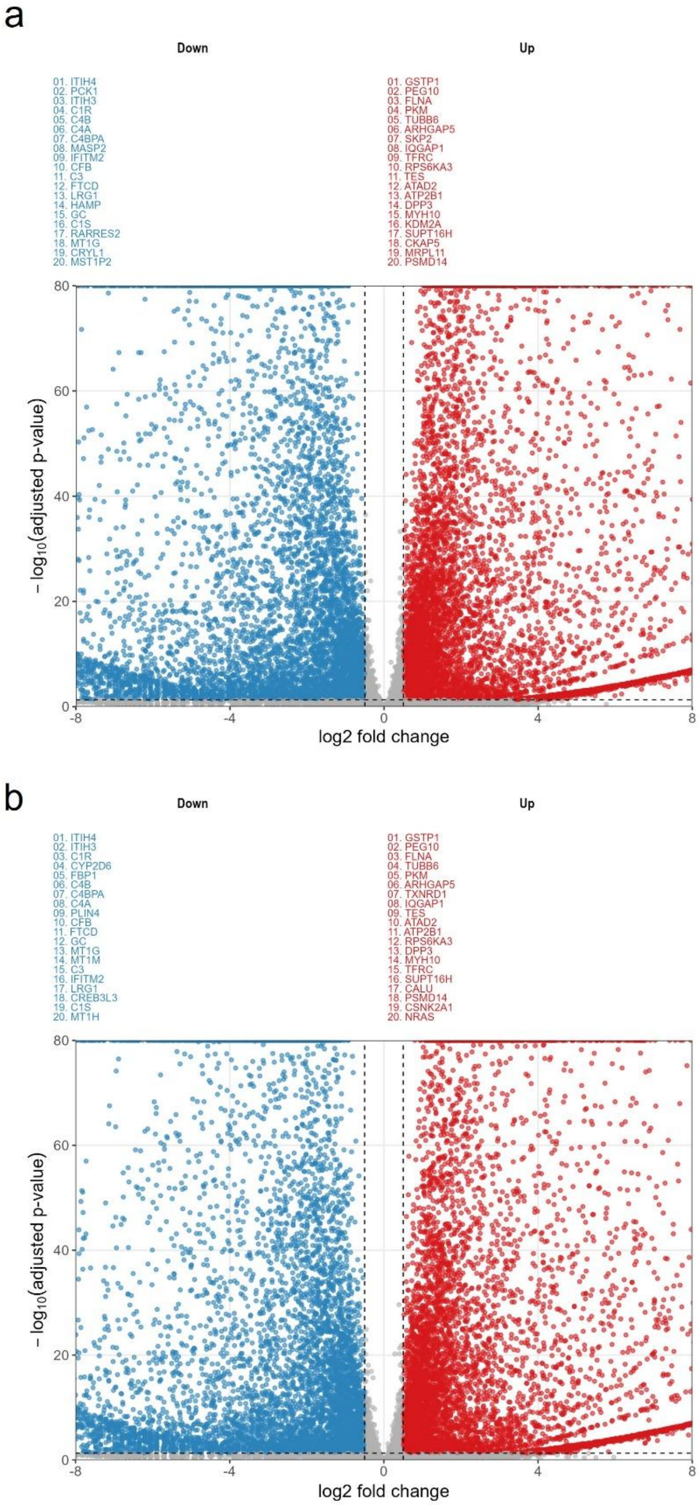


***Figure S4**. HuH7 benchmark differential expression relative to PHH. Full-size volcano plots showing differential expression of HuH7 cultured in (a) CDM and (b) FBS-SM relative to pooled PHH.* Axes and thresholds match the main DESeq2 criteria (FDR < 0.05 and |log2FC| > 0.5), with representative high-effect and/or highly significant genes annotated.

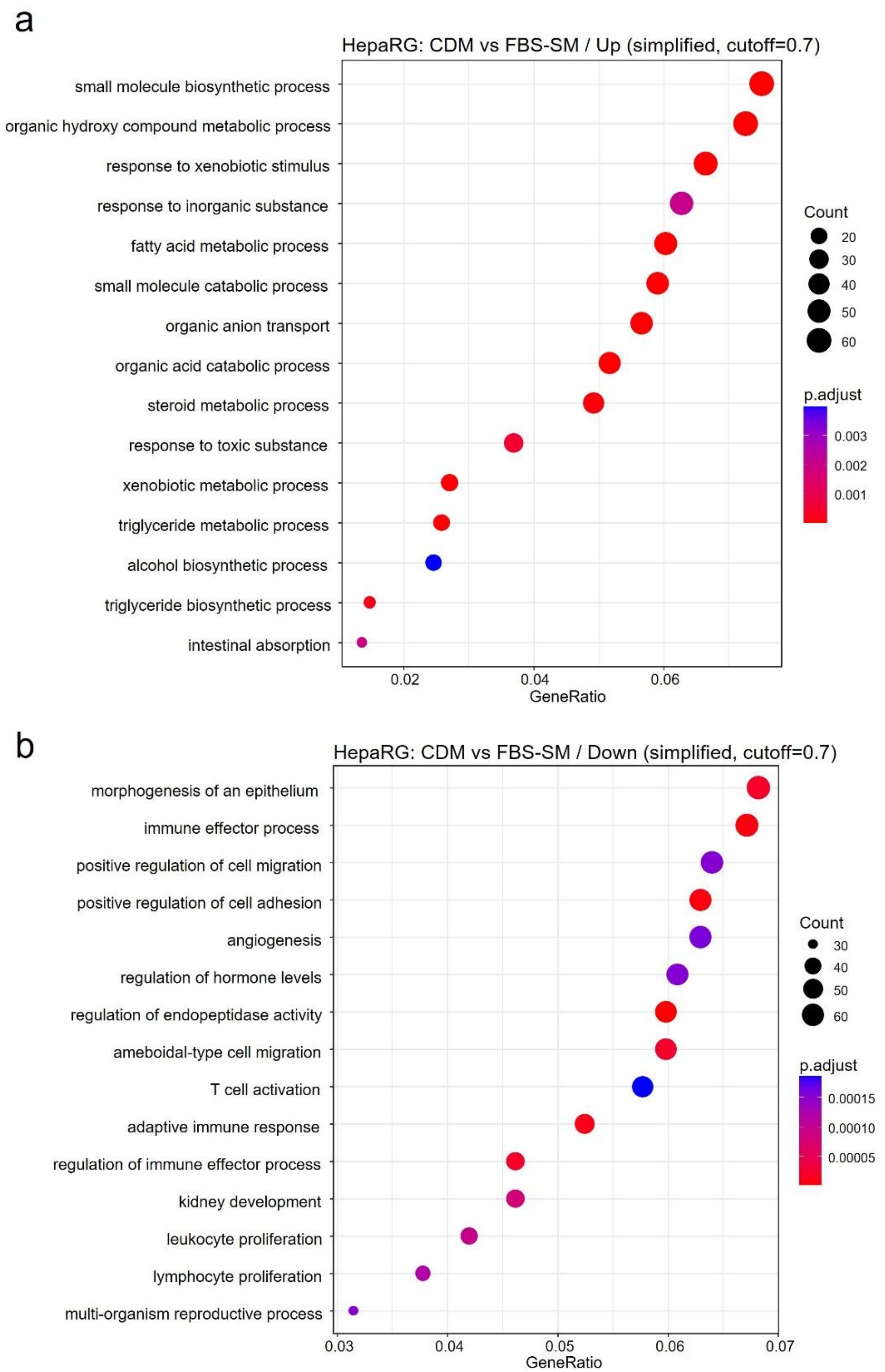


**Figure S5**. Direction-resolved GO:BP enrichment for HepaRG CDM versus FBS-SM. Simplified GO:BP enrichment dotplots for HepaRG genes (a) upregulated and (b) downregulated in CDM versus FBS-SM. Dot size denotes GeneRatio and dot color denotes adjusted P value.

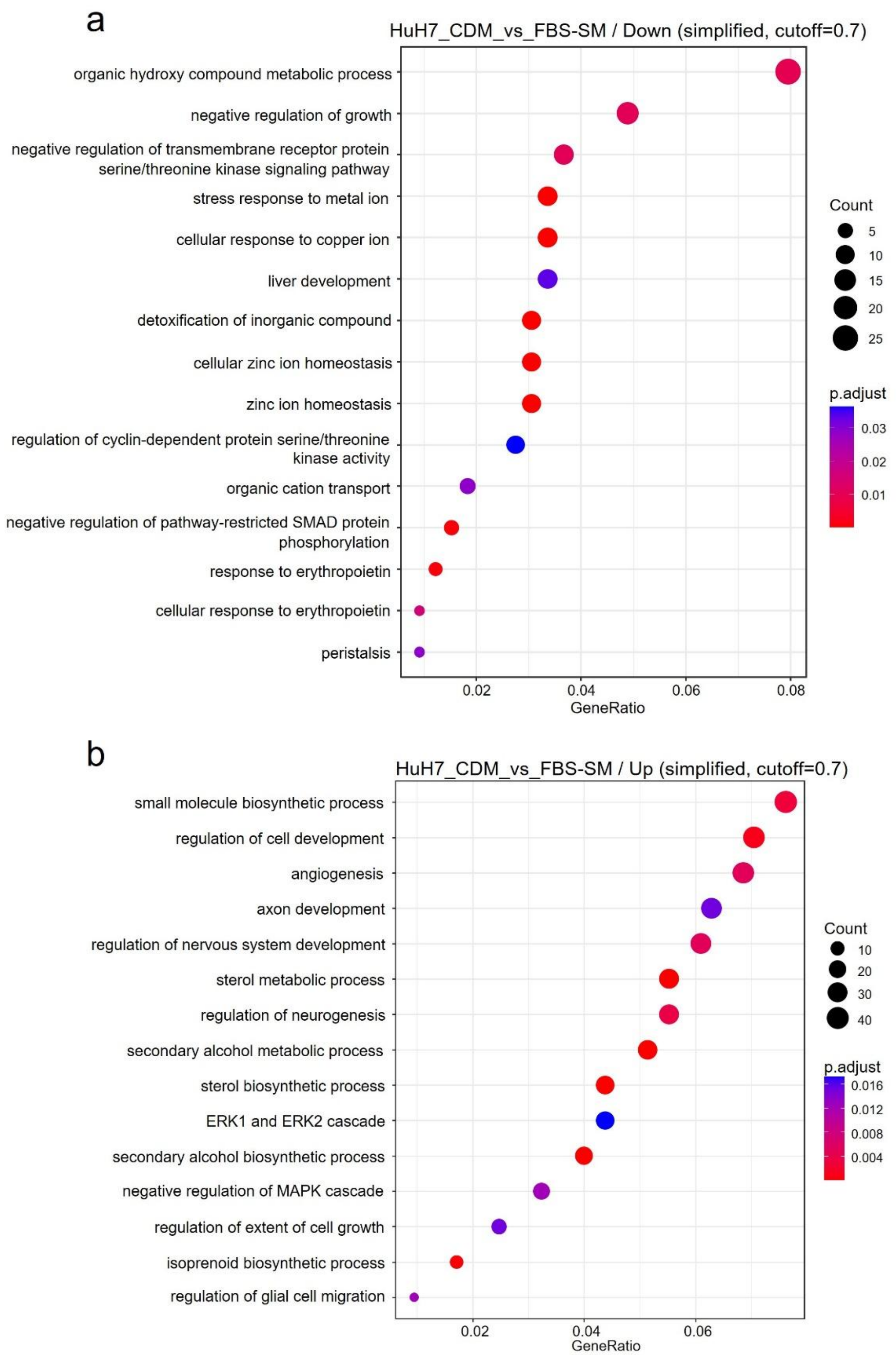


***Figure S6***. *Direction-resolved GO:BP enrichment for HuH7 CDM versus FBS-SM. Simplified GO:BP enrichment dotplots for HuH7 genes (a) upregulated and (b) downregulated in CDM versus FBS-SM. Dot size denotes GeneRatio and dot color denotes adjusted P value.*

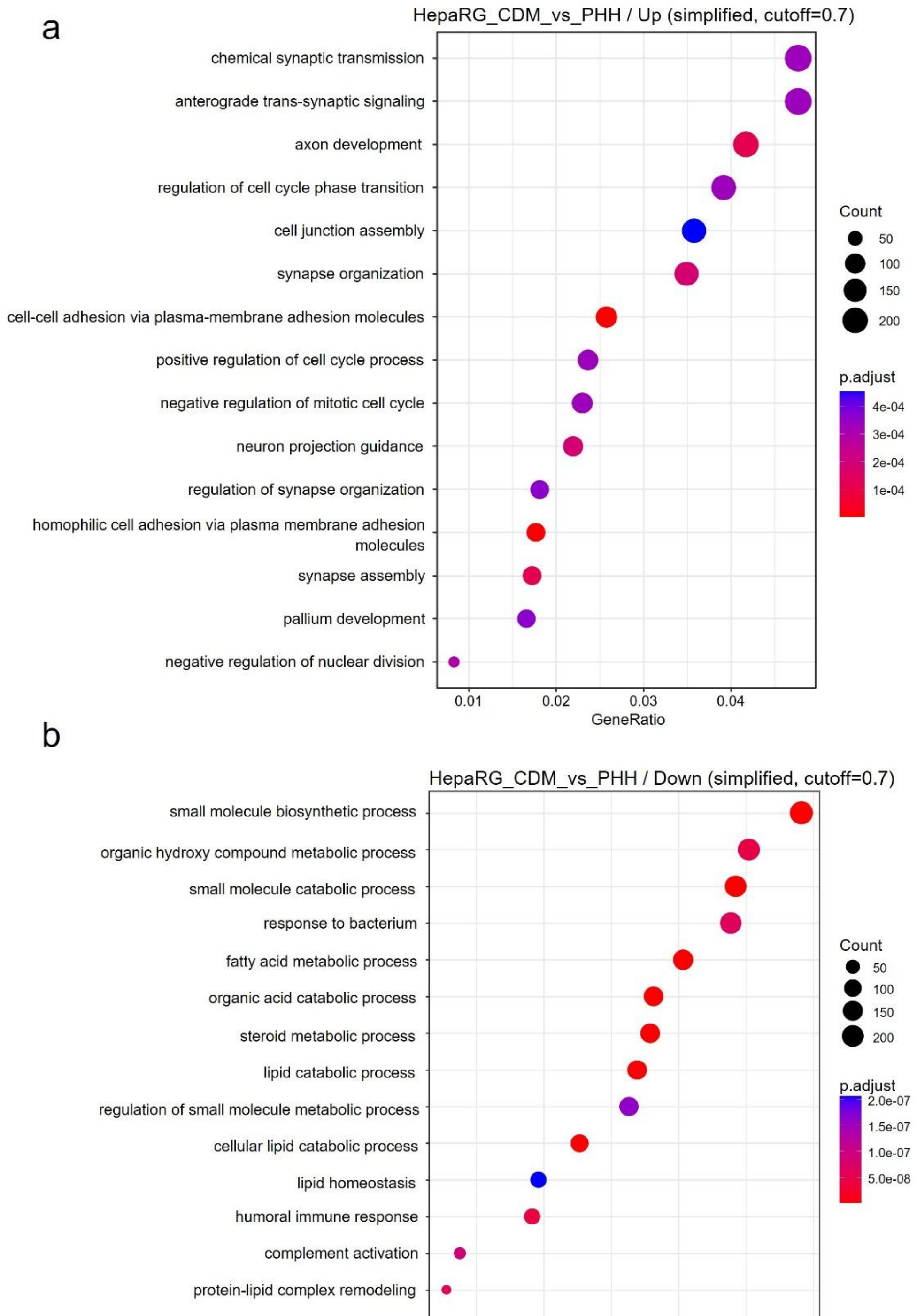


***Figure S7***. *Direction-resolved GO:BP enrichment for HepaRG in CDM benchmarking versus PHH. Simplified GO:BP enrichment dotplots for HepaRG differential expression (a) upregulated and (b) downregulated relative to PHH (direction-resolved).*

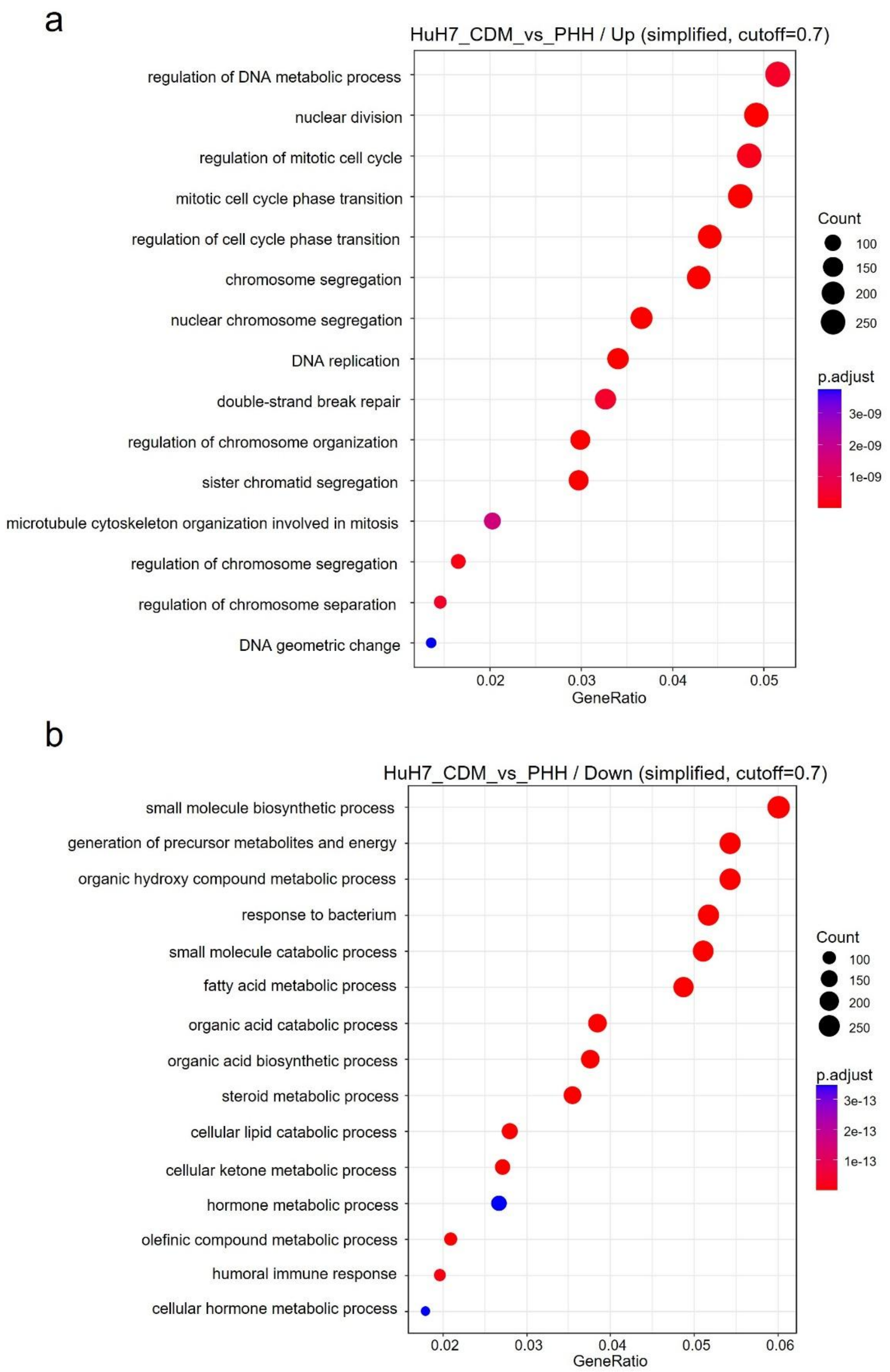


***Figure S8***. *Direction-resolved GO:BP enrichment for HuH7 in CDM benchmarking versus PHH. Simplified GO:BP enrichment dotplots for HuH7 differential expression (a) upregulated and (b) downregulated relative to PHH (direction-resolved).*

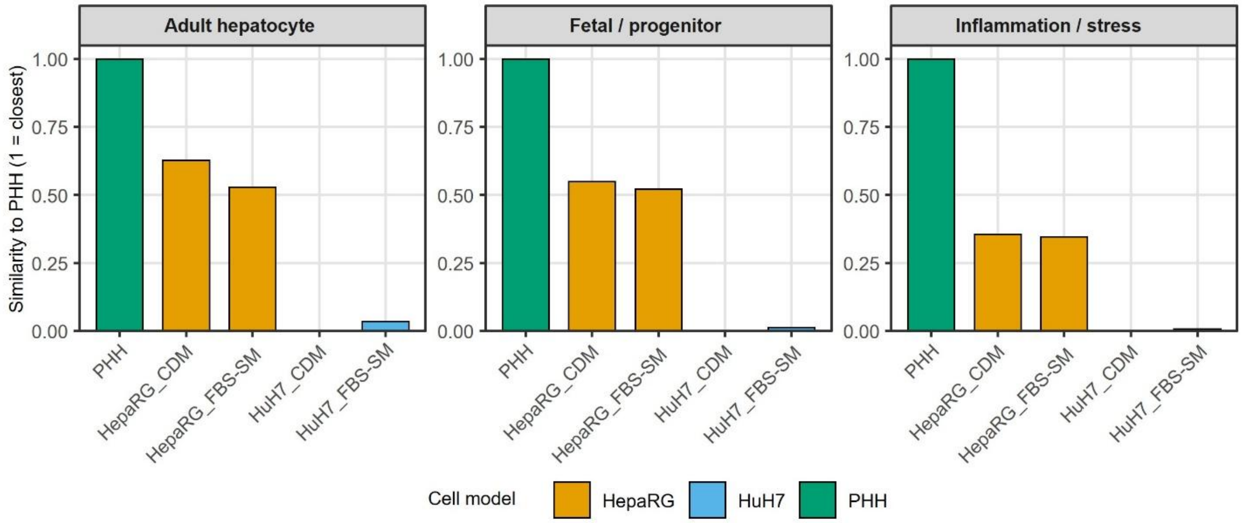


***Figure S9**. PHH-referenced similarity scores for hepatic identity marker categories. Quantitative similarity summaries benchmarking HepaRG and HuH7 (CDM and FBS-SM) to PHH across hepatic identity marker categories (adult hepatocyte, fetal/progenitor, inflammation/stress). Similarity scores were computed from gene-set expression deviations relative to PHH using an RMSE-based normalization (PHH fixed at 1).*

***Figure S10**. Full resolution ssGSEA pathway profiling benchmarked to PHH. High-resolution heatmap of ssGSEA scores (z-scored) computed from DESeq2 VST expression values for QuickGO-derived GO:BP term sets, **provided as a separate pdf file (Fig_S10_PHH_benchmark_pathways.pdf)**.*

1. Nessar, A. et al. *Promoting ethical and reproducible cell culture: implementing animal-free alternatives to teaching in molecular and cell biology.* **Volume 7 - 2025**, (2025).
